# A single-cell transcriptome atlas of the barley root apical meristem uncovers conserved and divergent roles of HvWOX5

**DOI:** 10.64898/2026.06.15.732315

**Authors:** Anika Dolata, Jasper Staut, Edgar Demesa-Arévalo, Tianyu Lan, Gabriele Buchmann, Pavel Solansky, Lea S. Berg, Michael T. Raissig, Marja Timmermans, Maria von Korff, Rüdiger Simon, Klaas Vandepoele, Yvonne Stahl

**Affiliations:** Institute for Developmental Genetics, Heinrich-Heine University, Universitätsstr. 1, D-40225 Düsseldorf, Germany; Faculty of Biosciences, Institute for Molecular Biosciences, Goethe University, D-60438 Frankfurt am Main, Germany; Ghent University, Department of Plant Biotechnology and Bioinformatics, Technologiepark 71, 9052 Ghent, Belgium; VIB-UGent Center for Plant Systems Biology, Technologiepark 71, 9052 Ghent, Belgium; VIB Center for AI & Computational Biology, VIB, Ghent, Belgium; Genetic Engineering Department, Irapuato Unit, Center for Research and Advanced Studies of the IPN (Cinvestav), Irapuato, Mexico; Institute of Plant Genetics, Heinrich-Heine University, Universitätsstr. 1, D-40225 Düsseldorf, Germany; Center for Plant Molecular Biology, Eberhard Karls University Tübingen, Tübingen 72076, Germany; Institute of Plant Sciences, University of Bern, 3013 Bern, Switzerland; Cluster of Excellence on Plant Sciences, “SMART Plants for Tomorrow’s Needs”, Heinrich-Heine-Universität Düsseldorf, 40225 Düsseldorf, Germany

**Keywords:** single-cell RNA sequencing, barley, root apical meristem, stem cell niche, quiescent centre, HvWOX5

## Abstract

Single-cell approaches have transformed plant developmental biology; however, cell-type-resolved resources for cereals remain limited. Here, we present a single-cell transcriptome atlas of the barley root apical meristem (RAM), which resolves 24 transcriptionally distinct cell populations. We assigned major root cell identities by integrating marker gene validation using Hybridization Chain Reaction (HCR) RNA fluorescence *in situ* hybridization and spatial transcriptomics with cross-species comparisons of published root atlases. Pseudotime analysis reconstructed developmental trajectories from the quiescent center to differentiating tissues, supporting the spatial and developmental organization of the atlas. We further demonstrated the utility of this resource by identifying *HvWOX5* expression in the quiescent center and metaxylem and showing that HvWOX5 loss-of-function mutants displayed reduced root and meristem lengths, altered stem cell niche homeostasis, and disrupted metaxylem organization. Taken together, this atlas provides a framework for dissecting barley root development and identifies HvWOX5 as a key regulator of RAM organization and metaxylem patterning.

## Introduction

The activity of the root apical meristem (RAM) of plants provides all cells necessary for building the entire root system. In the model plant *Arabidopsis thaliana* (Arabidopsis), RAM formation originates from a group of on average four to eight pluripotent stem cells, named the quiescent center (QC) (Baum et al. 2002; Dolan et al. 1993; Lu et al. 2021), which maintain the surrounding stem cells (also known as initials) in an undifferentiated state. These initials divide to generate transit-amplifying cells, which differentiate into distinct root tissues after several rounds of divisions. This sustains root growth and organ patterning throughout the plant’s whole life cycle. The resulting clonal lineages are organized in concentric layers, establishing a spatiotemporal gradient of cell differentiation along the longitudinal root axis, with the youngest cells forming at the QC position. This results in the formation of functionally and morphologically distinct root tissues that differ in their transcriptomes. These tissues are from the inside to the outside of the root: stele, endodermis, cortex and epidermis above the QC and the columella cells and lateral root cap below the QC (Dolan et al. 1993; van den Berg et al. 1997; Benfey and Scheres 2000).

Stem cell activity and cell fate specification of the root stem cell niche (SCN), which contains the QC and initials, are coordinated by a variety of hormonal cues, reactive oxygen species (ROS), and key developmental regulators, such as the conserved transcription factors (TFs) WUSCHEL-RELATED HOMEOBOX 5 (WOX5), important for QC and columella stem cell fate, the APETALA2-type TFs of the PLETHORA (PLT) family, master regulators of root development and important for maintaining the SCN, as well as the GRAS-domain TFs SHORTROOT (SHR) and SCARECROW (SCR), important for endodermis and cortex development (Drisch and Stahl 2015; Strotmann and Stahl 2021).

The organization of the RAM is conserved across monocots and dicots; however, there are remarkable differences in cell size and number. The *Hordeum vulgare* (barley) root shows a larger meristem, with an average of 30 QC cells and more cortex layers, forming the inner and outer cortex. These layers do not originate from a shared initial cell but are formed by the inner cortex/endodermis initial (CEI) and outer cortex initial (OCI) (Kirschner et al. 2017). Unlike the predominantly asymmetric formative divisions described in Arabidopsis, the enlarged barley QC is possibly maintained by symmetric and asymmetric cell divisions. This enlarged meristematic region suggests that the barley RAM has evolved an expanded and modular regulatory architecture that likely enhances its developmental robustness and plasticity. This may be of particular relevance for stress adaptation under changing environmental conditions, as plants are sessile organisms that must respond rapidly to external stimuli (García-Gómez et al. 2020). Barley, with its high tolerance to salinity and drought, is an interesting model for studying root development, especially with respect to climate change. However, the underlying gene regulatory network (GRN) of barley RAM maintenance remains elusive (Reviewed in Dresselhaus et al. 2025).

SCN regulation by phytohormones, such as auxin, has been described previously in Arabidopsis, where polar cell-to-cell auxin transport is mediated by the PIN gene family (Adamowski and Friml 2015; Blilou et al. 2005). In barley, a homologue of PIN1 has been shown to form a maximum concentration in the SCN, which maintains the stem cell pool in the RAM (Kirschner et al. 2018). In Arabidopsis, WOX5, a central regulator of the root SCN, is mainly expressed in the QC, where it maintains the surrounding initials in an undifferentiated state via non-cell-autonomous signaling. It also limits premature QC divisions by repressing cell cycle gene activity (Sarkar et al. 2007; Burkart et al. 2022). The barley WOX5 homologue was identified to be closest to the rice QUIESCENT-CENTER-SPECIFIC HOMEOBOX (QHB), which is expressed in the QC as well as in the metaxylem, suggesting a broader role for root maintenance due to its expanded expression domain (Chu et al. 2013; Kirschner et al. 2017). Mutation of QHB affects lateral root formation, resulting in fewer short lateral roots (S-type) and more long lateral roots (L-type) (Kawai et al. 2022). This indicates a general role of WOX5 in root formation also in crops; however, its function in main root growth remains largely elusive.

Members of the PLT gene family are essential for Arabidopsis root development. Their expression forms a gradient throughout the RAM, peaking in the SCN, which is mediated by auxin and local TF mobility. PLTs coordinate stem cell fate and tissue patterning in the root, possibly by forming complexes with other TFs (Aida et al. 2004; Galinha et al. 2007). Recently, it has been shown in Arabidopsis that PLT3 can directly interact with WOX5, regulating each other’s expression in a negative feedback loop (Burkart et al. 2022). This suggests complex regulatory and cell type-specific interactions to maintain SCN homeostasis. The cortex and endodermis cell fates are determined by the GRAS TFs, namely SHORTROOT (SHR) and SCARECROW (SCR). Here, SHR moves from the stele to the endodermis, where it interacts with SCR, which in turn prevents further SHR movement. This interaction is required for the regulation of the asymmetric division of the cortex/endodermis initials, which establishes the endodermis cell fate, as well as for QC specification and maintenance (Lim et al. 2000; Nakajima et al. 2001; Gallagher et al. 2004; Gallagher and Benfey 2009).

In recent years, single-cell RNA sequencing (scRNAseq) has emerged as a powerful technique for uncovering GRNs in various plant species. Root transcriptome atlases have been generated in Arabidopsis, as well as in cereals such as maize, rice, and wheat. This allowed the identification of root regulators in an unbiased manner and provided insights into RAM regulation at the molecular level (Zhang et al. 2019; Zhang et al. 2021; Wang et al. 2021; Serrano-Ron et al. 2021; Liu et al. 2021b; Ke et al. 2025; Denyer et al. 2019).

In this study, we developed a single-cell transcriptome atlas of the barley RAM. We identified 24 distinct clusters and their respective marker genes. We performed pseudotime analyses, uncovering a trajectory and distinct upregulated genes for various time points of cell type development. For final cluster annotation, we used cross-species annotations. This atlas allows the identification of cell type-specific genes, such as HvWOX5. We generated a loss-of-function mutant of barley WOX5 and observed phenotypic changes that partially resembled the previously described Arabidopsis phenotype but also shows novel phenotypes that affect the metaxylem.

## Results

### A single-cell transcriptome atlas of the barley RAM

To generate a single-cell atlas of the barley RAM, we enzymatically digested the first 2 mm of barley Golden Promise root tips six days after germination (6 DAG) to release protoplasts (Figure 1A-C). Protoplast viability was assessed using Calcein and DRAQ7 staining, and only protoplast samples with more than 70% viable cells were processed for sequencing. Subsequently, scRNAseq was performed using the microwell-based BD Rhapsody platform (Figure 1D) as previously described (Demesa-Arevalo et al. 2026). Across three biological replicates, we recovered approximately 5,000 cells, with an average of 4,851 genes and 24,854 transcripts per cell.

**Figure 1.**
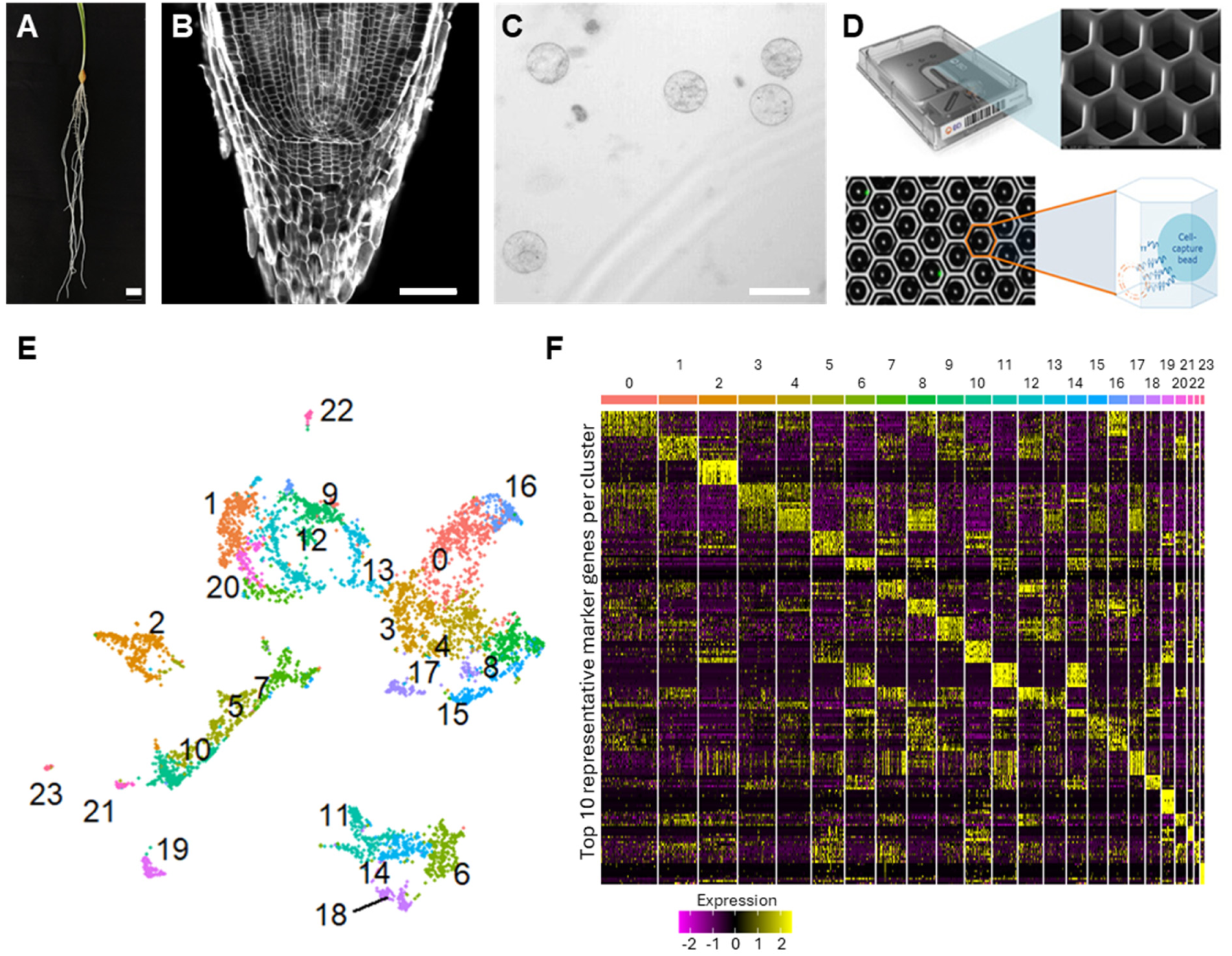
Workflow of the scRNA-seq pipeline of the barley RAM and cluster annotation. **A)** Barley roots were grown for six days. **(B)** Barley RAMs were harvested and **(C)** protoplasted, including viability testing. **D)** scRNA-seq was performed on the BD Rhapsody platform using cartridges with microwells and magnetic beads to capture single cells (modified from https://mdxk.co.kr/m21_06.php). **E)** Uniform manifold approximation and projection (UMAP) dimension reduction of approximately 5,000 single cells of the barley Golden Promise RAM showed 24 populations. **F)** Expression of the top 10 marker genes per cluster based on log2 fold change. Scalebars represent in **A)** 5 mm, in **B)** 100 µm, and in **C)** 50 µm.

To minimize possible protoplasting effects, we performed bulk RNA sequencing of protoplasted and intact 6 DAG barley RAMs in three biological replicates. Here, we identified 1,308 differentially expressed genes (DEGs) that were upregulated in response to protoplasting (Suppl. Table 1, Suppl. Figure 1A, A’), which were removed from the three independent scRNAseq data sets before downstream analyses. To reduce batch effects, we integrated the three replicates using the 7,500 most variable genes (Suppl. Figure 1B,C). Principal component analysis (PCA) and clustering with subsequent uniform manifold approximation and projection (UMAP) dimension reduction, identified and visualized 24 distinct populations containing between 32 and 494 cells, and their respective top10 cluster-specific marker genes (Figure 1E,F). Cluster-specific markers were selected for high specificity within the cluster of interest (high pct.1) and low presence in all other clusters (low pct.2; Suppl. Table 2) and were prioritized for subsequent validation by HCR RNA fluorescence *in situ* hybridization and spatial transcriptomics (Molecular Cartography).

### scRNAseq data analysis reveals specific marker genes for tissues in the RAM

We first assigned cluster identities using spatially resolved marker gene validation. HCR RNA *in situ* hybridization of *7HG0662370*, encoding an LRR-XI protein kinase, marked columella stem cells (CSCs) and parts of the vasculature (metaxylem, phloem) and supported the annotation of cluster 21 as CSCs (Figure 2A) together with expression of the ortholog of Arabidopsis *WOUND-INDUCED POLYPEPTIDE 5* (*WIP5*) (Suppl. Figure 2O). The putative protease inhibitor *1HG0004290* specifically labelled the outer cortex (Figure 2B), and its absence from the inner cortex supports the previously described separation of the inner and outer cortex lineages in barley (Kirschner et al. 2017); further cortex identities were assigned by molecular cartography data for *HvMND8* (Suppl. Figure 2A) and HCR *in situ* hybridization of *2HG0206040* (Suppl. Figure 2B). The *SCR* homologue *2HG0133890* (*HvSCR*) was exclusively detected in the endodermis and few QC cells by HCR RNA *in situ* hybridization (Figure 2C), consistent with previous *in situ* hybridization data (Kirschner 2017), supporting the annotation of clusters 1 and 20 as endodermis. Vascular identities were supported by the expression of *7HG0666250*, encoding a TCP transcription factor, *5HG0508470*, encoding a homeobox leucine-zipper protein (Figure 2D,E), *HvRA2*, *7HG0666250* (Suppl. Figure 2C,D), *HvMND1* (Suppl. Figure 2G) and the barley *SHR* homologue (Suppl. Figure 2I), which together annotated clusters 5, 7, and 10. Finally, spatial transcriptomics detected *3HG0301330* (*HvWOX5*) in metaxylem and, to a lesser extent, in the QC (Figure 2F), resembling the expression pattern of the rice QHB (Chu et al. 2013). *HvWOX5* was enriched in clusters 2 and 10 in the scRNAseq data. Because cluster 10 was independently supported as vascular/metaxylem tissue, we assigned the QC/niche identity to cluster 2. This is further supported by the published expression domains in the QC/niche of *HvSCR* (Figure 2C) and *HvSHR* (Suppl. Figure 2I) (Kirschner 2017), as well as orthologs of the Arabidopsis *JACKDAW* (*JKD*) (Suppl. Figure 2J) and *BRASSINOSTEROIDS AT VASCULAR AND ORGANIZING CENTER* (*BRAVO*) (Suppl. Figure 2L). We assigned xylem and phloem identity to the clusters 23 and 19 respectively, based on expression of *5HG0462580* (Xylem) (Suppl. Figure 2E) and the orthologs of Arabidopsis ALTERED PHLOEM DEVELOPMENT (*APL*) (Suppl. Figure 2M) and *VASCULAR RELATED NAC-DOMAIN PROTEIN 1*, *6* and *7* (*VND1,6,7*) (Suppl. Figure 2H,K,N). Together, these validations identified QC/niche, CSC, vascular, cortex, endodermal, and cell cycle-associated populations. However, several clusters remained unidentified, and we found one specific group of cells that was highly enriched in genes involved in the cell cycle (Suppl. Figure 2F).

**Figure 2.**
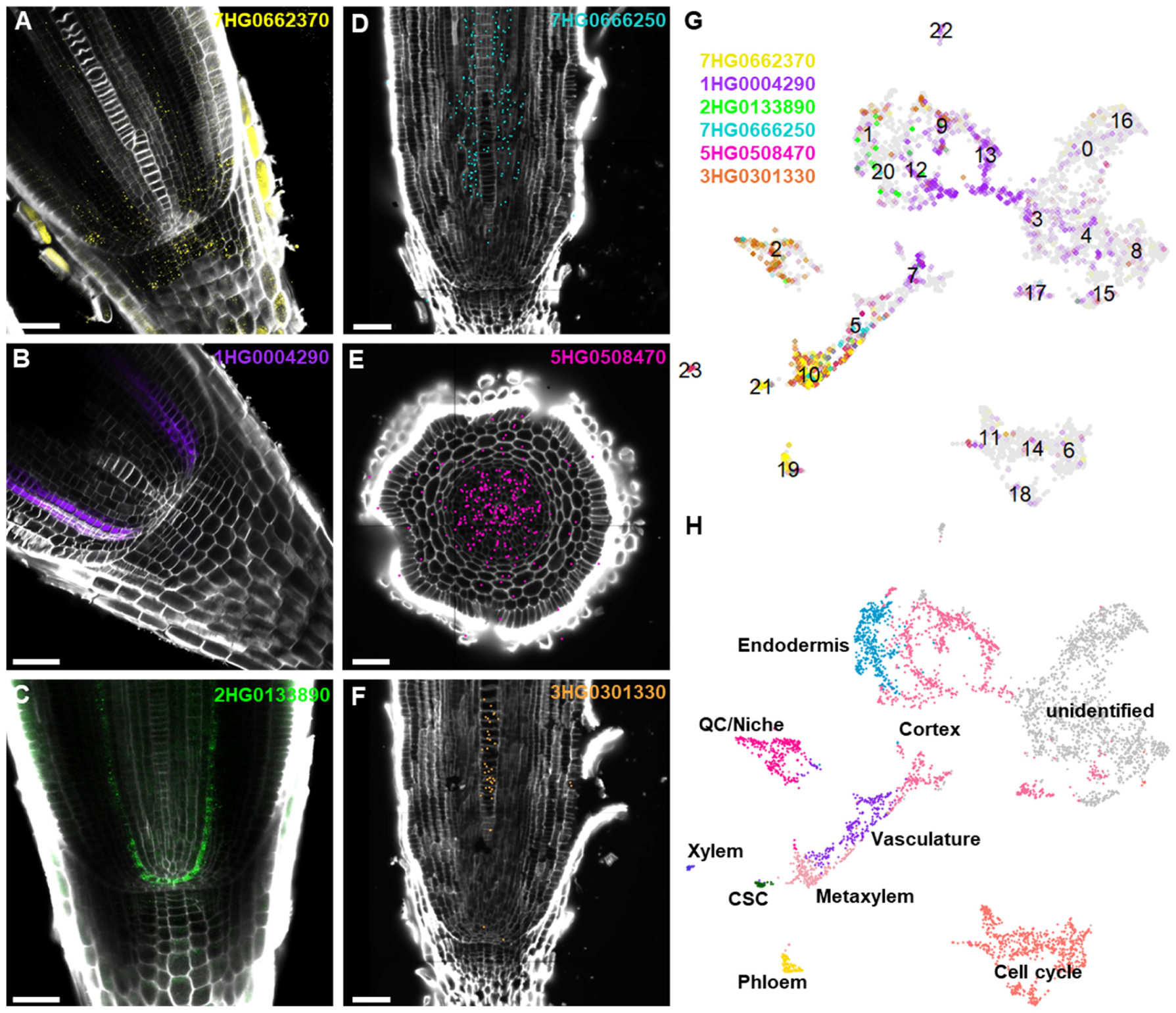
RNA *in situ* hybridization of marker genes for cluster annotation. Gene expression patterns in 6 DAG barley RAMs of the following genes by RNA-FISH data **A)** *7HG0662370*, **B)** *1HG0004290*, **C)** *2HG0133890* (*HvSCR*), or Molecular Cartography gene expression patterns of **D)** *7HG0666250*, **E)** *5HG0508470*, **and F)** *3HG0301330* (*HvWOX5*). Cell wall staining with SR2200 dye (white). Scalebars = 50 µm. **G)** Expression of **A-F)** in scRNAseq clusters. **H)** Cluster annotation based on marker gene expression.

### Cross-species comparisons allow identification of the remaining unidentified clusters

To annotate the missing clusters and to bring our data into a larger context, we performed cross-species comparisons between published data from Arabidopsis, maize, rice, tomato, and wheat (Wendrich et al. 2020; Ortiz-Ramírez et al. 2021; Liu et al. 2021a; Cantó-Pastor et al. 2024; Ke et al. 2025) (Figure 3A) with our barley RAM dataset (Figure 3B). For this, we inferred cluster-specific DEGs for Arabidopsis, maize, rice, tomato, and wheat. These DEG sets were then used to apply a cross-species annotation pipeline between barley and all other species as previously described in (Ke et al. 2025). We annotated the barley clusters as endodermis, cortex, LRC, epidermis, initials, stele, xylem, and phloem, covering almost all the expected cell types of the barley RAM. Comparisons of the cross-species annotations and the annotations derived from HCR RNA *in situ* hybridizations revealed some overlap; however, we also observed some variations between the predicted annotation and our marker-gene-based approach. We determined the quality of our cross-species comparison data by the number of species-specific annotations that agreed, ranging from very strong agreement (five out of five) to no agreement (two or fewer) (Suppl. Table 3). In cases where these comparisons show only weak agreement and contradict our *in situ* gene expression observations, we give priority to annotations based on our experimental verification. By combining data from our marker gene approach and cross-species comparisons, we were able to annotate all clusters of our scRNAseq dataset, except for cluster 22 (Figure 3C). By employing this method, we could identify all expected main cell types; however, the initial stem cells, which were annotated in the Arabidopsis and maize datasets, were only present in cluster 11, which we annotated as cell cycle (initials) (Figure 3C). For easy access to the generated data and its subsequent analyses as described above, we built an annotated web browser-based barley RAM transcriptome atlas available at https://www.zmbp-resources.uni-tuebingen.de/timmermans/barley_browser/.

**Figure 3.**
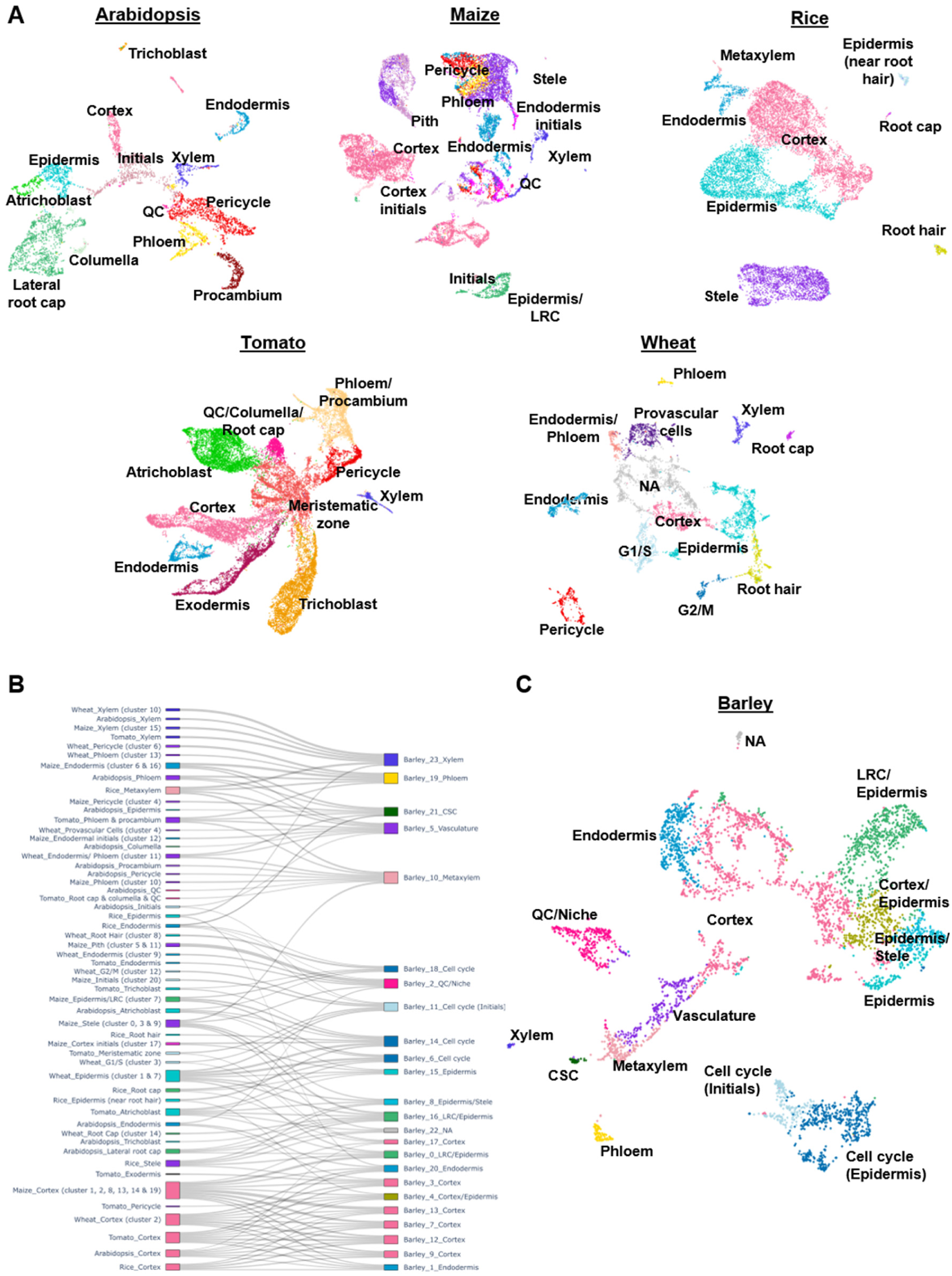
Cross-species comparison. **A)** Single-cell transcriptomes from Arabidopsis, maize, rice, tomato, and wheat were used for the final cluster annotation of the barley scRNAseq dataset. The transcriptome data derived from Wendrich et al., 2020, Cantó-Pastor et al., 2024, Ortiz-Ramírez et al., 2021, Liu et al., 2021 and Ke et al., 2025. **B)** Sankey-plot visualizing the clusters considered for barley RAM scRNAseq annotation. **C)** Final annotation of the barley RAM scRNAseq dataset.

### Pseudotime analysis for the detection of developmental trajectories

During scRNAseq, spatial and spatiotemporal information is lost. However, temporal dynamics can be inferred using pseudotime analysis, especially if knowledge of developmental transitions is available. In the RAM, it has been shown that all tissues originate from the QC, as described in Arabidopsis, where cells with meristematic identity appear earlier in pseudotime than differentiated cells (Denyer et al. 2019). We performed pseudotime analysis with monocle3 using a subset of our dataset, focusing on the QC/niche and vascular tissues, with the QC/niche cluster selected as a starting point (Figure 4A). According to our pseudotime analyses, starting from the QC/niche (cluster 2), the metaxylem develops (cluster 10), which is reasonable given its close position in relation to the QC. Vascular tissues (cluster 5) are slightly older in pseudotime and subsequently differentiate into xylem (cluster 23) and phloem (cluster 19). Plotting the expression of marker genes onto the pseudotime trajectories validated the proposed temporal dynamics, with the QC/niche being established first (Figure 4B), followed by the general vasculature (Figure 4B’), which later forms xylem (Figure 4B’’) and phloem (Figure 4B’’’). This pseudotime analysis of the barley RAM recapitulates developmental trajectories and highlights conserved developmental strategies across the roots of different plant species. As HvWOX5 appears early in development, peaking in the QC/niche and metaxylem cluster, we assumed an important role in RAM development and maintenance. To validate this proposed function, which has already been described in Arabidopsis (Sarkar et al. 2007; Burkart et al. 2022), we generated a *HvWOX5* loss-of-function mutant using CRISPR-Cas9.

**Figure 4.**
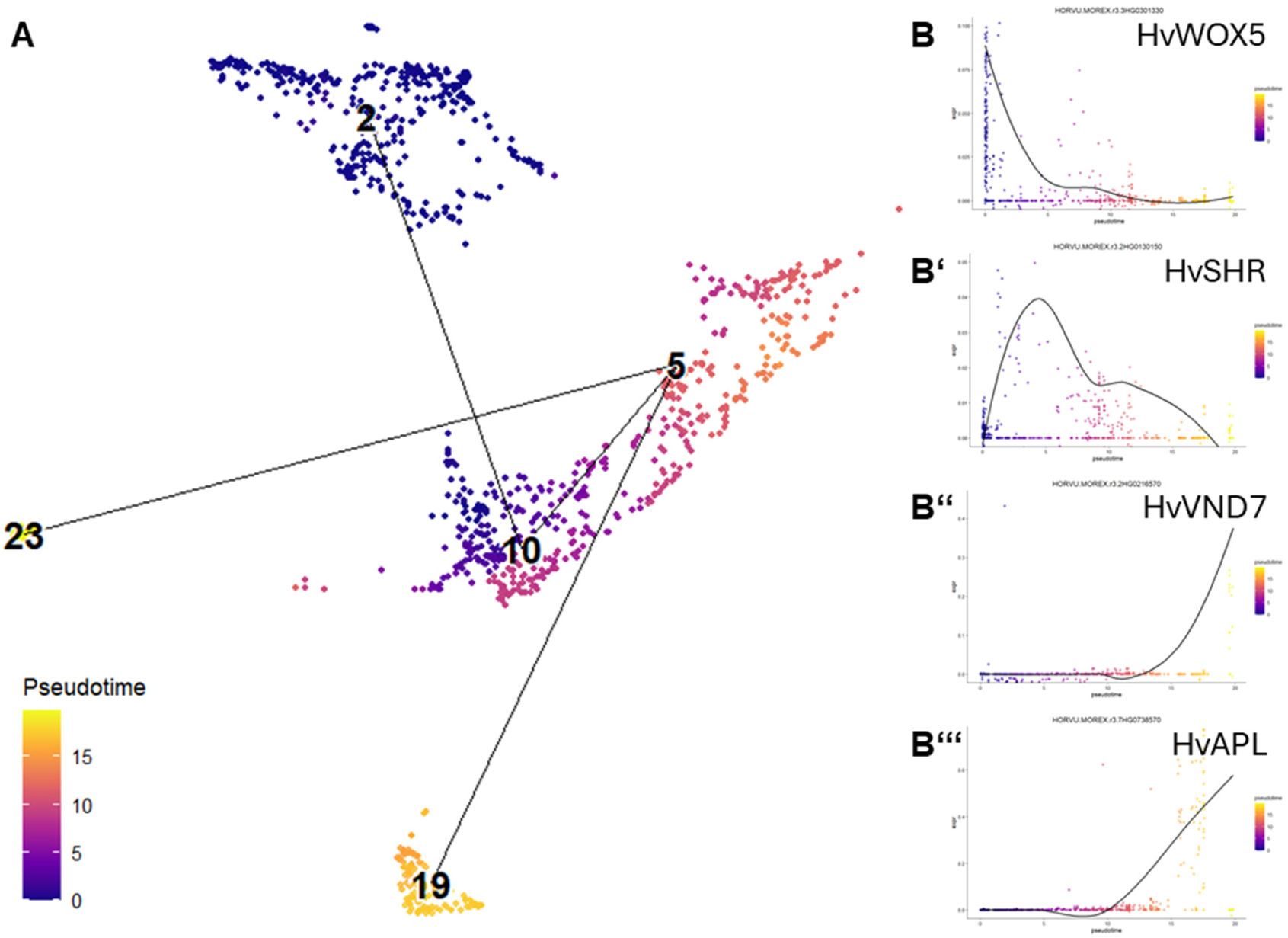
Pseudotime analysis to detect developmental trajectories. **A)** Pseudotime trajectories on a subset or the barley RAM scRNAseq dataset including QC/niche and vascular tissues. **B-B’’’)** Marker gene expression during pseudotime **for B)** *3HG0301330* (*HvWOX5*), **B’)** *2HG0130150* (*HvSHR*), **B’’)** *2HG0216570* (*HvVND7*) and **B’’’)** *7HG0738570* (*HvAPL*).

### HvWOX5 plays an essential role in barley RAM development and SCN homeostasis

The *HvWOX5* loss-of-function allele contains a four base pair deletion in the first exon, leading to an early stop codon after the first three amino acids (Suppl. Figure 3). First, we analyzed the root lengths of barley seedlings at 6 DAG. The *Hvwox5-*mutant shows a significant decrease in root length compared to wildtype Golden Promise Fast (GPF) (p = 1.08 x 10^−13^) (Figure 5A,B). As this phenotype may be caused by multiple factors, we quantified the length of the meristematic zone by measuring the distance between the QC region and the first cortex cell that doubled in size, which marks the end of the meristematic zone (Kirschner et al. 2017) (Figure 5C-E). Interestingly, *Hvwox5*-mutants show a significant decrease in meristematic zone length compared to the wildtype (p = 0.001251). To determine whether this shortened meristematic zone was caused by a defect in the QC region, we adjusted the previously described SCN-staining method (Burkart et al. 2022) for barley root sections (Figure 5F-G’’), allowing for the simultaneous quantification of QC divisions and CSC differentiation. Here, 5-ethynyl-2’-deoxyuridine (EdU) is incorporated into DNA during the S-phase, thereby marking cells that are about to undergo division or have just divided. The CSC layers can be determined by the absence of starch granules, which only appear in differentiated CCs. After 6 h of EdU treatment, the wildtype GPF showed, on average, two CSC layers and no QC divisions (Figure 5F’’). In *Hvwox5*-mutants, an increase in QC divisions was visible, accompanied by fewer CSC layers (Figure 5G’’), indicating a function of HvWOX5 in both QC maintenance and CSC differentiation. Additionally, we observed variations in metaxylem organization within the RAM of *Hvwox5-*mutants, which we classified into three categories: i) normal, one row of metaxylem cells; ii) division, an additional division of the metaxylem; and iii) double, additional divisions leading to a double metaxylem (Figure 5H-K). We quantified these metaxylem phenotypes by counting the roots that showed the respective phenotypes, leading to approximately 55% of wildtype GPF roots showing normal metaxylem and approximately 45% having an additional division. The *Hvwox5*-mutants display two rows of metaxylem in 50% of the roots. Approximately 40% of the roots showed additional divisions, and only 10% resembled a wild-type phenotype.

**Figure 5.**
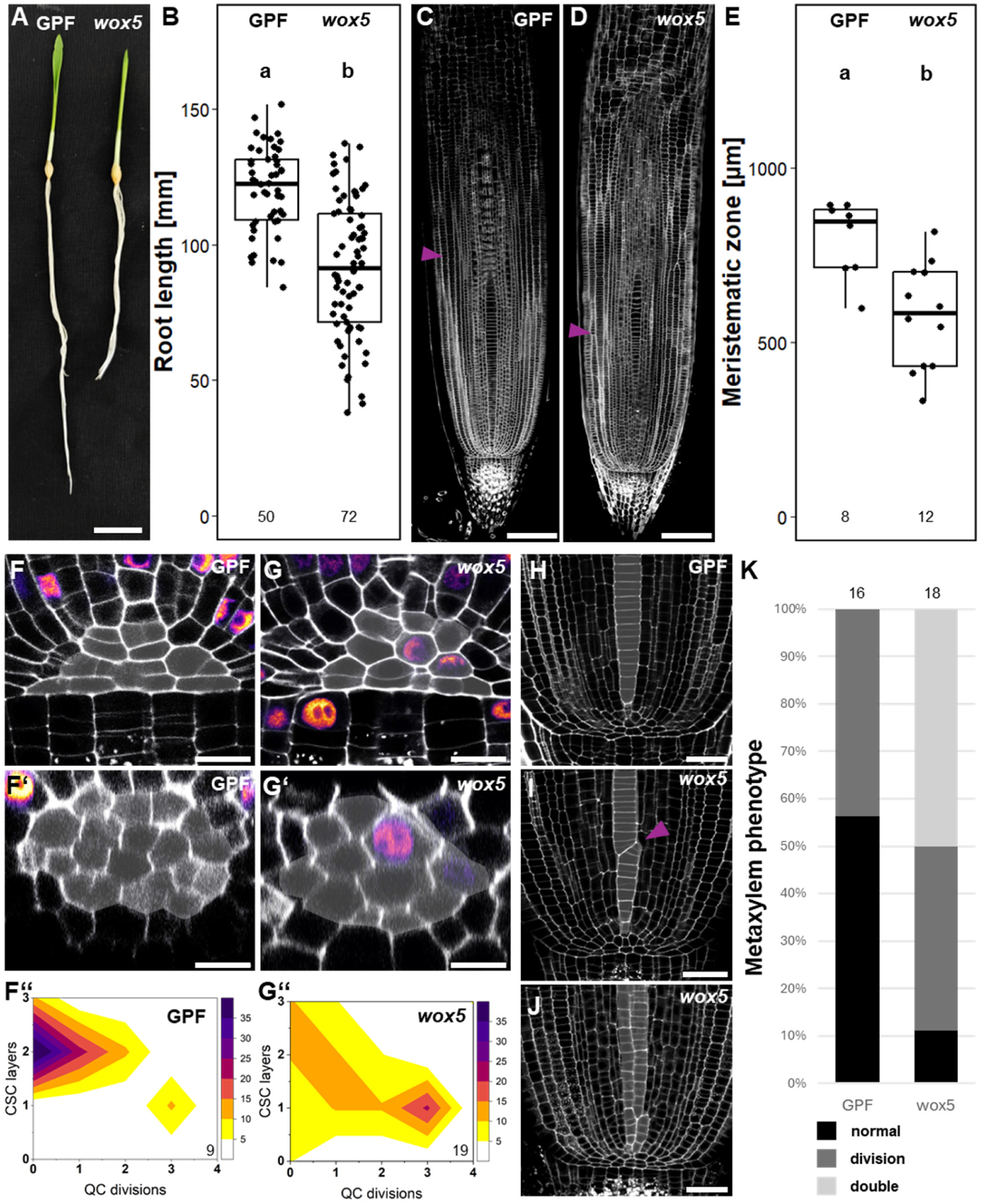
WOX5 function in the barley root. A, **B)** Root length measurements of barley seedlings 6 DAG, **A)** representative images of Golden Promise Fast (GPF) and *Hvwox5*. **B)** Quantification of root lengths of GPF and *Hvwox5*-seedlings, number of replicated is indicated below the boxplots. **C-E)** Measurement of meristematic zone length. Representative images of GPF **(C)** and *Hvwox5* **(D)** roots are shown. Purple arrows indicate the end of the meristematic zone. **E)** Quantification of meristematic zone length and number of replicates is indicated below the boxplots. **F-G’’)** EdU/mPSPI staining of GPF **(F-F’’)** and *Hvwox5* **(G-G’’)** roots. **F)** and **G)** show longitudinal sections, **F’)** and **G’)** optical cross-sections. **F’’)** and **G’’)** visualize QC divisions and CSC layers in 2D-plots. **H-K)** Quantification of metaxylem phenotypes, **H-J** show representative images of the respective phenotypes: **H)** normal metaxylem, **I)** division, **J)** double metaxylem. Division in **I)** is indicated by a purple arrow. **K)** Quantification of metaxylem phenotypes. The number of replicates is indicated above each bar plot. **L)** Quantification of root width; the number of replicates is indicated below the boxplots. Scale bars represent **A)** 2 mm, **C, D)** 200 µm, **F-G’)**, **H-J)** 20 µm.

Furthermore, we measured root width across the QC region. Interestingly, we observed significantly thinner roots in the *Hvwox5*-mutant compared to the wildtype (Suppl. Figure 4A,B). Additionally, we quantified the number of cell layers above and below the QC, demonstrating a significant decrease in the number of cell layers in the proximal part of the *Hvwox5-*mutant roots, but not in the distal part (Suppl. Figure 4A,C,D).

Taken together, these findings highlight the usability of this first barley RAM scRNAseq atlas to predict cell-type specific functional relevance of genes, as demonstrated here for HvWOX5.

## Discussion

scRNA-seq is a powerful tool for the unbiased characterization of transcriptional variations and similarities in cell populations in diverse biological systems. Given that clusters are mathematical groupings of cells based on similarities in their transcriptomes, their biological relevance is assigned by downstream analysis and cluster annotation. This is straightforward in well-studied model organisms such as Arabidopsis, where many validated markers are available. In less well-studied organisms such as barley, however, this remains more challenging.

Here, we established a transcriptome atlas of barley RAMs from three biological replicates and approximately 5,000 cells. We were able to identify 24 distinct clusters with their respective specific marker genes. Possible batch effects could be avoided by performing integration, after which all three replicates clustered in a similar pattern. Upregulated genes caused by protoplasting could be removed from our dataset by performing bulk RNA sequencing of protoplasts and undigested tissue; however, we may have overlooked some potentially relevant candidate genes as their expression might be affected by protoplasting. Additionally, the technology has a bias for smaller cells, as the well plate for BD Rhapsody cannot fit cells larger than 60 µm. Therefore, we may have lost large cells from the outer root cap layers. Nevertheless, the procedure is sufficient to capture all major cell types, including the pluripotent stem cells of the QC, indicating that approximately 5,000 cells are sufficient to cover the whole barley RAM. The expression of known marker genes for root tissues in our scRNAseq dataset helped to annotate cluster identities, combined with an unbiased approach selecting DEGs for each cluster and performing HCR RNA fluorescence *in situ* hybridization and spatial transcriptomics.

The specific expression of genes such as HvSCR, in line with its QC and endodermis expression pattern known from previously published *in situ* hybridizations (Kirschner 2017), suggests that the HCR RNA fluorescence *in situ* hybridization technique employed here achieves a high level of spatial resolution and specificity, highlighting the utility of this approach. The subsequent cluster identification supplemented by unbiased cross-species comparisons indicated that there is no need for *a priori* knowledge of specific marker genes for assigning cluster identities, allowing the identification of unique developmental regulators and downstream genes that give cells their distinct identities, resulting in different forms and functions. Furthermore, we identified the putative protease inhibitor *1HG0004290* as a novel marker for the outer cortex layers, showing distinct expression in this tissue, which makes it an interesting candidate for mutant generation and further functional studies.

We performed cross-species comparisons between our dataset and previously published data from Arabidopsis, maize, rice, tomato, and wheat to verify and further annotate our yet unidentified clusters and to place our data in a larger context. Notably, not all cell types that were expected were found in our annotation, such as the QC/niche, which we identified using our candidate gene approach. It has been previously reported that finding the very few QC cells, especially in Arabidopsis, can be difficult (Denyer et al. 2019). The absence of this cell type from some of the reference datasets may lead to misannotations of the barley scRNAseq dataset. Further difficulties can be caused by differences in the quality of the datasets used, such as different sequencing depths, number of cells, or under- or over-representation of certain cell types due to the method bias for smaller cells. Furthermore, while the genome conversions performed for maize and rice to assign orthologous groups to DEGs resulted in one-to-one conversions for the vast majority of genes, many-to-one conversions were also observed, resulting in cases where multiple orthologous groups get assigned to one gene. Discarding such genes led to a loss of DEGs, which is expected to cause a lower sensitivity in the analysis for rice and maize. Thus, we decided to prioritize the annotations validated by HCR RNA *in situ* hybridization and use the cross-species annotations only for the unidentified clusters. As described by (Ke et al. 2025), the xylem cluster is the most identifiable and transcriptionally distinct cluster. Cross-species comparisons validated cluster 23 of our scRNAseq dataset as the xylem cluster.

Pseudotime analysis allows for a deeper understanding of scRNAseq data, as it can add biological context to clustering, which is solely based on similarities in transcriptomes. By selecting the QC/niche cluster as a starting point, we ensured the biological validity of our analysis, as its role as the origin of all root tissues has been previously described (Reviewed in Dresselhaus et al. 2025). With pseudotime, we observed a trajectory from the undifferentiated QC/niche to the respective differentiated cell types. Notably, the more differentiated vascular tissues xylem and phloem appear later in pseudotime than the metaxylem, highlighting differentiation processes in the RAM. Plotting of previously identified marker genes onto our pseudotime data again confirmed the biological relevance, as the QC/niche marker appeared earlier in pseudotime than the markers for differentiated tissues, such as the xylem and phloem.

We were able to identify HvWOX5 expression in our scRNAseq dataset within the QC/niche and metaxylem clusters, similar to its rice ortholog (Chu et al. 2013). We generated a *Hvwox5* loss-of-function mutant, which shows shorter roots and meristematic zones, which has not been reported in other crop species; here, we show a short-root phenotype in barley *Hvwox5*-mutants that is more severe than previously described in Arabidopsis (Savina et al. 2020). This difference in severity could be caused by the additional expression of HvWOX5 in the metaxylem compared to the restricted expression domain of Arabidopsis AtWOX5 in the QC.

Hence, we conclude that not only the QC but also the metaxylem could be responsible for the overall root length as a consequence of the reduced meristem size. In addition to the novel function in metaxylem patterning, we observed previously not reported effects on root width and number of cell layers in *Hvwox5*-mutants, indicating a more versatile role of HvWOX5 in the barley RAM.

In Arabidopsis, *Atwox5*-mutants show more QC divisions with fewer CSC layers as a compensatory mechanism for distal stem cell loss (Stahl et al. 2009; Burkart et al. 2022; Strotmann et al. 2025). AtWOX5 influences differentiation of CSC non-cell-autonomously (Sarkar et al. 2007) together by interacting with AtPLT3 (Burkart et al. 2022). In the barley *Hvwox5*-mutant, we also observed more QC divisions and fewer CSC layers, indicating that the mechanisms described in Arabidopsis are conserved. In future studies, the potential interaction between HvPLT3 and HvWOX5 could be investigated in more detail to assess the conservation of molecular regulation in the barley RAM.

In summary, our single-cell transcriptome atlas of the barley RAM provides a framework for dissecting root development and identifies HvWOX5 as a regulator of both SCN homeostasis and metaxylem organization. Hereby, we showed both conserved but, importantly, also novel functions of WOX5 in crop root development.

## Materials and Methods

### Plant materials and growth conditions

Wildtype tissue was collected from the *Hordeum vulgare* cultivar Golden Promise for protoplast isolation and Golden Promise Fast (Buchmann et al. 2026) for root imaging and mutant generation. Seeds were stratified for two days at 12°C in the dark in Petri dishes lined with wet paper. Afterwards, they were sown on paper rolls and grown under long day conditions (16 h light, 8 h dark) at 21°C for 6 days. Root tips were cut at ∼1-2 mm with a scalpel for protoplasting and at ∼3-4 mm for HCR RNA fluorescence *in situ* hybridization and EdU/mPSPI-staining. We collected approximately 150 root tips per scRNA-seq experiment, and the overall experiment was performed three times.

### Protoplast isolation

The protoplasting protocol was adapted from (Demesa-Arevalo et al. 2026). Root tips were collected in 6-well plates containing 3 mL washing solution I (WSI; 0.4 M mannitol, 20 mM MES, 20 mM KCl, 10 mM CaCl2, 0.1% BSA, pH 5.7). For cell wall digestion, roots were cut into small pieces in 4 mL enzyme solution (0.4 M mannitol, 1.2% Cellulase R-10, 1.2% Cellulase RS, 0.36% Pectolyase Y-23, 0.4% Macerozyme R-10 (all from Duchefa), 20 mM MES, 20 mM KCl, pH 5.7, 10 mM CaCl2, 0.1% BSA, 0.06% β-mercaptoethanol) and incubated for 2.5 h while shaking. After digestion, the protoplast solution was added to a 40 µm strainer placed on a 50 mL falcon conical tube for centrifugation (250 *g*, 3 min, 4°C). The pellet was washed once with WSI and twice with WSII (0.6 M mannitol, 2 mM MES, 0.1% BSA, pH 5.7) with centrifugation between each wash step, as previously described. Protoplast quality was evaluated by staining with calcein AM (10 μM) and DRAQ7 (1.5 μM) staining (5 min at room temperature). Cell viability was scored using a BD Rhapsody scanner, and only samples exceeding 70% viability were used for mRNA capture and sequencing.

### Single-cell RNA sequencing

Approximately 20,000 cells were loaded per replicate for each sample. We used the mRNA Whole Transcriptome Analysis (WTA) kit from BD-Biosciences for library preparation. Libraries were sequenced using the Illumina NextSeq2000 platform, targeting 50,000 reads per cell. Raw scRNAseq data were processed using the BD WTA Rhapsody analysis pipeline. Gene reads were aligned to the *H. vulgare* cv. Morex V3 reference genome (https://doi.org/10.5447/ipk/2021/3). Single-cell RNA sequencing was performed as described in (Demesa-Arevalo et al. 2026).

### Subtraction of DEGs induced during protoplast isolation

Total RNA was extracted from three replicates of freshly protoplasted root tips and whole root tips using the Direct-zol RNA Miniprep Plus kit (Zymo Research). Bulk RNA sequencing was performed using the Illumina NextSeq platform (Biomarker Technologies). Raw sequencing reads were quality-checked using FastQC (v0.11.9). No trimming was required for raw reads. Transcript abundance was quantified at the isoform level using Salmon (v1.5.2) and mapped against the MorexV3 reference (Mascher et al. 2021). Quantified data were processed using the 3D RNA-seq pipeline to calculate transcript/gene-level abundance and transcripts per million (TPM) values (Guo et al. 2021). Low-expressed genes were filtered out based on a threshold of CPM > 1 in at least three samples for differential expression testing. To identify the protoplast effect, pairwise comparisons were made between intact tissue samples and fresh protoplasts. The false discovery rate (FDR) was performed in R (v4.3.1) using the edgeR package, followed by a Benjamini-Hochberg (BH) procedure (Robinson et al. 2010). Genes were considered significantly differentially expressed (DEGs) if they met an absolute log2 fold-change (|log2FC|) > 2. 1308 genes were removed from the scRNAseq matrix before identifying the most variable features for cluster analysis.

### UMAP and replicate integration

UMAP and integration were performed as described in (Demesa-Arevalo et al. 2026). For FindClusters, we applied a resolution of 1.5.

### Cluster-specific gene marker identification

Marker gene identification was performed as described in (Demesa-Arevalo et al. 2026).

### Molecular Cartography

Multiplex smRNA fish of barley root tips was performed as described in (Demesa-Arevalo et al. 2026) using the same probeset as described.

### HCR RNA fluorescence in situ hybridization

The HCR RNA fluorescence *in situ* hybridization protocol was adapted from (Berg et al. 2025). Modifications were as follows: Root tips were collected in 20 mL Fixative I in a 50 mL Falcon tube and fixed overnight at 8°C. During ethanol dehydration, the samples were microwaved at 180 W six times for 25 seconds. Samples were stored in 10 mL of 70% ethanol at 8°C. For sectioning, root tips were embedded in 6% agarose. Sectioning was performed using a Leica VT1000 Vibratome in 1x PBS with a section thickness of 50 µm. Subsequently, the sections were stored in DPBS at 8°C in 2 mL Eppendorf tubes. For multiplexing, amplifier B1 was paired with fluorophore 488, and amplifier B2 was paired with fluorophore 647. Cell walls were stained with SR2200. Imaging was performed using a Zeiss LSM 980 confocal microscope with a 32x water immersion objective. SR2200 was excited at 405 nm, fluorophore 488 at 488 nm, and fluorophore 647 at 639 nm. Image processing was performed using Fiji (Schindelin et al. 2012).

### Cross-species comparisons

Using a cross-species annotation pipeline presented by (Ke et al. 2025), the cluster annotations of five published single-cell root tip atlases (Wendrich et al. 2020; Cantó-Pastor et al. 2024; Liu et al. 2021a; Ortiz-Ramírez et al. 2021; Ke et al. 2025) were assigned to the clusters in barley. The processed and annotated atlases of Arabidopsis, tomato, and wheat were kindly provided by the original authors, whereas the processed rice (Nipponbare cultivar) and maize datasets were obtained from the Gene Expression Omnibus (GSE146035 and GSE173087, respectively). As no cell annotations were available for rice, the provided expression matrix was normalized, scaled, and clustered (FindClusters in Seurat v5.1.0 with resolution 0.3) using the top 30 principal components and 2000 most variable features. Clusters were annotated using the markers provided by the original publication. For the annotated datasets and the barley dataset, cluster-specific differentially expressed genes (DEGs) were inferred using FindAllMarkers in Seurat (v5.1.0) (Hao et al. 2024). Next, each gene was assigned an orthologous group using the orthology information provided by PLAZA Monocots v5.1 (van Bel et al. 2022), inferred using OrthoFinder (v1.1.10) (Emms and Kelly 2019). For rice, maize, and barley, gene identifiers were first converted to correspond with those included in PLAZA. For rice, a RAP-DB-to-IRGSP conversion file was used (https://rapdb.dna.affrc.go.jp/download/archive/RAP-MSU_2024-07-12.txt.gz), for maize, an AGP-v4-to-NAM-v5.0 conversion file (https://ftp.psb.ugent.be/pub/plaza/plaza_public_monocots_05/IdConversion/id_conversion.zma.csv.gz), and for barley, a Morex-V3-to-BPG-v1 conversion file (https://panbarlex.ipk-gatersleben.de/downloads/morex_gene_id_lift_over.tar.gz) was used. Because gene conversion mappings were not always one-to-one between annotations, some DEGs were assigned to multiple orthologous groups, in which case those genes were omitted from the analysis. An initial filtering was then applied to the DEGs, retaining only DEGs with an adjusted p-value smaller than 0.05, a log fold change larger than 0.1, or genes where the difference between the percentage of cells expressing the gene in the cluster compared to outside the cluster was larger than 5%. Next, the remaining DEGs were ranked based on log fold change, and the DEGs associated with the top 300 unique orthologous groups per cluster were retained as inputs for the cross-species annotation pipeline. The pipeline was applied to barley and each comparator species, yielding five predicted annotations for each barley cluster, which were aggregated into a final prediction with very strong agreement (all five predictions agreed), strong agreement (four predictions agreed), weak agreement (three predictions agreed), or no agreement (less than three predictions agreed).

### Pseudotime analysis

The integrated dataset was used for pseudotime analysis with monocle3 (v.1.4.23). We performed an analysis on a subset of the initial dataset, clustering with a resolution of 1.5 and a dimensionality of 25.

### Generation of the Hvwox5-CRISPR Cas9 mutant

Guide sequences were designed using <u>ECRISP</u>, inducing a double-strand break in the first exon of the gene of interest, leading to a 4 base pair deletion at position 7 from the start (Suppl. Figure 3). Single sgRNA strands were hybridized and inserted into the shuttle vector pMGE624 via *a Bpi*I cut/ligation reaction. Using *Bsa*I, the sgRNA was transferred to the recipient vector pMGE599 via a second cut/ligation reaction (Kumar et al. 2018). Golden Promise Fast embryos were transformed with the final vector according to the protocol from (Hensel et al. 2009). Successful transformation was tested by polymerase chain reaction (PCR) on DNA extracted from T0 plants, targeting Cas9. Cas9 was removed by segregation in T1 plants, homozygous T2 plants carrying the *Hvwox5*-Mutation, but no Cas9 were identified by amplification of the genomic HvWOX5 sequence including the targeted sequence and subsequent Sanger sequencing.

### Root length and meristematic zone measurements

For root length measurements, seedlings were imaged at 6 DAG. Measurements were performed in Fiji (Schindelin et al. 2012) using the Free Line (root length) or Straight Line Tool (meristematic zone). The meristematic zone was defined as the distance between the QC region and the first cortex cell that doubled in size. Values were documented in Excel, and statistical tests (t-test, α = 0.05) and plotting were performed in R.

### EdU/mPSPI-Staining and metaxylem phenotyping

The EdU/mPSPI-Staining protocol was adjusted from (Burkart et al. 2022). Seedlings were grown on paper rolls for 6 days. EdU treatment was performed in 50 mL Falcon tubes containing 20 mL EdU solution (final concentration 10 µM EdU) for 6 h. Root tips were fixed for a minimum of 24 h before sectioning, as previously described, to a thickness of 100 µm (see HCR RNA *in situ* hybridization). Images were acquired using a Zeiss LSM 980 confocal microscope with a 32x water-immersion objective. We recorded longitudinal views for CSC layer quantification and z-stacks to obtain optical cross-sections for QC division scoring. Quantification and visualization were performed as described in (Burkart et al. 2022) with Origin 2026.

Metaxylem phenotypes were analyzed in ZenBlue 3.11 and quantified and visualized using Excel. Representative images were modified for brightness and contrast using Fiji (Schindelin et al. 2012).

### Root width measurements

Root width was measured using the Straight Line Tool in Fiji (Schindelin et al. 2012). The width of the QC region was considered. The values were collected in Excel, and statistical tests (t-test, α = 0.05) and plotting were performed in R.

## Acknowledgements

We would like to acknowledge Tobias Lautwein for his help with the BD Rhapsody platform and Ali Eljebbawi for help with statistical analyses. This work was supported by the German Research Council (Deutsche Forschungsgemeinschaft, DFG) via Research Unit 5235 (CSCS: Cereal Stem Cell Systems) and Germany’s Excellence Strategy—EXC21 2048—Project ID: 390686111. Work in the lab of Y.S. was supported by the DFG (grant number: 448353073). The confocal microscope (Zeiss LSM980) was financed by a DFG grant to Y.S. (project number: 553896058).

## Author contribution statement

A.D carried out all experiments and most of the analysis. J.S. performed cross-species comparisons. E.D-A. helped with establishing the protoplasting protocol and data analysis. T.L. performed RNA sequencing analysis. G.B. helped with barley transformation. P.S. built the scRNAseq web-browser. L.S.B. helped with establishing the protocol for HCR RNA *in situ* hybridization. YS. acquired funding and supervised the project. A.D and Y.S. wrote the manuscript; all authors reviewed and edited the manuscript.

## Confict of interests statement

The authors declare no conflict of interest.

