## Supplementary Figures for "A single-cell transcriptome atlas of the barley root apical meristem uncovers conserved and divergent roles of HvWOX5"

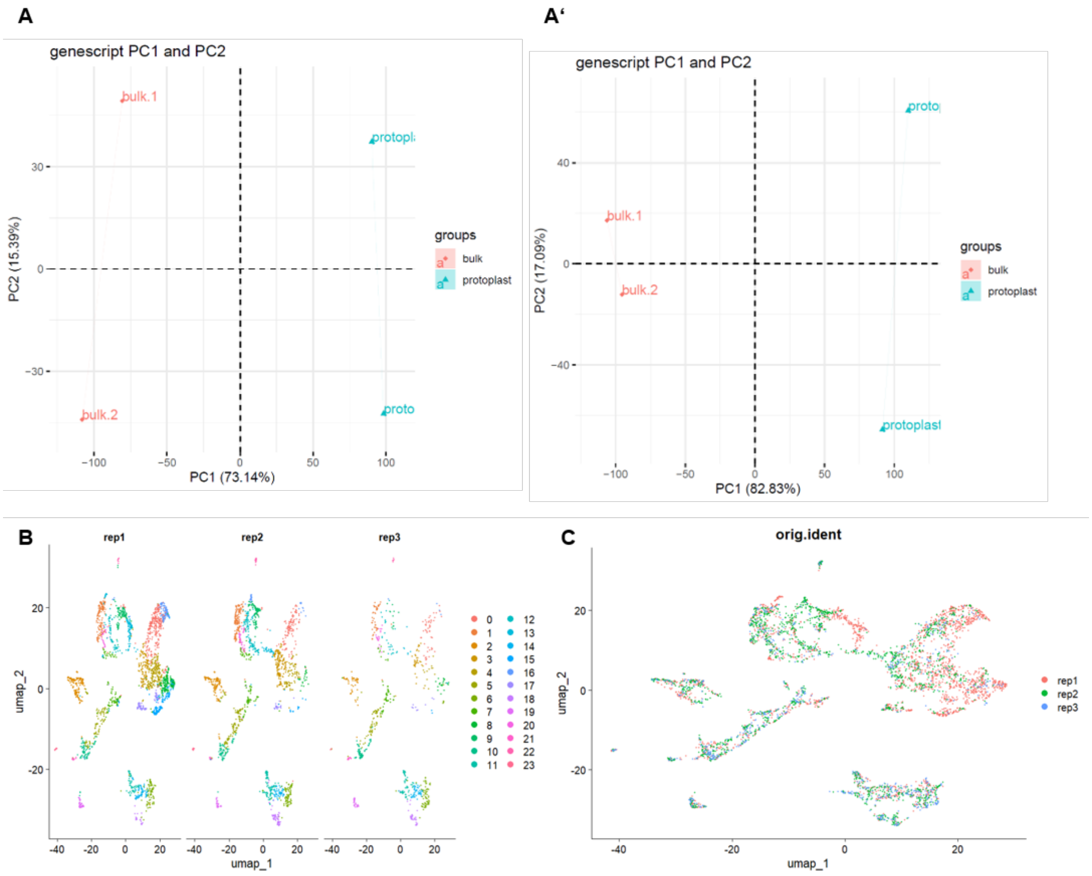


**Supplementary Figure 1. RNA sequencing for protoplasting effect removal and subsequent integration of scRNAseq replicates. A, A')** PC1 and PC2 of bulk RNA sequencing vs. protoplasted samples before **(A)** and after **(A')** batch effect removal. **B, C)** Clustering of three scRNAseq replicates after integration. **B)** clustering of individual replicates, **C)** overlap of all three replicates.

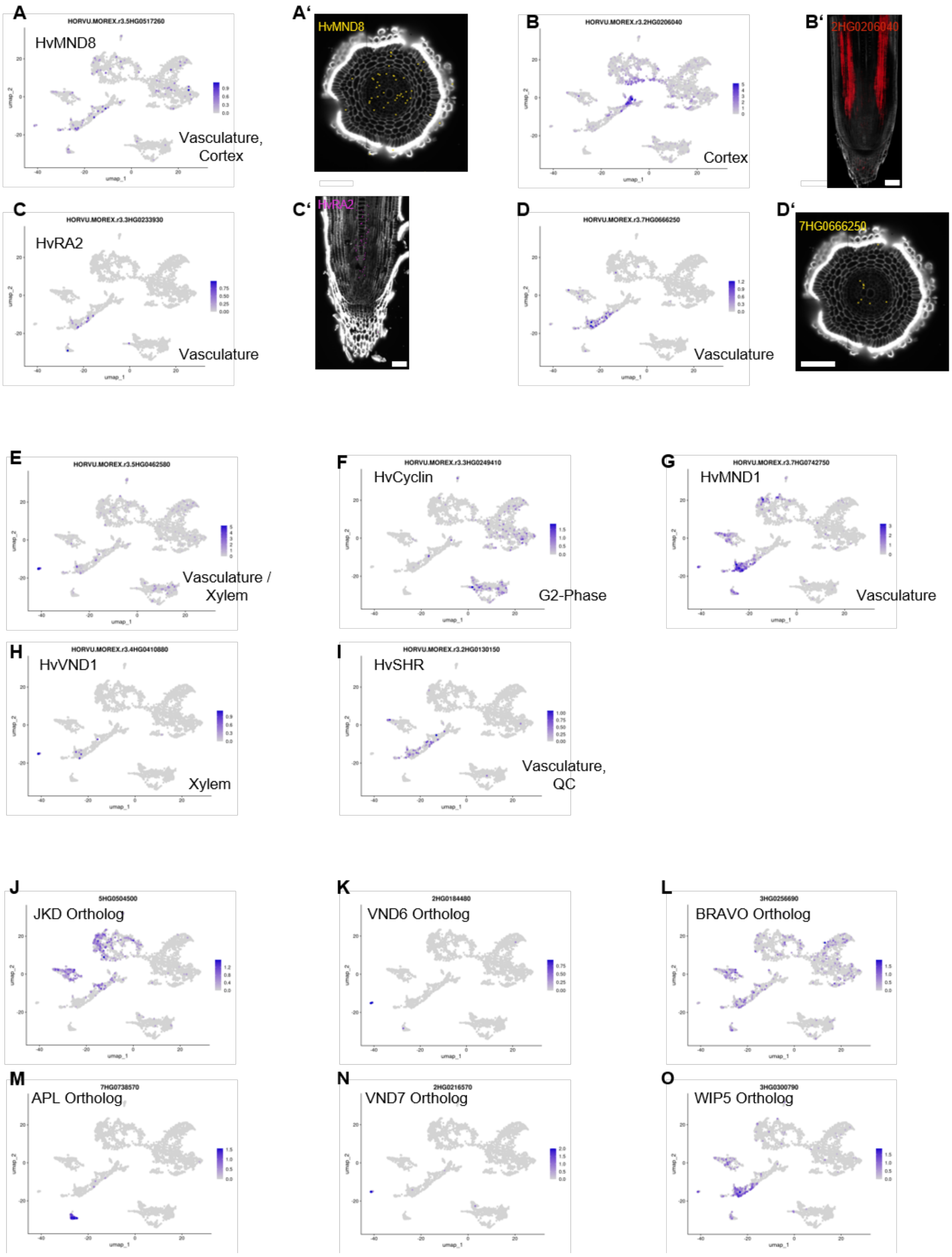


**Supplementary Figure 2. Marker gene expression for cluster annotation.** Gene expression patterns in 6 DAS barley RAMs of the following genes by Molecular Cartography/HCR *in situ* hybridization data **A,A')** *5HG0517260* (*HvMND8*), **B,B')** *2HG0206040*, **C,C')** *3HG0233930* (*HvRA2*) **D,D')** *7HG0666250*. Scale bars = 100 µm. **E-I)** Expression of marker genes from literature in our scRNAseq dataset. **E-H)** from Demesa-Arévalo *et al.* 2026, **I)** from Kirschner, 2017. **J-O)** expression of orthologs for cell type specific marker genes from Arabidopsis in the barley scRNAseq dataset. Orhologs were detected using ensembl plants.

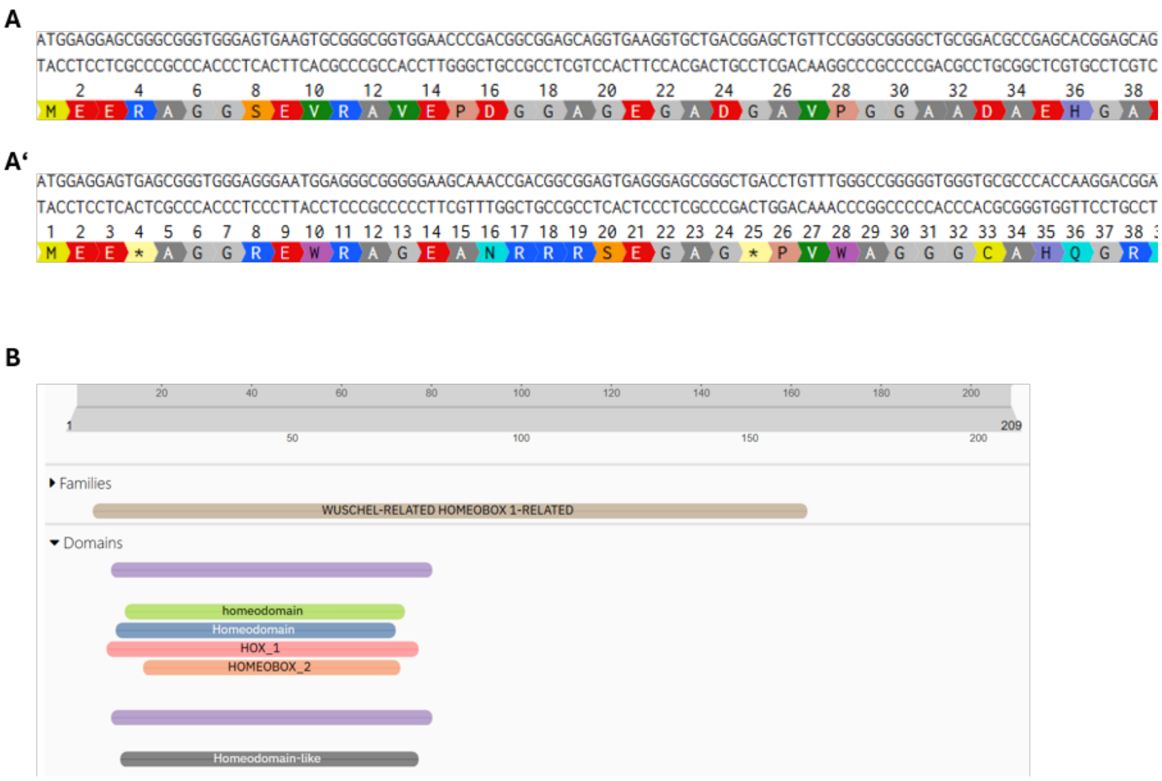


**Supplementary Figure 3. Sequencing and domain prediction of HvWOX5. A,A')** Sequences of wildtype HvWOX5 **(A)** and *Hvwox5* **(A'). B)** Domain prediction performed with InterPro (https://www.ebi.ac.uk/interpro/)

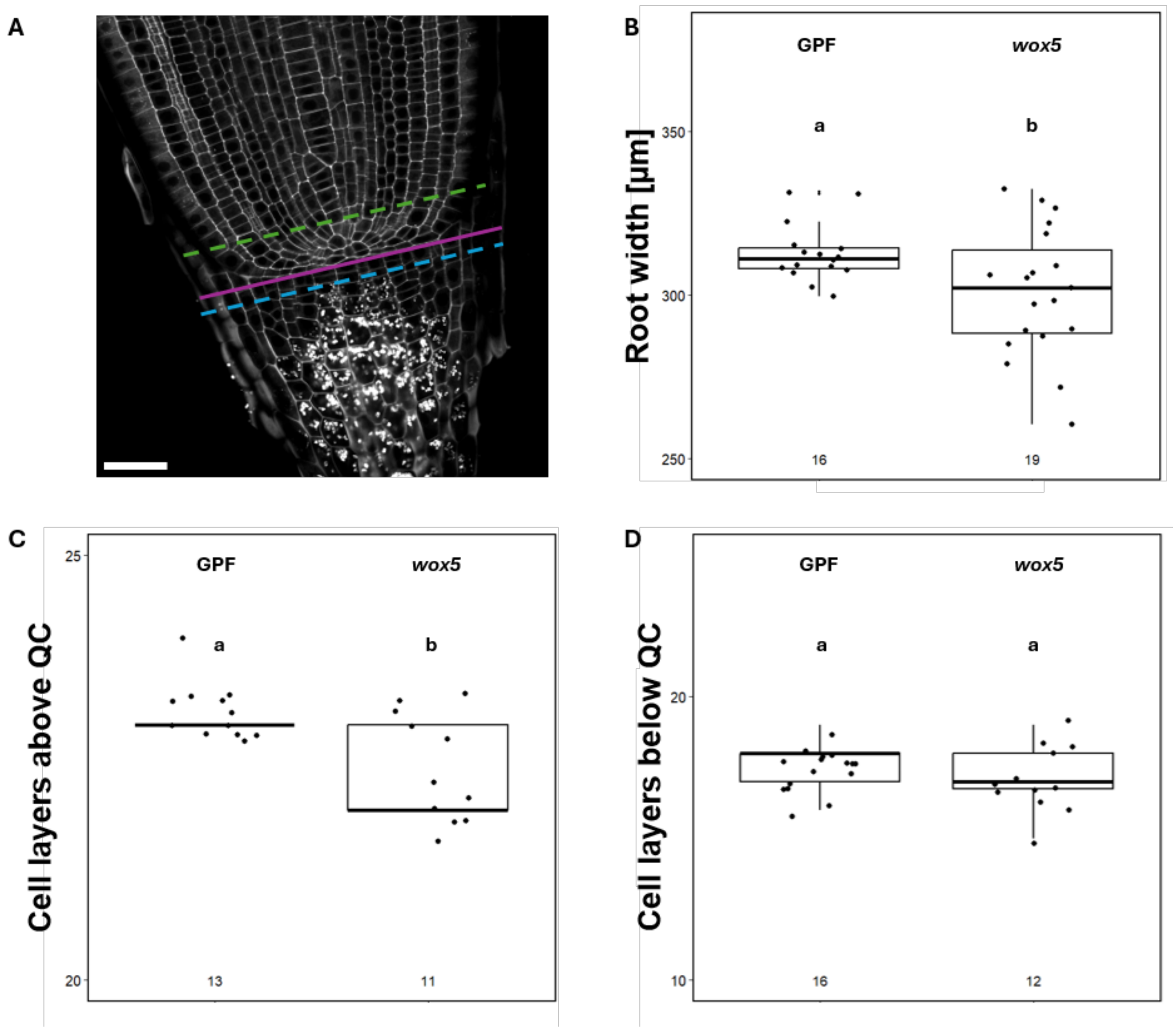


**Supplementary Figure 4. Phenotyping of the *Hvwox5*-mutant. A)** Schematic representation of zones for measuremens of root width (purple), cell layers above the QC (green) and cell layers below the QC (blue). Scale bar = 50 µm. **B)** Root width measurements, of GPF and *hvwox5*-roots. **C)** Number of cell layers above and **D)** below the QC in GPF and *hvwox5*-mutant roots. Statistical analyses were performed using t-tests, $\alpha = 0.05$. **B**: $p = 0.02562$, **C**: $p = 0.001647$, **D**: $p = 0.278$.
