## Supplementary Table 1 for "A single-cell transcriptome atlas of the barley root apical meristem uncovers conserved and divergent roles of HvWOX5"

**Supplementary Table 1. Genes upregulated during protoplasting.**

| Number | Gene.ID | Confidence | Biotype | Score | Description | annotation | classificati | Ontology | pfam | interpro |
|---|---|---|---|---|---|---|---|---|---|---|
| 1 | HORVU.MOREX.r3.1HG00022 | high | protein_cod | *** | 12-oxophyt | 4,5-didehyd | phytohormo | GO:000382 | PF00724 | IPR001155 |
| 2 | HORVU.MOREX.r3.1HG00022 | high | protein_cod | *** | 12-oxophyt | 4,5-didehyd | phytohormo | GO:000382 | PF00724 | IPR001155 |
| 3 | HORVU.MOREX.r3.1HG00023 | high | protein_cod | *** | 12-oxophyt | 4,5-didehyd | phytohormo | GO:000382 | PF00724 | IPR001155 |
| 4 | HORVU.MOREX.r3.1HG00041 | low | protein_cod | --* | Succinate d | alkyl hydrop | no mercato | GO:000588 | NA | NA |
| 5 | HORVU.MOREX.r3.1HG00049 | low | plastid_rela | *** | PGR5-like p | regulatory p | photosynthe | GO:001602 | NA | NA |
| 6 | HORVU.MOREX.r3.1HG00054 | high | protein_cod | *** | Glycosyltra | xylosyltrans | cell wall org | GO:001602 | PF04577 | IPR007657 |
| 7 | HORVU.MOREX.r3.1HG00056 | high | protein_cod | *** | Protein pho | clade C pho | protein mod | GO:000382 | PF00481 | IPR001932,IPR001932,IPR001932,IPR001932 |
| 8 | HORVU.MOREX.r3.1HG00057 | high | protein_cod | *** | Clathrin ass | enth domai | no mercato | GO:000554 | PF07651 | IPR011417,IPR008942 |
| 9 | HORVU.MOREX.r3.1HG00072 | high | protein_cod | *** | Zinc finger f | ZAT transcri | rna biosynth | GO:000367 | PF13912,PF | IPR007087,IPR007087 |
| 10 | HORVU.MOREX.r3.1HG00077 | low | non_coding | *** | Cytochrome | EC_1.14 oxi | enzyme clas | GO:000449 | PF00067 | IPR001128,IPR001128 |
| 11 | HORVU.MOREX.r3.1HG00084 | high | protein_cod | *** | 11S globuli | Globulin-typ | protein hom | GO:004573 | PF00190,PF | IPR011051,IPR006045,IPR006045 |
| 12 | HORVU.MOREX.r3.1HG00088 | high | protein_cod | --* | Clavata3/ES | CLE precurs | phytohormo | NA | NA | NA |
| 13 | HORVU.MOREX.r3.1HG00089 | high | protein_cod | *** | TOX high m | lysine speci | no mercato | NA | PF07795 | IPR012862 |
| 14 | HORVU.MOREX.r3.1HG00103 | high | protein_cod | *** | Non-specifi | arabinogala | cell wall org | GO:000686 | PF14368 | IPR016140,IPR016140 |
| 15 | HORVU.MOREX.r3.1HG00127 | high | protein_cod | *-* | Adenine/gu | cytokinin tr | phytohormo | GO:000534 | PF00860,PF | IPR006043,IPR006043 |
| 16 | HORVU.MOREX.r3.1HG00141 | high | protein_cod | *** | Transducin/ | substrate ad | protein hom | NA | PF00400,PF | IPR001680,IPR001680,IPR001680,IPR001680,IPR017986,IPR01798 |
| 17 | HORVU.MOREX.r3.1HG00145 | high | protein_cod | *** | Acyl-CoA de | isovaleryl-C | amino acid | GO:000399 | PF02771,PF | IPR009100,IPR013786,IPR006091,IPR009075,IPR009075 |
| 18 | HORVU.MOREX.r3.1HG00145 | high | protein_cod | *** | Peptide-N4- | peptide n4 r | no mercato | NA | PF12222 | IPR021102 |
| 19 | HORVU.MOREX.r3.1HG00154 | high | protein_cod | *** | Trihelix tran | TRIHELIX tra | rna biosynth | GO:000370 | NA | NA |
| 20 | HORVU.MOREX.r3.1HG00154 | high | protein_cod | *** | 2-oxoglutar | jasmonic ac | phytohormo | NA | PF14226,PF | IPR026992,IPR005123 |
| 21 | HORVU.MOREX.r3.1HG00164 | high | protein_cod | *** | Kinase, puta | DUF26 prot | protein mod | GO:000467 | PF01657,PF | IPR011009,IPR002902,IPR002902,IPR001245 |
| 22 | HORVU.MOREX.r3.1HG00179 | high | protein_cod | *** | Non-specifi | SnRK3 SNF1 | protein mod | NA | PF00069,PF | IPR000719,IPR004041,IPR011009 |
| 23 | HORVU.MOREX.r3.1HG00181 | high | protein_cod | *-* | Diacylglyce | cytosolic ac | lipid metab | GO:001674 | IPR012336 | |
| 24 | HORVU.MOREX.r3.1HG00182 | high | protein_cod | *** | Thioesteras | 1,4-dihydro | coenzyme n | NA | PF03061 | IPR003736,IPR006683,IPR029069 |
| 25 | HORVU.MOREX.r3.1HG00182 | high | protein_cod | *-* | Low temper | upf membra | no mercato | GO:001602 | PF01679 | IPR000612 |
| 26 | HORVU.MOREX.r3.1HG00193 | high | protein_cod | *** | Drebrin-like | drebrin prot | no mercato | NA | NA | NA |
| 27 | HORVU.MOREX.r3.1HG00204 | high | protein_cod | *** | Kinase fami | RLCK-VIIa re | protein mod | GO:000467 | PF07714 | IPR011009,IPR001245 |
| 28 | HORVU.MOREX.r3.1HG00209 | high | protein_cod | *** | Endo-1,31,4 | dlh domain | no mercato | GO:001678 | PF01738 | IPR029058,IPR002925 |
| 29 | HORVU.MOREX.r3.1HG00213 | high | protein_cod | *** | Glutathione | class phi gl | redox home | GO:001674 | PF02798,PF | IPR004045,IPR012336,IPR004046,IPR010987 |
| 30 | HORVU.MOREX.r3.1HG00213 | high | protein_cod | *** | Glutathione | class phi gl | redox home | GO:001674 | PF02798,PF | IPR010987,IPR012336,IPR004045,IPR004046 |
| 31 | HORVU.MOREX.r3.1HG00213 | high | protein_cod | *** | Glutathione | class phi gl | redox home | GO:001674 | PF00043,PF | IPR004046,IPR012336,IPR010987,IPR004045 |
| 32 | HORVU.MOREX.r3.1HG00219 | high | protein_cod | *** | 1-aminocyc | 1-aminocyc | phytohormo | GO:001649 | PF14226,PF | IPR026992,IPR005123 |
| 33 | HORVU.MOREX.r3.1HG00227 | high | protein_cod | *** | Rubber elon | lipid droplet | lipid metab | GO:000374 | PF05755 | IPR008802 |
| 34 | HORVU.MOREX.r3.1HG00238 | high | protein_cod | *** | Ethylene re | ethylene re | phytohormo | GO:000015 | PF02518,PF | IPR003594,IPR029016,IPR003018,IPR003594,IPR003661,IPR003661 |
| 35 | HORVU.MOREX.r3.1HG00239 | high | protein_cod | *-* | GATA transc | A/B-GATA-t | rna biosynth | GO:000370 | PF00320 | IPR000679 |
| 36 | HORVU.MOREX.r3.1HG00245 | high | protein_cod | *** | Universal st | usp domain | no mercato | GO:000695 | PF00582 | IPR006016 |
| 37 | HORVU.MOREX.r3.1HG00245 | high | protein_cod | --* | Transcriptio | bhlh domai | no mercato | NA | IPR011598 | |
| 38 | HORVU.MOREX.r3.1HG00255 | high | protein_cod | *-* | RING/U-box | RING-H2-cl | protein hom | NA | PF13639 | IPR001841 |
| 39 | HORVU.MOREX.r3.1HG00275 | high | protein_cod | *** | Plant/MSJ1 | not classifie | no mercato | NA | NA | NA |
| 40 | HORVU.MOREX.r3.1HG00276 | high | protein_cod | *** | Plant/MSJ1 | not classifie | no mercato | NA | NA | NA |
| 41 | HORVU.MOREX.r3.1HG00276 | high | protein_cod | *** | Plant/MSJ1 | not classifie | no mercato | NA | NA | NA |
| 42 | HORVU.MOREX.r3.1HG00288 | low | protein_cod | --* | Phosphorib | not classifie | no mercato | NA | NA | NA |
| 43 | HORVU.MOREX.r3.1HG00297 | high | protein_cod | *** | Malic enzym | cytosolic N | lipid metab | GO:000447 | PF03949,PF | IPR016040,IPR012302,IPR012301 |
| 44 | HORVU.MOREX.r3.1HG00391 | high | protein_cod | *** | RING finger | E3 ubiquitin | protein hom | GO:001602 | PF00097 | IPR018957 |
| 45 | HORVU.MOREX.r3.1HG00398 | high | protein_cod | *** | RING finger | E3 ubiquitin | protein hom | GO:000827 | PF05495,PF | IPR008913,IPR027370 |
| 46 | HORVU.MOREX.r3.1HG00407 | high | protein_cod | *** | Plant/T7N9- | D-2-hydroxy | amino acid | NA | PF07063 | IPR009770 |
| 47 | HORVU.MOREX.r3.1HG00414 | high | protein_cod | *** | Receptor pr | LRR-XII prot | protein mod | GO:000016 | PF08263,PF | IPR013210,IPR032675,IPR011009,IPR000719,IPR001611,IPR001611,IPR001611,IPR032675 |
| 48 | HORVU.MOREX.r3.1HG00415 | high | protein_cod | *** | Esterase/lip | phytyl ester | coenzyme n | GO:000414 | PF03982,PF | IPR007130,IPR029058,IPR022742 |
| 49 | HORVU.MOREX.r3.1HG00421 | high | protein_cod | *-* | PPPDE thiol | deubiquitin | protein hom | NA | PF05903 | IPR008580 |
| 50 | HORVU.MOREX.r3.1HG00432 | high | protein_cod | *** | MYB | quality prot | no mercato | GO:000367 | PF00249,PF | IPR001005,IPR001005,IPR009057 |
| 51 | HORVU.MOREX.r3.1HG00438 | high | protein_cod | *** | senescence | senescence | no mercato | NA | PF04520 | IPR007608 |
| 52 | HORVU.MOREX.r3.1HG00440 | high | protein_cod | *-* | Transmemb | thioredoxin | no mercato | GO:001602 | NA | NA |
| 53 | HORVU.MOREX.r3.1HG00444 | high | protein_cod | *** | Receptor-lik | DUF26 prot | protein mod | GO:000467 | PF01657,PF | IPR002902,IPR002902,IPR011009,IPR001245 |
| 54 | HORVU.MOREX.r3.1HG00461 | high | protein_cod | *** | Ser/Thr prot | metallopho | no mercato | GO:001678 | PF00149 | IPR029052,IPR029052,IPR004843 |
| 55 | HORVU.MOREX.r3.1HG00464 | high | protein_cod | *** | Plant basic | basic secret | no mercato | NA | PF04450 | IPR007541 |
| 56 | HORVU.MOREX.r3.1HG00473 | high | protein_cod | *** | CBS domai | cbs domain | no mercato | NA | PF00571 | IPR000644 |
| 57 | HORVU.MOREX.r3.1HG00475 | high | protein_cod | *** | RING finger | E3 ubiquitin | protein hom | NA | NA | NA |
| 58 | HORVU.MOREX.r3.1HG00479 | low | plastid_rela | *** | Thylakoid lu | thylakoid lu | no mercato | NA | PF00805,PF | IPR001646,IPR001646 |
| 59 | HORVU.MOREX.r3.1HG00495 | low | non_coding | *** | Cytochrome | FCC deform | coenzyme n | GO:000449 | PF00067 | IPR001128,IPR001128 |
| 60 | HORVU.MOREX.r3.1HG00499 | high | protein_cod | *** | Auxin-respo | auxin respo | no mercato | GO:000563 | PF02519 | IPR003676 |
| 61 | HORVU.MOREX.r3.1HG00500 | high | protein_cod | *** | Response t | regulatory p | nutrient upt | NA | NA | NA |
| 62 | HORVU.MOREX.r3.1HG00507 | high | protein_cod | *** | ATP-depend | EC_3.6 hydr | enzyme clas | GO:000552 | PF14363,PF | IPR025753,IPR003959,IPR027417 |
| 63 | HORVU.MOREX.r3.1HG00509 | high | protein_cod | *** | Trehalase | trehalase; E | carbohydra | GO:000382 | PF01204 | IPR008928,IPR001661 |
| 64 | HORVU.MOREX.r3.1HG00517 | high | protein_cod | *** | Glutathione | class tau gl | redox home | GO:000446 | PF00043,PF | IPR004046,IPR012336,IPR004045,IPR010987 |

| | | | | | | | | | | |
|---|---|---|---|---|---|---|---|---|---|---|
| 65 | HORVU.MOREX.r3.1HG00517 | high | protein_cod | *** | Glutathione | class tau gl | redox home | GO:001674 | PF13417 | IPR012336,IPR004045,IPR010987 |
| 66 | HORVU.MOREX.r3.1HG00518 | high | protein_cod | *** | Glutathione | class tau gl | redox home | GO:001674 | PF02798,PF | IPR004045,IPR010987,IPR004046,IPR01233 |
| 67 | HORVU.MOREX.r3.1HG00518 | high | protein_cod | *** | Glutathione | class tau gl | redox home | GO:001674 | PF13417 | IPR010987,IPR004045,IPR012336 |
| 68 | HORVU.MOREX.r3.1HG00518 | high | protein_cod | *** | Glutathione | class tau gl | redox home | GO:001674 | PF13417 | IPR004045,IPR012336,IPR010987 |
| 69 | HORVU.MOREX.r3.1HG00518 | high | protein_cod | *** | Glutathione | class tau gl | redox home | GO:001674 | PF13417 | IPR004045,IPR012336,IPR010987 |
| 70 | HORVU.MOREX.r3.1HG00518 | high | protein_cod | *** | Glutathione | class tau gl | redox home | GO:001674 | PF13417 | IPR004045,IPR012336,IPR010987 |
| 71 | HORVU.MOREX.r3.1HG00519 | high | protein_cod | *** | Glutathione | class tau gl | redox home | GO:001674 | PF13417 | IPR004045,IPR012336,IPR010987 |
| 72 | HORVU.MOREX.r3.1HG00529 | low | non_coding | *** | 60S riboson | pre-60S rib | protein bios | GO:000563 | PF04981 | IPR007064 |
| 73 | HORVU.MOREX.r3.1HG00530 | high | protein_cod | *** | PLATZ trans | PLATZ trans | rna biosynt | NA | PF04640 | IPR006734 |
| 74 | HORVU.MOREX.r3.1HG00534 | high | protein_cod | *** | Cinnamoyl- | 3beta hsd d | no mercato | NA | PF01370 | IPR016040,IPR001509 |
| 75 | HORVU.MOREX.r3.1HG00540 | high | protein_cod | *** | zinc finger B | not classifie | no mercato | NA | NA | NA |
| 76 | HORVU.MOREX.r3.1HG00545 | high | protein_cod | *** | Ring finger p | RING-H2-cl | protein hom | GO:001602 | PF13639 | IPR001841 |
| 77 | HORVU.MOREX.r3.1HG00548 | high | protein_cod | *** | Expansin-lik | alpha-like-c | cell wall org | GO:000557 | PF03330,PF | IPR009009,IPR007117,IPR007117,IPR00900 |
| 78 | HORVU.MOREX.r3.1HG00552 | high | protein_cod | *** | RING/U-box | RING-H2-cl | protein hom | GO:001602 | PF13639 | IPR001841 |
| 79 | HORVU.MOREX.r3.1HG00554 | low | non_coding | *** | Cytochrome | heme chape | cellular res | GO:000588 | PF03100 | IPR004329,IPR004329 |
| 80 | HORVU.MOREX.r3.1HG00560 | high | protein_cod | *** | Cortical cell | NA | NA | NA | PF14547 | IPR027923,IPR016140 |
| 81 | HORVU.MOREX.r3.1HG00564 | high | protein_cod | *-* | Hexosyltran | glucuronos | no mercato | GO:001602 | IPR029044 | |
| 82 | HORVU.MOREX.r3.1HG00574 | high | protein_cod | *** | Metacaspas | metacaspas | multi-proce | NA | PF06943 | IPR005735,IPR005735 |
| 83 | HORVU.MOREX.r3.1HG00574 | high | protein_cod | *** | Kinase fami | SnRK2 SNF1 | protein mod | GO:000016 | PF00069 | IPR011009,IPR000719 |
| 84 | HORVU.MOREX.r3.1HG00575 | high | protein_cod | *** | PRA1 family | pra family p | no mercato | GO:001602 | PF03208 | IPR004895 |
| 85 | HORVU.MOREX.r3.1HG00579 | high | protein_cod | *** | Sugar trans | monosacch | solute trans | GO:000521 | PF00083 | IPR005828,IPR003663,IPR020846 |
| 86 | HORVU.MOREX.r3.1HG00583 | high | protein_cod | *** | 2-oxoglutar | EC_1.14 oxi | enzyme clas | GO:001649 | PF14226,PF | IPR026992,IPR005123 |
| 87 | HORVU.MOREX.r3.1HG00589 | high | protein_cod | *** | DUF4228 d | duf 4228 do | no mercato | NA | PF14009 | IPR025322 |
| 88 | HORVU.MOREX.r3.1HG00592 | high | protein_cod | *** | Calcium bin | calcium bin | no mercato | GO:000550 | PF13499,PF | IPR002048,IPR002048,IPR011992 |
| 89 | HORVU.MOREX.r3.1HG00597 | high | protein_cod | *** | Diacylglyce | diacylglycer | lipid metab | GO:000016 | PF00130,PF | IPR002219,IPR001206,IPR016064,IPR00075 |
| 90 | HORVU.MOREX.r3.1HG00607 | high | protein_cod | *-* | Ethylene-re | transcriptio | external stir | GO:000367 | PF00847 | IPR016177,IPR001471 |
| 91 | HORVU.MOREX.r3.1HG00610 | high | protein_cod | *-* | RING/U-box | RING-H2-cl | protein hom | GO:001602 | PF13639 | IPR001841 |
| 92 | HORVU.MOREX.r3.1HG00612 | high | protein_cod | *-* | Flagellar bic | nucleolar pr | no mercato | NA | NA | NA |
| 93 | HORVU.MOREX.r3.1HG00623 | high | protein_cod | *** | Dehydratior | subgroup EI | rna biosynt | GO:000367 | PF00847 | IPR001471,IPR016177 |
| 94 | HORVU.MOREX.r3.1HG00628 | low | non_coding | *** | Cytochrome | EC_1.14 oxi | enzyme clas | GO:000449 | PF00067 | IPR001128,IPR001128 |
| 95 | HORVU.MOREX.r3.1HG00633 | high | protein_cod | *** | Eukaryotic a | Pepsin-type | protein hom | GO:000419 | PF14541,PF | IPR032799,IPR021109,IPR032861 |
| 96 | HORVU.MOREX.r3.1HG00638 | high | protein_cod | *** | Glycosyltrar | EC_2.4 glyc | enzyme clas | GO:000815 | PF00201 | IPR002213 |
| 97 | HORVU.MOREX.r3.1HG00639 | high | protein_cod | *** | CASP-like p | casp proteir | no mercato | NA | PF04535 | IPR006702,IPR006459 |
| 98 | HORVU.MOREX.r3.1HG00640 | high | protein_cod | *** | Rhomboid f | Rhomboid-t | protein hom | GO:000425 | PF00641,PF | IPR001876,IPR001876,IPR022764 |
| 99 | HORVU.MOREX.r3.1HG00640 | low | protein_cod | --* | Elongation f | not classifie | no mercato | GO:000016 | NA | NA |
| 100 | HORVU.MOREX.r3.1HG00648 | high | protein_cod | *-* | ATP-depend | EC_3.6 hydr | enzyme clas | GO:000016 | PF00004,PF | IPR003959,IPR003959,IPR025753,IPR027417,IPR027417 |
| 101 | HORVU.MOREX.r3.1HG00651 | high | protein_cod | *** | NAC domai | NAC transc | rna biosynt | GO:000367 | PF02365 | IPR003441,IPR003441 |
| 102 | HORVU.MOREX.r3.1HG00660 | high | protein_cod | --* | with no lysir | not classifie | no mercato | NA | NA | NA |
| 103 | HORVU.MOREX.r3.1HG00665 | high | protein_cod | *** | F-box family | substrate ac | protein hom | NA | PF13516,PF | IPR001611,IPR001611,IPR001611 |
| 104 | HORVU.MOREX.r3.1HG00669 | high | protein_cod | *-* | Octicosape | pb domain | no mercato | GO:000467 | PF00564 | IPR000270 |
| 105 | HORVU.MOREX.r3.1HG00671 | high | protein_cod | -** | Wound-ind | bowman bir | no mercato | GO:000486 | NA | NA |
| 106 | HORVU.MOREX.r3.1HG00682 | low | protein_cod | --* | RNA-bindin | not classifie | no mercato | NA | NA | NA |
| 107 | HORVU.MOREX.r3.1HG00683 | high | protein_cod | --* | Nuclear por | not classifie | no mercato | NA | NA | NA |
| 108 | HORVU.MOREX.r3.1HG00690 | high | protein_cod | *** | Glycerol-3- | NAD-depen | lipid metab | GO:000436 | PF01210,PF | IPR011128,IPR008927,IPR006109,IPR016040,IPR01604 |
| 109 | HORVU.MOREX.r3.1HG00690 | high | protein_cod | *** | Calcium-tra | P2B-type ca | solute trans | GO:000016 | PF12515,PF | IPR023299,IPR024750,IPR006068,IPR001757,IPR001757,IPR008250,IPR023214,IPR023214,IPR004014,IPR0064 |
| 110 | HORVU.MOREX.r3.1HG00693 | high | protein_cod | *** | Cysteine pr | Cystatin pro | protein hom | GO:000486 | PF16845 | IPR000010 |
| 111 | HORVU.MOREX.r3.1HG00694 | high | protein_cod | *** | Aldehyde de | hydroxycinn | cell wall org | GO:000815 | PF00171 | IPR015590,IPR016161 |
| 112 | HORVU.MOREX.r3.1HG00703 | high | protein_cod | *** | Protein kina | hydrogen pe | redox home | GO:000016 | PF08263,PF | IPR032675,IPR013210,IPR011009,IPR001245 |
| 113 | HORVU.MOREX.r3.1HG00710 | high | protein_cod | *** | Expansin | alpha-class | cell wall org | GO:000557 | PF01357,PF | IPR009009,IPR007117,IPR009009,IPR007117 |
| 114 | HORVU.MOREX.r3.1HG00712 | high | protein_cod | *** | U-box doma | E3 ubiquitin | protein hom | GO:000484 | PF00514,PF | IPR016024,IPR000225,IPR003613 |
| 115 | HORVU.MOREX.r3.1HG00714 | high | protein_cod | *** | TOM1-like p | ubiquitin ad | vesicle traff | GO:000562 | PF03127,PF | IPR004152,IPR002014,IPR008942 |
| 116 | HORVU.MOREX.r3.1HG00720 | high | protein_cod | *** | Phosphatid | upf protein | no mercato | NA | PF01161 | IPR008914,IPR005247,IPR008914 |
| 117 | HORVU.MOREX.r3.1HG00721 | high | protein_cod | *** | Ammonium | ammonium | solute trans | GO:000851 | PF00909 | IPR024041,IPR001905,IPR024041 |
| 118 | HORVU.MOREX.r3.1HG00721 | high | protein_cod | *** | cotton fiber | duf 761 don | no mercato | NA | PF05553 | IPR008480 |
| 119 | HORVU.MOREX.r3.1HG00740 | high | protein_cod | *** | Phosphatid | inositol pho | lipid metab | GO:001602 | PF14360 | IPR025749 |
| 120 | HORVU.MOREX.r3.1HG00752 | high | protein_cod | *** | Fructose-1, | cytosolic fru | carbohydra | GO:000573 | PF00316 | IPR033391 |
| 121 | HORVU.MOREX.r3.1HG00757 | high | protein_cod | *** | Subtilisin-li | protease *( | protein hom | GO:000425 | PF05922,PF | IPR010259,IPR009020,IPR000209,IPR000209,IPR000209,IPR00020 |
| 122 | HORVU.MOREX.r3.1HG00766 | high | protein_cod | *** | Receptor-lik | SCREW pep | phytohorm | GO:000016 | PF07714,PF | IPR032675,IPR011009,IPR001245,IPR013210,IPR001611,IPR00161 |
| 123 | HORVU.MOREX.r3.1HG00772 | high | protein_cod | *** | Hemoglobir | goshi non s | no mercato | GO:000534 | PF00042 | IPR000971,IPR009050 |
| 124 | HORVU.MOREX.r3.1HG00777 | high | protein_cod | *** | Calcium-tra | P2B-type ca | solute trans | GO:000016 | PF00122,PF | IPR001757,IPR001757,IPR008250,IPR006068,IPR023214,IPR023214,IPR006408,IPR0040 |
| 125 | HORVU.MOREX.r3.1HG00779 | low | protein_cod | --- | Transcriptio | CLE precurs | phytohorm | NA | NA | NA |
| 126 | HORVU.MOREX.r3.1HG00779 | high | protein_cod | --* | sequence-s | not classifie | no mercato | NA | NA | NA |
| 127 | HORVU.MOREX.r3.1HG00781 | high | protein_cod | *** | 2-oxoglutar | EC_1.14 oxi | enzyme clas | GO:001649 | PF03171,PF | IPR005123,IPR026992 |
| 128 | HORVU.MOREX.r3.1HG00789 | high | protein_cod | *** | p-loop cont | zeta toxin d | no mercato | GO:000552 | PF06414 | IPR027417,IPR027417,IPR010488 |
| 129 | HORVU.MOREX.r3.1HG00795 | high | protein_cod | *** | Protein kina | EC_2.7 tran | enzyme clas | GO:000016 | PF00069 | IPR011009,IPR000719 |

| | | | | | | | | | | |
|---|---|---|---|---|---|---|---|---|---|---|
| 130 | HORVU.MOREX.r3.1HG00796 | high | protein_cod | *** | Sec14p-like | phosphoino | multi-proce | GO:001602 | PF00650,PF | IPR001251,IPR001251,IPR011074,IPR011074 |
| 131 | HORVU.MOREX.r3.1HG00811 | high | protein_cod | *** | Type I inosit | type-I inosit | multi-proce | GO:001678 | PF03372 | IPR005135,IPR005135,IPR005135 |
| 132 | HORVU.MOREX.r3.1HG00811 | high | protein_cod | *** | Choline/eth | choline kina | lipid metab | GO:001630 | IPR011009 | |
| 133 | HORVU.MOREX.r3.1HG00812 | high | protein_cod | *** | Serine/thre | SD-2 protei | protein moc | GO:000016 | PF00069,PF | IPR000719,IPR001480,IPR000858,IPR003609,IPR011009,IPR00148 |
| 134 | HORVU.MOREX.r3.1HG00819 | high | protein_cod | *** | senescence | not classifie | no mercato | NA | PF04520 | IPR007608 |
| 135 | HORVU.MOREX.r3.1HG00820 | high | protein_cod | *** | Heat shock | HSF transcr | rna biosynth | GO:000367 | PF00447 | IPR011991,IPR000232 |
| 136 | HORVU.MOREX.r3.1HG00827 | high | protein_cod | *** | Late embryo | lea 2 domai | no mercato | GO:001602 | PF03168 | IPR004864 |
| 137 | HORVU.MOREX.r3.1HG00836 | high | protein_cod | *** | RING/U-box | RING-H2-cl | protein hom | NA | PF13639 | IPR001841 |
| 138 | HORVU.MOREX.r3.1HG00836 | high | protein_cod | *-* | Protein PLA | protein plas | no mercato | GO:000716 | PF05701 | IPR008545 |
| 139 | HORVU.MOREX.r3.1HG00839 | high | protein_cod | *** | 1,2-dihydro | acireducton | amino acid | GO:000550 | PF03079 | IPR004313,IPR011051 |
| 140 | HORVU.MOREX.r3.1HG00857 | high | protein_cod | *** | Core-2/I-bra | core i branc | no mercato | GO:000837 | PF02485 | IPR003406 |
| 141 | HORVU.MOREX.r3.1HG00862 | high | protein_cod | --* | NAD(P)-bin | 4fe 4s ferre | no mercato | NA | NA | NA |
| 142 | HORVU.MOREX.r3.1HG00864 | high | protein_cod | *** | Auxin-respo | transcriptio | phytohormo | GO:000563 | PF02309 | IPR033389 |
| 143 | HORVU.MOREX.r3.1HG00868 | high | protein_cod | *** | Pirin-like pr | transcriptio | rna biosynth | NA | PF02678,PF | IPR003829,IPR008778,IPR011051 |
| 144 | HORVU.MOREX.r3.1HG00878 | high | protein_cod | *** | Glutaredoxi | glutaredoxir | protein moc | GO:000562 | PF00462 | IPR012336,IPR002109,IPR011905 |
| 145 | HORVU.MOREX.r3.1HG00885 | low | non_coding | *** | 60S ribosom | 60s ribosom | no mercato | GO:000584 | NA | NA |
| 146 | HORVU.MOREX.r3.1HG00887 | high | protein_cod | *** | WRKY trans | transcriptio | rna biosynth | GO:000367 | PF03106 | IPR003657,IPR003657 |
| 147 | HORVU.MOREX.r3.1HG00894 | high | protein_cod | *** | Aspartic pro | Pepsin-type | protein hom | GO:000419 | PF05184,PF | IPR007856,IPR033121,IPR008138,IPR021109,IPR021109,IPR011001 |
| 148 | HORVU.MOREX.r3.1HG00901 | high | protein_cod | *** | DUF1645 fa | histone lysir | no mercato | NA | PF07816 | IPR012442 |
| 149 | HORVU.MOREX.r3.1HG00910 | high | protein_cod | *** | 2-oxoglutar | quality prot | no mercato | GO:001649 | PF03171 | IPR005123 |
| 150 | HORVU.MOREX.r3.1HG00910 | high | protein_cod | --* | DNA-direct | kow domain | no mercato | GO:000237 | NA | NA |
| 151 | HORVU.MOREX.r3.1HG00917 | high | protein_cod | *** | DUF1645 fa | not classifie | no mercato | NA | PF07816 | IPR012442 |
| 152 | HORVU.MOREX.r3.1HG00918 | high | protein_cod | *** | Cinnamoyl- | 3beta hsd d | no mercato | GO:000385 | PF01073 | IPR002225,IPR016040 |
| 153 | HORVU.MOREX.r3.1HG00924 | high | protein_cod | *** | WRKY trans | transcriptio | rna biosynth | GO:000367 | PF03106 | IPR003657,IPR003657 |
| 154 | HORVU.MOREX.r3.1HG00929 | high | protein_cod | *** | FAD-binding | EC_1.1 oxid | enzyme clas | GO:000382 | PF08031,PF | IPR016166,IPR012951,IPR006094 |
| 155 | HORVU.MOREX.r3.1HG00930 | high | protein_cod | *** | Indole-3-ac | jasmonoyl-a | phytohormo | NA | PF03321 | IPR004993 |
| 156 | HORVU.MOREX.r3.1HG00948 | high | protein_cod | --* | ATP-depend | dde tnp don | no mercato | GO:000016 | NA | NA |
| 157 | HORVU.MOREX.r3.1HG00951 | high | protein_cod | *** | UDP-glucos | UDP-D-gluc | carbohydrat | GO:000397 | PF16363 | IPR016040,IPR016040,IPR005886 |
| 158 | HORVU.MOREX.r3.2HG00961 | low | protein_cod | --* | WEB family | alkyl hydrop | no mercato | NA | NA | NA |
| 159 | HORVU.MOREX.r3.2HG00989 | high | protein_cod | *** | Glutamyl-tR | glutamyl trn | no mercato | GO:001674 | PF01425 | IPR023631,IPR023631 |
| 160 | HORVU.MOREX.r3.2HG00991 | high | protein_cod | *** | 12-oxophyt | 4,5-didehyd | phytohormo | GO:000382 | PF00724 | IPR001155 |
| 161 | HORVU.MOREX.r3.2HG00998 | high | protein_cod | *** | Heat shock | chaperone | protein hom | GO:000552 | PF00183,PF | IPR001404,IPR003594,IPR003594,IPR020568 |
| 162 | HORVU.MOREX.r3.2HG01005 | high | protein_cod | *** | Glycosyltra | nucleotid tr | no mercato | GO:000013 | PF03407 | IPR005069 |
| 163 | HORVU.MOREX.r3.2HG01012 | low | protein_cod | --* | Trafficking p | ring type do | no mercato | GO:000551 | NA | NA |
| 164 | HORVU.MOREX.r3.2HG01012 | low | protein_cod | --* | RNA binding | ring type do | no mercato | NA | NA | NA |
| 165 | HORVU.MOREX.r3.2HG01016 | high | protein_cod | *-* | Receptor-lik | WAK/WAKL | protein moc | GO:000016 | PF13947,PF | IPR011009,IPR025287,IPR001245 |
| 166 | HORVU.MOREX.r3.2HG01063 | high | protein_cod | *** | Glycosyltra | EC_2.4 glyc | enzyme clas | GO:000815 | PF00201 | IPR002213 |
| 167 | HORVU.MOREX.r3.2HG01064 | high | protein_cod | *** | Glycosyltra | EC_2.4 glyc | enzyme clas | GO:000815 | PF00201 | IPR002213 |
| 168 | HORVU.MOREX.r3.2HG01076 | high | protein_cod | *** | Protein kina | RLCK-VIIa r | protein moc | GO:000016 | PF07714 | IPR011009,IPR001245 |
| 169 | HORVU.MOREX.r3.2HG01079 | high | protein_cod | *** | MLO-like pr | calmodulin- | solute trans | GO:000551 | PF03094 | IPR004326 |
| 170 | HORVU.MOREX.r3.2HG01101 | high | protein_cod | *** | Transcriptio | bHLH trans | rna biosynth | GO:004698 | NA | NA |
| 171 | HORVU.MOREX.r3.2HG01114 | high | protein_cod | *** | NAC domai | NAC transc | rna biosynth | GO:000367 | PF02365 | IPR003441,IPR003441 |
| 172 | HORVU.MOREX.r3.2HG01129 | high | protein_cod | *-* | Ethylene-re | subgroup E | rna biosynth | GO:000367 | PF00847 | IPR016177,IPR001471 |
| 173 | HORVU.MOREX.r3.2HG01132 | high | protein_cod | *** | Hypoxia-res | component | cellular resp | GO:001602 | PF04588 | IPR007667 |
| 174 | HORVU.MOREX.r3.2HG01133 | high | protein_cod | *** | Universal st | usp domain | no mercato | GO:000695 | PF00582 | IPR006016 |
| 175 | HORVU.MOREX.r3.2HG01134 | high | protein_cod | *** | Universal st | usp domain | no mercato | GO:000695 | PF00582 | IPR006016 |
| 176 | HORVU.MOREX.r3.2HG01140 | low | non_coding | --* | 50S ribosom | sh domain c | no mercato | GO:000373 | NA | NA |
| 177 | HORVU.MOREX.r3.2HG01152 | high | protein_cod | *** | Tubby-like F | stress-resp | external stir | GO:003509 | PF01167,PF | IPR000007,IPR001810,IPR001810,IPR025659 |
| 178 | HORVU.MOREX.r3.2HG01158 | low | protein_cod | --* | PHD finger t | not classifie | no mercato | NA | NA | NA |
| 179 | HORVU.MOREX.r3.2HG01158 | high | protein_cod | --* | Mannose-bi | not classifie | no mercato | NA | NA | NA |
| 180 | HORVU.MOREX.r3.2HG01163 | high | protein_cod | *** | WD40 repea | anapc4 wd4 | no mercato | NA | PF00400,PF | IPR017986,IPR001680,IPR001680,IPR001680,IPR001680,IPR001680,IPR001680,IPR00168 |
| 181 | HORVU.MOREX.r3.2HG01164 | high | protein_cod | *** | Early respor | protein early | no mercato | NA | NA | NA |
| 182 | HORVU.MOREX.r3.2HG01172 | high | protein_cod | *** | Transmemb | trna m 4 x o | no mercato | GO:001602 | NA | NA |
| 183 | HORVU.MOREX.r3.2HG01173 | high | protein_cod | *** | Glycosyltra | xylosyltrans | cell wall org | GO:001602 | PF04577 | IPR007657 |
| 184 | HORVU.MOREX.r3.2HG01185 | high | protein_cod | *** | Glycosyltra | EC_2.4 glyc | enzyme clas | GO:000815 | PF00201 | IPR002213 |
| 185 | HORVU.MOREX.r3.2HG01187 | low | non_coding | *** | Cytochrome | brassinoste | phytohormo | GO:000449 | PF00067 | IPR001128,IPR001128 |
| 186 | HORVU.MOREX.r3.2HG01196 | high | protein_cod | *** | Transcriptio | component | rna biosynth | GO:000374 | PF08711 | IPR017923,IPR017923 |
| 187 | HORVU.MOREX.r3.2HG01202 | high | protein_cod | *** | Pyrrolidone | pyrrolidone- | protein moc | GO:000582 | PF01470 | IPR016125 |
| 188 | HORVU.MOREX.r3.2HG01208 | high | protein_cod | *** | tolB protein | dppiv n dom | no mercato | NA | PF07676,PF | IPR011659,IPR011659,IPR011659 |
| 189 | HORVU.MOREX.r3.2HG01213 | low | non_coding | *** | Cytochrome | EC_1.14 oxi | enzyme clas | GO:000449 | PF00067 | IPR001128,IPR001128 |
| 190 | HORVU.MOREX.r3.2HG01214 | low | non_coding | *** | Cytochrome | EC_1.14 oxi | enzyme clas | GO:000449 | PF00067 | IPR001128,IPR001128 |
| 191 | HORVU.MOREX.r3.2HG01216 | high | protein_cod | --* | Ribosome-r | not classifie | no mercato | GO:000016 | NA | NA |
| 192 | HORVU.MOREX.r3.2HG01221 | high | protein_cod | *-* | RING/U-box | e3 ubiquitin | no mercato | NA | PF13639 | IPR001841 |
| 193 | HORVU.MOREX.r3.2HG01223 | high | protein_cod | *** | Ribonuclea | T2-type RNa | rna process | GO:000372 | PF00445 | IPR001568,IPR001568 |
| 194 | HORVU.MOREX.r3.2HG01224 | high | protein_cod | *-* | Myb factor | MYB class-F | rna biosynth | GO:000367 | PF00249,PF | IPR001005,IPR001005,IPR009057 |

| | | | | | | | | | | |
|---|---|---|---|---|---|---|---|---|---|---|
| 195 | HORVU.MOREX.r3.2HG012252 | low | protein_cod | --* | SAUR-like a | glyco hydro | no mercator | GO:000973 | NA | NA |
| 196 | HORVU.MOREX.r3.2HG012335 | high | protein_cod | *** | Trehalose 6 | trehalose-6- | carbohydrat | GO:000382 | PF02358 | IPR003337,IPR003337,IPR023214,IPR006379 |
| 197 | HORVU.MOREX.r3.2HG012445 | high | protein_cod | *** | calmodulin- | c2h2 and c2 | no mercator | NA | PF07816 | IPR012442 |
| 198 | HORVU.MOREX.r3.2HG012551 | high | protein_cod | *** | SPX domain | regulatory p | nutrient upt | GO:001603 | PF03105 | IPR004331 |
| 199 | HORVU.MOREX.r3.2HG012584 | high | protein_cod | *** | Kinase fami | RLCK-VIIa re | protein moc | GO:000467 | PF07714 | IPR011009,IPR001245 |
| 200 | HORVU.MOREX.r3.2HG012641 | high | protein_cod | *-* | Nuclear trar | component | rna biosynth | GO:000563 | PF00808 | IPR003958,IPR009072 |
| 201 | HORVU.MOREX.r3.2HG012645 | high | protein_cod | *** | Sulfotransfe | EC_2.8 tran | enzyme clas | NA | PF00685 | IPR027417,IPR000863 |
| 202 | HORVU.MOREX.r3.2HG012646 | high | protein_cod | *** | Sulfotransfe | EC_2.8 tran | enzyme clas | NA | PF00685 | IPR027417,IPR000863 |
| 203 | HORVU.MOREX.r3.2HG012688 | high | protein_cod | *-* | Ninja-family | transcriptio | rna biosynth | GO:000716 | PF16136,PF | IPR032310,IPR032308,IPR012463 |
| 204 | HORVU.MOREX.r3.2HG012733 | high | protein_cod | *** | B12D protei | nadh ubiqui | no mercator | GO:001602 | PF06522 | IPR010530 |
| 205 | HORVU.MOREX.r3.2HG012742 | high | protein_cod | *** | Dihydroflav | bifunctional | no mercator | GO:000382 | PF01370 | IPR016040,IPR001509 |
| 206 | HORVU.MOREX.r3.2HG012860 | high | protein_cod | *** | Receptor kir | LRR-Xb prot | protein moc | GO:000016 | PF00069,PF | IPR032675,IPR000719,IPR001611,IPR032675,IPR013210,IPR011009 |
| 207 | HORVU.MOREX.r3.2HG012902 | high | protein_cod | *** | Short-chain | short chain | no mercator | NA | IPR016040 | |
| 208 | HORVU.MOREX.r3.2HG013021 | high | protein_cod | *** | ZCF37, puta | not classifie | no mercator | GO:001602 | NA | NA |
| 209 | HORVU.MOREX.r3.2HG013024 | high | protein_cod | *** | Transferase | hydroxycinn | cell wall org | GO:001674 | PF02458 | IPR003480 |
| 210 | HORVU.MOREX.r3.2HG013059 | high | protein_cod | *-* | Kinase fami | EC_2.7 tran | enzyme clas | GO:000016 | PF00069 | IPR011009,IPR000719 |
| 211 | HORVU.MOREX.r3.2HG013074 | high | protein_cod | *** | Kinase fami | MAP3K-WNI | multi-proce | GO:000467 | PF00069 | IPR000719,IPR011009 |
| 212 | HORVU.MOREX.r3.2HG013243 | high | protein_cod | --* | Transcriptio | protein free | no mercator | NA | NA | NA |
| 213 | HORVU.MOREX.r3.2HG013345 | high | protein_cod | *** | Remorin | remorin c dc | no mercator | NA | PF03763 | IPR005516 |
| 214 | HORVU.MOREX.r3.2HG013494 | high | protein_cod | *-* | Heat shock | HSF transcr | rna biosynth | GO:000367 | PF00447 | IPR011991,IPR000232 |
| 215 | HORVU.MOREX.r3.2HG013512 | high | protein_cod | *** | F-box protei | substrate ac | protein hom | NA | NA | NA |
| 216 | HORVU.MOREX.r3.2HG013531 | high | protein_cod | *** | Sugar trans | monosacch | solute trans | GO:000521 | PF00083 | IPR005828,IPR003663,IPR020846 |
| 217 | HORVU.MOREX.r3.2HG013590 | high | protein_cod | *** | Equilibrative | adenosine p | nucleotide r | GO:000533 | PF01733 | IPR020846,IPR002259 |
| 218 | HORVU.MOREX.r3.2HG013629 | high | protein_cod | *** | Cyclin famil | NA | NA | GO:000007 | PF08613 | IPR013922 |
| 219 | HORVU.MOREX.r3.2HG013640 | high | protein_cod | *-* | Plant cadmi | protein plan | no mercator | GO:001602 | PF04749 | IPR006461,IPR006461 |
| 220 | HORVU.MOREX.r3.2HG013832 | high | protein_cod | --* | Dihydroxy-a | acidic chitin | external stir | GO:000382 | PF00704 | IPR017853,IPR001223 |
| 221 | HORVU.MOREX.r3.2HG013834 | high | protein_cod | *** | Xylanase inl | acidic chitin | external stir | GO:000597 | PF00704 | IPR001223,IPR017853 |
| 222 | HORVU.MOREX.r3.2HG013873 | high | protein_cod | *** | Receptor-lik | DUF26 prote | protein moc | GO:000467 | PF01657,PF | IPR002902,IPR002902,IPR000719,IPR011009 |
| 223 | HORVU.MOREX.r3.2HG013934 | high | protein_cod | *** | Protein tran | assembly fa | protein bios | GO:000374 | PF01253 | IPR001950,IPR001950,IPR005874 |
| 224 | HORVU.MOREX.r3.2HG014017 | high | protein_cod | *** | Ras-related | ras protein r | no mercator | GO:000392 | PF00071 | IPR027417,IPR001806 |
| 225 | HORVU.MOREX.r3.2HG014029 | high | protein_cod | *** | Glucose-6-p | phosphome | solute trans | GO:001602 | PF03151 | IPR004696,IPR004853 |
| 226 | HORVU.MOREX.r3.2HG014119 | high | protein_cod | *** | senescence | senescence | no mercator | NA | PF04520 | IPR007608 |
| 227 | HORVU.MOREX.r3.2HG014162 | high | protein_cod | *** | Autophagy-r | autophagos | protein hom | GO:000691 | PF02991 | IPR029071,IPR004241 |
| 228 | HORVU.MOREX.r3.2HG014174 | high | protein_cod | *** | DNA/RNA h | chromatin r | chromatin o | GO:000367 | PF00176,PF | IPR027417,IPR027417,IPR027417,IPR000330,IPR001650,IPR027417,IPR027417 |
| 229 | HORVU.MOREX.r3.2HG014203 | high | protein_cod | *** | Phosphoade | (phospho)a | nutrient upt | GO:000382 | PF00085,PF | IPR004508,IPR013766,IPR012336,IPR002500 |
| 230 | HORVU.MOREX.r3.2HG014515 | high | protein_cod | *** | Receptor-lik | CIF/TWS1-p | phytohormo | GO:000016 | PF00069,PF | IPR000719,IPR011009,IPR032675,IPR032675,IPR032675,IPR013210,IPR001611,IPR001611,IPR001611,IPR00161 |
| 231 | HORVU.MOREX.r3.2HG015218 | high | protein_cod | *-* | GCK domair | gck domain | no mercator | NA | PF07802 | IPR012891 |
| 232 | HORVU.MOREX.r3.2HG015233 | high | protein_cod | *** | Receptor-lik | URK-1 prote | protein moc | GO:000467 | PF07714 | IPR001245,IPR011009 |
| 233 | HORVU.MOREX.r3.2HG015261 | high | protein_cod | *-* | Glutaredoxi | glutaredoxir | no mercator | GO:000562 | PF00462 | IPR002109,IPR012336 |
| 234 | HORVU.MOREX.r3.2HG015297 | high | protein_cod | *** | Transporter | inositol tran | solute trans | GO:000521 | PF00083,PF | IPR005828,IPR005828,IPR020846,IPR020846,IPR003663 |
| 235 | HORVU.MOREX.r3.2HG015323 | high | protein_cod | *** | Non-specifi | SnRK3 SNF1 | protein moc | NA | PF00069,PF | IPR000719,IPR004041,IPR011009 |
| 236 | HORVU.MOREX.r3.2HG015721 | high | protein_cod | *** | Sugar trans | monosacch | solute trans | GO:000521 | PF00083 | IPR005828,IPR020846,IPR020846,IPR003663 |
| 237 | HORVU.MOREX.r3.2HG015869 | high | protein_cod | *** | WRKY trans | transcriptio | rna biosynth | GO:000367 | PF03106,PF | IPR003657,IPR018872,IPR003657 |
| 238 | HORVU.MOREX.r3.2HG015910 | high | protein_cod | *** | Core-2/I-bra | beta glucurc | no mercator | GO:000837 | PF02485 | IPR003406 |
| 239 | HORVU.MOREX.r3.2HG016013 | high | protein_cod | *-* | Basic helix-l | bHLH class- | rna biosynth | GO:004698 | PF00010 | IPR011598,IPR011598 |
| 240 | HORVU.MOREX.r3.2HG016056 | high | protein_cod | *** | Peptidylprol | peptidyl-pro | protein moc | GO:000041 | PF00515,PF | IPR001440,IPR001179,IPR001179,IPR001179,IPR011990 |
| 241 | HORVU.MOREX.r3.2HG016079 | high | protein_cod | --* | Alcohol deh | plastidial ox | redox home | GO:000402 | PF08240 | IPR016040,IPR011032,IPR013154 |
| 242 | HORVU.MOREX.r3.2HG016094 | high | protein_cod | *** | LURP-one-li | lurp domain | no mercator | NA | PF04525 | IPR007612,IPR025659 |
| 243 | HORVU.MOREX.r3.2HG016232 | high | protein_cod | *** | F-box protei | mynd type d | no mercator | NA | PF01753 | IPR002893,IPR001810 |
| 244 | HORVU.MOREX.r3.2HG016293 | high | protein_cod | *** | Nodulin-like | UMF23-type | solute trans | GO:001602 | PF06813 | IPR010658,IPR020846,IPR020846,IPR020846,IPR020846 |
| 245 | HORVU.MOREX.r3.2HG016374 | high | protein_cod | *** | Glycerophos | glycerophos | no mercator | GO:000662 | PF03009 | IPR030395,IPR017946,IPR017946 |
| 246 | HORVU.MOREX.r3.2HG016375 | high | protein_cod | *** | Hydroxyprol | bar domain | no mercator | NA | NA | NA |
| 247 | HORVU.MOREX.r3.2HG016388 | high | protein_cod | --* | Jasmonate z | co-regulator | rna biosynth | NA | PF09425,PF | IPR018467,IPR010399 |
| 248 | HORVU.MOREX.r3.2HG016416 | high | protein_cod | *-* | Trihelix tran | TRIHELIX tra | rna biosynth | NA | NA | NA |
| 249 | HORVU.MOREX.r3.2HG016434 | high | protein_cod | *** | Ethylene-res | subgroup EF | rna biosynth | GO:000367 | PF00847 | IPR016177,IPR001471 |
| 250 | HORVU.MOREX.r3.2HG016447 | high | protein_cod | *** | Nudix hydro | nudix hydrol | no mercator | GO:001678 | PF00293 | IPR015797,IPR000086 |
| 251 | HORVU.MOREX.r3.2HG016470 | high | protein_cod | *** | Potassium t | potassium c | solute trans | GO:000681 | PF02705 | IPR003855,IPR003855 |
| 252 | HORVU.MOREX.r3.2HG016520 | high | protein_cod | *** | Short-chain | short chain | no mercator | GO:001649 | IPR016040 | |
| 253 | HORVU.MOREX.r3.2HG016537 | high | protein_cod | --* | Target of rap | genome ass | no mercator | GO:000554 | NA | NA |
| 254 | HORVU.MOREX.r3.2HG016545 | high | protein_cod | *** | Arogenate d | arogenate d | amino acid | GO:000466 | PF00800 | IPR001086 |
| 255 | HORVU.MOREX.r3.2HG016706 | high | protein_cod | *** | Protease inl | aai domain | no mercator | GO:000650 | PF14368 | IPR016140,IPR016140 |
| 256 | HORVU.MOREX.r3.2HG016735 | high | protein_cod | *** | U-box doma | E3 ubiquitin | protein hom | GO:000484 | PF04564 | IPR016024,IPR003613 |
| 257 | HORVU.MOREX.r3.2HG016856 | high | protein_cod | *** | Late embryo | microtubule | cell wall org | GO:001602 | PF03168 | IPR004864 |
| 258 | HORVU.MOREX.r3.2HG016858 | high | protein_cod | *** | Thioredoxin | thioredoxin | photosynthe | GO:000562 | PF00085 | IPR013766,IPR005746,IPR012336 |
| 259 | HORVU.MOREX.r3.2HG016877 | high | protein_cod | *** | Scarecrow t | GRAS transc | rna biosynth | GO:000370 | PF03514 | IPR005202 |

| | | | | | | | | | | |
|---|---|---|---|---|---|---|---|---|---|---|
| 260 | HORVU.MOREX.r3.2HG016879 | high | protein_cod | *** | Haloacid de | phosphosug | carbohydrat | GO:000815 | PF13419 | IPR023214,IPR023214,IPR006439 |
| 261 | HORVU.MOREX.r3.2HG016988 | high | protein_cod | *** | T-complex p | t complex p | no mercator | NA | PF05794 | IPR008862 |
| 262 | HORVU.MOREX.r3.2HG017057 | high | protein_cod | *** | AGAMOUS-l | MADS/AGL t | rna biosynth | GO:000097 | PF01486,PF | IPR002487,IPR002100,IPR002100 |
| 263 | HORVU.MOREX.r3.2HG017058 | high | protein_cod | *** | 12-oxophyt | 4,5-didehyd | phytohormo | GO:000382 | PF00724 | IPR001155 |
| 264 | HORVU.MOREX.r3.2HG017117 | high | protein_cod | *** | F-box protei | f box domai | no mercator | NA | PF12937 | IPR001810,IPR001810 |
| 265 | HORVU.MOREX.r3.2HG017174 | high | protein_cod | *** | Reductase 1 | EC_1.1 oxid | enzyme clas | GO:001649 | PF00248 | IPR023210,IPR023210 |
| 266 | HORVU.MOREX.r3.2HG017201 | high | protein_cod | *** | transmemb | gag 3 domai | no mercator | NA | NA | NA |
| 267 | HORVU.MOREX.r3.2HG017208 | high | protein_cod | --* | RING/FYVE/ | not classifie | no mercator | NA | NA | NA |
| 268 | HORVU.MOREX.r3.2HG017211 | high | protein_cod | *** | Dihydrofola | fsh domain | no mercator | NA | PF03959 | IPR005645,IPR029058 |
| 269 | HORVU.MOREX.r3.2HG017214 | high | protein_cod | *** | RING/U-box | RING-H2-cl | protein hom | GO:001602 | PF13639 | IPR001841 |
| 270 | HORVU.MOREX.r3.2HG017218 | high | protein_cod | *-* | Ring finger p | RING-H2-cl | protein hom | NA | PF13639 | IPR001841 |
| 271 | HORVU.MOREX.r3.2HG017340 | high | protein_cod | *** | NAC domain | NAC transcr | rna biosynth | GO:000367 | PF02365 | IPR003441,IPR003441 |
| 272 | HORVU.MOREX.r3.2HG017405 | high | protein_cod | *** | 1-deoxy-D-x | aldedh dom | no mercator | GO:001602 | NA | NA |
| 273 | HORVU.MOREX.r3.2HG017465 | high | protein_cod | *** | WPP domai | outer nuclea | cytoskeleto | GO:001602 | NA | NA |
| 274 | HORVU.MOREX.r3.2HG017518 | high | protein_cod | *** | 2-oxoglutar | indole-3-ac | phytohormo | GO:001649 | PF03171,PF | IPR005123,IPR026992 |
| 275 | HORVU.MOREX.r3.2HG017538 | high | protein_cod | *** | Sec14p-like | phosphoino | multi-proce | NA | PF00650,PF | IPR001251,IPR001251,IPR011074,IPR011074 |
| 276 | HORVU.MOREX.r3.2HG017558 | high | protein_cod | *** | Aldo/keto re | EC_1.1 oxid | enzyme clas | NA | PF00248 | IPR023210,IPR023210 |
| 277 | HORVU.MOREX.r3.2HG017604 | high | protein_cod | *** | Transducin/ | protein jing | no mercator | NA | PF00400,PF | IPR017986,IPR017986,IPR001680,IPR001680,IPR001680,IPR001680,IPR00168 |
| 278 | HORVU.MOREX.r3.2HG017618 | high | protein_cod | *** | Peptide met | methionine | redox home | GO:000811 | PF01625 | IPR002569,IPR002569,IPR002569 |
| 279 | HORVU.MOREX.r3.2HG017625 | high | protein_cod | *** | BTB/POZ an | substrate a | protein hom | GO:000371 | PF00651,PF | IPR000197,IPR000210,IPR011333,IPR011333,IPR000197 |
| 280 | HORVU.MOREX.r3.2HG017642 | high | protein_cod | *** | Bag family n | ubiquitin do | no mercator | NA | PF00240 | IPR000626,IPR029071 |
| 281 | HORVU.MOREX.r3.2HG017644 | high | protein_cod | *** | Bag family n | ubiquitin do | no mercator | NA | PF00240 | IPR029071,IPR000626 |
| 282 | HORVU.MOREX.r3.2HG017732 | high | protein_cod | *** | Kinase fami | AGC-VIIIb/p | protein mod | GO:000016 | PF00069,PF | IPR000719,IPR000719,IPR011009,IPR01100 |
| 283 | HORVU.MOREX.r3.2HG017783 | high | protein_cod | *** | Transporter | inositol tran | solute trans | GO:000521 | PF00083 | IPR020846,IPR005828,IPR003663 |
| 284 | HORVU.MOREX.r3.2HG017802 | high | protein_cod | *** | Chitinase | acidic endo | no mercator | GO:000456 | PF00187,PF | IPR001002,IPR000726,IPR000726,IPR023346,IPR001002 |
| 285 | HORVU.MOREX.r3.2HG017806 | high | protein_cod | *** | transmemb | duf 642 don | no mercator | NA | PF04862,PF | IPR006946,IPR006946,IPR008979 |
| 286 | HORVU.MOREX.r3.2HG017932 | high | protein_cod | *** | BTB/POZ do | substrate a | protein hom | GO:005126 | PF02214 | IPR011044,IPR011044,IPR011044,IPR011333,IPR003131 |
| 287 | HORVU.MOREX.r3.2HG017964 | high | protein_cod | *** | Leucine-rich | dhhc domai | no mercator | NA | PF13855,PF | IPR001611,IPR032675,IPR001611,IPR001611,IPR001611,IPR001611,IPR00161 |
| 288 | HORVU.MOREX.r3.2HG018050 | high | protein_cod | *** | Glutaredoxi | glutaredoxi | protein mod | GO:000562 | PF00462 | IPR011899,IPR012336,IPR002109 |
| 289 | HORVU.MOREX.r3.2HG018068 | high | protein_cod | *** | Zinc finger ( | anaphase p | no mercator | NA | PF14624,PF | IPR032838,IPR002035,IPR001841,IPR002035 |
| 290 | HORVU.MOREX.r3.2HG018082 | high | protein_cod | *** | Ammonium | ammonium | solute trans | GO:000851 | PF00909 | IPR001905,IPR024041,IPR024041 |
| 291 | HORVU.MOREX.r3.2HG018240 | high | protein_cod | *** | Aquaporin | plasma mer | solute trans | GO:001526 | PF00230 | IPR000425,IPR023271,IPR000425 |
| 292 | HORVU.MOREX.r3.2HG018252 | low | plastid_rela | *** | Oxygen-evo | zinc ribbon | no mercator | GO:000550 | PF05757 | IPR023222,IPR008797 |
| 293 | HORVU.MOREX.r3.2HG018273 | high | protein_cod | *** | Glycosyltra | EC_2.4 glyc | enzyme clas | GO:000815 | PF00201 | IPR002213 |
| 294 | HORVU.MOREX.r3.2HG018281 | high | protein_cod | *** | Yellow strip | iron chelato | solute trans | GO:001602 | PF03169 | IPR004813,IPR004813 |
| 295 | HORVU.MOREX.r3.2HG018284 | high | protein_cod | *** | Yellow strip | iron chelato | solute trans | GO:001602 | PF03169 | IPR004813,IPR004813 |
| 296 | HORVU.MOREX.r3.2HG018299 | high | protein_cod | *** | UDP-glycos | EC_2.4 glyc | enzyme clas | GO:000815 | PF00201 | IPR002213 |
| 297 | HORVU.MOREX.r3.2HG018338 | high | protein_cod | *-* | GRAM doma | gram domai | no mercator | NA | PF02893 | IPR004182 |
| 298 | HORVU.MOREX.r3.2HG018383 | high | protein_cod | *** | E3 ubiquitin | E3 ubiquitin | protein hom | GO:001687 | PF00097 | IPR018957 |
| 299 | HORVU.MOREX.r3.2HG018393 | high | protein_cod | *** | Lectin recep | L-Lectin pro | protein mod | GO:000467 | PF00069,PF | IPR013320,IPR011009,IPR000719,IPR00122 |
| 300 | HORVU.MOREX.r3.2HG018394 | high | protein_cod | *** | Lectin recep | L-Lectin pro | protein mod | GO:000467 | PF00139,PF | IPR001220,IPR011009,IPR000719,IPR01332 |
| 301 | HORVU.MOREX.r3.2HG018402 | high | protein_cod | *** | Short-chain | EC_1.1 oxid | enzyme clas | NA | PF00106 | IPR016040,IPR016040,IPR002347 |
| 302 | HORVU.MOREX.r3.2HG018408 | high | protein_cod | *** | MYB transcr | MYB class-F | rna biosynth | GO:000367 | PF00249,PF | IPR009057,IPR001005,IPR001005 |
| 303 | HORVU.MOREX.r3.2HG018413 | high | protein_cod | *** | MYB transcr | MYB class-F | rna biosynth | GO:000367 | PF00249,PF | IPR001005,IPR001005,IPR009057 |
| 304 | HORVU.MOREX.r3.2HG018455 | high | protein_cod | *** | SAUR-like a | auxin respo | no mercator | GO:000973 | PF02519 | IPR003676 |
| 305 | HORVU.MOREX.r3.2HG018484 | high | protein_cod | *** | GATA transc | A/B-GATA-ty | rna biosynth | GO:000367 | PF00320 | IPR000679 |
| 306 | HORVU.MOREX.r3.2HG018489 | high | protein_cod | *** | SH3 domain | auxiliary reg | vesicle traff | NA | PF14604 | IPR001452,IPR001452 |
| 307 | HORVU.MOREX.r3.2HG018493 | high | protein_cod | *** | Zinc finger, I | BBX class-I\ | rna biosynth | GO:000562 | PF00643,PF | IPR000315,IPR000315 |
| 308 | HORVU.MOREX.r3.2HG018532 | high | protein_cod | *** | Trihelix tran | TRIHELIX tra | rna biosynth | NA | NA | NA |
| 309 | HORVU.MOREX.r3.2HG018588 | high | protein_cod | --* | Purine nucle | regulatory p | multi-proce | GO:000382 | NA | NA |
| 310 | HORVU.MOREX.r3.2HG018615 | high | protein_cod | *** | 2-oxoglutar | type-I flavor | secondary r | GO:001649 | PF03171,PF | IPR005123,IPR026992 |
| 311 | HORVU.MOREX.r3.2HG018745 | high | protein_cod | *** | Rhomboid-l | Rhomboid-t | protein hom | GO:000425 | PF01694 | IPR022764 |
| 312 | HORVU.MOREX.r3.2HG018879 | high | protein_cod | *** | Glycosyltra | zeatin O-glu | phytohormo | GO:000815 | PF00201 | IPR002213 |
| 313 | HORVU.MOREX.r3.2HG018885 | high | protein_cod | *** | Glutathione | glutathione | redox home | NA | PF00255 | IPR012336,IPR000889 |
| 314 | HORVU.MOREX.r3.2HG018901 | high | protein_cod | *** | Glycerol-3-p | organic pho | solute trans | GO:001602 | PF07690 | IPR011701,IPR020846 |
| 315 | HORVU.MOREX.r3.2HG018944 | high | protein_cod | *** | Cyclin famil | regulatory p | cell division | GO:000007 | PF08613 | IPR013763,IPR013922 |
| 316 | HORVU.MOREX.r3.2HG019021 | high | protein_cod | *** | Oleosin | Oleosin-typ | lipid metab | GO:000581 | PF01277 | IPR000136 |
| 317 | HORVU.MOREX.r3.2HG019026 | high | protein_cod | *** | SHR5-recep | LRR-VIII-2 p | protein mod | GO:000467 | PF00069,PF | IPR000719,IPR032675,IPR021720,IPR01100 |
| 318 | HORVU.MOREX.r3.2HG019062 | high | protein_cod | --* | Ceramide ki | not classifie | no mercator | GO:000395 | NA | NA |
| 319 | HORVU.MOREX.r3.2HG019063 | high | protein_cod | *** | Lecithin:ch | lecithin cho | no mercator | GO:000662 | PF02450 | IPR003386,IPR029058,IPR029058,IPR02905 |
| 320 | HORVU.MOREX.r3.2HG019089 | high | protein_cod | *-* | Armadillo/b | cbm domain | no mercator | GO:000563 | IPR016024 | |
| 321 | HORVU.MOREX.r3.2HG019144 | high | protein_cod | *** | Vacuolar so | lytic vacuole | vesicle traff | GO:000550 | PF02225,PF | IPR003137,IPR026823 |
| 322 | HORVU.MOREX.r3.2HG019149 | high | protein_cod | *** | Protein kina | MAP3K-RAF | multi-proce | GO:000467 | PF07714 | IPR011009,IPR001245 |
| 323 | HORVU.MOREX.r3.2HG019197 | high | protein_cod | *** | Protein pho | clade F pho | protein mod | GO:000382 | PF00481 | IPR001932,IPR001932 |
| 324 | HORVU.MOREX.r3.2HG019284 | high | protein_cod | *** | Leucine-rich | plant intrac | no mercator | NA | PF00560,PF | IPR001611,IPR032675,IPR001611 |

| | | | | | | | | | | |
|---|---|---|---|---|---|---|---|---|---|---|
| 325 | HORVU.MOREX.r3.2HG01928 | high | protein_cod | *** | WRKY trans | transcriptio | rna biosynth | GO:000367 | PF10533,PF | IPR018872,IPR003657,IPR003657 |
| 326 | HORVU.MOREX.r3.2HG01932 | high | protein_cod | *-* | Trihelix tran | TRIHELIX tra | rna biosynth | NA | NA | NA |
| 327 | HORVU.MOREX.r3.2HG01936 | high | protein_cod | *** | Ubiquinol o | alternative c | cellular resp | GO:000991 | PF01786 | IPR002680 |
| 328 | HORVU.MOREX.r3.2HG01936 | high | protein_cod | *** | Ubiquinol o | alternative c | cellular resp | GO:000991 | PF01786 | IPR002680 |
| 329 | HORVU.MOREX.r3.2HG01936 | high | protein_cod | *** | Ubiquinol o | alternative c | cellular resp | GO:000991 | PF01786 | IPR002680 |
| 330 | HORVU.MOREX.r3.2HG01937 | high | protein_cod | *** | Hypoxia-res | component | cellular resp | GO:001602 | PF04588 | IPR007667 |
| 331 | HORVU.MOREX.r3.2HG01942 | high | protein_cod | *** | Seed specif | late embryo | no mercator | NA | NA | NA |
| 332 | HORVU.MOREX.r3.2HG01945 | high | protein_cod | *** | Vacuolar ca | cation antip | solute trans | GO:000681 | PF01699,PF | IPR004713,IPR004798,IPR004837,IPR004837 |
| 333 | HORVU.MOREX.r3.2HG01946 | high | protein_cod | *** | Autophagy-r | autophagos | protein hom | GO:000691 | PF02991 | IPR004241,IPR029071 |
| 334 | HORVU.MOREX.r3.2HG01948 | high | protein_cod | *-* | Cysteine/Hi | nucleoredo | redox home | NA | PF13905,PF | IPR012336,IPR012336,IPR012336,IPR004146,IPR012336,IPR012336,IPR01233 |
| 335 | HORVU.MOREX.r3.2HG01952 | high | protein_cod | *** | Receptor-lik | cold-respon | external stir | GO:000016 | PF07714 | IPR011009,IPR001245 |
| 336 | HORVU.MOREX.r3.2HG01954 | low | protein_cod | --* | Eukaryotic t | not classifie | no mercator | GO:000173 | NA | NA |
| 337 | HORVU.MOREX.r3.2HG01959 | high | protein_cod | *-* | Calcium-de | c2 domain c | no mercator | NA | PF00168 | IPR000008,IPR000008 |
| 338 | HORVU.MOREX.r3.2HG01959 | high | protein_cod | *** | F-box protei | f box domai | no mercator | NA | PF00646,PF | IPR001810,IPR025886,IPR001810 |
| 339 | HORVU.MOREX.r3.2HG01961 | high | protein_cod | *** | Lectin recep | EC_2.7 tran | enzyme clas | GO:000467 | PF00139,PF | IPR013320,IPR011009,IPR001220,IPR00071 |
| 340 | HORVU.MOREX.r3.2HG01963 | high | protein_cod | *** | senescence | regulatory p | multi-proce | NA | PF04570 | IPR007650 |
| 341 | HORVU.MOREX.r3.2HG01964 | high | protein_cod | *-* | senescence | regulatory p | multi-proce | NA | PF04570 | IPR007650 |
| 342 | HORVU.MOREX.r3.2HG01965 | high | protein_cod | *-* | senescence | regulatory p | multi-proce | NA | PF04570 | IPR007650 |
| 343 | HORVU.MOREX.r3.2HG01965 | high | protein_cod | *-* | senescence | regulatory p | multi-proce | NA | PF04570 | IPR007650 |
| 344 | HORVU.MOREX.r3.2HG01969 | high | protein_cod | *** | U-box doma | E3 ubiquitin | protein hom | GO:000484 | PF04564 | IPR003613,IPR016024 |
| 345 | HORVU.MOREX.r3.2HG01972 | high | protein_cod | --* | disease resi | not classifie | no mercator | NA | NA | NA |
| 346 | HORVU.MOREX.r3.2HG01973 | high | protein_cod | *** | S-norcoclau | bet v 1 dom | no mercator | GO:000695 | PF00407 | IPR000916 |
| 347 | HORVU.MOREX.r3.2HG01983 | high | protein_cod | *** | Protein tran | assembly fa | protein bios | GO:000374 | PF01253 | IPR001950,IPR001950 |
| 348 | HORVU.MOREX.r3.2HG01985 | high | protein_cod | --* | Matrilin-3 | not classifie | no mercator | GO:000150 | NA | NA |
| 349 | HORVU.MOREX.r3.2HG02003 | high | protein_cod | *** | Beta-fructof | cell wall aci | carbohydrat | GO:000455 | PF00251,PF | IPR013148,IPR013189,IPR013320,IPR013320,IPR02329 |
| 350 | HORVU.MOREX.r3.2HG02003 | high | protein_cod | *** | Beta-fructof | cell wall aci | carbohydrat | GO:000455 | PF08244,PF | IPR023296,IPR013189,IPR013320,IPR013320,IPR01314 |
| 351 | HORVU.MOREX.r3.2HG02007 | low | non_coding | *** | Cytochrome | ferulate 5-h | cell wall org | GO:000449 | PF00067 | IPR001128,IPR001128 |
| 352 | HORVU.MOREX.r3.2HG02016 | high | protein_cod | *** | SAUR-like a | auxin respo | no mercator | GO:000973 | PF02519 | IPR003676 |
| 353 | HORVU.MOREX.r3.2HG02021 | high | protein_cod | *** | Transcriptio | bHLH trans | rna biosynth | NA | NA | NA |
| 354 | HORVU.MOREX.r3.2HG02022 | high | protein_cod | *** | Kelch repea | substrate ac | external stir | NA | PF01344,PF | IPR006652,IPR001810,IPR001810 |
| 355 | HORVU.MOREX.r3.2HG02028 | high | protein_cod | --* | PRLI-interac | not classifie | no mercator | NA | NA | NA |
| 356 | HORVU.MOREX.r3.2HG02030 | low | protein_cod | NA | NA | glycine rich | no mercator | NA | NA | NA |
| 357 | HORVU.MOREX.r3.2HG02032 | high | protein_cod | *** | Hydroxycinn | EC_2.3 acyl | enzyme clas | GO:001674 | PF02458 | IPR003480 |
| 358 | HORVU.MOREX.r3.2HG02032 | high | protein_cod | --* | quiescin-su | ctenidin | no mercator | NA | NA | NA |
| 359 | HORVU.MOREX.r3.2HG02038 | high | protein_cod | *** | Nuclease S | bifunctional | chromatin o | GO:000367 | PF02265 | IPR003154,IPR008947 |
| 360 | HORVU.MOREX.r3.2HG02045 | high | protein_cod | *** | Cysteine pr | Papain-type | protein hom | GO:000650 | PF00396,PF | IPR000118,IPR000668,IPR013201 |
| 361 | HORVU.MOREX.r3.2HG02048 | high | protein_cod | *** | Ethylene-re | subgroup EF | rna biosynth | GO:000367 | PF00847 | IPR001471,IPR016177 |
| 362 | HORVU.MOREX.r3.2HG02069 | high | protein_cod | *** | transcriptio | TGA-type tra | rna biosynth | NA | PF00170,PF | IPR004827,IPR025422 |
| 363 | HORVU.MOREX.r3.2HG02074 | high | protein_cod | *** | Wound-resp | not classifie | no mercator | NA | PF12609 | IPR022251 |
| 364 | HORVU.MOREX.r3.2HG02075 | high | protein_cod | *** | Wound-resp | not classifie | no mercator | NA | PF12609 | IPR022251 |
| 365 | HORVU.MOREX.r3.2HG02093 | high | protein_cod | *** | Receptor-lik | EC_2.7 tran | enzyme clas | GO:000016 | PF00069 | IPR000719,IPR011009 |
| 366 | HORVU.MOREX.r3.2HG02095 | high | protein_cod | *** | Dof-like zinc | DOF transcr | rna biosynth | GO:000367 | PF02701 | IPR003851 |
| 367 | HORVU.MOREX.r3.2HG02099 | high | protein_cod | *** | Late embryo | lea 2 domai | no mercator | GO:000573 | PF03168 | IPR004864 |
| 368 | HORVU.MOREX.r3.2HG02100 | high | protein_cod | *** | Transcriptio | MYBR-R-typ | rna biosynth | GO:000367 | PF00249 | IPR009057,IPR006447,IPR009057,IPR001005 |
| 369 | HORVU.MOREX.r3.2HG02106 | high | protein_cod | *** | Acetyl-coen | EC_6.2 ligas | enzyme clas | GO:000016 | PF00501,PF | IPR000873,IPR025110 |
| 370 | HORVU.MOREX.r3.2HG02107 | high | protein_cod | --* | Chromatin s | not classifie | no mercator | NA | NA | NA |
| 371 | HORVU.MOREX.r3.2HG02119 | low | non_coding | *** | Cytochrome | EC_1.14 oxi | enzyme clas | GO:000449 | PF00067 | IPR001128,IPR001128 |
| 372 | HORVU.MOREX.r3.2HG02129 | high | protein_cod | *** | Acyl-protein | protein dea | protein mod | GO:001678 | PF02230 | IPR003140,IPR029058 |
| 373 | HORVU.MOREX.r3.2HG02130 | high | protein_cod | *-* | Protein F12 | not classifie | no mercator | NA | NA | NA |
| 374 | HORVU.MOREX.r3.2HG02130 | high | protein_cod | *** | Blue copper | phytocyanin | no mercator | GO:000905 | PF02298 | IPR008972,IPR003245 |
| 375 | HORVU.MOREX.r3.2HG02131 | high | protein_cod | *** | Blue copper | phytocyanin | no mercator | GO:000905 | PF02298 | IPR008972,IPR003245 |
| 376 | HORVU.MOREX.r3.2HG02137 | low | non_coding | --* | ATP-depend | not classifie | no mercator | GO:000016 | NA | NA |
| 377 | HORVU.MOREX.r3.2HG02141 | high | protein_cod | *** | CoA ligase | crotonobeta | no mercator | GO:000382 | PF00501,PF | IPR000873,IPR025110 |
| 378 | HORVU.MOREX.r3.2HG02146 | high | protein_cod | *** | Glutathione | class tau glu | redox home | GO:001674 | PF13417 | IPR012336,IPR004045,IPR010987 |
| 379 | HORVU.MOREX.r3.2HG02146 | high | protein_cod | *** | Glutathione | class tau glu | redox home | GO:001674 | PF13417,PF | IPR012336,IPR010987,IPR004045,IPR004046 |
| 380 | HORVU.MOREX.r3.2HG02170 | high | protein_cod | *** | C2 calcium/ | ft interacting | no mercator | GO:001602 | PF00168,PF | IPR000008,IPR000008,IPR000008,IPR000008,IPR000008,IPR000008,IPR013583,IPR000008,IPR00000 |
| 381 | HORVU.MOREX.r3.3HG02193 | low | protein_cod | --* | Glutamate r | not classifie | no mercator | GO:000497 | NA | NA |
| 382 | HORVU.MOREX.r3.3HG02211 | high | protein_cod | *** | Glycosyltra | xylan alpha- | cell wall org | GO:001602 | PF04577 | IPR007657 |
| 383 | HORVU.MOREX.r3.3HG02212 | high | protein_cod | *** | Glycosyltra | xylan alpha- | cell wall org | GO:001602 | PF04577 | IPR007657 |
| 384 | HORVU.MOREX.r3.3HG02216 | low | protein_cod | NA | NA | porr domain | no mercator | NA | NA | NA |
| 385 | HORVU.MOREX.r3.3HG02219 | high | protein_cod | *** | Protein NEG | regulatory p | external stir | GO:000551 | PF15699 | IPR031425 |
| 386 | HORVU.MOREX.r3.3HG02221 | low | protein_cod | *-* | Abscisic aci | neoxanthin | secondary r | GO:001602 | PF14108 | IPR025461 |
| 387 | HORVU.MOREX.r3.3HG02223 | high | protein_cod | *** | Mannose-6- | phosphoma | carbohydrat | GO:000447 | PF01238 | IPR011051,IPR001250,IPR001250 |
| 388 | HORVU.MOREX.r3.3HG02223 | high | protein_cod | -** | Trypsin inhil | Bowman-Bi | protein hom | GO:000486 | PF00228,PF | IPR000877,IPR000877,IPR000877,IPR000877,IPR000877 |
| 389 | HORVU.MOREX.r3.3HG02231 | low | protein_cod | --* | aberrant roc | EC_2.4 glyc | enzyme clas | NA | NA | NA |

| | | | | | | | | | | |
|---|---|---|---|---|---|---|---|---|---|---|
| 390 | HORVU.MOREX.r3.3HG02232( | high | protein_cod | *** | Expansin | NA | NA | GO:000557 | PF01357,PF | IPR007117,IPR009009,IPR007117,IPR00900 |
| 391 | HORVU.MOREX.r3.3HG02243( | low | non_coding | *-* | mediator of | xylosyltrans | no mercato | NA | NA | NA |
| 392 | HORVU.MOREX.r3.3HG02246 | high | protein_cod | *** | Sarcoplasm | histidine ric | no mercato | NA | NA | NA |
| 393 | HORVU.MOREX.r3.3HG02248 | high | protein_cod | *-* | Basic 7S glo | Pepsin-type | protein hom | GO:000419 | PF14541,PF | IPR032799,IPR021109,IPR032861 |
| 394 | HORVU.MOREX.r3.3HG02253 | high | protein_cod | *** | Heat shock | class-C-I sn | protein hom | NA | PF00011 | IPR002068,IPR008978 |
| 395 | HORVU.MOREX.r3.3HG02253 | high | protein_cod | *** | Protein kina | LRK10-1-lik | protein moc | NA | PF14380,PF | IPR032872,IPR000719,IPR025287,IPR01100 |
| 396 | HORVU.MOREX.r3.3HG02261 | high | protein_cod | *** | Heat shock | class-C-I sn | protein hom | NA | PF00011 | IPR002068,IPR008978 |
| 397 | HORVU.MOREX.r3.3HG02261 | high | protein_cod | *** | Heat shock | class-C-I sn | protein hom | NA | PF00011 | IPR002068,IPR008978 |
| 398 | HORVU.MOREX.r3.3HG02279 | high | protein_cod | *** | Glycosyltra | EC_2.4 glyc | enzyme clas | NA | PF00201 | IPR002213 |
| 399 | HORVU.MOREX.r3.3HG02281 | high | protein_cod | *** | MYB transcr | MYBR-R-typ | rna biosynt | GO:000367 | PF00249 | IPR009057,IPR006447,IPR009057,IPR00100 |
| 400 | HORVU.MOREX.r3.3HG02289 | high | protein_cod | --* | Sulfate ade | gtd binding | no mercato | GO:000016 | NA | NA |
| 401 | HORVU.MOREX.r3.3HG02294( | high | protein_cod | *** | Receptor-lik | EC_2.7 tran | enzyme clas | GO:000016 | PF00069,PF | IPR000719,IPR001938,IPR011009,IPR00193 |
| 402 | HORVU.MOREX.r3.3HG02315 | high | protein_cod | *** | F-box protei | f box domai | no mercato | NA | PF00646,PF | IPR001810,IPR001810,IPR025886 |
| 403 | HORVU.MOREX.r3.3HG02336 | high | protein_cod | *** | SH3 domair | regulatory p | protein hom | GO:004687 | PF01363,PF | IPR011011,IPR000306,IPR007461 |
| 404 | HORVU.MOREX.r3.3HG02340( | high | protein_cod | *** | Xyloglucan | EC_2.4 glyc | enzyme clas | GO:000455 | PF06955,PF | IPR010713,IPR013320,IPR000757 |
| 405 | HORVU.MOREX.r3.3HG02340 | high | protein_cod | *** | Inhibitor pro | PR6 proteas | protein hom | GO:000486 | PF00280 | IPR000864,IPR000864 |
| 406 | HORVU.MOREX.r3.3HG02340 | high | protein_cod | --* | microtubule | not classifie | no mercato | NA | NA | NA |
| 407 | HORVU.MOREX.r3.3HG02341 | low | non_coding | *** | Cytochrome | EC_1.14 oxi | enzyme clas | GO:000449 | PF00067 | IPR001128,IPR001128 |
| 408 | HORVU.MOREX.r3.3HG02347 | high | protein_cod | *** | Nuclease | ssDNA/dsD | chromatin o | GO:000367 | PF00565 | IPR016071,IPR016071,IPR016071 |
| 409 | HORVU.MOREX.r3.3HG02352 | low | protein_cod | --* | Anaphase-p | not classifie | no mercato | GO:001602 | NA | NA |
| 410 | HORVU.MOREX.r3.3HG02357 | high | protein_cod | -** | FMN-depen | nad p h dey | no mercato | GO:000582 | PF03358 | IPR029039,IPR005025 |
| 411 | HORVU.MOREX.r3.3HG02359 | high | protein_cod | *** | transcriptio | regulatory p | plant reproc | NA | PF14144 | IPR025422 |
| 412 | HORVU.MOREX.r3.3HG02361 | high | protein_cod | *** | Glutaredoxi | glutaredoxir | protein moc | GO:000562 | PF00462 | IPR002109,IPR012336 |
| 413 | HORVU.MOREX.r3.3HG02366 | high | protein_cod | *** | ATP-depenc | ATP-depend | cellular resp | GO:000016 | PF00365 | IPR000023,IPR000023 |
| 414 | HORVU.MOREX.r3.3HG02366 | high | protein_cod | *** | Myb transcr | MYB class-F | rna biosynt | GO:000367 | PF00249,PF | IPR009057,IPR001005,IPR001005 |
| 415 | HORVU.MOREX.r3.3HG02368 | high | protein_cod | *** | 2-oxoglutar | EC_1.14 oxi | enzyme clas | GO:001649 | PF14226,PF | IPR026992,IPR005123 |
| 416 | HORVU.MOREX.r3.3HG02369( | high | protein_cod | *** | MYB-RELATI | MYBR-R-typ | rna biosynt | GO:000367 | PF00249 | IPR006447,IPR009057,IPR001005 |
| 417 | HORVU.MOREX.r3.3HG02370 | high | protein_cod | *** | Harbinger tr | dde tnp dor | no mercato | NA | PF13359 | IPR027806 |
| 418 | HORVU.MOREX.r3.3HG02373( | high | protein_cod | *** | WRKY famil | transcriptio | rna biosynt | GO:000367 | PF03106 | IPR003657,IPR003657 |
| 419 | HORVU.MOREX.r3.3HG02375 | high | protein_cod | *** | Heat-shock | class-C-II sr | protein hom | NA | PF00011 | IPR008978,IPR002068 |
| 420 | HORVU.MOREX.r3.3HG02377 | high | protein_cod | *** | Heat shock | class-C-II sr | protein hom | NA | PF00011 | IPR008978,IPR002068 |
| 421 | HORVU.MOREX.r3.3HG02379 | high | protein_cod | *-* | Type I inosit | type-I inosit | multi-proce | GO:001678 | IPR005135,IPR005135 | |
| 422 | HORVU.MOREX.r3.3HG02382( | high | protein_cod | *** | Transcriptio | TRIHELIX tr | rna biosynt | NA | NA | NA |
| 423 | HORVU.MOREX.r3.3HG02383 | high | protein_cod | --* | Hemoglobir | pmei domai | no mercato | GO:000927 | NA | NA |
| 424 | HORVU.MOREX.r3.3HG02391 | high | protein_cod | *** | Glycosyltra | EC_2.4 glyc | enzyme clas | GO:000815 | PF00201 | IPR002213 |
| 425 | HORVU.MOREX.r3.3HG02399( | high | protein_cod | *** | Auxin-respo | transcriptio | phytohormo | GO:000563 | PF02309 | IPR033389 |
| 426 | HORVU.MOREX.r3.3HG02400 | high | protein_cod | *** | Protein kina | WAK/WAKL | protein moc | GO:000467 | PF00069,PF | IPR000719,IPR025287,IPR011009 |
| 427 | HORVU.MOREX.r3.3HG02405 | high | protein_cod | *** | Glycosyltra | EC_2.4 glyc | enzyme clas | GO:000815 | PF00201 | IPR002213 |
| 428 | HORVU.MOREX.r3.3HG02405 | high | protein_cod | *** | Glycosyltra | EC_2.4 glyc | enzyme clas | GO:000815 | PF00201 | IPR002213 |
| 429 | HORVU.MOREX.r3.3HG02409 | high | protein_cod | *** | Plant/MSJ1 | not classifie | no mercato | NA | NA | NA |
| 430 | HORVU.MOREX.r3.3HG02416 | low | non_coding | --* | DNA-directe | class tau gl | redox home | GO:000367 | NA | NA |
| 431 | HORVU.MOREX.r3.3HG02417 | low | plastid_rela | *** | Triose phos | phosphome | solute trans | GO:000950 | PF03151 | IPR004853,IPR004696 |
| 432 | HORVU.MOREX.r3.3HG02420 | high | protein_cod | *** | Alpha/beta- | abhydrolase | no mercato | GO:000815 | PF07859 | IPR029058,IPR013094 |
| 433 | HORVU.MOREX.r3.3HG02430 | low | non_coding | *** | Cytochrome | cinnamate 4 | secondary n | GO:000449 | PF00067 | IPR001128,IPR001128 |
| 434 | HORVU.MOREX.r3.3HG02442 | high | protein_cod | *** | Alpha/beta- | hydrolase 4 | no mercato | GO:001602 | PF12146 | IPR022742,IPR029058 |
| 435 | HORVU.MOREX.r3.3HG02452 | high | protein_cod | *** | Cysteine pro | EC_3.4 hydr | enzyme clas | GO:000650 | PF00112,PF | IPR000668,IPR013201 |
| 436 | HORVU.MOREX.r3.3HG02454 | high | protein_cod | --* | Cellulose sy | signaling pe | no mercato | GO:000588 | NA | NA |
| 437 | HORVU.MOREX.r3.3HG02460( | low | non_coding | *** | Cytochrome | sterol C-22 | lipid metabo | GO:000449 | PF00067 | IPR001128,IPR001128 |
| 438 | HORVU.MOREX.r3.3HG02461 | high | protein_cod | *** | Heme-bindi | accessory h | coenzyme n | NA | PF04832 | IPR006917,IPR011256 |
| 439 | HORVU.MOREX.r3.3HG02466 | high | protein_cod | --* | DUF4228 do | not classifie | no mercato | NA | NA | NA |
| 440 | HORVU.MOREX.r3.3HG02473 | high | protein_cod | --* | Soul heme-l | tsa wollemi | no mercato | NA | PF10184 | IPR018790,IPR032710 |
| 441 | HORVU.MOREX.r3.3HG02476 | high | protein_cod | *** | Mitochondri | manganese | solute trans | GO:001602 | PF00153,PF | IPR018108,IPR018108,IPR018108,IPR023395,IPR02339 |
| 442 | HORVU.MOREX.r3.3HG02481 | high | protein_cod | *-* | Late embryo | why domain | no mercato | GO:000926 | PF03168 | IPR004864 |
| 443 | HORVU.MOREX.r3.3HG02484 | high | protein_cod | *** | Short-chain | component | coenzyme n | GO:000953 | PF00106 | IPR002347,IPR016040 |
| 444 | HORVU.MOREX.r3.3HG02489 | high | protein_cod | *** | Thioesteras | 4hbt domai | no mercato | NA | PF03061 | IPR029069,IPR006683 |
| 445 | HORVU.MOREX.r3.3HG02522 | high | protein_cod | --* | Transcriptio | expressed p | no mercato | GO:004698 | NA | NA |
| 446 | HORVU.MOREX.r3.3HG02536 | high | protein_cod | *** | Late embryo | late embryo | no mercato | GO:000695 | PF03242 | IPR004926 |
| 447 | HORVU.MOREX.r3.3HG02540 | high | protein_cod | *** | Kinase fami | RLCK-VIIa r | protein moc | GO:000018 | PF00069 | IPR011009,IPR000719 |
| 448 | HORVU.MOREX.r3.3HG02550 | high | protein_cod | --* | transmemb | PSY precurs | phytohormo | NA | NA | NA |
| 449 | HORVU.MOREX.r3.3HG02554 | high | protein_cod | --* | Disease res | not classifie | no mercato | NA | NA | NA |
| 450 | HORVU.MOREX.r3.3HG02559( | high | protein_cod | *** | Kinase fami | EC_2.7 tran | enzyme clas | GO:000016 | PF07714 | IPR011009,IPR001245 |
| 451 | HORVU.MOREX.r3.3HG02588 | high | protein_cod | *** | F-box protei | substrate ac | protein hom | NA | PF00646 | IPR001810,IPR001810 |
| 452 | HORVU.MOREX.r3.3HG02647 | low | protein_cod | --* | Abscisic aci | gram domai | no mercato | NA | NA | NA |
| 453 | HORVU.MOREX.r3.3HG02678 | low | plastid_rela | *-* | Protein CHL | protein chu | no mercato | NA | NA | NA |
| 454 | HORVU.MOREX.r3.3HG02684 | high | protein_cod | *** | Cytokinin ril | cytokinin ph | phytohormo | NA | PF03641 | IPR005269,IPR031100 |

| | | | | | | | | | | |
|---|---|---|---|---|---|---|---|---|---|---|
| 455 | HORVU.MOREX.r3.3HG02702 | high | protein_cod | *-* | ATP-depend | EC_3.6 hydr | enzyme clas | GO:000016 | PF14363,PF | IPR025753,IPR003959,IPR027417 |
| 456 | HORVU.MOREX.r3.3HG02708 | high | protein_cod | *** | Pleiotropic | subfamily A | solute trans | GO:000016 | PF00005,PF | IPR003439,IPR003439,IPR013525,IPR013525,IPR027417,IPR013581,IPR027417,IPR02948 |
| 457 | HORVU.MOREX.r3.3HG02737 | low | non_coding | *** | Cytochrome | EC_1.14 oxi | enzyme clas | GO:000449 | PF00067 | IPR001128,IPR001128 |
| 458 | HORVU.MOREX.r3.3HG02740 | high | protein_cod | --* | DWNN dom | not classifie | no mercato | NA | NA | NA |
| 459 | HORVU.MOREX.r3.3HG02746 | high | protein_cod | *** | Cinnamoyl- | 3beta hsd d | no mercato | GO:000385 | PF01073 | IPR016040,IPR002225 |
| 460 | HORVU.MOREX.r3.3HG02746 | high | protein_cod | *** | DUF1645 fa | not classifie | no mercato | NA | PF07816 | IPR012442 |
| 461 | HORVU.MOREX.r3.3HG02747 | high | protein_cod | *** | Carbonic ar | beta-type ca | photosynthe | GO:000408 | PF00484 | IPR001765,IPR001765 |
| 462 | HORVU.MOREX.r3.3HG02750 | high | protein_cod | *-* | ATP-depend | EC_3.6 hydr | enzyme clas | GO:000016 | PF00004,PF | IPR027417,IPR003959,IPR025753 |
| 463 | HORVU.MOREX.r3.3HG02752 | low | plastid_rela | *** | Protein CHL | protein chu | no mercato | GO:001251 | NA | NA |
| 464 | HORVU.MOREX.r3.3HG02757 | high | protein_cod | *** | Potassium | voltage-gate | solute trans | GO:000521 | PF12796,PF | IPR018490,IPR020683,IPR020683,IPR000595,IPR005821,IPR021789,IPR02068 |
| 465 | HORVU.MOREX.r3.3HG02781 | high | protein_cod | *** | WRKY trans | transcriptio | rna biosynth | GO:000367 | PF03106 | IPR003657,IPR003657 |
| 466 | HORVU.MOREX.r3.3HG02785 | high | protein_cod | *** | Hydroxyacyl | sulfur dioxy | amino acid | GO:001678 | PF00753 | IPR001279,IPR001279 |
| 467 | HORVU.MOREX.r3.3HG02786 | high | protein_cod | *** | Glutaredoxi | glutaredoxir | protein mod | GO:000562 | PF00462 | IPR002109,IPR011905,IPR012336 |
| 468 | HORVU.MOREX.r3.3HG02800 | high | protein_cod | --* | Heavy meta | hma domair | no mercato | NA | PF00403 | IPR006121,IPR006121 |
| 469 | HORVU.MOREX.r3.3HG02806 | high | protein_cod | --* | Uridine kina | glycerate kir | photosynthe | GO:000016 | IPR027417 | |
| 470 | HORVU.MOREX.r3.3HG02808 | high | protein_cod | *-* | Myb transcr | MYB class-F | rna biosynth | GO:000367 | PF00249,PF | IPR001005,IPR001005,IPR009057 |
| 471 | HORVU.MOREX.r3.3HG02815 | high | protein_cod | *** | Glutathione | class tau glu | redox home | GO:001674 | PF13417 | IPR004045,IPR010987,IPR012336 |
| 472 | HORVU.MOREX.r3.3HG02817 | high | protein_cod | --* | splicing fact | RNA splicing | rna process | NA | PF00076,PF | IPR012677,IPR000504,IPR000504 |
| 473 | HORVU.MOREX.r3.3HG02826 | high | protein_cod | *** | ABC transpo | subfamily A | solute trans | GO:000016 | PF00005,PF | IPR003439,IPR003439,IPR027417,IPR011527,IPR011527,IPR027417,IPR011527,IPR01152 |
| 474 | HORVU.MOREX.r3.3HG02829 | low | protein_cod | --* | Single-stran | MAP3K-MEK | multi-proce | NA | NA | NA |
| 475 | HORVU.MOREX.r3.3HG02834 | high | protein_cod | *** | Kelch repea | substrate(C | secondary r | NA | PF00646 | IPR001810,IPR001810 |
| 476 | HORVU.MOREX.r3.3HG02840 | high | protein_cod | *** | Cytokinin ril | cytokinin ph | phytohormo | GO:000969 | PF03641 | IPR005269,IPR031100 |
| 477 | HORVU.MOREX.r3.3HG02848 | low | plastid_rela | *** | WEB family | web family | no mercato | NA | NA | NA |
| 478 | HORVU.MOREX.r3.3HG02850 | high | protein_cod | *** | S-adenosyl | methyltrans | no mercato | GO:000815 | PF08241 | IPR029063,IPR013216 |
| 479 | HORVU.MOREX.r3.3HG02855 | high | protein_cod | *** | Serine acety | serine O-ac | amino acid | GO:000573 | PF06426,PF | IPR010493,IPR011004,IPR001451,IPR005881 |
| 480 | HORVU.MOREX.r3.3HG02863 | high | protein_cod | *-* | senescence | not classifie | no mercato | NA | PF04520 | IPR007608 |
| 481 | HORVU.MOREX.r3.3HG02865 | high | protein_cod | -** | F-box family | f box domai | no mercato | NA | PF08387,PF | IPR006566,IPR001810,IPR001810 |
| 482 | HORVU.MOREX.r3.3HG02866 | high | protein_cod | *** | WRKY trans | transcriptio | rna biosynth | GO:000367 | PF03106 | IPR003657,IPR003657 |
| 483 | HORVU.MOREX.r3.3HG02867 | high | protein_cod | *-* | RING/FYVE/ | ring ch type | no mercato | GO:000827 | PF12906 | IPR011016 |
| 484 | HORVU.MOREX.r3.3HG02870 | high | protein_cod | *** | Glycosyltrar | EC_2.4 glyc | enzyme clas | GO:000815 | PF00201 | IPR002213 |
| 485 | HORVU.MOREX.r3.3HG02870 | high | protein_cod | *** | Glycosyltrar | EC_2.4 glyc | enzyme clas | GO:000815 | PF00201 | IPR002213 |
| 486 | HORVU.MOREX.r3.3HG02878 | high | protein_cod | *** | Receptor-lik | SCREW pep | phytohormo | GO:000016 | PF07714,PF | IPR001245,IPR013210,IPR032675,IPR011009,IPR001611,IPR001611,IPR001611,IPR03267 |
| 487 | HORVU.MOREX.r3.3HG02884 | high | protein_cod | *** | Protein kina | MAP3K-RAF | multi-proce | GO:000467 | PF07714 | IPR001245,IPR011009 |
| 488 | HORVU.MOREX.r3.3HG02888 | high | protein_cod | *** | MACPF dom | regulatory p | multi-proce | NA | PF01823 | IPR020864 |
| 489 | HORVU.MOREX.r3.3HG02890 | high | protein_cod | *** | Coiled-coil | coiled coil d | no mercato | NA | PF05670 | IPR008532 |
| 490 | HORVU.MOREX.r3.3HG02893 | high | protein_cod | *-* | Vascular pla | VOZ transcr | rna biosynth | NA | NA | NA |
| 491 | HORVU.MOREX.r3.3HG02898 | high | protein_cod | *** | Gibberellin | gibberellin 2 | phytohormo | GO:001649 | PF03171,PF | IPR005123,IPR026992 |
| 492 | HORVU.MOREX.r3.3HG02899 | high | protein_cod | *** | Heavy meta | hma domair | no mercato | GO:003000 | PF00403 | IPR006121,IPR006121 |
| 493 | HORVU.MOREX.r3.3HG02904 | high | protein_cod | *** | Ankyrin repe | pgg domain | no mercato | NA | PF12796 | IPR020683,IPR020683 |
| 494 | HORVU.MOREX.r3.3HG02919 | high | protein_cod | *** | Peroxidase | EC_1.11 oxi | enzyme clas | GO:000460 | PF00141 | IPR010255,IPR002016 |
| 495 | HORVU.MOREX.r3.3HG02920 | high | protein_cod | *** | Tyrosine de | aromatic an | amino acid | GO:000382 | PF00282 | IPR002129,IPR015424 |
| 496 | HORVU.MOREX.r3.3HG02922 | high | protein_cod | *** | Mal d 1-ass | fra a 1 ssoci | no mercato | NA | NA | NA |
| 497 | HORVU.MOREX.r3.3HG02924 | high | protein_cod | *** | Core-2/I-bra | beta glucurc | no mercato | GO:000837 | PF02485 | IPR003406 |
| 498 | HORVU.MOREX.r3.3HG02930 | high | protein_cod | *** | BnaA05g07 | verprolin | no mercato | NA | NA | NA |
| 499 | HORVU.MOREX.r3.3HG02934 | high | protein_cod | *** | Expansin | alpha-class | cell wall org | GO:000557 | PF01357,PF | IPR007117,IPR009009,IPR007117,IPR00900 |
| 500 | HORVU.MOREX.r3.3HG02934 | high | protein_cod | *** | Expansin | NA | NA | GO:000557 | PF01357,PF | IPR007117,IPR007117,IPR009009,IPR00900 |
| 501 | HORVU.MOREX.r3.3HG02934 | high | protein_cod | *** | Expansin | NA | NA | GO:000557 | PF01357,PF | IPR007117,IPR007117,IPR009009,IPR00900 |
| 502 | HORVU.MOREX.r3.3HG02934 | high | protein_cod | *** | Expansin | alpha-class | cell wall org | GO:000557 | PF03330,PF | IPR009009,IPR007117,IPR007117,IPR00900 |
| 503 | HORVU.MOREX.r3.3HG02935 | high | protein_cod | *** | Protein kina | L-Lectin pro | protein mod | GO:000467 | PF00139,PF | IPR013320,IPR001220,IPR000719,IPR01100 |
| 504 | HORVU.MOREX.r3.3HG02935 | high | protein_cod | *** | Secretory ca | regulatory p | vesicle traff | NA | PF04144 | IPR007273 |
| 505 | HORVU.MOREX.r3.3HG02936 | high | protein_cod | *** | Vacuolar so | component | vesicle traff | GO:000081 | PF03997 | IPR007143 |
| 506 | HORVU.MOREX.r3.3HG02940 | high | protein_cod | *** | Serine/threo | SD-1 protein | protein mod | GO:000016 | PF07714,PF | IPR001245,IPR001480,IPR003609,IPR001480,IPR000858,IPR01100 |
| 507 | HORVU.MOREX.r3.3HG02943 | high | protein_cod | *** | Breast carci | component | protein hom | GO:004259 | PF12490 | IPR022175,IPR017986,IPR017986,IPR01798 |
| 508 | HORVU.MOREX.r3.3HG02943 | high | protein_cod | *** | Peroxidase | EC_1.11 oxi | enzyme clas | GO:000460 | PF00141 | IPR002016,IPR010255 |
| 509 | HORVU.MOREX.r3.3HG02946 | high | protein_cod | --* | Iron-sulfur c | porphobilin | no mercato | GO:000573 | NA | NA |
| 510 | HORVU.MOREX.r3.3HG02947 | low | protein_cod | --* | alpha/beta- | not classifie | no mercato | NA | NA | NA |
| 511 | HORVU.MOREX.r3.3HG02949 | high | protein_cod | *** | Ethylene-re | subgroup EF | rna biosynth | GO:000367 | PF00847 | IPR016177,IPR001471 |
| 512 | HORVU.MOREX.r3.3HG02951 | high | protein_cod | *-* | Oxidoreduc | not classifie | no mercato | NA | NA | NA |
| 513 | HORVU.MOREX.r3.3HG02953 | high | protein_cod | *** | Auxin efflux | auxin transp | phytohormo | GO:000973 | PF03547,PF | IPR004776,IPR004776 |
| 514 | HORVU.MOREX.r3.3HG02956 | high | protein_cod | *** | F-box family | substrate ac | protein hom | NA | PF00646 | IPR001810,IPR001810 |
| 515 | HORVU.MOREX.r3.3HG02964 | high | protein_cod | *** | Glycerophos | glycerophos | no mercato | GO:000662 | PF03009 | IPR030395,IPR017946,IPR017946 |
| 516 | HORVU.MOREX.r3.3HG02965 | high | protein_cod | *** | Ribosomal p | ribosomal p | no mercato | GO:000373 | NA | NA |
| 517 | HORVU.MOREX.r3.3HG02966 | high | protein_cod | *** | NAC domain | NAC transcr | rna biosynth | GO:000367 | PF02365 | IPR003441,IPR003441 |
| 518 | HORVU.MOREX.r3.3HG02974 | low | non_coding | *** | Cytochrome | cinnamate 4 | secondary r | GO:000449 | PF00067 | IPR001128,IPR001128 |
| 519 | HORVU.MOREX.r3.3HG02976 | high | protein_cod | *** | Kinase fami | RLCK-II rece | protein mod | GO:000016 | PF07714 | IPR011009,IPR001245 |

| | | | | | | | | | | |
|---|---|---|---|---|---|---|---|---|---|---|
| 520 | HORVU.MOREX.r3.3HG02977 | high | protein_cod | *** | Expansin | alpha-class | cell wall org | GO:00055 | PF03330,PF | IPR009009,IPR007117,IPR007117,IPR00900 |
| 521 | HORVU.MOREX.r3.3HG02978 | high | protein_cod | *** | U-box dom | E3 ubiquitin | protein hom | GO:000484 | PF00514,PF | IPR000225,IPR016024,IPR003613 |
| 522 | HORVU.MOREX.r3.3HG02978 | high | protein_cod | *** | BAG family | ubiquitin do | no mercato | NA | PF00240 | IPR029071,IPR000626 |
| 523 | HORVU.MOREX.r3.3HG02979 | high | protein_cod | *** | U-box dom | E3 ubiquitin | protein hom | NA | PF00514,PF | IPR000225,IPR000225,IPR000225,IPR016024 |
| 524 | HORVU.MOREX.r3.3HG02979 | high | protein_cod | *** | Jasmonate | EC_2.1 tran | enzyme clas | GO:000816 | PF03492 | IPR029063,IPR005299 |
| 525 | HORVU.MOREX.r3.3HG02986 | high | protein_cod | *** | Myb transcr | MYB class-F | rna biosynth | GO:000098 | PF00249,PF | IPR009057,IPR001005,IPR001005 |
| 526 | HORVU.MOREX.r3.3HG02990 | high | protein_cod | *** | Protein NRT | anion trans | solute trans | GO:001602 | PF00854,PF | IPR020846,IPR020846,IPR020846,IPR000109,IPR00010 |
| 527 | HORVU.MOREX.r3.3HG02994 | high | protein_cod | *-* | Ethylene-re | subgroup E | rna biosynth | GO:000367 | PF00847 | IPR016177,IPR001471 |
| 528 | HORVU.MOREX.r3.3HG02996 | high | protein_cod | --* | Late embry | 2 3 bisphos | no mercato | NA | NA | NA |
| 529 | HORVU.MOREX.r3.3HG02998 | high | protein_cod | *** | Actin | actin filame | cytoskeleto | GO:000016 | PF00022 | IPR004000 |
| 530 | HORVU.MOREX.r3.3HG02998 | high | protein_cod | *** | U-box dom | E3 ubiquitin | protein hom | GO:000484 | PF04564 | IPR003613,IPR016024 |
| 531 | HORVU.MOREX.r3.3HG02999 | high | protein_cod | *** | Early respo | pam domai | no mercato | NA | NA | NA |
| 532 | HORVU.MOREX.r3.3HG03000 | low | protein_cod | --* | Helicase pr | EC_2.4 glyc | enzyme clas | NA | NA | NA |
| 533 | HORVU.MOREX.r3.3HG03002 | high | protein_cod | *** | MYB transc | MYBR-R-typ | rna biosynth | GO:000367 | PF00249,PF | IPR006447,IPR009057,IPR009057,IPR001005,IPR00100 |
| 534 | HORVU.MOREX.r3.3HG03005 | high | protein_cod | *** | Glucan end | EC_3.2 glyc | enzyme clas | GO:000455 | PF00332 | IPR017853,IPR000490 |
| 535 | HORVU.MOREX.r3.3HG03006 | high | protein_cod | *** | Acidic endo | acidic chitin | external stir | GO:000455 | PF00704 | IPR017853,IPR001223 |
| 536 | HORVU.MOREX.r3.3HG03008 | low | non_coding | *** | Cytochrom | jasmonoyl- | phytohorm | GO:000550 | PF00067 | IPR001128,IPR001128 |
| 537 | HORVU.MOREX.r3.3HG03011 | high | protein_cod | *** | Nematode r | nematode r | no mercato | GO:000695 | PF07231,PF | IPR009869,IPR009743 |
| 538 | HORVU.MOREX.r3.3HG03013 | high | protein_cod | *** | Glycerol-3- | glycerol-3-p | lipid metab | GO:000815 | PF01553 | IPR023214,IPR002123 |
| 539 | HORVU.MOREX.r3.3HG03017 | high | protein_cod | *** | 3-oxo-5-alp | steroid 5-al | phytohorm | GO:000386 | PF02544 | IPR001104 |
| 540 | HORVU.MOREX.r3.3HG03018 | high | protein_cod | *** | Soul heme- | accessory h | coenzyme n | GO:000577 | PF04832 | IPR006917,IPR011256,IPR011256 |
| 541 | HORVU.MOREX.r3.3HG03021 | high | protein_cod | *** | Protein pho | clade A pho | protein mod | GO:000382 | PF00481 | IPR001932,IPR001932,IPR001932 |
| 542 | HORVU.MOREX.r3.3HG03022 | high | protein_cod | *** | Avr9/Cf-9 ra | dna integrity | no mercato | NA | PF05003,PF | IPR007700,IPR021864 |
| 543 | HORVU.MOREX.r3.3HG03026 | high | protein_cod | *** | Calcium-de | c2 domain | no mercato | NA | PF00168 | IPR000008,IPR000008 |
| 544 | HORVU.MOREX.r3.3HG03036 | high | protein_cod | --* | RNAse E/G- | not classifie | no mercato | NA | NA | NA |
| 545 | HORVU.MOREX.r3.3HG03037 | high | protein_cod | *** | 2-oxoglutar | jasmonic ac | phytohorm | NA | PF14226,PF | IPR026992,IPR005123 |
| 546 | HORVU.MOREX.r3.3HG03039 | high | protein_cod | *** | Trichome bi | xylan O-ace | cell wall org | NA | PF13839,PF | IPR026057,IPR025846 |
| 547 | HORVU.MOREX.r3.3HG03047 | high | protein_cod | *** | RING finger | E3 ubiquitin | protein hom | NA | NA | NA |
| 548 | HORVU.MOREX.r3.3HG03051 | high | protein_cod | *** | Acid phosp | acid phosp | no mercato | GO:000460 | PF02681 | IPR003832 |
| 549 | HORVU.MOREX.r3.3HG03054 | high | protein_cod | *** | Late embry | late embryo | no mercato | GO:000695 | PF03242 | IPR004926 |
| 550 | HORVU.MOREX.r3.3HG03056 | high | protein_cod | *** | Receptor-li | LRR-III prot | protein mod | GO:000016 | PF08263,PF | IPR013210,IPR011009,IPR000719,IPR001611,IPR03267 |
| 551 | HORVU.MOREX.r3.3HG03066 | high | protein_cod | --* | Pectate lya | not classifie | no mercato | GO:001682 | NA | NA |
| 552 | HORVU.MOREX.r3.3HG03066 | high | protein_cod | --* | Cyclophilin- | ethylene sig | phytohorm | NA | NA | NA |
| 553 | HORVU.MOREX.r3.3HG03070 | high | protein_cod | *** | NAC domai | NAC transc | rna biosynth | GO:000367 | PF02365 | IPR003441,IPR003441 |
| 554 | HORVU.MOREX.r3.3HG03074 | high | protein_cod | *** | Acyl-[acyl-c | delta-9 stea | lipid metab | GO:000663 | PF03405 | IPR005067,IPR009078 |
| 555 | HORVU.MOREX.r3.3HG03081 | high | protein_cod | *** | BTB/POZ an | substrate a | protein hom | GO:000371 | PF00651,PF | IPR000197,IPR000210,IPR011333,IPR000197 |
| 556 | HORVU.MOREX.r3.3HG03081 | high | protein_cod | *** | BTB/POZ an | substrate a | protein hom | GO:000371 | PF02135,PF | IPR000197,IPR011333,IPR011333,IPR000197,IPR00021 |
| 557 | HORVU.MOREX.r3.3HG03083 | high | protein_cod | --* | Cysteine-ric | gnk homolo | no mercato | GO:000467 | PF01657 | IPR002902 |
| 558 | HORVU.MOREX.r3.3HG03084 | high | protein_cod | *-* | cysteine-ric | gnk homolo | no mercato | NA | PF01657,PF | IPR002902,IPR002902 |
| 559 | HORVU.MOREX.r3.3HG03093 | high | protein_cod | *** | Protease in | aai domain | no mercato | GO:000650 | PF14368 | IPR016140,IPR016140 |
| 560 | HORVU.MOREX.r3.3HG03093 | high | protein_cod | *** | Protein kina | MAP3K-RAF | multi-proce | GO:000467 | PF13426,PF | IPR011009,IPR000014,IPR001245,IPR000014,IPR00001 |
| 561 | HORVU.MOREX.r3.3HG03094 | high | protein_cod | *** | DUF506 fan | dna directe | no mercato | NA | PF04720 | IPR006502,IPR006502 |
| 562 | HORVU.MOREX.r3.3HG03095 | high | protein_cod | *-* | Loricrin | not classifie | no mercato | NA | NA | NA |
| 563 | HORVU.MOREX.r3.3HG03103 | high | protein_cod | --* | 10 kDa cha | f box domai | no mercato | GO:000552 | NA | NA |
| 564 | HORVU.MOREX.r3.3HG03104 | high | protein_cod | *** | Glycosyltra | nucleotid tr | no mercato | GO:000013 | PF03407 | IPR005069 |
| 565 | HORVU.MOREX.r3.3HG03111 | high | protein_cod | *** | Transmemb | not classifie | no mercato | GO:001602 | NA | NA |
| 566 | HORVU.MOREX.r3.3HG03115 | high | protein_cod | *** | SAUR-like a | auxin respo | no mercato | GO:000973 | PF02519 | IPR003676 |
| 567 | HORVU.MOREX.r3.3HG03116 | high | protein_cod | *** | Glutathione | class tau gl | redox home | GO:001674 | PF02798 | IPR004045,IPR012336,IPR010987 |
| 568 | HORVU.MOREX.r3.3HG03117 | high | protein_cod | *** | NAC domai | NAC transc | rna biosynth | GO:000367 | PF02365 | IPR003441,IPR003441 |
| 569 | HORVU.MOREX.r3.3HG03117 | high | protein_cod | *** | YGL010w-li | 2 hydroxy p | no mercato | GO:001602 | PF06127 | IPR009305 |
| 570 | HORVU.MOREX.r3.3HG03117 | high | protein_cod | *** | Kinase fami | CDK9 prote | protein mod | GO:000467 | PF00069 | IPR011009,IPR000719 |
| 571 | HORVU.MOREX.r3.3HG03130 | high | protein_cod | *** | Thiamin pyr | thiamine di | coenzyme n | GO:000016 | PF04265,PF | IPR007371,IPR007373,IPR006282,IPR007371,IPR00737 |
| 572 | HORVU.MOREX.r3.3HG03131 | low | protein_cod | --* | Ornithine c | not classifie | no mercato | GO:000458 | NA | NA |
| 573 | HORVU.MOREX.r3.3HG03138 | high | protein_cod | *-* | PsbP family | psbp domai | no mercato | GO:000550 | PF01789 | IPR002683,IPR016123,IPR016123 |
| 574 | HORVU.MOREX.r3.3HG03139 | high | protein_cod | *** | Methyl este | ab hydrolas | no mercato | NA | PF12697 | IPR029058,IPR000073 |
| 575 | HORVU.MOREX.r3.3HG03139 | high | protein_cod | *** | Methyl este | ab hydrolas | no mercato | NA | PF12697 | IPR029058,IPR000073 |
| 576 | HORVU.MOREX.r3.3HG03140 | high | protein_cod | *** | Methyl este | ab hydrolas | no mercato | NA | PF12697 | IPR029058,IPR000073 |
| 577 | HORVU.MOREX.r3.3HG03142 | high | protein_cod | *** | 2-oxoglutar | oxidoreduct | phytohorm | GO:001649 | PF03171,PF | IPR005123,IPR026992 |
| 578 | HORVU.MOREX.r3.3HG03147 | high | protein_cod | *** | Lipase | lipase 3 don | no mercato | GO:000662 | PF01764 | IPR002921,IPR029058 |
| 579 | HORVU.MOREX.r3.3HG03148 | low | protein_cod | --* | 26S proteas | not classifie | no mercato | GO:000050 | NA | NA |
| 580 | HORVU.MOREX.r3.3HG03161 | low | protein_cod | --* | Voltage-dep | Pepsin-type | protein hom | GO:000016 | NA | NA |
| 581 | HORVU.MOREX.r3.3HG03202 | high | protein_cod | *** | Glutathione | class tau gl | redox home | GO:001674 | PF00043,PF | IPR004046,IPR010987,IPR004045,IPR01233 |
| 582 | HORVU.MOREX.r3.3HG03220 | high | protein_cod | *** | Calcium-bir | calcium sen | multi-proce | GO:000550 | PF13499,PF | IPR002048,IPR002048,IPR011992 |
| 583 | HORVU.MOREX.r3.3HG03221 | high | protein_cod | *** | Calcium-bir | calcium sen | multi-proce | GO:000550 | PF13833,PF | IPR002048,IPR011992,IPR002048 |
| 584 | HORVU.MOREX.r3.3HG03221 | high | protein_cod | *** | Calcium-bir | calcium sen | multi-proce | GO:000550 | PF13499,PF | IPR011992,IPR002048,IPR002048 |

| | | | | | | | | | | |
|---|---|---|---|---|---|---|---|---|---|---|
| 585 | HORVU.MOREX.r3.3HG032252 | high | protein_cod | *** | U-box doma | u box domai | no mercato | NA | IPR016024 | |
| 586 | HORVU.MOREX.r3.3HG032265 | high | protein_cod | *** | NAD kinase | NAD kinase; | coenzyme n | GO:000395 | PF01513 | IPR016064,IPR002504 |
| 587 | HORVU.MOREX.r3.3HG032329 | low | protein_cod | --* | P-loop cont | g type lectin | no mercato | NA | NA | NA |
| 588 | HORVU.MOREX.r3.3HG032331 | high | protein_cod | *** | Serine/threo | g type lectin | no mercato | GO:000016 | PF01453 | IPR001480,IPR001480 |
| 589 | HORVU.MOREX.r3.3HG032358 | high | protein_cod | --* | Disease res | not classifie | no mercato | NA | NA | NA |
| 590 | HORVU.MOREX.r3.3HG032359 | low | non_coding | --* | 50S ribosom | not classifie | no mercato | GO:000373 | NA | NA |
| 591 | HORVU.MOREX.r3.3HG032407 | high | protein_cod | *** | Beta-1,3-glu | EC_3.2 glyc | enzyme clas | GO:000455 | PF00332 | IPR017853,IPR000490 |
| 592 | HORVU.MOREX.r3.3HG032445 | high | protein_cod | *** | Progesteron | s 8 oxocitro | no mercato | GO:000385 | IPR016040 | |
| 593 | HORVU.MOREX.r3.3HG032522 | low | protein_cod | --* | DegP protea | oberon cc d | no mercato | NA | NA | NA |
| 594 | HORVU.MOREX.r3.3HG032525 | high | protein_cod | *** | Peroxidase | EC_1.11 oxi | enzyme clas | GO:000460 | PF00141 | IPR010255,IPR002016 |
| 595 | HORVU.MOREX.r3.3HG032618 | low | protein_cod | --* | HIT zinc fing | not classifie | no mercato | NA | NA | NA |
| 596 | HORVU.MOREX.r3.3HG032931 | high | protein_cod | *** | Protein kina | RLCK-VIIa re | protein moc | GO:000016 | PF07714 | IPR011009,IPR001245 |
| 597 | HORVU.MOREX.r3.3HG032943 | high | protein_cod | *** | Plant-specif | 1 >3 beta glu | no mercato | NA | PF04720 | IPR006502,IPR006502 |
| 598 | HORVU.MOREX.r3.3HG032974 | low | non_coding | *** | Cytochrome | EC_1.14 oxi | enzyme clas | GO:000449 | PF00067 | IPR001128,IPR001128 |
| 599 | HORVU.MOREX.r3.3HG033003 | high | protein_cod | *** | Soluble inor | cytosolic py | multi-proce | GO:000028 | PF00719 | IPR008162,IPR008162 |
| 600 | HORVU.MOREX.r3.4HG033146 | high | protein_cod | *** | Calcium-de | c2 domain c | no mercato | NA | PF00168 | IPR000008,IPR000008 |
| 601 | HORVU.MOREX.r3.4HG033189 | high | protein_cod | *** | Haloacid de | pyrimidine p | coenzyme n | GO:000815 | PF13419 | IPR023214,IPR023214,IPR006439 |
| 602 | HORVU.MOREX.r3.4HG033354 | high | protein_cod | *** | Acetyltransf | n acetyltran | no mercato | GO:001674 | PF13302 | IPR000182,IPR016181 |
| 603 | HORVU.MOREX.r3.4HG033401 | high | protein_cod | --* | double-stra | branched ch | no mercato | NA | NA | NA |
| 604 | HORVU.MOREX.r3.4HG033516 | high | protein_cod | *** | Hedgehog-in | gsdh domai | no mercato | GO:000382 | PF07995 | IPR011041,IPR012938 |
| 605 | HORVU.MOREX.r3.4HG033532 | high | protein_cod | *** | transmemb | nfact r 1 dor | no mercato | NA | PF04398 | IPR007493,IPR007493 |
| 606 | HORVU.MOREX.r3.4HG033582 | high | protein_cod | *** | F-box protei | substrate ac | protein hom | NA | PF01476,PF | IPR018392,IPR018392,IPR001810,IPR001810 |
| 607 | HORVU.MOREX.r3.4HG033590 | high | protein_cod | --* | staurospori | not classifie | no mercato | NA | NA | NA |
| 608 | HORVU.MOREX.r3.4HG033591 | high | protein_cod | *** | Haloacid de | xanthosine r | nucleotide r | GO:000815 | IPR023214,IPR006439,IPR010237 | |
| 609 | HORVU.MOREX.r3.4HG033592 | low | protein_cod | *-* | Splicing fact | RNA splicing | rna process | GO:000035 | PF08799 | IPR014906,IPR014906 |
| 610 | HORVU.MOREX.r3.4HG033596 | high | protein_cod | *** | Haloacid de | xanthosine r | nucleotide r | GO:000815 | IPR010237,IPR023214,IPR006439 | |
| 611 | HORVU.MOREX.r3.4HG033626 | high | protein_cod | --* | sensitive to | not classifie | no mercato | NA | NA | NA |
| 612 | HORVU.MOREX.r3.4HG033687 | high | protein_cod | *** | Microsomal | microsomal | no mercato | GO:001602 | PF01124 | IPR001129 |
| 613 | HORVU.MOREX.r3.4HG033689 | high | protein_cod | *** | Basic blue p | phytocyanin | no mercato | GO:000905 | PF02298 | IPR008972,IPR003245 |
| 614 | HORVU.MOREX.r3.4HG033700 | high | protein_cod | *** | Regulator of | regulatory p | vesicle traff | GO:001503 | PF03398 | IPR005061 |
| 615 | HORVU.MOREX.r3.4HG033704 | high | protein_cod | *** | Accelerated | sphingosine | lipid metabo | GO:000573 | PF08718 | IPR014830,IPR014830 |
| 616 | HORVU.MOREX.r3.4HG033724 | high | protein_cod | *** | Glutamine s | cytosolic glu | nutrient upt | GO:000016 | PF03951,PF | IPR008147,IPR008147,IPR008146 |
| 617 | HORVU.MOREX.r3.4HG033783 | low | plastid_rela | *** | UV-B-induc | uv b induce | no mercato | NA | PF05542 | IPR008479 |
| 618 | HORVU.MOREX.r3.4HG033802 | high | protein_cod | *** | Alpha/beta- | hydrolase 4 | no mercato | GO:000662 | PF12146 | IPR022742,IPR029058 |
| 619 | HORVU.MOREX.r3.4HG033808 | high | protein_cod | *** | Peptide tran | anion transp | solute trans | GO:000521 | PF00854 | IPR000109,IPR020846,IPR020846 |
| 620 | HORVU.MOREX.r3.4HG033851 | low | protein_cod | --* | P-loop cont | 50s ribosom | no mercato | NA | NA | NA |
| 621 | HORVU.MOREX.r3.4HG033862 | high | protein_cod | *-* | Erythronate | not classifie | no mercato | GO:000815 | NA | NA |
| 622 | HORVU.MOREX.r3.4HG033864 | high | protein_cod | *** | Anthocyanin | EC_2.3 acyl | enzyme clas | GO:001674 | PF02458 | IPR003480 |
| 623 | HORVU.MOREX.r3.4HG033867 | high | protein_cod | --* | marker for o | expressed p | no mercato | NA | NA | NA |
| 624 | HORVU.MOREX.r3.4HG033993 | high | protein_cod | *** | Endoplasmi | thiol-disulfi | protein moc | GO:000375 | PF04137 | IPR007266 |
| 625 | HORVU.MOREX.r3.4HG033996 | high | protein_cod | -** | serine-type | not classifie | no mercato | NA | NA | NA |
| 626 | HORVU.MOREX.r3.4HG034003 | high | protein_cod | --* | beta-fructof | rnase h type | no mercato | NA | NA | NA |
| 627 | HORVU.MOREX.r3.4HG034004 | high | protein_cod | *** | BnaCnng07 | rnase h type | no mercato | NA | NA | NA |
| 628 | HORVU.MOREX.r3.4HG034055 | high | protein_cod | *** | Sorghum bic | histidinol ph | no mercato | NA | NA | NA |
| 629 | HORVU.MOREX.r3.4HG034068 | high | protein_cod | *** | Kinase PfkB | myo-inosito | multi-proce | NA | PF00294 | IPR029056,IPR011611 |
| 630 | HORVU.MOREX.r3.4HG034187 | high | protein_cod | --* | T-box transc | not classifie | no mercato | NA | NA | NA |
| 631 | HORVU.MOREX.r3.4HG034260 | high | protein_cod | *** | MAR-bindin | cyclin n tern | no mercato | NA | PF05542,PF | IPR008479,IPR008479 |
| 632 | HORVU.MOREX.r3.4HG034280 | high | protein_cod | *** | Zinc-finger p | ZAT transcri | rna biosynth | GO:000367 | PF13912,PF | IPR007087,IPR007087 |
| 633 | HORVU.MOREX.r3.4HG034286 | high | protein_cod | *** | Kinase fami | SnRK2 SNF1 | protein moc | GO:000016 | PF00069 | IPR011009,IPR000719 |
| 634 | HORVU.MOREX.r3.4HG034368 | high | protein_cod | *** | F-box/LRR p | substrate ac | protein hom | NA | PF13516,PF | IPR001611,IPR001611,IPR001611 |
| 635 | HORVU.MOREX.r3.4HG034399 | high | protein_cod | *** | F-box/kelch | substrate(P. | secondary n | NA | PF01344,PF | IPR001810,IPR006652,IPR001810 |
| 636 | HORVU.MOREX.r3.4HG034405 | high | protein_cod | *** | F-box/kelch | substrate(P. | secondary n | NA | PF00646,PF | IPR001810,IPR001810,IPR006652 |
| 637 | HORVU.MOREX.r3.4HG034406 | low | protein_cod | --* | Glucose-6-p | not classifie | no mercato | GO:000434 | NA | NA |
| 638 | HORVU.MOREX.r3.4HG034506 | low | protein_cod | --* | PHYTOENE | not classifie | no mercato | NA | NA | NA |
| 639 | HORVU.MOREX.r3.4HG034514 | high | protein_cod | *-* | Kinase, puta | Crinkly-like | protein moc | GO:000467 | PF00069 | IPR000719,IPR009091,IPR011009 |
| 640 | HORVU.MOREX.r3.4HG034584 | high | protein_cod | *** | Receptor le | EC_2.7 tran | enzyme clas | GO:000467 | PF00069,PF | IPR000719,IPR011009,IPR013320,IPR001220 |
| 641 | HORVU.MOREX.r3.4HG034661 | high | protein_cod | *-* | patatin-like | not classifie | no mercato | NA | NA | NA |
| 642 | HORVU.MOREX.r3.4HG034717 | high | protein_cod | *** | 1-aminocyc | 1-aminocyc | phytohormo | GO:001649 | PF14226,PF | IPR026992,IPR005123 |
| 643 | HORVU.MOREX.r3.4HG034736 | high | protein_cod | *** | NAC domain | NAC transcr | rna biosynth | GO:000367 | PF02365 | IPR003441,IPR003441 |
| 644 | HORVU.MOREX.r3.4HG034978 | high | protein_cod | *-* | F-box/LRR-r | f box domai | no mercato | NA | PF13516 | IPR001611 |
| 645 | HORVU.MOREX.r3.4HG034979 | low | protein_cod | --* | Peroxin-6 | f box domai | no mercato | GO:000552 | NA | NA |
| 646 | HORVU.MOREX.r3.4HG035077 | high | protein_cod | *** | Non-specifi | SnRK3 SNF1 | protein moc | GO:000016 | PF00069,PF | IPR000719,IPR011009,IPR004041 |
| 647 | HORVU.MOREX.r3.4HG035094 | high | protein_cod | *** | Basic blue p | phytocyanin | no mercato | GO:000905 | PF02298 | IPR008972,IPR003245 |
| 648 | HORVU.MOREX.r3.4HG035107 | low | protein_cod | --* | STRUBBELIG | proline rich | no mercato | NA | NA | NA |
| 649 | HORVU.MOREX.r3.4HG035132 | high | protein_cod | *** | F-box protei | f box domai | no mercato | GO:001602 | PF12937 | IPR001810,IPR001810 |

| | | | | | | | | | | |
|---|---|---|---|---|---|---|---|---|---|---|
| 650 | HORVU.MOREX.r3.4HG03517 | high | protein_cod | --* | Calcium-ac | not classifie | no mercato | GO:000166 | NA | NA |
| 651 | HORVU.MOREX.r3.4HG03518 | high | protein_cod | *** | Peroxidase | EC_1.11 oxi | enzyme clas | GO:000460 | PF00141 | IPR002016,IPR010255 |
| 652 | HORVU.MOREX.r3.4HG03520 | high | protein_cod | *** | Saposin B d | saposin b ty | no mercato | GO:000662 | PF03489,PF | IPR008138,IPR011001,IPR007856,IPR007856,IPR011001 |
| 653 | HORVU.MOREX.r3.4HG03527 | high | protein_cod | *** | WRKY trans | transcriptio | rna biosynth | GO:000367 | PF03106 | IPR003657,IPR003657 |
| 654 | HORVU.MOREX.r3.4HG03561 | high | protein_cod | *** | Calcium-bir | calcium sen | multi-proce | GO:000550 | PF13202,PF | IPR002048,IPR011992,IPR002048 |
| 655 | HORVU.MOREX.r3.4HG03562 | high | protein_cod | *** | Serine/threc | SD-2 protei | protein moc | GO:000016 | PF01453,PF | IPR001480,IPR011009,IPR000858,IPR001480,IPR003609,IPR00071 |
| 656 | HORVU.MOREX.r3.4HG03564 | high | protein_cod | *** | Non-specifi | SnRK3 SNF | protein moc | GO:000016 | PF00069,PF | IPR011009,IPR000719,IPR028375,IPR004041 |
| 657 | HORVU.MOREX.r3.4HG03565 | high | protein_cod | *** | Protein ZIN | metal chela | solute trans | GO:000521 | PF07690 | IPR011701,IPR020846 |
| 658 | HORVU.MOREX.r3.4HG03573 | high | protein_cod | *** | Proline tran | proline tran | amino acid | GO:001602 | PF01490 | IPR013057 |
| 659 | HORVU.MOREX.r3.4HG03579 | high | protein_cod | *** | Metallothio | metallothio | external stir | GO:004687 | PF01439 | IPR000347 |
| 660 | HORVU.MOREX.r3.4HG03611 | high | protein_cod | *** | 3-ketoacyl- | 3-ketoacyl- | lipid metab | GO:000382 | PF08392,PF | IPR016039,IPR016039,IPR013601,IPR016039,IPR013747 |
| 661 | HORVU.MOREX.r3.4HG03756 | high | protein_cod | *-* | Heavy meta | hma domair | no mercato | GO:003000 | PF00403 | IPR006121,IPR006121 |
| 662 | HORVU.MOREX.r3.4HG03785 | low | transposon_ | *-* | Retrotransp | duf 4219 do | no mercato | GO:000367 | NA | NA |
| 663 | HORVU.MOREX.r3.4HG03792 | high | protein_cod | *** | Calcium bin | calcium sen | multi-proce | GO:000550 | PF13499,PF | IPR002048,IPR002048,IPR011992 |
| 664 | HORVU.MOREX.r3.4HG03797 | high | protein_cod | *** | NAC domai | NAC transc | rna biosynth | GO:000367 | PF02365 | IPR003441,IPR003441 |
| 665 | HORVU.MOREX.r3.4HG03797 | high | protein_cod | *** | Thioredoxin | atypical thic | redox home | GO:000562 | PF00085 | IPR013766,IPR012336 |
| 666 | HORVU.MOREX.r3.4HG03797 | high | protein_cod | *** | Phospholipi | active comp | lipid metab | GO:000016 | PF16209,PF | IPR032631,IPR023299,IPR023299,IPR023214,IPR023214,IPR008250,IPR032630,IPR001757,IPR001757,IPR00653 |
| 667 | HORVU.MOREX.r3.4HG03801 | high | protein_cod | *** | Chaperone | sp dnaj cha | no mercato | NA | PF00226 | IPR001623,IPR001623 |
| 668 | HORVU.MOREX.r3.4HG03802 | high | protein_cod | *-* | serine-rich | serine rich p | no mercato | NA | NA | NA |
| 669 | HORVU.MOREX.r3.4HG03809 | high | protein_cod | *** | MYB transc | MYB class-F | rna biosynth | GO:000367 | PF00249,PF | IPR009057,IPR001005,IPR001005 |
| 670 | HORVU.MOREX.r3.4HG03817 | high | protein_cod | *** | Calmodulin | calcium bin | no mercato | GO:000550 | PF13499,PF | IPR002048,IPR002048,IPR011992 |
| 671 | HORVU.MOREX.r3.4HG03826 | high | protein_cod | *** | Xyloglucan | 1,6-alpha-x | cell wall org | GO:001602 | PF05637 | IPR008630 |
| 672 | HORVU.MOREX.r3.4HG03827 | high | protein_cod | *** | Universal st | usp domain | no mercato | GO:000577 | PF00582 | IPR006016 |
| 673 | HORVU.MOREX.r3.4HG03827 | high | protein_cod | *** | Multidrug re | multidrug re | no mercato | NA | PF14009 | IPR025322 |
| 674 | HORVU.MOREX.r3.4HG03832 | high | protein_cod | -** | Peroxisoma | peroxisoma | cell division | GO:000367 | PF05648 | IPR008733 |
| 675 | HORVU.MOREX.r3.4HG03835 | high | protein_cod | *** | Pathogenes | bet v 1 dom | no mercato | GO:000695 | PF00407 | IPR000916 |
| 676 | HORVU.MOREX.r3.4HG03836 | high | protein_cod | *** | Short-chain | short chain | no mercato | NA | IPR016040 | |
| 677 | HORVU.MOREX.r3.4HG03838 | high | protein_cod | *** | DUF538 fan | cpsase sm | no mercato | NA | PF04398 | IPR007493,IPR007493 |
| 678 | HORVU.MOREX.r3.4HG03839 | high | protein_cod | *** | DUF1262 fa | toxin 10 dor | no mercato | NA | PF06880 | IPR010683 |
| 679 | HORVU.MOREX.r3.4HG03847 | high | protein_cod | *** | Asparagine | glutamine-d | amino acid | NA | PF13537,PF | IPR017932,IPR006426,IPR001962,IPR02905 |
| 680 | HORVU.MOREX.r3.4HG03849 | high | protein_cod | *** | Omega-3 fa | delta-12/de | lipid metab | GO:000662 | PF00487,PF | IPR005804,IPR021863 |
| 681 | HORVU.MOREX.r3.4HG03856 | low | protein_cod | --* | Hydrophobi | stress induc | no mercato | GO:001602 | NA | NA |
| 682 | HORVU.MOREX.r3.4HG03859 | high | protein_cod | *** | RING/U-box | RING-H2-cl | protein hom | GO:001602 | PF13639 | IPR001841 |
| 683 | HORVU.MOREX.r3.4HG03860 | high | protein_cod | *-* | transmemb | cchc type d | no mercato | NA | NA | NA |
| 684 | HORVU.MOREX.r3.4HG03860 | high | protein_cod | *** | Glutathione | class lambd | redox home | GO:001674 | PF13417 | IPR004045,IPR010987,IPR012336 |
| 685 | HORVU.MOREX.r3.4HG03861 | high | protein_cod | *** | Glutathione | class lambd | redox home | GO:001674 | PF13417 | IPR012336,IPR010987,IPR004045 |
| 686 | HORVU.MOREX.r3.4HG03870 | high | protein_cod | *** | Zinc finger f | ZAT transcri | rna biosynth | GO:000367 | PF13912,PF | IPR007087,IPR007087 |
| 687 | HORVU.MOREX.r3.4HG03884 | high | protein_cod | *** | Haloacid de | haloacid de | no mercato | GO:000815 | IPR010237,IPR023214,IPR023214,IPR006439 | |
| 688 | HORVU.MOREX.r3.4HG03886 | high | protein_cod | *** | RING/U-box | RING-H2-cl | protein hom | NA | PF06547,PF | IPR010543,IPR001841 |
| 689 | HORVU.MOREX.r3.4HG03887 | high | protein_cod | *** | Hsp70 nucl | nucleotide | protein hom | NA | PF08609 | IPR013918,IPR016024 |
| 690 | HORVU.MOREX.r3.4HG03888 | high | protein_cod | *** | Avr9/Cf-9 ra | duf 668 dom | no mercato | NA | PF11961,PF | IPR021864,IPR007700 |
| 691 | HORVU.MOREX.r3.4HG03889 | high | protein_cod | *** | Beta-1,3-N- | legb domair | no mercato | GO:001602 | PF04646 | IPR006740 |
| 692 | HORVU.MOREX.r3.4HG03890 | high | protein_cod | *** | Protein pho | clade A pho | protein moc | GO:000382 | PF00481 | IPR001932,IPR001932,IPR001932 |
| 693 | HORVU.MOREX.r3.4HG03892 | high | protein_cod | *** | lipase, puta | GTPase effe | multi-proce | NA | PF04788 | IPR006873 |
| 694 | HORVU.MOREX.r3.4HG03893 | high | protein_cod | *** | Heat-shock | class-C-I sn | protein hom | NA | PF00011 | IPR002068,IPR008978 |
| 695 | HORVU.MOREX.r3.4HG03894 | high | protein_cod | *** | Heat-shock | class-C-I sn | protein hom | NA | PF00011 | IPR008978,IPR002068 |
| 696 | HORVU.MOREX.r3.4HG03894 | high | protein_cod | *** | Maternal ef | 60s ribosom | no mercato | NA | NA | NA |
| 697 | HORVU.MOREX.r3.4HG03895 | high | protein_cod | *** | Transmemb | membrane | no mercato | GO:001602 | PF06127 | IPR009305 |
| 698 | HORVU.MOREX.r3.4HG03895 | high | protein_cod | *-* | RNA-bindin | RNA splicin | rna process | GO:000367 | PF00076 | IPR000504,IPR012677 |
| 699 | HORVU.MOREX.r3.4HG03896 | high | protein_cod | *** | Josephin, p | ubiquitinyl h | no mercato | GO:000484 | PF02099 | IPR006155 |
| 700 | HORVU.MOREX.r3.4HG03910 | high | protein_cod | *** | Phosphoen | phosphoen | carbohydra | GO:000461 | PF01293 | IPR001272,IPR008210,IPR001272 |
| 701 | HORVU.MOREX.r3.4HG03911 | high | protein_cod | *** | Patatin | phospholip | lipid metab | GO:000662 | PF01734 | IPR016035,IPR002641 |
| 702 | HORVU.MOREX.r3.4HG03912 | high | protein_cod | --* | Clathrin hea | calcium bin | no mercato | NA | NA | NA |
| 703 | HORVU.MOREX.r3.4HG03914 | high | protein_cod | *** | alpha/beta- | abhydrolase | no mercato | NA | PF07859 | IPR029058,IPR013094 |
| 704 | HORVU.MOREX.r3.4HG03916 | high | protein_cod | *** | Calcium-bir | SCS-clade | multi-proce | GO:000550 | PF13202 | IPR002048,IPR011992,IPR011992 |
| 705 | HORVU.MOREX.r3.4HG03916 | high | protein_cod | -** | Oxidoreduc | dna cytosin | no mercato | GO:001602 | NA | NA |
| 706 | HORVU.MOREX.r3.4HG03922 | high | protein_cod | *** | Heat shock | class-P sma | protein hom | NA | PF00011 | IPR002068,IPR008978 |
| 707 | HORVU.MOREX.r3.4HG03924 | high | protein_cod | *** | Armadillo re | regulatory p | multi-proce | NA | PF00514 | IPR016024,IPR016024,IPR000225 |
| 708 | HORVU.MOREX.r3.4HG03928 | high | protein_cod | *** | Senescence | senescence | no mercato | NA | PF06911 | IPR009686 |
| 709 | HORVU.MOREX.r3.4HG03929 | high | protein_cod | *** | Kinase fami | RLCK-VI rec | protein moc | GO:000467 | PF00069,PF | IPR000719,IPR011009,IPR006016 |
| 710 | HORVU.MOREX.r3.4HG03930 | high | protein_cod | *** | U-box doma | E3 ubiquitin | protein hom | GO:000484 | PF04564 | IPR003613,IPR016024 |
| 711 | HORVU.MOREX.r3.4HG03936 | high | protein_cod | *** | Glutamate | glutamate d | amino acid | GO:000382 | PF00282 | IPR015424,IPR002129,IPR010107 |
| 712 | HORVU.MOREX.r3.4HG03945 | high | protein_cod | *** | Poly [ADP-ri | ADP-ribosyl | protein moc | GO:000395 | PF12174 | IPR022003 |
| 713 | HORVU.MOREX.r3.4HG03949 | high | protein_cod | *** | Non-symbic | goshi non s | no mercato | GO:000534 | PF00042 | IPR000971,IPR009050 |
| 714 | HORVU.MOREX.r3.4HG03949 | high | protein_cod | *** | Kinase, puta | WAK/WAKL | protein moc | GO:000016 | PF00069 | IPR011009,IPR000719 |

| | | | | | | | | | | |
|---|---|---|---|---|---|---|---|---|---|---|
| 715 | HORVU.MOREX.r3.4HG039513 | high | protein_cod | *** | Cyclin, puta | regulatory p | cell division | GO:000563 | PF00134,PF | IPR006671,IPR004367,IPR013763,IPR013763 |
| 716 | HORVU.MOREX.r3.4HG039523 | high | protein_cod | *** | Major facilit | UMF15-type | solute trans | GO:001602 | PF07690 | IPR020846,IPR011701 |
| 717 | HORVU.MOREX.r3.4HG039554 | high | protein_cod | *** | Glutamine s | cytosolic glu | nutrient upt | GO:000016 | PF00120,PF | IPR008146,IPR008147,IPR008147 |
| 718 | HORVU.MOREX.r3.4HG039613 | high | protein_cod | *** | Sugar trans | monosacch | solute trans | GO:000521 | PF00083 | IPR003663,IPR005828,IPR020846 |
| 719 | HORVU.MOREX.r3.4HG039701 | high | protein_cod | *** | Beta-glucos | EC_3.2 glyc | enzyme clas | GO:000455 | PF00232 | IPR017853,IPR001360 |
| 720 | HORVU.MOREX.r3.4HG039709 | high | protein_cod | --* | Envelope gl | alkyl hydrop | no mercato | GO:001602 | NA | NA |
| 721 | HORVU.MOREX.r3.4HG039757 | high | protein_cod | *** | Glucan end | EC_3.2 glyc | enzyme clas | GO:000455 | PF07983,PF | IPR012946,IPR017853,IPR000490 |
| 722 | HORVU.MOREX.r3.4HG039816 | high | protein_cod | --* | S-adenosyl | cchc type d | no mercato | GO:000209 | NA | NA |
| 723 | HORVU.MOREX.r3.4HG039857 | high | protein_cod | *** | Sigma facto | strigolacton | phytohormo | GO:001678 | PF12697 | IPR000073,IPR029058 |
| 724 | HORVU.MOREX.r3.4HG039896 | high | protein_cod | *** | Bifunctional | not classifie | no mercato | GO:001602 | NA | NA |
| 725 | HORVU.MOREX.r3.4HG039924 | high | protein_cod | *** | Homeobox | HD-ZIP I/II-t | rna biosynth | GO:000367 | PF00046 | IPR009057,IPR001356 |
| 726 | HORVU.MOREX.r3.4HG039970 | high | protein_cod | *** | Serine acet | serine O-ac | amino acid | GO:000573 | PF06426,PF | IPR011004,IPR005881,IPR010493,IPR001451 |
| 727 | HORVU.MOREX.r3.4HG039997 | low | non_coding | *** | Cytochrome | cytochrome | no mercato | GO:001602 | PF04526,PF | IPR005018,IPR006593 |
| 728 | HORVU.MOREX.r3.4HG040058 | high | protein_cod | *-* | Protein pho | protein pho | no mercato | GO:000382 | PF07228 | IPR001932,IPR001932 |
| 729 | HORVU.MOREX.r3.4HG040078 | high | protein_cod | *** | Heavy meta | hma domain | no mercato | GO:003000 | PF00403,PF | IPR006121,IPR006121,IPR006121,IPR006121 |
| 730 | HORVU.MOREX.r3.4HG040086 | high | protein_cod | *** | D-tagatose- | expressed p | no mercato | NA | NA | NA |
| 731 | HORVU.MOREX.r3.4HG040095 | high | protein_cod | *** | 3-hydroxybe | EC_1.14 oxi | enzyme clas | GO:007194 | PF01494 | IPR002938,IPR023753,IPR023753 |
| 732 | HORVU.MOREX.r3.4HG040098 | high | protein_cod | *** | 3-hydroxybe | EC_1.14 oxi | enzyme clas | GO:007194 | PF01494 | IPR002938,IPR023753,IPR023753 |
| 733 | HORVU.MOREX.r3.4HG040099 | high | protein_cod | *** | 3-hydroxybe | EC_1.14 oxi | enzyme clas | GO:007194 | PF01494 | IPR002938,IPR023753,IPR023753 |
| 734 | HORVU.MOREX.r3.4HG040107 | high | protein_cod | *** | Expansin | alpha-class | cell wall org | GO:000557 | PF01357,PF | IPR009009,IPR007117,IPR007117,IPR009009 |
| 735 | HORVU.MOREX.r3.4HG040117 | high | protein_cod | *** | Blue copper | phytocyanin | no mercato | GO:000905 | PF02298 | IPR008972,IPR003245 |
| 736 | HORVU.MOREX.r3.4HG040118 | high | protein_cod | *** | Blue copper | phytocyanin | no mercato | GO:000905 | PF02298 | IPR008972,IPR003245 |
| 737 | HORVU.MOREX.r3.4HG040119 | high | protein_cod | *** | Blue copper | phytocyanin | no mercato | GO:000905 | PF02298 | IPR008972,IPR003245 |
| 738 | HORVU.MOREX.r3.4HG040120 | high | protein_cod | *** | Blue copper | phytocyanin | no mercato | GO:000905 | PF02298 | IPR003245,IPR008972 |
| 739 | HORVU.MOREX.r3.4HG040121 | high | protein_cod | *** | Blue copper | phytocyanin | no mercato | GO:000905 | PF02298 | IPR008972,IPR003245 |
| 740 | HORVU.MOREX.r3.4HG040198 | high | protein_cod | *** | Sulfate tran | sulfate tran | solute trans | GO:000827 | PF00916,PF | IPR011547,IPR001902,IPR002645,IPR002645 |
| 741 | HORVU.MOREX.r3.4HG040231 | high | protein_cod | *** | DUF506 fan | duf 506 fam | no mercato | NA | PF04720 | IPR006502,IPR006502 |
| 742 | HORVU.MOREX.r3.4HG040310 | high | protein_cod | *** | DUF1997 fa | exonucleas | no mercato | NA | PF09366 | IPR018971 |
| 743 | HORVU.MOREX.r3.4HG040311 | high | protein_cod | *** | plant/protei | duf 3110 do | no mercato | NA | PF11360 | IPR021503 |
| 744 | HORVU.MOREX.r3.4HG040312 | high | protein_cod | *-* | Late embryo | not classifie | no mercato | NA | PF02987 | IPR004238 |
| 745 | HORVU.MOREX.r3.4HG040362 | high | protein_cod | *** | Sucrose tra | sugar trans | solute trans | GO:000588 | IPR005989,IPR020846 | |
| 746 | HORVU.MOREX.r3.4HG040381 | high | protein_cod | --* | Ascorbate-s | not classifie | no mercato | GO:000573 | NA | NA |
| 747 | HORVU.MOREX.r3.4HG040468 | high | protein_cod | *** | NAD(P)H de | EC_1.6 oxid | enzyme clas | GO:000395 | PF03358 | IPR005025,IPR029039,IPR010089 |
| 748 | HORVU.MOREX.r3.4HG040524 | high | protein_cod | *** | Jasmonate | component | phytohormo | NA | PF06200,PF | IPR010399,IPR018467 |
| 749 | HORVU.MOREX.r3.4HG040561 | low | protein_cod | --* | Voltage-dep | ap erf doma | no mercato | GO:000521 | NA | NA |
| 750 | HORVU.MOREX.r3.4HG040599 | high | protein_cod | *** | Transmemb | phytocyanin | no mercato | GO:001602 | NA | NA |
| 751 | HORVU.MOREX.r3.4HG040632 | low | protein_cod | --* | RING/FYVE/ | ubiquitin ca | no mercato | GO:004687 | NA | NA |
| 752 | HORVU.MOREX.r3.4HG040638 | high | protein_cod | *** | GEM-like pr | gram domai | no mercato | NA | PF02893 | IPR004182 |
| 753 | HORVU.MOREX.r3.4HG040639 | high | protein_cod | *** | Purine perm | organic cati | solute trans | GO:000521 | NA | NA |
| 754 | HORVU.MOREX.r3.4HG040642 | high | protein_cod | *** | Purine perm | organic cati | solute trans | GO:000521 | NA | NA |
| 755 | HORVU.MOREX.r3.4HG040645 | high | protein_cod | *** | Purine perm | organic cati | solute trans | GO:000521 | NA | NA |
| 756 | HORVU.MOREX.r3.4HG040671 | high | protein_cod | *-* | Homeobox | HD-ZIP I/II-t | rna biosynth | GO:000367 | PF00046,PF | IPR009057,IPR001356,IPR003106 |
| 757 | HORVU.MOREX.r3.4HG040672 | low | protein_cod | --* | nuclear pol | not classifie | no mercato | NA | NA | NA |
| 758 | HORVU.MOREX.r3.4HG040698 | high | protein_cod | *** | Ras-related | H-class RAB | vesicle traff | GO:000392 | PF00071 | IPR005225,IPR027417,IPR001806 |
| 759 | HORVU.MOREX.r3.4HG040762 | low | plastid_rela | *** | Pheophorbi | pheophorbi | coenzyme n | GO:000821 | PF00355,PF | IPR017941,IPR017941,IPR013626 |
| 760 | HORVU.MOREX.r3.4HG040790 | high | protein_cod | --* | Na(+)/H(+) | putative sili | solute trans | GO:000588 | PF03600 | IPR004680 |
| 761 | HORVU.MOREX.r3.4HG040806 | high | protein_cod | *** | Glycosyltra | EC_2.4 glyc | enzyme clas | GO:000815 | PF00201 | IPR002213 |
| 762 | HORVU.MOREX.r3.4HG040880 | high | protein_cod | --* | Peptide met | peptide met | no mercato | GO:000697 | IPR011057 | |
| 763 | HORVU.MOREX.r3.4HG040944 | high | protein_cod | *** | LysM domai | chitin recep | external stir | GO:001602 | PF01476,PF | IPR018392,IPR018392,IPR018392,IPR018392 |
| 764 | HORVU.MOREX.r3.4HG040987 | high | protein_cod | *** | Expansin-lik | alpha-like-c | cell wall org | GO:000557 | PF03330,PF | IPR009009,IPR007117,IPR007117,IPR009009 |
| 765 | HORVU.MOREX.r3.4HG041121 | high | protein_cod | *** | Beta-carote | carotenoid | secondary n | GO:000550 | PF04116 | IPR006694 |
| 766 | HORVU.MOREX.r3.4HG041209 | high | protein_cod | *** | C3HC4-type | voltage dep | no mercato | NA | PF17123,PF | IPR001841,IPR002035,IPR002035,IPR032838 |
| 767 | HORVU.MOREX.r3.4HG041212 | high | protein_cod | *** | Zinc finger ( | voltage dep | no mercato | NA | PF14624,PF | IPR032838,IPR002035,IPR002035,IPR002035,IPR001841 |
| 768 | HORVU.MOREX.r3.4HG041215 | high | protein_cod | *** | Beta-amyla | beta amylas | carbohydrat | NA | PF01373 | IPR001554,IPR017853 |
| 769 | HORVU.MOREX.r3.4HG041234 | high | protein_cod | *** | Alpha/beta | mental dom | no mercato | GO:001602 | NA | NA |
| 770 | HORVU.MOREX.r3.4HG041265 | high | protein_cod | *** | TVP38/TMEI | transmemb | no mercato | GO:001602 | PF09335 | IPR032816 |
| 771 | HORVU.MOREX.r3.4HG041391 | high | protein_cod | *** | Cell wall inv | cell wall aci | carbohydrat | GO:000455 | PF00251,PF | IPR013320,IPR013320,IPR013148,IPR013189,IPR023296 |
| 772 | HORVU.MOREX.r3.4HG041400 | high | protein_cod | *** | Short-chain | short chain | no mercato | NA | PF00106 | IPR002347,IPR016040 |
| 773 | HORVU.MOREX.r3.4HG041403 | high | protein_cod | *** | Short-chain | short chain | no mercato | NA | PF00106,PF | IPR016040,IPR002347,IPR002347 |
| 774 | HORVU.MOREX.r3.4HG041533 | low | non_coding | *** | Cytochrome | cinnamate 4 | secondary n | GO:000449 | PF00067 | IPR001128,IPR001128 |
| 775 | HORVU.MOREX.r3.4HG041716 | high | protein_cod | *** | Flavin-conta | EC_1.8 oxid | enzyme clas | GO:000449 | PF00743,PF | IPR023753,IPR020946,IPR020946,IPR023753 |
| 776 | HORVU.MOREX.r3.4HG041741 | high | protein_cod | --* | Cold regula | night-time r | multi-proce | GO:000563 | NA | NA |
| 777 | HORVU.MOREX.r3.5HG042048 | high | protein_cod | *** | BEL1-like h | BEL transcri | rna biosynth | GO:000367 | PF05920,PF | IPR008422,IPR006563,IPR009057 |
| 778 | HORVU.MOREX.r3.5HG042131 | high | protein_cod | *** | Receptor pr | PIP/PIPL pe | phytohormo | GO:000016 | PF08263,PF | IPR013210,IPR000719,IPR032675,IPR032675,IPR001611,IPR001611,IPR032675,IPR001611,IPR011009 |
| 779 | HORVU.MOREX.r3.5HG042213 | high | protein_cod | *** | Ras family p | F-class RAB | vesicle traff | GO:000392 | PF00071 | IPR027417,IPR005225,IPR001806 |

| | | | | | | | | | | |
|---|---|---|---|---|---|---|---|---|---|---|
| 780 | HORVU.MOREX.r3.5HG04240 | high | protein_cod | *** | Phytoene sy | phytoene sy | secondary n | GO:000431 | PF00494 | IPR002060,IPR008949 |
| 781 | HORVU.MOREX.r3.5HG04245 | high | protein_cod | *** | Glutathione | class phi glu | redox home | GO:001674 | PF00043,PF | IPR004046,IPR010987,IPR012336,IPR004045 |
| 782 | HORVU.MOREX.r3.5HG04248 | high | protein_cod | *** | Glutathione | class phi glu | redox home | GO:001674 | PF00043,PF | IPR004046,IPR012336,IPR010987,IPR004045 |
| 783 | HORVU.MOREX.r3.5HG04251 | high | protein_cod | *** | Cationic am | cationic am | solute trans | GO:001602 | PF13520,PF | IPR002293,IPR029485 |
| 784 | HORVU.MOREX.r3.5HG04262 | high | protein_cod | *** | GTP binding | ribosome as | protein bios | GO:000552 | PF01926 | IPR027417,IPR006073 |
| 785 | HORVU.MOREX.r3.5HG04267 | high | protein_cod | *** | Cation/H(+) | proton:mon | solute trans | GO:000681 | PF00999 | IPR006153 |
| 786 | HORVU.MOREX.r3.5HG04278 | low | protein_cod | --* | Sterile alpha | not classifie | no mercator | NA | NA | NA |
| 787 | HORVU.MOREX.r3.5HG04331 | high | protein_cod | *** | Glycosyltrar | EC_2.4 glyc | enzyme clas | GO:000815 | NA | NA |
| 788 | HORVU.MOREX.r3.5HG04358 | low | protein_cod | --* | Leucine-rich | NA | NA | NA | NA | NA |
| 789 | HORVU.MOREX.r3.5HG04392 | high | protein_cod | *** | Zinc finger p | ZFP transcri | rna biosynth | GO:000367 | NA | NA |
| 790 | HORVU.MOREX.r3.5HG04408 | high | protein_cod | *** | APOLLO | DNA exonuc | chromatin o | GO:000367 | PF00929 | IPR012337,IPR013520 |
| 791 | HORVU.MOREX.r3.5HG04422 | high | protein_cod | *** | Receptor-lik | EC_2.7 tran | enzyme clas | GO:000467 | PF12819,PF | IPR024788,IPR032675,IPR001245,IPR001611,IPR01100 |
| 792 | HORVU.MOREX.r3.5HG04424 | high | protein_cod | *** | UNC93-like | protein unc | no mercator | GO:001602 | PF05978 | IPR020846,IPR010291 |
| 793 | HORVU.MOREX.r3.5HG04439 | low | protein_cod | --* | Peroxisome | not classifie | no mercator | GO:000110 | NA | NA |
| 794 | HORVU.MOREX.r3.5HG04440 | high | protein_cod | *** | Pathogenes | bet v 1 dom | no mercator | GO:000695 | PF00407 | IPR000916 |
| 795 | HORVU.MOREX.r3.5HG04441 | high | protein_cod | *** | Pathogenes | bet v 1 dom | no mercator | GO:000695 | NA | NA |
| 796 | HORVU.MOREX.r3.5HG04442 | high | protein_cod | *** | Pathogenes | bet v 1 dom | no mercator | GO:000695 | PF00407 | IPR000916 |
| 797 | HORVU.MOREX.r3.5HG04446 | high | protein_cod | *** | Universal st | usp domain | no mercator | GO:000695 | PF00582 | IPR006016 |
| 798 | HORVU.MOREX.r3.5HG04464 | high | protein_cod | *** | GDSL ester | gdsl esteras | no mercator | GO:001678 | PF00657 | IPR001087,IPR013830,IPR013830,IPR01383 |
| 799 | HORVU.MOREX.r3.5HG04486 | high | protein_cod | *-* | Plant calmo | cam binding | no mercator | NA | PF07839 | IPR012417 |
| 800 | HORVU.MOREX.r3.5HG04507 | low | non_coding | *** | ATP-depend | chaperone | protein hom | GO:000016 | PF02151,PF | IPR004176,IPR001943,IPR019489,IPR027417,IPR027417,IPR004176,IPR004176,IPR003959,IPR00395 |
| 801 | HORVU.MOREX.r3.5HG04536 | high | protein_cod | *** | Pyruvate de | subunit alpl | amino acid | GO:000473 | PF00676 | IPR029061,IPR001017 |
| 802 | HORVU.MOREX.r3.5HG04550 | high | protein_cod | *-* | Ethylene-re | subgroup EF | rna biosynth | GO:000367 | PF00847 | IPR016177,IPR001471 |
| 803 | HORVU.MOREX.r3.5HG04582 | low | non_coding | --* | 30S ribosom | proline rich | no mercator | GO:000373 | NA | NA |
| 804 | HORVU.MOREX.r3.5HG04582 | high | protein_cod | *** | HVA22-like | hva protein | no mercator | GO:001602 | PF03134 | IPR004345 |
| 805 | HORVU.MOREX.r3.5HG04588 | high | protein_cod | *-* | senescence | oj000126 1 | no mercator | NA | PF04520 | IPR007608 |
| 806 | HORVU.MOREX.r3.5HG04594 | high | protein_cod | *** | Plant/T32M | duf 668 don | no mercator | NA | PF05003,PF | IPR007700,IPR021864 |
| 807 | HORVU.MOREX.r3.5HG04612 | high | protein_cod | *** | Calcium-bir | calcium sen | multi-proce | GO:000550 | PF13202,PF | IPR002048,IPR011992,IPR002048 |
| 808 | HORVU.MOREX.r3.5HG04619 | low | non_coding | *-* | Cytochrome | EC_1.14 oxi | enzyme clas | GO:000382 | PF00067 | IPR001128,IPR001128 |
| 809 | HORVU.MOREX.r3.5HG04634 | high | protein_cod | *-* | NAC domai | NAC transcr | rna biosynth | GO:000367 | PF02365 | IPR003441,IPR003441 |
| 810 | HORVU.MOREX.r3.5HG04634 | high | protein_cod | *** | NAC domai | NAC transcr | rna biosynth | GO:000367 | PF02365 | IPR003441,IPR003441 |
| 811 | HORVU.MOREX.r3.5HG04641 | high | protein_cod | *** | BTB/POZ do | substrate a | protein hom | NA | PF00651,PF | IPR011333,IPR000210,IPR027356 |
| 812 | HORVU.MOREX.r3.5HG04644 | low | protein_cod | --* | apyrase 1 | not classifie | no mercator | NA | NA | NA |
| 813 | HORVU.MOREX.r3.5HG04645 | high | protein_cod | *** | Methyltrans | engb type g | no mercator | NA | NA | NA |
| 814 | HORVU.MOREX.r3.5HG04648 | high | protein_cod | *** | Glycosyltrar | EC_2.4 glyc | enzyme clas | GO:000815 | PF00201 | IPR002213 |
| 815 | HORVU.MOREX.r3.5HG04653 | high | protein_cod | *** | Nodulin-like | UMF23-type | solute trans | GO:001602 | PF06813 | IPR010658,IPR020846,IPR020846 |
| 816 | HORVU.MOREX.r3.5HG04654 | high | protein_cod | *** | LOB domair | AS2/LOB tra | rna biosynth | NA | PF03195 | IPR004883 |
| 817 | HORVU.MOREX.r3.5HG04655 | high | protein_cod | *** | Ferritin | iron storage | nutrient upt | NA | PF00210 | IPR008331,IPR009078 |
| 818 | HORVU.MOREX.r3.5HG04656 | low | protein_cod | --* | S-adenosyl- | tyrosinase c | no mercator | NA | NA | NA |
| 819 | HORVU.MOREX.r3.5HG04658 | high | protein_cod | *** | Calcium-bir | calcium bin | no mercator | GO:000550 | PF13833,PF | IPR002048,IPR011992,IPR002048 |
| 820 | HORVU.MOREX.r3.5HG04658 | high | protein_cod | *** | Asparagine | glutamine-d | amino acid | GO:000016 | PF13537,PF | IPR029055,IPR006426,IPR017932,IPR001962 |
| 821 | HORVU.MOREX.r3.5HG04673 | high | protein_cod | *** | Autophagy-r | autophagos | protein hom | GO:000691 | PF02991 | IPR029071,IPR004241 |
| 822 | HORVU.MOREX.r3.5HG04677 | high | protein_cod | --* | nodulin MtN | not classifie | no mercator | NA | NA | NA |
| 823 | HORVU.MOREX.r3.5HG04683 | high | protein_cod | *-* | Inosine-5'-n | regulatory p | redox home | NA | PF00571,PF | IPR000644,IPR000644 |
| 824 | HORVU.MOREX.r3.5HG04686 | high | protein_cod | *** | ATP-depend | atp depend | no mercator | NA | NA | NA |
| 825 | HORVU.MOREX.r3.5HG04688 | high | protein_cod | *** | Plastid-lipid | Fibrillin plas | lipid metab | NA | PF04755 | IPR006843 |
| 826 | HORVU.MOREX.r3.5HG04697 | high | protein_cod | *-* | RING/U-box | ring type do | no mercator | NA | PF13639 | IPR001841 |
| 827 | HORVU.MOREX.r3.5HG04715 | high | protein_cod | *** | Ubiquitin-co | E2 MUB ubi | protein hom | GO:000016 | PF00179 | IPR000608,IPR016135 |
| 828 | HORVU.MOREX.r3.5HG04720 | high | protein_cod | *** | Transcriptio | bZIP class-S | rna biosynth | GO:000370 | PF00170 | IPR004827 |
| 829 | HORVU.MOREX.r3.5HG04728 | high | protein_cod | *** | Aquaporin | plasma mer | solute trans | GO:001526 | PF00230 | IPR000425,IPR023271,IPR000425 |
| 830 | HORVU.MOREX.r3.5HG04736 | high | protein_cod | *** | Protein pho | clade A pho | protein moc | GO:000382 | PF00481 | IPR001932,IPR001932 |
| 831 | HORVU.MOREX.r3.5HG04740 | high | protein_cod | *** | Pleiotropic | subfamily A | solute trans | GO:000016 | PF08370,PF | IPR027417,IPR013581,IPR013525,IPR027417,IPR003439,IPR003439,IPR02948 |
| 832 | HORVU.MOREX.r3.5HG04758 | high | protein_cod | --* | SKP1-like pr | ppiase cycl | no mercator | GO:000651 | NA | NA |
| 833 | HORVU.MOREX.r3.5HG04759 | high | protein_cod | --* | cotton fiber | not classifie | no mercator | NA | PF05553 | IPR008480 |
| 834 | HORVU.MOREX.r3.5HG04760 | high | protein_cod | *** | Molybdopte | catalytic co | coenzyme n | GO:000573 | PF02391 | IPR003448,IPR003448 |
| 835 | HORVU.MOREX.r3.5HG04761 | high | protein_cod | *** | Chaperone | chaperone | no mercator | NA | PF00226 | IPR001623,IPR001623 |
| 836 | HORVU.MOREX.r3.5HG04761 | high | protein_cod | *** | Chaperone | sp dnaj cha | no mercator | NA | PF00226 | IPR001623,IPR001623 |
| 837 | HORVU.MOREX.r3.5HG04762 | high | protein_cod | *** | Glutathione | class tau glu | redox home | GO:001674 | PF13417 | IPR004045,IPR010987,IPR012336 |
| 838 | HORVU.MOREX.r3.5HG04777 | high | protein_cod | *** | Homeobox | HD-ZIP I/II-t | rna biosynth | GO:000367 | PF00046,PF | IPR001356,IPR003106,IPR009057 |
| 839 | HORVU.MOREX.r3.5HG04778 | high | protein_cod | *** | Endo-1,3(4) | endo 4 beta | no mercator | GO:005286 | PF03639 | IPR005200 |
| 840 | HORVU.MOREX.r3.5HG04783 | high | protein_cod | *** | Zinc finger A | an type dom | no mercator | GO:000827 | PF01428,PF | IPR000058,IPR000058 |
| 841 | HORVU.MOREX.r3.5HG04788 | high | protein_cod | *** | Trehalose-6 | EC_2.4 glyc | enzyme clas | GO:000382 | PF02358,PF | IPR006379,IPR023214,IPR003337,IPR003337,IPR00183 |
| 842 | HORVU.MOREX.r3.5HG04790 | high | protein_cod | *** | Alcohol deh | EC_1.1 oxid | enzyme clas | GO:000827 | PF00107,PF | IPR011032,IPR016040,IPR013149,IPR013154 |
| 843 | HORVU.MOREX.r3.5HG04790 | high | protein_cod | --* | Galactose o | not classifie | no mercator | NA | NA | NA |
| 844 | HORVU.MOREX.r3.5HG04798 | high | protein_cod | --* | Protein BIG | protein big | no mercator | NA | NA | NA |

| | | | | | | | | | | |
|---|---|---|---|---|---|---|---|---|---|---|
| 845 | HORVU.MOREX.r3.5HG048004 | high | protein_cod | *** | MTD1 | regulatory p | multi-proce | NA | NA | NA |
| 846 | HORVU.MOREX.r3.5HG048006 | high | protein_cod | *** | Protein GAM | karyogamy f | plant reproc | GO:001602 | NA | NA |
| 847 | HORVU.MOREX.r3.5HG048019 | low | protein_cod | --* | Peroxisome | not classifie | no mercator | GO:000030 | NA | NA |
| 848 | HORVU.MOREX.r3.5HG048041 | high | protein_cod | *** | Copper tran | copper catio | solute trans | GO:000537 | PF04145 | IPR007274 |
| 849 | HORVU.MOREX.r3.5HG048085 | high | protein_cod | *** | SAUR-like a | saur auxin r | no mercator | GO:000973 | PF02519 | IPR003676 |
| 850 | HORVU.MOREX.r3.5HG048089 | high | protein_cod | *** | Dirigent pro | dirigent prot | no mercator | GO:000557 | PF03018 | IPR004265 |
| 851 | HORVU.MOREX.r3.5HG048098 | low | protein_cod | --* | Spermidine | auxin respo | no mercator | GO:000016 | NA | NA |
| 852 | HORVU.MOREX.r3.5HG048129 | high | protein_cod | *** | RING/U-box | signal trans | phytohormo | NA | PF13639 | IPR001841 |
| 853 | HORVU.MOREX.r3.5HG048158 | high | protein_cod | *** | Hexosyltran | hexosyltrans | no mercator | GO:000013 | PF01762 | IPR002659 |
| 854 | HORVU.MOREX.r3.5HG048161 | high | protein_cod | *** | Flavin-conta | flavin-deper | phytohormo | GO:000449 | IPR023753,IPR023753,IPR023753 | |
| 855 | HORVU.MOREX.r3.5HG048239 | high | protein_cod | *** | Tetraspanin | regulatory p | vesicle traff | GO:001602 | PF00335 | IPR018499 |
| 856 | HORVU.MOREX.r3.5HG048255 | high | protein_cod | *** | Chaperone | regulatory p | external stir | NA | IPR001305 | |
| 857 | HORVU.MOREX.r3.5HG048406 | high | protein_cod | *** | WRKY trans | transcriptio | rna biosynth | GO:000367 | PF03106 | IPR003657,IPR003657 |
| 858 | HORVU.MOREX.r3.5HG048415 | high | protein_cod | *** | Polynucleot | deadenylas | rna process | NA | PF04857,PF | IPR012337,IPR006941,IPR006941 |
| 859 | HORVU.MOREX.r3.5HG048463 | high | protein_cod | --* | Homeodom | PSY precurs | phytohormo | NA | NA | NA |
| 860 | HORVU.MOREX.r3.5HG048469 | high | protein_cod | *-* | Calcium-bir | calcium sen | multi-proce | GO:000550 | PF13499 | IPR002048,IPR011992 |
| 861 | HORVU.MOREX.r3.5HG048561 | low | non_coding | *** | Cytochrome | EC_1.14 oxi | enzyme clas | GO:000449 | PF00067 | IPR001128,IPR001128 |
| 862 | HORVU.MOREX.r3.5HG048564 | low | non_coding | *** | Cytochrome | EC_1.14 oxi | enzyme clas | GO:000449 | PF00067 | IPR001128,IPR001128 |
| 863 | HORVU.MOREX.r3.5HG048592 | high | protein_cod | *** | Ripening-rel | kiwellin | no mercator | NA | IPR009009 | |
| 864 | HORVU.MOREX.r3.5HG048594 | high | protein_cod | *** | Hexosyltran | beta-1,3-ga | cell wall org | GO:000013 | PF01762,PF | IPR002659,IPR025298 |
| 865 | HORVU.MOREX.r3.5HG048609 | high | protein_cod | *** | Phosphatidy | regulatory p | plant organ | GO:001602 | PF02493,PF | IPR003409,IPR003409,IPR003409,IPR003409,IPR003409,IPR003409,IPR003409 |
| 866 | HORVU.MOREX.r3.5HG048620 | high | protein_cod | *** | Mitochondri | phosphate t | solute trans | GO:001602 | PF00153,PF | IPR018108,IPR018108,IPR023395 |
| 867 | HORVU.MOREX.r3.5HG048622 | high | protein_cod | *** | Serine/threo | EC_2.7 tran | enzyme clas | GO:000016 | PF01453,PF | IPR001480,IPR001480,IPR011009,IPR011009,IPR000719 |
| 868 | HORVU.MOREX.r3.5HG048653 | high | protein_cod | *** | Ethylene-re | subgroup EF | rna biosynth | GO:000367 | PF00847 | IPR001471,IPR016177 |
| 869 | HORVU.MOREX.r3.5HG048703 | high | protein_cod | *** | Alpha/beta- | abhydrolase | no mercator | GO:000815 | PF07859 | IPR013094,IPR029058 |
| 870 | HORVU.MOREX.r3.5HG048707 | high | protein_cod | *** | Alpha/beta- | abhydrolase | no mercator | GO:000815 | PF07859 | IPR029058,IPR013094 |
| 871 | HORVU.MOREX.r3.5HG048709 | high | protein_cod | *** | Alpha/beta- | abhydrolase | no mercator | GO:000815 | PF07859 | IPR029058,IPR013094 |
| 872 | HORVU.MOREX.r3.5HG048796 | low | protein_cod | --* | At4g40080 | enth domair | no mercator | GO:000554 | NA | NA |
| 873 | HORVU.MOREX.r3.5HG048798 | high | protein_cod | *** | Ring finger p | RING-H2-cl | protein hom | GO:001602 | PF13639 | IPR001841 |
| 874 | HORVU.MOREX.r3.5HG048804 | high | protein_cod | *** | Blue copper | phytocyanin | no mercator | GO:000905 | PF02298 | IPR003245,IPR008972 |
| 875 | HORVU.MOREX.r3.5HG048830 | high | protein_cod | *** | ABC transp | subfamily A | solute trans | GO:000016 | PF00005,PF | IPR003439,IPR027417,IPR013525 |
| 876 | HORVU.MOREX.r3.5HG048865 | low | protein_cod | *-* | Dof zinc fing | DOF transcr | rna biosynth | GO:000367 | PF02701 | IPR003851 |
| 877 | HORVU.MOREX.r3.5HG048897 | high | protein_cod | *** | Cellulose sy | cellulose sy | no mercator | GO:001602 | PF03552,PF | IPR005150,IPR005150,IPR029044,IPR029044 |
| 878 | HORVU.MOREX.r3.5HG048906 | high | protein_cod | *** | F-box protei | f box domai | no mercator | NA | PF00646 | IPR001810,IPR017451,IPR001810 |
| 879 | HORVU.MOREX.r3.5HG049038 | high | protein_cod | *** | Calcium-bir | SRC1-clade | multi-proce | GO:000550 | PF13499 | IPR011992,IPR002048 |
| 880 | HORVU.MOREX.r3.5HG049058 | high | protein_cod | *** | Zinc finger A | zinc finger a | no mercator | GO:000367 | PF01754,PF | IPR002653,IPR000058 |
| 881 | HORVU.MOREX.r3.5HG049079 | high | protein_cod | *** | Cobyric acid | histone h3 2 | no mercator | NA | NA | NA |
| 882 | HORVU.MOREX.r3.5HG049147 | high | protein_cod | *** | DUF4228 do | hth type tra | no mercator | NA | PF14009 | IPR025322 |
| 883 | HORVU.MOREX.r3.5HG049251 | high | protein_cod | *** | N-methyl-L- | pipecolate c | external stir | GO:001649 | PF01266 | IPR006076,IPR023753,IPR023753 |
| 884 | HORVU.MOREX.r3.5HG049272 | high | protein_cod | *** | MTD1 | regulatory p | multi-proce | NA | NA | NA |
| 885 | HORVU.MOREX.r3.5HG049297 | high | protein_cod | *** | Auxin-induc | b561 and do | no mercator | GO:000367 | PF04526 | IPR005018 |
| 886 | HORVU.MOREX.r3.5HG049299 | high | protein_cod | *** | Auxin-induc | b561 and do | no mercator | GO:000367 | PF04526 | IPR005018 |
| 887 | HORVU.MOREX.r3.5HG049336 | high | protein_cod | *** | Chymotryps | PR6 proteas | protein hom | GO:000486 | PF00280 | IPR000864,IPR000864 |
| 888 | HORVU.MOREX.r3.5HG049407 | high | protein_cod | *** | Purple acid | purple acid | no mercator | GO:000399 | PF00149,PF | IPR004843,IPR029052,IPR015914,IPR025733,IPR008963 |
| 889 | HORVU.MOREX.r3.5HG049431 | high | protein_cod | *** | Transmemb | not classifie | no mercator | GO:001602 | NA | NA |
| 890 | HORVU.MOREX.r3.5HG049452 | high | protein_cod | *** | SAUR-like a | indole aceti | no mercator | GO:000973 | PF02519 | IPR003676 |
| 891 | HORVU.MOREX.r3.5HG049479 | high | protein_cod | *** | NAC domain | NAC transcr | rna biosynth | GO:000367 | PF02365 | IPR003441,IPR003441 |
| 892 | HORVU.MOREX.r3.5HG049536 | high | protein_cod | *** | Beta-glucos | EC_3.2 glyc | enzyme clas | GO:000455 | PF00232 | IPR001360,IPR017853 |
| 893 | HORVU.MOREX.r3.5HG049564 | high | protein_cod | *** | Solute carri | solute trans | solute trans | GO:001602 | PF06027 | IPR009262 |
| 894 | HORVU.MOREX.r3.5HG049586 | high | protein_cod | *** | glycosyltran | xylosyltrans | cell wall org | NA | PF03016 | IPR004263 |
| 895 | HORVU.MOREX.r3.5HG049591 | high | protein_cod | *** | Respiratory | NADPH-oxid | redox home | GO:000460 | PF01794,PF | IPR017938,IPR017938,IPR013130,IPR013121,IPR013623,IPR011992,IPR013112 |
| 896 | HORVU.MOREX.r3.5HG049662 | high | protein_cod | *** | Glutathione | maleylaceto | amino acid | GO:000382 | PF13417 | IPR005955,IPR012336,IPR004045,IPR010987 |
| 897 | HORVU.MOREX.r3.5HG049688 | high | protein_cod | *** | Glycosyltrar | EC_2.4 glyc | enzyme clas | GO:000815 | PF00201 | IPR002213 |
| 898 | HORVU.MOREX.r3.5HG049796 | high | protein_cod | *** | BON1-asso | bon associa | no mercator | NA | NA | NA |
| 899 | HORVU.MOREX.r3.5HG049797 | high | protein_cod | *** | Ethylene-re | subgroup EF | rna biosynth | GO:000367 | PF00847 | IPR001471,IPR016177 |
| 900 | HORVU.MOREX.r3.5HG049799 | high | protein_cod | *** | Heat shock | HSF transcr | rna biosynth | GO:000367 | PF00447 | IPR000232,IPR011991 |
| 901 | HORVU.MOREX.r3.5HG049823 | high | protein_cod | *** | GTPase-acti | Rab GTPase | vesicle traff | NA | PF00566 | IPR000195,IPR000195,IPR000195,IPR000195,IPR000195 |
| 902 | HORVU.MOREX.r3.5HG049848 | high | protein_cod | *** | Glycosyltrar | EC_2.4 glyc | enzyme clas | GO:000815 | PF00201 | IPR002213 |
| 903 | HORVU.MOREX.r3.5HG049882 | high | protein_cod | *** | Late embryo | lea 2 domai | no mercator | NA | PF03168 | IPR004864 |
| 904 | HORVU.MOREX.r3.5HG049903 | low | non_coding | *** | ATP-depend | chaperone c | protein hom | GO:000016 | PF00004,PF | IPR003959,IPR019489,IPR027417,IPR004176,IPR004176,IPR027417,IPR004176,IPR003959 |
| 905 | HORVU.MOREX.r3.5HG049934 | high | protein_cod | *** | Kinase fami | RLCK-VIIa r | protein mod | GO:000016 | PF07714 | IPR001245,IPR011009 |
| 906 | HORVU.MOREX.r3.5HG049976 | high | protein_cod | *** | Cysteine pro | Papain-type | protein hom | GO:000650 | PF00112,PF | IPR000668,IPR013201 |
| 907 | HORVU.MOREX.r3.5HG050142 | high | protein_cod | *** | Ornithine de | ornithine de | polyamine r | GO:000382 | PF00278,PF | IPR029066,IPR009006,IPR022643,IPR022644 |
| 908 | HORVU.MOREX.r3.5HG050147 | high | protein_cod | *** | Receptor-lik | DUF26 prot | protein mod | GO:000467 | PF00069,PF | IPR000719,IPR011009,IPR002902,IPR002902 |
| 909 | HORVU.MOREX.r3.5HG050150 | high | protein_cod | *** | Glucan end | EC_3.2 glyc | enzyme clas | GO:000455 | PF07983,PF | IPR012946,IPR000490,IPR017853 |

| | | | | | | | | | | |
|---|---|---|---|---|---|---|---|---|---|---|
| 910 | HORVU.MOREX.r3.5HG050167 | high | protein_cod | *** | Ankyrin repe | pgg domain | no mercato | GO:001602 | PF12796,PF | IPR020683,IPR020683,IPR020683,IPR020683,IPR020683,IPR026961 |
| 911 | HORVU.MOREX.r3.5HG050301 | high | protein_cod | *** | Flavin-conta | containing r | no mercato | GO:000449 | PF00743 | IPR020946,IPR023753,IPR023753 |
| 912 | HORVU.MOREX.r3.5HG050442 | low | non_coding | *** | Cytochrome | EC_1.14 oxi | enzyme clas | GO:000449 | PF00067 | IPR001128,IPR001128 |
| 913 | HORVU.MOREX.r3.5HG050453 | high | protein_cod | *** | Protein ELC | component | vesicle traff | GO:000646 | PF05743,PF | IPR008883,IPR016135,IPR017916 |
| 914 | HORVU.MOREX.r3.5HG050494 | high | protein_cod | *** | Purine perm | organic cati | solute trans | GO:000521 | NA | NA |
| 915 | HORVU.MOREX.r3.5HG050602 | high | protein_cod | *-* | Receptor-lik | WAK/WAKL | protein mod | GO:000016 | PF07714 | IPR001245,IPR011009 |
| 916 | HORVU.MOREX.r3.5HG050623 | high | protein_cod | *** | Cysteine pr | EC_3.4 hydr | enzyme clas | GO:000650 | PF08246,PF | IPR013201,IPR000668 |
| 917 | HORVU.MOREX.r3.5HG050629 | high | protein_cod | *-* | Splicing fac | large subun | rna process | GO:000367 | PF00076 | IPR000504,IPR012677,IPR012677,IPR006529,IPR012677 |
| 918 | HORVU.MOREX.r3.5HG050669 | low | protein_cod | --* | glycine/prol | NA | NA | NA | NA | NA |
| 919 | HORVU.MOREX.r3.5HG050679 | high | protein_cod | *** | Calcineurin | metallopho | no mercato | GO:001602 | PF00149 | IPR029052,IPR029052,IPR004843 |
| 920 | HORVU.MOREX.r3.5HG050797 | low | protein_cod | --* | Eukaryotic t | transcriptio | rna biosynth | GO:000173 | NA | NA |
| 921 | HORVU.MOREX.r3.5HG050827 | high | protein_cod | *** | Aminotrans | ornithine an | amino acid | GO:000382 | PF00202 | IPR010164,IPR015424,IPR005814 |
| 922 | HORVU.MOREX.r3.5HG050873 | high | protein_cod | *** | F-box family | f box domai | no mercato | NA | PF12937 | IPR001810,IPR001810 |
| 923 | HORVU.MOREX.r3.5HG050912 | high | protein_cod | *** | Auxin-respo | transcriptio | phytohormo | GO:000563 | PF02309,PF | IPR033389,IPR033389 |
| 924 | HORVU.MOREX.r3.5HG050921 | high | protein_cod | *** | Lipoxygenas | 9-lipoxygen | redox home | GO:000663 | PF01477,PF | IPR001024,IPR013819,IPR001024,IPR013819 |
| 925 | HORVU.MOREX.r3.5HG050939 | high | protein_cod | *** | LEM3 (Ligan | regulatory c | lipid metab | GO:001602 | PF03381 | IPR005045 |
| 926 | HORVU.MOREX.r3.5HG050982 | high | protein_cod | *** | ATP sulfuryl | ATP sulfuryl | coenzyme n | GO:000010 | PF14306,PF | IPR025980,IPR024951,IPR015947,IPR002650 |
| 927 | HORVU.MOREX.r3.5HG051024 | high | protein_cod | *** | Dirigent pro | arath diriger | no mercato | GO:000557 | PF03018 | IPR004265 |
| 928 | HORVU.MOREX.r3.5HG051025 | high | protein_cod | *** | Dirigent pro | arath diriger | no mercato | GO:000557 | PF03018 | IPR004265 |
| 929 | HORVU.MOREX.r3.5HG051046 | high | protein_cod | *** | Maltase-glu | dna directe | no mercato | GO:001602 | NA | NA |
| 930 | HORVU.MOREX.r3.5HG051047 | high | protein_cod | *** | Cytosolic Fe | assembly fa | coenzyme n | NA | PF02256,PF | IPR009016,IPR003149,IPR004108 |
| 931 | HORVU.MOREX.r3.5HG051088 | low | protein_cod | --* | Heavy meta | not classifie | no mercato | NA | NA | NA |
| 932 | HORVU.MOREX.r3.5HG051231 | low | non_coding | *** | Cytochrome | EC_1.14 oxi | enzyme clas | GO:000449 | PF00067 | IPR001128,IPR001128 |
| 933 | HORVU.MOREX.r3.5HG051258 | high | protein_cod | *-* | myosin-binc | gtd binding | no mercato | NA | PF04576 | IPR007656 |
| 934 | HORVU.MOREX.r3.5HG051280 | high | protein_cod | *** | Two-compo | GARP subgr | rna biosynth | GO:000367 | PF00249 | IPR006447,IPR001005,IPR009057 |
| 935 | HORVU.MOREX.r3.5HG051289 | high | protein_cod | *-* | Nuclear por | nucleoporin | protein tran | NA | NA | NA |
| 936 | HORVU.MOREX.r3.5HG051292 | high | protein_cod | --* | ARGONAUT | genome ass | no mercato | NA | PF12274 | IPR022059 |
| 937 | HORVU.MOREX.r3.5HG051313 | high | protein_cod | *** | Thioredoxin | protein invo | photosynthe | GO:000562 | PF00085 | IPR013766,IPR012336 |
| 938 | HORVU.MOREX.r3.5HG051319 | high | protein_cod | *** | Cold acclim | cold-respon | external stir | GO:001602 | PF05562 | IPR008892 |
| 939 | HORVU.MOREX.r3.5HG051334 | high | protein_cod | --* | Basic-leucir | not classifie | no mercato | NA | NA | NA |
| 940 | HORVU.MOREX.r3.5HG051396 | high | protein_cod | *** | Erythronate | protein dom | no mercato | GO:001602 | NA | NA |
| 941 | HORVU.MOREX.r3.5HG051429 | high | protein_cod | --* | Chaperone | component | protein tran | GO:000552 | IPR001623 | |
| 942 | HORVU.MOREX.r3.5HG051468 | low | protein_cod | --* | Phenylalani | not classifie | no mercato | GO:000004 | NA | NA |
| 943 | HORVU.MOREX.r3.5HG051505 | high | protein_cod | *-* | Ribose-5-ph | ribose phos | no mercato | GO:000475 | NA | NA |
| 944 | HORVU.MOREX.r3.5HG051671 | high | protein_cod | *** | Glutathione | class tau gl | redox home | GO:001674 | PF13417 | IPR004045,IPR012336,IPR010987 |
| 945 | HORVU.MOREX.r3.5HG051683 | high | protein_cod | *** | Dehydrin | dehydrin dh | no mercato | GO:000695 | PF00257 | IPR000167 |
| 946 | HORVU.MOREX.r3.5HG051705 | high | protein_cod | *** | Expp1 prote | diphthamid | protein bios | NA | NA | NA |
| 947 | HORVU.MOREX.r3.5HG051725 | high | protein_cod | *** | Alpha/beta- | abhydrolase | no mercato | GO:000815 | PF07859 | IPR029058,IPR013094 |
| 948 | HORVU.MOREX.r3.5HG051806 | high | protein_cod | *** | Glycosyltra | EC_2.4 glyc | enzyme clas | GO:000815 | PF00201 | IPR002213 |
| 949 | HORVU.MOREX.r3.5HG051813 | high | protein_cod | *** | Acetyltransf | n acetyltran | no mercato | GO:001674 | PF13302 | IPR000182,IPR016181 |
| 950 | HORVU.MOREX.r3.5HG051856 | high | protein_cod | *-* | Leucine-rich | LRR-domain | cell wall org | NA | PF08263 | IPR032675,IPR013210 |
| 951 | HORVU.MOREX.r3.5HG051912 | high | protein_cod | *** | CASP-like p | casp protein | no mercato | GO:000588 | PF04535 | IPR006702,IPR006459 |
| 952 | HORVU.MOREX.r3.5HG052083 | high | protein_cod | *** | Actin cross- | duf 569 don | no mercato | NA | PF04601 | IPR008999,IPR007679 |
| 953 | HORVU.MOREX.r3.5HG052094 | high | protein_cod | *-* | ATP-depend | EC_3.6 hydr | enzyme clas | GO:000016 | PF14363,PF | IPR027417,IPR025753,IPR003959 |
| 954 | HORVU.MOREX.r3.5HG052121 | low | protein_cod | --* | HCO3- tran | not classifie | no mercato | NA | NA | NA |
| 955 | HORVU.MOREX.r3.5HG052251 | high | protein_cod | --* | Galactose o | not classifie | no mercato | NA | NA | NA |
| 956 | HORVU.MOREX.r3.5HG052305 | high | protein_cod | *** | Phosphate t | phosphate t | solute trans | GO:000531 | PF00083 | IPR004738,IPR020846,IPR005828 |
| 957 | HORVU.MOREX.r3.5HG052401 | high | protein_cod | *** | Glycosyltra | EC_2.4 glyc | enzyme clas | GO:000815 | PF00201 | IPR002213 |
| 958 | HORVU.MOREX.r3.5HG052406 | high | protein_cod | *** | Remorin | remorin c d | no mercato | NA | PF03763 | IPR005516 |
| 959 | HORVU.MOREX.r3.5HG052413 | high | protein_cod | *** | Triacylglyce | Patatin-type | lipid metab | GO:000480 | PF11815,PF | IPR021771,IPR016035,IPR002641 |
| 960 | HORVU.MOREX.r3.5HG052414 | high | protein_cod | --* | 7-cyano-7-c | not classifie | no mercato | GO:000016 | NA | NA |
| 961 | HORVU.MOREX.r3.5HG052463 | high | protein_cod | *-* | Ethylene-re | transcriptio | phytohormo | GO:000367 | PF00847 | IPR016177,IPR001471 |
| 962 | HORVU.MOREX.r3.5HG052480 | low | non_coding | --* | RNA polyme | fe2og dioxy | no mercato | NA | NA | NA |
| 963 | HORVU.MOREX.r3.5HG052482 | high | protein_cod | *** | DCD (Devel | dcd domain | no mercato | NA | PF10539 | IPR013989 |
| 964 | HORVU.MOREX.r3.5HG052518 | high | protein_cod | *** | LURP-one-li | protein lurp | no mercato | NA | PF04525 | IPR007612,IPR025659 |
| 965 | HORVU.MOREX.r3.5HG052587 | high | protein_cod | *-* | Armadillo/b | cbm domain | no mercato | GO:000563 | IPR016024 | |
| 966 | HORVU.MOREX.r3.5HG052594 | high | protein_cod | *** | Zinc finger p | ZAT transcri | rna biosynth | GO:000367 | PF13912,PF | IPR007087,IPR007087 |
| 967 | HORVU.MOREX.r3.5HG052599 | high | protein_cod | *** | Zinc finger p | ZAT transcri | rna biosynth | GO:000367 | PF13912,PF | IPR007087,IPR007087 |
| 968 | HORVU.MOREX.r3.5HG052601 | high | protein_cod | *** | Zinc finger p | ZAT transcri | rna biosynth | GO:000367 | PF13912,PF | IPR007087,IPR007087 |
| 969 | HORVU.MOREX.r3.5HG052603 | high | protein_cod | *** | Zinc finger p | ZAT transcri | rna biosynth | GO:000367 | PF13912,PF | IPR007087,IPR007087 |
| 970 | HORVU.MOREX.r3.5HG052604 | high | protein_cod | *** | Zinc finger p | ZAT transcri | rna biosynth | GO:000367 | PF13912,PF | IPR007087,IPR007087 |
| 971 | HORVU.MOREX.r3.5HG052605 | high | protein_cod | *** | Zinc finger p | ZAT transcri | rna biosynth | GO:000367 | PF13912,PF | IPR007087,IPR007087 |
| 972 | HORVU.MOREX.r3.5HG052606 | high | protein_cod | *** | Zinc finger p | ZAT transcri | rna biosynth | GO:000367 | PF13912,PF | IPR007087,IPR007087 |
| 973 | HORVU.MOREX.r3.5HG052609 | high | protein_cod | *** | Zinc finger p | ZAT transcri | rna biosynth | GO:000367 | PF13912,PF | IPR007087,IPR007087 |
| 974 | HORVU.MOREX.r3.5HG052611 | high | protein_cod | *** | Zinc finger p | ZAT transcri | rna biosynth | GO:000367 | PF13912,PF | IPR007087,IPR007087 |

| | | | | | | | | | | |
|---|---|---|---|---|---|---|---|---|---|---|
| 975 | HORVU.MOREX.r3.5HG05261 | high | protein_cod | *** | Actin depol | actin-depol | cytoskeleto | GO:00037 | PF00241 | IPR002108 |
| 976 | HORVU.MOREX.r3.5HG05262 | high | protein_cod | *** | Phosphatas | clade C pho | protein mod | GO:000382 | PF00481 | IPR001932,IPR001932,IPR001932 |
| 977 | HORVU.MOREX.r3.5HG05265 | high | protein_cod | *** | NAD(P)-bin | nad p bd do | no mercato | NA | PF13460 | IPR016040,IPR016040 |
| 978 | HORVU.MOREX.r3.5HG05272 | high | protein_cod | --* | Trypsin inhi | Bowman-Bi | protein hom | GO:000486 | PF00228 | IPR000877,IPR000877 |
| 979 | HORVU.MOREX.r3.5HG05273 | high | protein_cod | *-* | Protein MID | mcafunc do | no mercato | GO:000716 | NA | NA |
| 980 | HORVU.MOREX.r3.5HG05274 | high | protein_cod | *** | rRNA N-glyc | rrna n | no mercato | NA | NA | NA |
| 981 | HORVU.MOREX.r3.5HG05274 | high | protein_cod | *** | Pectin acet | pectin acet | cell wall org | GO:000557 | PF03283 | IPR029058,IPR029058,IPR029058,IPR00496 |
| 982 | HORVU.MOREX.r3.5HG05277 | high | protein_cod | *** | Receptor pr | RLCK-IV rec | protein mod | GO:000016 | PF07714 | IPR001245,IPR011009 |
| 983 | HORVU.MOREX.r3.5HG05278 | high | protein_cod | *** | GRAM doma | gram domai | no mercato | NA | PF02893 | IPR004182 |
| 984 | HORVU.MOREX.r3.5HG05283 | high | protein_cod | *** | High affinity | regulatory f | nutrient upt | GO:001016 | PF16974 | IPR016605 |
| 985 | HORVU.MOREX.r3.5HG05285 | high | protein_cod | *** | Epoxide hyd | epoxide hyd | cell wall org | GO:000382 | PF12697 | IPR029058,IPR000073 |
| 986 | HORVU.MOREX.r3.5HG05286 | high | protein_cod | *** | Protein PLA | protein plan | no mercato | GO:001602 | PF04749 | IPR006461,IPR006461 |
| 987 | HORVU.MOREX.r3.5HG05291 | high | protein_cod | *** | F-box doma | f box domai | no mercato | GO:000015 | PF12937 | IPR001810,IPR001810 |
| 988 | HORVU.MOREX.r3.5HG05296 | high | protein_cod | *** | Phosphate t | phosphate t | solute trans | GO:000531 | PF00083 | IPR020846,IPR005828,IPR004738 |
| 989 | HORVU.MOREX.r3.5HG05297 | high | protein_cod | *** | Deaminase- | guanosine d | nucleotide r | GO:000382 | PF00383 | IPR002125,IPR016193 |
| 990 | HORVU.MOREX.r3.5HG05302 | high | protein_cod | *** | peptidyl-pr | peptidyl pro | no mercato | NA | NA | NA |
| 991 | HORVU.MOREX.r3.5HG05310 | high | protein_cod | *** | Glutathione | class phi gl | redox home | GO:001674 | PF02798,PF | IPR004045,IPR012336,IPR010987,IPR00404 |
| 992 | HORVU.MOREX.r3.5HG05321 | high | protein_cod | *** | Expansin | NA | NA | GO:000557 | PF01357,PF | IPR007117,IPR009009,IPR009009,IPR00711 |
| 993 | HORVU.MOREX.r3.5HG05321 | high | protein_cod | *** | Expansin | NA | NA | GO:000557 | PF01357,PF | IPR007117,IPR009009,IPR007117,IPR00900 |
| 994 | HORVU.MOREX.r3.5HG05321 | high | protein_cod | *** | Expansin | NA | NA | GO:000557 | PF01357,PF | IPR007117,IPR009009,IPR007117,IPR00900 |
| 995 | HORVU.MOREX.r3.5HG05322 | high | protein_cod | *** | Patatin | phospholip | lipid metab | GO:000662 | PF01734 | IPR002641,IPR016035 |
| 996 | HORVU.MOREX.r3.5HG05332 | high | protein_cod | *** | Pirin-like pr | transcriptio | rna biosynt | GO:001602 | PF02678,PF | IPR003829,IPR011051,IPR008778 |
| 997 | HORVU.MOREX.r3.5HG05362 | high | protein_cod | *** | Xyloglucan | EC_2.4 glyc | enzyme cla | GO:000455 | PF00722,PF | IPR000757,IPR013320,IPR010713 |
| 998 | HORVU.MOREX.r3.5HG05369 | high | protein_cod | *** | Cytokinin ri | cytokinin ph | phytohorm | GO:000969 | PF03641 | IPR005269,IPR031100 |
| 999 | HORVU.MOREX.r3.5HG05381 | high | protein_cod | *** | Glutathione | class theta | redox home | GO:001674 | PF02798 | IPR010987,IPR012336,IPR004045 |
| 1000 | HORVU.MOREX.r3.5HG05381 | high | protein_cod | *** | Pleiotropic | subfamily A | solute trans | GO:000016 | PF08370,PF | IPR013581,IPR029481,IPR027417,IPR003439,IPR003439,IPR027417,IPR013525,IPR01352 |
| 1001 | HORVU.MOREX.r3.6HG05400 | high | protein_cod | *** | O-methyltra | caffeic acid | cell wall org | GO:000816 | PF00891,PF | IPR011991,IPR029063,IPR001077,IPR012967 |
| 1002 | HORVU.MOREX.r3.6HG05401 | high | protein_cod | *** | Rhodanese- | rhodanese | no mercato | GO:001674 | PF00581 | IPR001763,IPR001763 |
| 1003 | HORVU.MOREX.r3.6HG05414 | high | protein_cod | *** | Glycosyltra | EC_2.4 glyc | enzyme cla | GO:000815 | PF00201 | IPR002213 |
| 1004 | HORVU.MOREX.r3.6HG05431 | high | protein_cod | *** | Germin-like | germin prot | no mercato | GO:000557 | PF00190 | IPR006045,IPR011051 |
| 1005 | HORVU.MOREX.r3.6HG05433 | high | protein_cod | *** | High affinity | nitrate trans | solute trans | GO:001602 | PF07690 | IPR020846,IPR011701 |
| 1006 | HORVU.MOREX.r3.6HG05435 | high | protein_cod | *** | Expansin pr | expansin b1 | no mercato | GO:000557 | PF03330,PF | IPR007117,IPR009009,IPR007117,IPR00900 |
| 1007 | HORVU.MOREX.r3.6HG05435 | high | protein_cod | *** | Expansin pr | NA | NA | GO:000557 | PF03330,PF | IPR009009,IPR007117,IPR007117,IPR00900 |
| 1008 | HORVU.MOREX.r3.6HG05435 | high | protein_cod | *** | High affinity | nitrate trans | solute trans | GO:001602 | PF07690 | IPR020846,IPR011701 |
| 1009 | HORVU.MOREX.r3.6HG05443 | high | protein_cod | *-* | MYB transc | MYB class-F | rna biosynt | GO:000367 | PF00249,PF | IPR001005,IPR001005,IPR009057 |
| 1010 | HORVU.MOREX.r3.6HG05479 | high | protein_cod | --* | ATP synthas | transmemb | no mercato | GO:000016 | NA | NA |
| 1011 | HORVU.MOREX.r3.6HG05480 | high | protein_cod | *** | Acetyl-coen | EC_6.2 ligas | enzyme cla | GO:000016 | PF00501,PF | IPR000873,IPR025110 |
| 1012 | HORVU.MOREX.r3.6HG05480 | high | protein_cod | *** | Glutathione | gst c termin | no mercato | GO:000436 | PF13409 | IPR004045,IPR010987,IPR012336,IPR01233 |
| 1013 | HORVU.MOREX.r3.6HG05488 | high | protein_cod | --* | transcriptio | not classifie | no mercato | NA | NA | NA |
| 1014 | HORVU.MOREX.r3.6HG05488 | high | protein_cod | *** | Calcium-bir | ef hand dom | no mercato | GO:000550 | PF13833 | IPR011992,IPR002048 |
| 1015 | HORVU.MOREX.r3.6HG05516 | high | protein_cod | *** | AT hook mo | AHL clade-E | rna biosynt | GO:000367 | PF03479 | IPR005175 |
| 1016 | HORVU.MOREX.r3.6HG05516 | high | protein_cod | *** | BAX inhibito | Prgrammed | multi-proce | GO:001602 | NA | NA |
| 1017 | HORVU.MOREX.r3.6HG05522 | high | protein_cod | *** | Polyubiquiti | UBQ ubiquit | protein hom | NA | PF00240,PF | IPR029071,IPR029071,IPR029071,IPR029071,IPR000626,IPR000626,IPR000626,IPR000626,IPR000626,IPR02907 |
| 1018 | HORVU.MOREX.r3.6HG05525 | high | protein_cod | *** | Response t | regulatory p | nutrient upt | NA | NA | NA |
| 1019 | HORVU.MOREX.r3.6HG05542 | high | protein_cod | *** | F-box family | not classifie | no mercato | NA | NA | NA |
| 1020 | HORVU.MOREX.r3.6HG05547 | low | non_coding | *** | Cytochrom | EC_1.14 oxi | enzyme cla | GO:000449 | PF00067 | IPR001128,IPR001128 |
| 1021 | HORVU.MOREX.r3.6HG05548 | high | protein_cod | *** | Phosphopa | phosphopa | coenzyme n | GO:000382 | PF01467 | IPR004821,IPR004821 |
| 1022 | HORVU.MOREX.r3.6HG05558 | high | protein_cod | *** | Plant regula | NLP transcr | rna biosynt | GO:000367 | PF00564,PF | IPR000270,IPR003035 |
| 1023 | HORVU.MOREX.r3.6HG05570 | high | protein_cod | *** | Calmodulin | arath iq don | no mercato | NA | NA | NA |
| 1024 | HORVU.MOREX.r3.6HG05576 | high | protein_cod | *** | Polyubiquiti | UBQ ubiquit | protein hom | NA | PF00240,PF | IPR029071,IPR029071,IPR029071,IPR029071,IPR029071,IPR000626,IPR000626,IPR000626,IPR000626,IPR00062 |
| 1025 | HORVU.MOREX.r3.6HG05583 | high | protein_cod | *** | Purple acid | EC_3.1 hydr | enzyme cla | GO:000399 | PF16656,PF | IPR029052,IPR015914,IPR025733,IPR008963,IPR00484 |
| 1026 | HORVU.MOREX.r3.6HG05583 | high | protein_cod | *-* | Calcium-de | c2 domain | no mercato | GO:000367 | PF00168 | IPR000008,IPR000008 |
| 1027 | HORVU.MOREX.r3.6HG05593 | high | protein_cod | *** | Polyubiquiti | UBQ ubiquit | protein hom | GO:000975 | PF00240,PF | IPR029071,IPR029071,IPR000626,IPR000626,IPR000626,IPR000626,IPR029071,IPR02907 |
| 1028 | HORVU.MOREX.r3.6HG05599 | high | protein_cod | *** | Apoptosis-i | pyr redox 2 | no mercato | GO:001649 | PF07992 | IPR023753,IPR023753 |
| 1029 | HORVU.MOREX.r3.6HG05600 | low | protein_cod | --* | phospholip | not classifie | no mercato | NA | NA | NA |
| 1030 | HORVU.MOREX.r3.6HG05600 | high | protein_cod | *** | DUF1644 fa | c2h2 type d | no mercato | NA | PF07800 | IPR012866 |
| 1031 | HORVU.MOREX.r3.6HG05601 | high | protein_cod | *** | Endoglucan | endoglucan | no mercato | GO:000027 | PF00759 | IPR001701,IPR008928,IPR008928 |
| 1032 | HORVU.MOREX.r3.6HG05611 | high | protein_cod | *** | Leucine-ric | lrrnt 2 doma | no mercato | NA | PF08263,PF | IPR013210,IPR001611,IPR032675,IPR001611,IPR03267 |
| 1033 | HORVU.MOREX.r3.6HG05658 | high | protein_cod | *** | Polyubiquiti | UBQ ubiquit | protein hom | NA | PF00240,PF | IPR029071,IPR029071,IPR000626,IPR000626,IPR000626,IPR000626,IPR000626,IPR029071,IPR029071,IPR02907 |
| 1034 | HORVU.MOREX.r3.6HG05658 | high | protein_cod | *** | Polyubiquiti | UBQ ubiquit | protein hom | NA | PF00240,PF | IPR029071,IPR000626,IPR000626,IPR000626,IPR029071,IPR029071 |
| 1035 | HORVU.MOREX.r3.6HG05658 | high | protein_cod | *** | Polyubiquiti | UBQ ubiquit | protein hom | NA | PF00240,PF | IPR029071,IPR029071,IPR000626,IPR000626,IPR000626,IPR000626,IPR000626,IPR000626,IPR029071,IPR029071,IPR029071,IPR02907 |
| 1036 | HORVU.MOREX.r3.6HG05673 | high | protein_cod | *** | Testis-expre | smp ltd dom | no mercato | GO:000828 | NA | NA |
| 1037 | HORVU.MOREX.r3.6HG05674 | high | protein_cod | *** | Flotillin-like | adapter pro | vesicle traff | NA | PF01145 | IPR001107,IPR001107 |
| 1038 | HORVU.MOREX.r3.6HG05693 | high | protein_cod | *** | Plant/T31B | dnmt rfd do | no mercato | NA | PF11443 | IPR002035,IPR024553 |
| 1039 | HORVU.MOREX.r3.6HG05706 | high | protein_cod | *** | BZIP transc | bZIP class-C | rna biosynt | GO:000370 | PF00170 | IPR004827 |

| | | | | | | | | | | |
|---|---|---|---|---|---|---|---|---|---|---|
| 1040 | HORVU.MOREX.r3.6HG05710 | high | protein_cod | *** | VQ motif far | regulatory p | rna biosynth | NA | PF05678 | IPR008889 |
| 1041 | HORVU.MOREX.r3.6HG05731 | high | protein_cod | *** | Copper-trar | P1B-type he | solute trans | GO:000016 | PF00122,PF | IPR001757,IPR008250,IPR006122,IPR006122,IPR006121,IPR023214,IPR023214,IPR006121,IPR027256,IPR006121,IPR006121,IPR00612 |
| 1042 | HORVU.MOREX.r3.6HG05741 | high | protein_cod | *** | U-box doma | E3 ubiquitin | protein hom | GO:000484 | PF04564 | IPR016024,IPR003613 |
| 1043 | HORVU.MOREX.r3.6HG05742 | high | protein_cod | --* | 60 kDa chap | tmv respons | no mercator | GO:000016 | PF14009 | IPR025322 |
| 1044 | HORVU.MOREX.r3.6HG05743 | high | protein_cod | *** | Receptor-lik | LysM protei | protein mod | GO:000467 | PF07714,PF | IPR001245,IPR018392,IPR011009,IPR018392 |
| 1045 | HORVU.MOREX.r3.6HG05744 | high | protein_cod | *** | Receptor-lik | LysM protei | protein mod | GO:000467 | PF01476,PF | IPR011009,IPR018392,IPR018392,IPR001245,IPR018392 |
| 1046 | HORVU.MOREX.r3.6HG05748 | high | protein_cod | *** | Amino acid | solute trans | solute trans | GO:001602 | PF01490 | IPR013057 |
| 1047 | HORVU.MOREX.r3.6HG05750 | high | protein_cod | *** | RPM1-intera | avrrpt cleav | no mercator | NA | PF05627,PF | IPR008700,IPR008700 |
| 1048 | HORVU.MOREX.r3.6HG05754 | high | protein_cod | *** | Glycosyltrar | hydroxycinn | cell wall org | GO:000815 | PF00201 | IPR002213 |
| 1049 | HORVU.MOREX.r3.6HG05801 | high | protein_cod | *** | Protein pho | clade E pho | protein mod | GO:000382 | PF00481 | IPR001932,IPR001932,IPR001932 |
| 1050 | HORVU.MOREX.r3.6HG05842 | high | protein_cod | *** | MACPF dom | macpf doma | no mercator | NA | PF01823 | IPR020864 |
| 1051 | HORVU.MOREX.r3.6HG05911 | high | protein_cod | *** | High affinity | regulatory fa | nutrient upt | GO:001016 | PF16974 | IPR016605 |
| 1052 | HORVU.MOREX.r3.6HG05911 | high | protein_cod | *** | High affinity | regulatory fa | nutrient upt | GO:001016 | PF16974 | IPR016605 |
| 1053 | HORVU.MOREX.r3.6HG05928 | high | protein_cod | *** | B-box zinc fi | BBX class-I | rna biosynth | GO:000562 | PF00643 | IPR000315 |
| 1054 | HORVU.MOREX.r3.6HG05929 | high | protein_cod | *** | Chitinase fa | acidic endo | no mercator | GO:000456 | PF00187,PF | IPR001002,IPR001002,IPR000726,IPR02334 |
| 1055 | HORVU.MOREX.r3.6HG05953 | high | protein_cod | *** | Zinc finger ( | anaphase p | no mercator | NA | PF17123,PF | IPR001841,IPR002035,IPR032838,IPR00203 |
| 1056 | HORVU.MOREX.r3.6HG05955 | high | protein_cod | *** | Ammonium | ammonium | solute trans | GO:000851 | PF00909 | IPR024041,IPR001905,IPR024041 |
| 1057 | HORVU.MOREX.r3.6HG05958 | low | protein_cod | --* | UDP-glucos | auxin respo | no mercator | NA | NA | NA |
| 1058 | HORVU.MOREX.r3.6HG05963 | high | protein_cod | --* | Heme-resp | not classifie | no mercator | GO:000098 | NA | NA |
| 1059 | HORVU.MOREX.r3.6HG05964 | high | protein_cod | *** | MYB | MYB class-F | rna biosynth | GO:000367 | PF00249,PF | IPR001005,IPR001005,IPR009057 |
| 1060 | HORVU.MOREX.r3.6HG05967 | high | protein_cod | *** | Aquaporin | plasma mer | solute trans | GO:001526 | PF00230 | IPR000425,IPR000425,IPR023271 |
| 1061 | HORVU.MOREX.r3.6HG05985 | high | protein_cod | *** | MYB transcr | MYB class-F | rna biosynth | GO:000367 | PF00249,PF | IPR009057,IPR001005,IPR001005 |
| 1062 | HORVU.MOREX.r3.6HG05991 | high | protein_cod | *** | Autophagy-r | accessory c | protein hom | GO:000691 | PF10033 | IPR018731 |
| 1063 | HORVU.MOREX.r3.6HG05994 | high | protein_cod | *** | Kinase fami | RLCK-VIIa r | protein mod | GO:000016 | PF07714 | IPR001245,IPR011009 |
| 1064 | HORVU.MOREX.r3.6HG06004 | high | protein_cod | *** | Alpha/beta- | duf 676 dom | no mercator | GO:001678 | PF05057 | IPR007751,IPR029058 |
| 1065 | HORVU.MOREX.r3.6HG06009 | high | protein_cod | *** | Aquaporin | tonoplast in | solute trans | GO:001526 | PF00230 | IPR000425,IPR023271,IPR000425 |
| 1066 | HORVU.MOREX.r3.6HG06011 | high | protein_cod | *** | Expansin pr | expansin b1 | no mercator | GO:000557 | PF03330,PF | IPR009009,IPR007117,IPR007117,IPR00900 |
| 1067 | HORVU.MOREX.r3.6HG06012 | high | protein_cod | *** | Sulfotransfe | EC_2.8 tran | enzyme clas | GO:000814 | PF00685 | IPR000863,IPR027417 |
| 1068 | HORVU.MOREX.r3.6HG06015 | high | protein_cod | *** | Cysteine sy | EC_2.5 tran | enzyme clas | GO:000412 | PF00291 | IPR005856,IPR001926,IPR005859,IPR00192 |
| 1069 | HORVU.MOREX.r3.6HG06019 | high | protein_cod | *** | Glutathione | glutathione | redox home | GO:000460 | PF00255 | IPR012336,IPR000889 |
| 1070 | HORVU.MOREX.r3.6HG06029 | high | protein_cod | *** | Kinase fami | RLCK-VIIa r | protein mod | GO:000016 | PF07714 | IPR001245,IPR011009 |
| 1071 | HORVU.MOREX.r3.6HG06030 | high | protein_cod | *** | Amino acid | amino acid | solute trans | GO:001602 | PF01490 | IPR013057 |
| 1072 | HORVU.MOREX.r3.6HG06030 | high | protein_cod | *-* | F-box family | duf 295 dom | no mercator | NA | PF03478 | IPR005174 |
| 1073 | HORVU.MOREX.r3.6HG06046 | high | protein_cod | *** | Hydroxycinr | EC_2.3 acyl | enzyme clas | GO:001674 | PF02458 | IPR003480 |
| 1074 | HORVU.MOREX.r3.6HG06049 | high | protein_cod | *** | F-box-like p | f box domai | no mercator | NA | PF12937 | IPR001810,IPR001810 |
| 1075 | HORVU.MOREX.r3.6HG06052 | low | transposon_ | *** | Transposon | osjnbb0016 | no mercator | NA | PF05340 | IPR008004 |
| 1076 | HORVU.MOREX.r3.6HG06052 | high | protein_cod | -** | EG5651 | not classifie | no mercator | NA | NA | NA |
| 1077 | HORVU.MOREX.r3.6HG06063 | high | protein_cod | *** | Peptide tran | anion transp | solute trans | GO:000521 | PF00854 | IPR000109,IPR020846,IPR02084 |
| 1078 | HORVU.MOREX.r3.6HG06069 | high | protein_cod | *** | Trihelix tran | TRIHELIX tra | rna biosynth | NA | NA | NA |
| 1079 | HORVU.MOREX.r3.6HG06070 | high | protein_cod | *** | Glycosyltrar | EC_2.4 glyc | enzyme clas | GO:000815 | PF00201 | IPR002213 |
| 1080 | HORVU.MOREX.r3.6HG06071 | high | protein_cod | *** | chitin synth | protein mod | no mercator | NA | PF06749 | IPR009606 |
| 1081 | HORVU.MOREX.r3.6HG06075 | high | protein_cod | *** | Adenine nuc | usp domain | no mercator | NA | PF00582 | IPR006016 |
| 1082 | HORVU.MOREX.r3.6HG06089 | high | protein_cod | *** | Lectin recep | EC_2.7 tran | enzyme clas | GO:000016 | PF00139,PF | IPR001220,IPR011009,IPR013320,IPR00124 |
| 1083 | HORVU.MOREX.r3.6HG06096 | high | protein_cod | *** | Cobyric acid | histone h3 2 | no mercator | NA | NA | NA |
| 1084 | HORVU.MOREX.r3.6HG06098 | low | protein_cod | -** | Transmemb | not classifie | no mercator | GO:001602 | NA | NA |
| 1085 | HORVU.MOREX.r3.6HG06099 | high | protein_cod | *** | Rho GDP-di | guanine nuc | multi-proce | GO:000509 | PF02115 | IPR000406,IPR014756 |
| 1086 | HORVU.MOREX.r3.6HG06102 | high | protein_cod | *** | Eukaryotic a | Pepsin-type | protein hom | GO:000419 | PF14543,PF | IPR021109,IPR032861,IPR032799 |
| 1087 | HORVU.MOREX.r3.6HG06122 | low | protein_cod | --* | ESAT-6-like | not classifie | no mercator | GO:000551 | NA | NA |
| 1088 | HORVU.MOREX.r3.6HG06124 | high | protein_cod | *** | transmemb | protein kina | no mercator | NA | NA | NA |
| 1089 | HORVU.MOREX.r3.6HG06129 | high | protein_cod | --* | Bifunctional | not classifie | no mercator | GO:000382 | NA | NA |
| 1090 | HORVU.MOREX.r3.6HG06130 | high | protein_cod | *** | DUF1005 fa | nuclear fact | no mercator | NA | PF06219 | IPR010410 |
| 1091 | HORVU.MOREX.r3.6HG06132 | high | protein_cod | *** | Blue copper | phytocyanin | no mercator | GO:000905 | PF02298 | IPR003245,IPR008972 |
| 1092 | HORVU.MOREX.r3.6HG06133 | high | protein_cod | *** | Chalcone sy | EC_2.3 acyl | enzyme clas | GO:000382 | PF02797,PF | IPR012328,IPR001099,IPR016039,IPR01603 |
| 1093 | HORVU.MOREX.r3.6HG06134 | high | protein_cod | *** | MACPF dom | macpf doma | no mercator | NA | PF01823 | IPR020864 |
| 1094 | HORVU.MOREX.r3.6HG06141 | high | protein_cod | *** | TPR repeat- | inactive tpr | no mercator | GO:004280 | PF13414,PF | IPR011990,IPR011990,IPR011990,IPR011990,IPR019734 |
| 1095 | HORVU.MOREX.r3.6HG06163 | high | protein_cod | *** | U-box doma | E3 ubiquitin | protein hom | GO:000484 | PF04564 | IPR003613,IPR016024 |
| 1096 | HORVU.MOREX.r3.6HG06168 | high | protein_cod | *** | Blue copper | phytocyanin | no mercator | GO:000905 | PF02298 | IPR003245,IPR008972 |
| 1097 | HORVU.MOREX.r3.6HG06169 | high | protein_cod | *** | Heat-shock | class-M-I sn | protein hom | NA | PF00011 | IPR008978,IPR002068 |
| 1098 | HORVU.MOREX.r3.6HG06170 | high | protein_cod | *** | Lipid transfe | aai domain | no mercator | GO:000686 | PF14368 | IPR016140,IPR016140 |
| 1099 | HORVU.MOREX.r3.6HG06171 | high | protein_cod | *** | Glycosyltrar | EC_2.4 glyc | enzyme clas | GO:000815 | PF00201 | IPR002213 |
| 1100 | HORVU.MOREX.r3.6HG06171 | high | protein_cod | *** | Glycosyltrar | EC_2.4 glyc | enzyme clas | GO:000815 | PF00201 | IPR002213 |
| 1101 | HORVU.MOREX.r3.6HG06172 | high | protein_cod | *-* | RNA-binding | glycine rich | no mercator | GO:000367 | PF00076 | IPR000504,IPR012677 |
| 1102 | HORVU.MOREX.r3.6HG06175 | high | protein_cod | *** | Ethylene-re | subgroup EF | rna biosynth | GO:000367 | PF00847 | IPR001471,IPR016177 |
| 1103 | HORVU.MOREX.r3.6HG06177 | high | protein_cod | *** | 2-oxoglutara | EC_1.14 oxi | enzyme clas | NA | PF03171,PF | IPR005123,IPR026992 |
| 1104 | HORVU.MOREX.r3.6HG06180 | high | protein_cod | --* | UDP-N-acet | thioredoxin | no mercator | GO:000382 | NA | NA |

| | | | | | | | | | | |
|---|---|---|---|---|---|---|---|---|---|---|
| 1105 | HORVU.MOREX.r3.6HG061812 | high | protein_cod | *** | Temperatur | lipocln cyto | no mercato | GO:000521 | PF08212 | IPR000566,IPR011038 |
| 1106 | HORVU.MOREX.r3.6HG061882 | high | protein_cod | *** | Protein kina | MAP3K-MEK | multi-proce | GO:000467 | PF00069 | IPR011009,IPR000719 |
| 1107 | HORVU.MOREX.r3.6HG061892 | high | protein_cod | *** | Chaperone | chaperone p | no mercato | GO:001602 | NA | NA |
| 1108 | HORVU.MOREX.r3.6HG061941 | high | protein_cod | *** | Flavonol syr | flavonol syn | secondary r | GO:001649 | PF03171,PF | IPR005123,IPR026992 |
| 1109 | HORVU.MOREX.r3.6HG062020 | high | protein_cod | *** | Kinase fami | RLCK-VIIa re | protein moc | GO:000467 | PF07714 | IPR001245,IPR011009 |
| 1110 | HORVU.MOREX.r3.6HG062072 | high | protein_cod | *** | Ribonuclea | T2-type RNa | rna process | GO:000372 | PF00445 | IPR001568,IPR001568 |
| 1111 | HORVU.MOREX.r3.6HG062193 | high | protein_cod | *** | Hexosyltran | hexosyltran | no mercato | GO:000013 | PF01762 | IPR002659 |
| 1112 | HORVU.MOREX.r3.6HG062270 | high | protein_cod | *** | Protein CYP | vid domain | no mercato | NA | PF08553 | IPR013863 |
| 1113 | HORVU.MOREX.r3.6HG062271 | high | protein_cod | *** | Dehydrin | dehydrin dh | no mercato | GO:000695 | PF00257,PF | IPR000167,IPR000167,IPR000167,IPR000167,IPR000167,IPR000167,IPR000167,IPR000167,IPR00016 |
| 1114 | HORVU.MOREX.r3.6HG062274 | high | protein_cod | *** | Protein DET | metabolite t | solute trans | GO:000685 | PF01554,PF | IPR002528,IPR002528,IPR002528 |
| 1115 | HORVU.MOREX.r3.6HG062292 | high | protein_cod | *** | Trehalose-6 | EC_2.4 glyc | enzyme clas | GO:000382 | PF00982,PF | IPR023214,IPR001830,IPR006379,IPR003337,IPR003337 |
| 1116 | HORVU.MOREX.r3.6HG062295 | low | protein_cod | --* | ER membra | not classifie | no mercato | GO:001602 | NA | NA |
| 1117 | HORVU.MOREX.r3.6HG062455 | high | protein_cod | *** | Alpha/beta- | protein de-S | protein moc | GO:001678 | IPR029058 | |
| 1118 | HORVU.MOREX.r3.6HG062532 | high | protein_cod | *** | Protein pho | DBP phosph | rna biosynth | GO:000382 | PF00481 | IPR001932,IPR001932 |
| 1119 | HORVU.MOREX.r3.6HG062563 | high | protein_cod | *** | Serine-rich | serine rich p | no mercato | NA | NA | NA |
| 1120 | HORVU.MOREX.r3.6HG062616 | high | protein_cod | *** | Alcohol deh | EC_1.1 oxid | enzyme clas | GO:000827 | PF00107,PF | IPR013149,IPR016040,IPR013154,IPR011032 |
| 1121 | HORVU.MOREX.r3.6HG062700 | high | protein_cod | *** | Auxin-respo | transcriptio | phytohormo | GO:000563 | PF02309,PF | IPR033389,IPR033389 |
| 1122 | HORVU.MOREX.r3.6HG062707 | high | protein_cod | *** | 3(2),5-bisph | phosphoad | coenzyme n | GO:000679 | PF00459 | IPR000760,IPR006239 |
| 1123 | HORVU.MOREX.r3.6HG062760 | high | protein_cod | *** | Protein kina | regulatory P | photosynthe | NA | PF00069 | IPR011009,IPR000719 |
| 1124 | HORVU.MOREX.r3.6HG062789 | low | non_coding | *** | Cytochrome | EC_1.14 oxi | enzyme clas | GO:000449 | PF00067 | IPR001128,IPR001128 |
| 1125 | HORVU.MOREX.r3.6HG062825 | high | protein_cod | *** | Cinnamoyl | cinnamoyl- | cell wall org | GO:000382 | PF01370 | IPR001509,IPR016040 |
| 1126 | HORVU.MOREX.r3.6HG062832 | high | protein_cod | *** | Germin-like | germin prot | no mercato | GO:000557 | PF00190 | IPR011051,IPR006045 |
| 1127 | HORVU.MOREX.r3.6HG062865 | low | protein_cod | *-* | BAG family | bag domain | no mercato | GO:005108 | PF02179 | IPR003103,IPR003103 |
| 1128 | HORVU.MOREX.r3.6HG062866 | high | protein_cod | *** | PPPDE thiol | deubiquitin | protein hom | NA | PF05903 | IPR008580 |
| 1129 | HORVU.MOREX.r3.6HG063025 | high | protein_cod | *** | Kinase fami | LysM protei | protein moc | GO:000467 | PF00069 | IPR000719,IPR011009 |
| 1130 | HORVU.MOREX.r3.6HG063065 | high | protein_cod | *** | Protein DET | metabolite t | solute trans | GO:000685 | PF01554,PF | IPR002528,IPR002528,IPR002528 |
| 1131 | HORVU.MOREX.r3.6HG063104 | high | protein_cod | *** | Mesoderm i | cargo recep | protein hom | GO:001602 | NA | NA |
| 1132 | HORVU.MOREX.r3.6HG063148 | low | non_coding | *** | Cytochrome | EC_1.14 oxi | enzyme clas | GO:000449 | PF00067 | IPR001128,IPR001128 |
| 1133 | HORVU.MOREX.r3.6HG063350 | high | protein_cod | *** | Response re | A-type ARR | phytohormo | GO:000016 | PF00072 | IPR011006,IPR001789 |
| 1134 | HORVU.MOREX.r3.7HG063472 | high | protein_cod | *** | RNA-binding | meiosis pro | no mercato | GO:000367 | PF04059,PF | IPR012677,IPR007201,IPR012677,IPR000504,IPR00050 |
| 1135 | HORVU.MOREX.r3.7HG063538 | high | protein_cod | *** | Acid inverta | vacuolar aci | carbohydra | GO:000455 | PF00251,PF | IPR023296,IPR013148,IPR013320,IPR021792,IPR01318 |
| 1136 | HORVU.MOREX.r3.7HG063657 | high | protein_cod | *** | Kelch repea | substrate ac | protein hom | NA | PF00646,PF | IPR001810,IPR001810,IPR006652,IPR006652 |
| 1137 | HORVU.MOREX.r3.7HG064112 | high | protein_cod | *** | Glutathione | class tau gl | redox home | GO:001674 | PF02798 | IPR004045,IPR012336,IPR010987 |
| 1138 | HORVU.MOREX.r3.7HG064264 | high | protein_cod | *** | Glycosyltra | EC_2.4 glyc | enzyme clas | GO:000815 | PF00201 | IPR002213 |
| 1139 | HORVU.MOREX.r3.7HG064578 | high | protein_cod | --* | Avr9/Cf-9 ra | not classifie | no mercato | NA | NA | NA |
| 1140 | HORVU.MOREX.r3.7HG064798 | high | protein_cod | *** | Coiled-coil | duf 2052 do | no mercato | NA | PF09747 | IPR018613 |
| 1141 | HORVU.MOREX.r3.7HG064799 | high | protein_cod | *** | Acid phosph | acid phosph | protein hom | GO:000399 | PF03767 | IPR023214,IPR005519,IPR010028 |
| 1142 | HORVU.MOREX.r3.7HG064812 | high | protein_cod | *** | Glutathione | class tau gl | redox home | GO:001674 | PF00043,PF | IPR010987,IPR004046,IPR004045,IPR01233 |
| 1143 | HORVU.MOREX.r3.7HG064846 | high | protein_cod | *** | Glycosyltra | EC_2.4 glyc | enzyme clas | NA | PF00201 | IPR002213 |
| 1144 | HORVU.MOREX.r3.7HG064865 | high | protein_cod | *** | Auxin repres | dormancy a | no mercato | NA | PF05564 | IPR008406 |
| 1145 | HORVU.MOREX.r3.7HG064902 | high | protein_cod | *** | Carbonic ar | alpha-type | photosynthe | GO:000408 | PF00194 | IPR001148,IPR001148 |
| 1146 | HORVU.MOREX.r3.7HG064924 | high | protein_cod | --* | protein kina | protein cad | no mercato | NA | NA | NA |
| 1147 | HORVU.MOREX.r3.7HG065107 | low | protein_cod | --* | Armadillo re | transcriptio | no mercato | GO:000563 | NA | NA |
| 1148 | HORVU.MOREX.r3.7HG065326 | high | protein_cod | *** | Transferase | EC_2.3 acyl | enzyme clas | GO:001674 | PF02458 | IPR003480 |
| 1149 | HORVU.MOREX.r3.7HG065426 | high | protein_cod | *** | Subtilisin-li | subtilisin pr | no mercato | GO:000425 | PF00082,PF | IPR000209,IPR000209,IPR000209,IPR000209,IPR01025 |
| 1150 | HORVU.MOREX.r3.7HG065664 | high | protein_cod | *** | Calcium-bir | calcium sen | multi-proce | GO:000550 | PF13405,PF | IPR002048,IPR002048,IPR011992 |
| 1151 | HORVU.MOREX.r3.7HG065667 | high | protein_cod | *** | Raffinose sy | galactinol-s | carbohydra | NA | PF05691 | IPR017853,IPR017853,IPR017853,IPR008811 |
| 1152 | HORVU.MOREX.r3.7HG065805 | high | protein_cod | *** | Acyl-coenzy | acyl CoA oxi | lipid metab | GO:000167 | PF02770,PF | IPR006091,IPR009075,IPR002655,IPR009100,IPR009075,IPR00907 |
| 1153 | HORVU.MOREX.r3.7HG065872 | high | protein_cod | *** | DUF868 fan | duf 868 fam | no mercato | NA | PF05910 | IPR008586 |
| 1154 | HORVU.MOREX.r3.7HG065928 | high | protein_cod | *** | Low temper | plasma mer | no mercato | GO:001602 | PF01679 | IPR000612 |
| 1155 | HORVU.MOREX.r3.7HG065971 | high | protein_cod | *** | ATP-depend | atp depend | no mercato | NA | NA | NA |
| 1156 | HORVU.MOREX.r3.7HG066072 | high | protein_cod | *** | Ring finger p | RING-H2-cl | protein hom | GO:001602 | PF13639 | IPR001841 |
| 1157 | HORVU.MOREX.r3.7HG066085 | high | protein_cod | *-* | LL-diamino | not classifie | no mercato | GO:000848 | NA | NA |
| 1158 | HORVU.MOREX.r3.7HG066157 | high | protein_cod | *** | Chaperone | chaperone p | no mercato | NA | PF00226 | IPR001623,IPR001623 |
| 1159 | HORVU.MOREX.r3.7HG066191 | high | protein_cod | *** | Zinc-finger p | ZAT transcri | rna biosynth | GO:000367 | PF13912,PF | IPR007087,IPR007087 |
| 1160 | HORVU.MOREX.r3.7HG066213 | high | protein_cod | *** | DUF1677 fa | ribosomal l | no mercato | NA | PF07911 | IPR012876 |
| 1161 | HORVU.MOREX.r3.7HG066238 | high | protein_cod | *** | Pathogenes | bet v 1 dom | no mercato | GO:000695 | PF00407 | IPR000916 |
| 1162 | HORVU.MOREX.r3.7HG066242 | high | protein_cod | *** | Kinase-like | RLCK-VIIa re | protein moc | GO:000467 | PF07714 | IPR011009,IPR001245 |
| 1163 | HORVU.MOREX.r3.7HG066303 | low | protein_cod | --* | DNA-bindin | not classifie | no mercato | NA | NA | NA |
| 1164 | HORVU.MOREX.r3.7HG066327 | high | protein_cod | *** | RNA ligase/ | rna ligase c | no mercato | GO:000411 | IPR009097 | |
| 1165 | HORVU.MOREX.r3.7HG066387 | high | protein_cod | *** | Transmemb | tf b3 domai | no mercato | GO:001602 | PF05705 | IPR029058,IPR029058,IPR008547 |
| 1166 | HORVU.MOREX.r3.7HG066399 | high | protein_cod | *** | Scarecrow t | GRAS trans | rna biosynth | GO:000370 | PF03514 | IPR005202 |
| 1167 | HORVU.MOREX.r3.7HG066403 | high | protein_cod | *-* | AR781, phe | not classifie | no mercato | NA | PF07816 | IPR012442 |
| 1168 | HORVU.MOREX.r3.7HG066425 | high | protein_cod | *** | 12-oxophyt | 4,5-didehyd | phytohormo | GO:000382 | PF00724 | IPR001155 |
| 1169 | HORVU.MOREX.r3.7HG066427 | high | protein_cod | *** | 12-oxophyt | 4,5-didehyd | phytohormo | GO:000382 | PF00724 | IPR001155 |

| | | | | | | | | | | |
|---|---|---|---|---|---|---|---|---|---|---|
| 1170 | HORVU.MOREX.r3.7HG066428 | high | protein_cod | *** | Pentatricop | pentatricop | no mercato | NA | PF13041,PF | IPR002885,IPR002885,IPR002885,IPR002885,IPR011990,IPR002885,IPR002885,IPR002885,IPR002885,IPR002885,IPR002885,IPR00288 |
| 1171 | HORVU.MOREX.r3.7HG066470 | high | protein_cod | *** | RING/U-box | RING-H2-cl | protein hom | GO:001602 | PF13639 | IPR001841 |
| 1172 | HORVU.MOREX.r3.7HG066484 | high | protein_cod | *** | Heat-shock | shsp domai | no mercato | NA | PF00011 | IPR008978,IPR002068 |
| 1173 | HORVU.MOREX.r3.7HG066518 | high | protein_cod | *** | Glycosyltra | EC_2.4 glyc | enzyme clas | GO:000815 | PF00201 | IPR002213 |
| 1174 | HORVU.MOREX.r3.7HG066599 | high | protein_cod | *** | Shikimate k | shikimate ki | amino acid | GO:001630 | PF01202 | IPR031322,IPR027417 |
| 1175 | HORVU.MOREX.r3.7HG066655 | high | protein_cod | *** | Amino acid | amino acid | solute trans | GO:001602 | PF01490 | IPR013057 |
| 1176 | HORVU.MOREX.r3.7HG066674 | high | protein_cod | *** | Glycosyltra | EC_2.4 glyc | enzyme clas | GO:000815 | PF00201 | IPR002213 |
| 1177 | HORVU.MOREX.r3.7HG066682 | high | protein_cod | *** | RING/U-box | RING-H2-cl | protein hom | NA | PF13639 | IPR001841 |
| 1178 | HORVU.MOREX.r3.7HG066708 | high | protein_cod | *** | Serine/threo | bhlh myc n | no mercato | GO:001630 | PF14215 | IPR025610 |
| 1179 | HORVU.MOREX.r3.7HG066762 | high | protein_cod | *** | Peroxidase | EC_1.11 oxi | enzyme clas | GO:000460 | PF00141 | IPR010255,IPR002016 |
| 1180 | HORVU.MOREX.r3.7HG066763 | high | protein_cod | *** | Peroxidase | NA | NA | GO:000460 | PF00141 | IPR010255,IPR002016 |
| 1181 | HORVU.MOREX.r3.7HG066766 | high | protein_cod | *** | Peroxidase | EC_1.11 oxi | enzyme clas | GO:000460 | PF00141 | IPR010255,IPR002016 |
| 1182 | HORVU.MOREX.r3.7HG066768 | high | protein_cod | *** | Peroxidase | NA | NA | GO:000460 | PF00141 | IPR010255,IPR002016 |
| 1183 | HORVU.MOREX.r3.7HG066804 | high | protein_cod | *-* | CDP-diacylg | peptidase m | no mercato | GO:001674 | NA | NA |
| 1184 | HORVU.MOREX.r3.7HG066821 | high | protein_cod | *** | Coiled-coil | chitin elicito | no mercato | NA | NA | NA |
| 1185 | HORVU.MOREX.r3.7HG066888 | high | protein_cod | *** | Pathogenes | CAPE precu | phytohormo | GO:000557 | PF00188 | IPR014044,IPR014044 |
| 1186 | HORVU.MOREX.r3.7HG066890 | high | protein_cod | *** | Pathogenes | CAPE precu | phytohormo | GO:000557 | PF00188 | IPR014044,IPR014044 |
| 1187 | HORVU.MOREX.r3.7HG066894 | high | protein_cod | *** | U-box doma | E3 ubiquitin | protein hom | GO:000484 | PF04564 | IPR003613,IPR016024 |
| 1188 | HORVU.MOREX.r3.7HG067035 | high | protein_cod | *** | Plastid mov | tify domain | no mercato | NA | PF14009 | IPR025322 |
| 1189 | HORVU.MOREX.r3.7HG067069 | high | protein_cod | *** | HIPL1 prote | gsdh domai | no mercato | GO:000382 | PF07995 | IPR011041,IPR011041,IPR011041,IPR012938 |
| 1190 | HORVU.MOREX.r3.7HG067090 | high | protein_cod | *** | Aspartic pro | Pepsin-type | protein hom | GO:000419 | PF14543,PF | IPR032861,IPR032799,IPR021109 |
| 1191 | HORVU.MOREX.r3.7HG067489 | high | protein_cod | *** | Vacuolar pr | regulatory c | vesicle traff | NA | NA | NA |
| 1192 | HORVU.MOREX.r3.7HG067531 | low | protein_cod | --* | N-lysine me | not classifie | no mercato | NA | NA | NA |
| 1193 | HORVU.MOREX.r3.7HG067539 | high | protein_cod | *** | Thioredoxin | duf 953 dom | no mercato | NA | PF06110 | IPR012336,IPR010357 |
| 1194 | HORVU.MOREX.r3.7HG067575 | low | protein_cod | *-* | BAG family | bag domain | no mercato | GO:005108 | PF02179 | IPR003103,IPR003103 |
| 1195 | HORVU.MOREX.r3.7HG067593 | low | protein_cod | --* | molybdenu | not classifie | no mercato | NA | NA | NA |
| 1196 | HORVU.MOREX.r3.7HG067604 | high | protein_cod | --* | Decaprenyl- | not classifie | no mercato | GO:000028 | NA | NA |
| 1197 | HORVU.MOREX.r3.7HG067613 | high | protein_cod | *** | Diacylglyce | acyl-CoA:di | lipid metab | GO:001602 | PF03982 | IPR007130 |
| 1198 | HORVU.MOREX.r3.7HG067619 | low | protein_cod | --* | BAG family | bag domain | no mercato | GO:005108 | NA | NA |
| 1199 | HORVU.MOREX.r3.7HG067620 | low | protein_cod | --* | Dynamin-1 | bag domain | no mercato | GO:000016 | NA | NA |
| 1200 | HORVU.MOREX.r3.7HG067748 | high | protein_cod | *** | Protein O-li | xylan alpha- | cell wall org | GO:001602 | PF04577 | IPR007657 |
| 1201 | HORVU.MOREX.r3.7HG067757 | high | protein_cod | *** | Glycosyltra | egf domain | no mercato | GO:001602 | PF04577 | IPR007657 |
| 1202 | HORVU.MOREX.r3.7HG067799 | high | protein_cod | *** | Transducin/ | substrate ad | protein hom | NA | PF00400 | IPR001680,IPR011047 |
| 1203 | HORVU.MOREX.r3.7HG067813 | high | protein_cod | *** | Caffeoyl-Co | caffeoyl-Co. | cell wall org | GO:000816 | PF01596 | IPR029063,IPR002935 |
| 1204 | HORVU.MOREX.r3.7HG067814 | high | protein_cod | --* | Cortactin-bi | caffeoyl coa | no mercato | GO:000573 | NA | NA |
| 1205 | HORVU.MOREX.r3.7HG067853 | high | protein_cod | *-* | Ubiquitin-as | ubiquitin an | no mercato | NA | PF08325,PF | IPR013536,IPR018997,IPR029071,IPR018997 |
| 1206 | HORVU.MOREX.r3.7HG067952 | high | protein_cod | *** | Zinc finger A | zinc finger a | no mercato | GO:000367 | PF01754,PF | IPR002653,IPR000058 |
| 1207 | HORVU.MOREX.r3.7HG068055 | high | protein_cod | *** | AT hook mo | AHL clade-E | rna biosynth | NA | PF03479 | IPR005175 |
| 1208 | HORVU.MOREX.r3.7HG068195 | high | protein_cod | *** | Chorismate | chorismate | amino acid | GO:000410 | IPR008238,IPR020822 | |
| 1209 | HORVU.MOREX.r3.7HG068222 | high | protein_cod | --* | NAD(P)-bin | regulatory p | multi-proce | NA | NA | NA |
| 1210 | HORVU.MOREX.r3.7HG068229 | low | protein_cod | --* | NAC (No Ap | SIM-type cy | cell division | NA | NA | NA |
| 1211 | HORVU.MOREX.r3.7HG068250 | low | non_coding | *** | Cytochrome | cytochrome | no mercato | GO:001602 | PF04526,PF | IPR005018,IPR006593 |
| 1212 | HORVU.MOREX.r3.7HG068482 | high | protein_cod | *** | NAC domai | NAC transcr | rna biosynth | GO:000367 | PF02365 | IPR003441,IPR003441 |
| 1213 | HORVU.MOREX.r3.7HG068523 | high | protein_cod | *-* | Ethylene-re | subgroup EF | rna biosynth | GO:000367 | PF00847 | IPR001471,IPR016177 |
| 1214 | HORVU.MOREX.r3.7HG068537 | high | protein_cod | *** | Stress up-re | stress up re | no mercato | NA | PF07712 | IPR011692 |
| 1215 | HORVU.MOREX.r3.7HG068658 | high | protein_cod | *-* | Transcriptio | bZIP class-I | rna biosynth | GO:000370 | PF07716 | IPR004827 |
| 1216 | HORVU.MOREX.r3.7HG068679 | high | protein_cod | *** | Alcohol deh | sorbitol deh | carbohydrat | GO:000393 | PF08240,PF | IPR011032,IPR016040,IPR013154,IPR013149 |
| 1217 | HORVU.MOREX.r3.7HG068746 | high | protein_cod | *** | Chaperone | dnaj domair | no mercato | NA | PF00226 | IPR001623,IPR001623 |
| 1218 | HORVU.MOREX.r3.7HG068794 | high | protein_cod | *** | DUF241 dor | not classifie | no mercato | NA | PF03087 | IPR004320 |
| 1219 | HORVU.MOREX.r3.7HG068883 | low | transposon_ | *-* | Retrotransp | proline rich | no mercato | GO:000367 | NA | NA |
| 1220 | HORVU.MOREX.r3.7HG068887 | high | protein_cod | *** | Monodehyd | monodehyd | redox home | GO:001649 | PF07992 | IPR023753,IPR023753,IPR016156,IPR023753 |
| 1221 | HORVU.MOREX.r3.7HG069024 | high | protein_cod | *** | 2-oxoglutar | EC_1.14 oxi | enzyme clas | GO:001649 | PF14226,PF | IPR026992,IPR005123 |
| 1222 | HORVU.MOREX.r3.7HG070003 | high | protein_cod | *** | Nitrate tran | anion transp | solute trans | GO:000521 | PF00854 | IPR020846,IPR020846,IPR020846,IPR000109 |
| 1223 | HORVU.MOREX.r3.7HG070207 | high | protein_cod | *** | Blue copper | phytocyanin | no mercato | GO:000905 | PF02298 | IPR003245,IPR008972 |
| 1224 | HORVU.MOREX.r3.7HG070284 | high | protein_cod | *** | Zinc transp | metal cation | solute trans | GO:001602 | PF01544 | IPR002523 |
| 1225 | HORVU.MOREX.r3.7HG070346 | high | protein_cod | *** | Amino acid | amino acid | solute trans | GO:001602 | PF01490 | IPR013057 |
| 1226 | HORVU.MOREX.r3.7HG070349 | high | protein_cod | *** | Amino acid | amino acid | solute trans | GO:001602 | PF01490 | IPR013057 |
| 1227 | HORVU.MOREX.r3.7HG070481 | high | protein_cod | *** | L-gulonolac | L-gulono-1,4 | redox home | GO:000388 | PF01565,PF | IPR006094,IPR007173,IPR016166,IPR010030 |
| 1228 | HORVU.MOREX.r3.7HG070725 | high | protein_cod | *** | Tubby-like F | stress-resp | external stir | NA | PF00646,PF | IPR001810,IPR001810,IPR025659,IPR025659,IPR025659,IPR000007 |
| 1229 | HORVU.MOREX.r3.7HG070726 | low | protein_cod | --* | 2-oxoglutar | not classifie | no mercato | NA | NA | NA |
| 1230 | HORVU.MOREX.r3.7HG070835 | high | protein_cod | *-* | Crooked ne | not classifie | no mercato | GO:000639 | IPR011990 | |
| 1231 | HORVU.MOREX.r3.7HG070896 | high | protein_cod | *** | VQ motif fa | regulatory p | rna biosynth | NA | PF05678 | IPR008889 |
| 1232 | HORVU.MOREX.r3.7HG070958 | high | protein_cod | *** | ADP-ribosyl | small GTPas | vesicle traff | GO:000016 | PF00025 | IPR027417,IPR005225,IPR006689 |
| 1233 | HORVU.MOREX.r3.7HG071056 | high | protein_cod | *** | WRKY trans | transcriptio | rna biosynth | GO:000367 | PF03106 | IPR003657,IPR003657 |
| 1234 | HORVU.MOREX.r3.7HG071189 | high | protein_cod | *** | O-methyltra | caffeic acid | cell wall org | GO:000816 | PF08100,PF | IPR012967,IPR029063,IPR011991,IPR001077 |

| | | | | | | | | | | |
|---|---|---|---|---|---|---|---|---|---|---|
| 1235 | HORVU.MOREX.r3.7HG071245 | high | protein_cod | *** | Protein kina | MAP3K-MEK | multi-proce | GO:0000166 | PF00069 | IPR011009,IPR000719 |
| 1236 | HORVU.MOREX.r3.7HG071258 | high | protein_cod | *** | Ribonucleas | endoribonu | no mercato | GO:000452 | PF00636 | IPR000999,IPR000999 |
| 1237 | HORVU.MOREX.r3.7HG071337 | low | non_coding | *** | Cytochrome | EC_1.14 oxi | enzyme clas | GO:0004497 | PF00067 | IPR001128,IPR001128 |
| 1238 | HORVU.MOREX.r3.7HG071385 | high | protein_cod | --* | multidrug re | mapk kinase | no mercato | NA | PF15697 | IPR031421 |
| 1239 | HORVU.MOREX.r3.7HG071394 | high | protein_cod | *** | GTP-binding | GTPase *(R | protein tran | GO:0000166 | PF00071 | IPR005225,IPR027417,IPR001806 |
| 1240 | HORVU.MOREX.r3.7HG071414 | high | protein_cod | *** | 3-ketoacyl-( | 3-ketoacyl-( | lipid metab | GO:000382 | PF08541,PF | IPR016039,IPR016039,IPR013747,IPR013601 |
| 1241 | HORVU.MOREX.r3.7HG071435 | high | protein_cod | *** | Acyl-[acyl-c | palmitoyl-A | lipid metab | GO:000662 | PF12590,PF | IPR021113,IPR029069,IPR029069,IPR002864 |
| 1242 | HORVU.MOREX.r3.7HG071475 | high | protein_cod | *-* | Kelch repea | substrate(P | secondary r | NA | PF01344,PF | IPR006652,IPR006652,IPR001810 |
| 1243 | HORVU.MOREX.r3.7HG071497 | high | protein_cod | *** | Multiprotein | transcriptio | rna biosynth | GO:0003677 | PF08523,PF | IPR010982,IPR013729,IPR001387 |
| 1244 | HORVU.MOREX.r3.7HG071539 | low | protein_cod | --* | Tetratricope | formin prote | no mercato | GO:000367 | NA | NA |
| 1245 | HORVU.MOREX.r3.7HG071549 | high | protein_cod | --* | SWIB/MDM2 | early noduli | no mercato | NA | NA | NA |
| 1246 | HORVU.MOREX.r3.7HG071582 | high | protein_cod | *** | Transmemb | phd type do | no mercato | GO:001602 | PF07343 | IPR009943 |
| 1247 | HORVU.MOREX.r3.7HG071589 | high | protein_cod | *** | Nitrate trans | abscisic aci | phytohormo | NA | PF00854 | IPR000109,IPR020846,IPR020846 |
| 1248 | HORVU.MOREX.r3.7HG071595 | high | protein_cod | *** | cysteine-ric | not classifie | no mercato | NA | PF12734 | IPR028144 |
| 1249 | HORVU.MOREX.r3.7HG071664 | high | protein_cod | *** | Xyloglucan ( | xyloglucan e | cell wall org | GO:000455 | PF06955,PF | IPR013320,IPR010713,IPR000757 |
| 1250 | HORVU.MOREX.r3.7HG071782 | high | protein_cod | *** | Abscisic aci | receptor co | phytohormo | NA | PF10604 | IPR019587 |
| 1251 | HORVU.MOREX.r3.7HG071795 | high | protein_cod | *-* | Oxidative st | regulatory p | multi-proce | NA | NA | NA |
| 1252 | HORVU.MOREX.r3.7HG071813 | high | protein_cod | *** | Amino acid | amino acid t | solute trans | GO:001602 | PF01490 | IPR013057 |
| 1253 | HORVU.MOREX.r3.7HG071893 | high | protein_cod | *** | Peroxidase | EC_1.11 oxi | enzyme clas | GO:000460 | PF00141 | IPR002016,IPR010255 |
| 1254 | HORVU.MOREX.r3.7HG072161 | high | protein_cod | *-* | senescence | regulatory p | multi-proce | NA | PF04570 | IPR007650 |
| 1255 | HORVU.MOREX.r3.7HG072164 | high | protein_cod | *** | Kinase fami | RLCK-VIIa re | protein mod | GO:0000166 | PF00069 | IPR011009,IPR000719 |
| 1256 | HORVU.MOREX.r3.7HG072166 | high | protein_cod | --* | Serine/thre | not classifie | no mercato | GO:000013 | NA | NA |
| 1257 | HORVU.MOREX.r3.7HG072183 | high | protein_cod | *** | Germin-like | germin prot | no mercato | GO:0005570 | PF00190 | IPR011051,IPR006045 |
| 1258 | HORVU.MOREX.r3.7HG072256 | high | protein_cod | *** | Pleiotropic ( | subfamily A | solute trans | GO:0000166 | PF14510,PF | IPR029481,IPR013525,IPR013525,IPR027417,IPR003439,IPR003439,IPR013581,IPR02741 |
| 1259 | HORVU.MOREX.r3.7HG072324 | high | protein_cod | *** | TMV respon | tmv respons | no mercato | NA | PF14009 | IPR025322 |
| 1260 | HORVU.MOREX.r3.7HG072326 | high | protein_cod | *** | TMV respon | tmv respons | no mercato | NA | PF14009 | IPR025322 |
| 1261 | HORVU.MOREX.r3.7HG072354 | high | protein_cod | *** | Cinnamoyl- | phaseic acid | phytohormo | GO:000382 | PF01370 | IPR016040,IPR001509 |
| 1262 | HORVU.MOREX.r3.7HG072388 | low | non_coding | *** | Cytochrome | EC_1.14 oxi | enzyme clas | GO:0004497 | PF00067 | IPR001128,IPR001128 |
| 1263 | HORVU.MOREX.r3.7HG072475 | low | non_coding | *** | Cytochrome | 2-hydroxyis | secondary r | GO:0003677 | PF00067 | IPR001128,IPR001128 |
| 1264 | HORVU.MOREX.r3.7HG072537 | high | protein_cod | *** | Kelch repea | substrate ac | phytohormo | NA | NA | NA |
| 1265 | HORVU.MOREX.r3.7HG072544 | high | protein_cod | *-* | Pre-mRNA-s | not classifie | no mercato | NA | NA | NA |
| 1266 | HORVU.MOREX.r3.7HG072709 | high | protein_cod | --* | tRNA (guani | not classifie | no mercato | GO:0006400 | NA | NA |
| 1267 | HORVU.MOREX.r3.7HG072950 | high | protein_cod | *** | Zinc finger p | LSU process | protein bios | GO:0003676 | PF12756 | IPR015880 |
| 1268 | HORVU.MOREX.r3.7HG072976 | high | protein_cod | *** | Receptor pr | EC_2.7 tran | enzyme clas | GO:000467 | PF00069 | IPR000719,IPR011009 |
| 1269 | HORVU.MOREX.r3.7HG073053 | high | protein_cod | -** | Copper tran | copper chap | nutrient upt | NA | PF00403 | IPR006121,IPR006121 |
| 1270 | HORVU.MOREX.r3.7HG073087 | high | protein_cod | *** | Glycosyltrar | xylosyltrans | cell wall org | GO:000013 | PF03360 | IPR029044,IPR005027 |
| 1271 | HORVU.MOREX.r3.7HG073122 | high | protein_cod | *** | 2-oxoglutar | EC_1.14 oxi | enzyme clas | GO:001649 | PF03171,PF | IPR005123,IPR026992 |
| 1272 | HORVU.MOREX.r3.7HG073200 | high | protein_cod | *** | Wound-resp | not classifie | no mercato | NA | PF12609 | IPR022251 |
| 1273 | HORVU.MOREX.r3.7HG073202 | low | non_coding | *** | Cytochrome | EC_1.14 oxi | enzyme clas | GO:0004497 | PF00067 | IPR001128,IPR001128 |
| 1274 | HORVU.MOREX.r3.7HG073212 | high | protein_cod | *** | Glycosyltrar | EC_2.4 glyc | enzyme clas | GO:000815 | PF00201 | IPR002213 |
| 1275 | HORVU.MOREX.r3.7HG073286 | high | protein_cod | *** | Polyubiquiti | UBQ ubiquit | protein hom | NA | PF00240,PF | IPR029071,IPR029071,IPR000626,IPR000626,IPR000626,IPR000626,IPR029071,IPR02907 |
| 1276 | HORVU.MOREX.r3.7HG073302 | high | protein_cod | *** | Polyubiquiti | UBQ ubiquit | protein hom | NA | PF00240,PF | IPR029071,IPR029071,IPR029071,IPR029071,IPR000626,IPR000626,IPR000626,IPR000626,IPR000626,IPR02907 |
| 1277 | HORVU.MOREX.r3.7HG073307 | high | protein_cod | *** | Polyubiquiti | UBQ ubiquit | protein hom | NA | PF00240,PF | IPR000626,IPR000626,IPR000626,IPR000626,IPR000626,IPR000626,IPR029071,IPR029071,IPR029071,IPR029071,IPR029071,IPR02907 |
| 1278 | HORVU.MOREX.r3.7HG073311 | high | protein_cod | *** | Polyubiquiti | UBQ ubiquit | protein hom | NA | PF00240,PF | IPR029071,IPR029071,IPR000626,IPR000626,IPR000626,IPR000626,IPR000626,IPR000626,IPR029071,IPR029071,IPR02907 |
| 1279 | HORVU.MOREX.r3.7HG073348 | high | protein_cod | *** | Carbonic ar | beta-type ca | photosynthe | GO:0004089 | PF00484 | IPR001765,IPR001765 |
| 1280 | HORVU.MOREX.r3.7HG073349 | high | protein_cod | *** | Carbonic ar | beta-type ca | photosynthe | GO:0004089 | PF00484 | IPR001765,IPR001765 |
| 1281 | HORVU.MOREX.r3.7HG073752 | high | protein_cod | *** | Kinase-like | RLCK-VIIa re | protein mod | GO:000467 | PF07714 | IPR001245,IPR011009 |
| 1282 | HORVU.MOREX.r3.7HG073757 | high | protein_cod | *** | Elongation c | fatty acid el | lipid metab | GO:001602 | PF01151 | IPR002076 |
| 1283 | HORVU.MOREX.r3.7HG073801 | high | protein_cod | *** | VQ motif far | regulatory p | rna biosynth | NA | PF05678 | IPR008889 |
| 1284 | HORVU.MOREX.r3.7HG073929 | high | protein_cod | *-* | Serine/argin | serine threo | no mercato | NA | NA | NA |
| 1285 | HORVU.MOREX.r3.7HG074020 | high | protein_cod | --* | Pectin lyase | osjnba0041 | no mercato | NA | NA | NA |
| 1286 | HORVU.MOREX.r3.7HG074050 | high | protein_cod | *** | Kelch repea | substrate ac | external stir | GO:000958 | PF00646 | IPR001810,IPR001810 |
| 1287 | HORVU.MOREX.r3.7HG074211 | high | protein_cod | *** | Heptahelica | heptahelica | no mercato | GO:001602 | PF03006 | IPR004254 |
| 1288 | HORVU.MOREX.r3.7HG074244 | high | protein_cod | *** | Chitinase | basic chitin | external stir | GO:000456 | PF00182,PF | IPR023346,IPR001002,IPR000726,IPR001002 |
| 1289 | HORVU.MOREX.r3.7HG074252 | high | protein_cod | *** | Dihydroflav | epimerase c | no mercato | NA | PF01370 | IPR001509,IPR016040 |
| 1290 | HORVU.MOREX.r3.7HG074280 | high | protein_cod | *-* | Chaperone | sp dnaj chap | no mercato | NA | PF00226 | IPR001623,IPR001623 |
| 1291 | HORVU.MOREX.r3.7HG074451 | high | protein_cod | --* | Disease res | nb arc doma | no mercato | NA | PF00931 | IPR032675,IPR002182,IPR027417 |
| 1292 | HORVU.MOREX.r3.7HG074460 | high | protein_cod | *** | Carbonic ar | beta-type ca | photosynthe | GO:0004089 | PF00484 | IPR001765,IPR001765 |
| 1293 | HORVU.MOREX.r3.7HG074494 | high | protein_cod | *** | Calcium-bir | calcium sen | multi-proce | GO:0005509 | PF13833,PF | IPR002048,IPR002048,IPR011992 |
| 1294 | HORVU.MOREX.r3.7HG074495 | high | protein_cod | *** | Calcium-bir | calcium bin | no mercato | GO:0005509 | PF13833,PF | IPR002048,IPR002048,IPR011992 |
| 1295 | HORVU.MOREX.r3.7HG074521 | high | protein_cod | *** | NF-X1-type | NF-X transc | rna biosynth | GO:0003700 | PF01422,PF | IPR000967,IPR000967,IPR000967,IPR000967,IPR000967,IPR000967,IPR000967,IPR00096 |
| 1296 | HORVU.MOREX.r3.7HG074579 | high | protein_cod | *** | Ubiquitin sy | cue domain | no mercato | NA | NA | NA |
| 1297 | HORVU.MOREX.r3.7HG074664 | high | protein_cod | *** | 1-aminocyc | 1-aminocyc | phytohormo | GO:001649 | PF14226,PF | IPR026992,IPR005123 |
| 1298 | HORVU.MOREX.r3.7HG074667 | high | protein_cod | *** | DUF668 fan | protein psk | no mercato | GO:0005880 | PF11961,PF | IPR021864,IPR007700 |
| 1299 | HORVU.MOREX.r3.7HG074668 | high | protein_cod | *-* | Chaperone | sp dnaj chap | no mercato | NA | PF00226 | IPR001623,IPR001623 |

| | | | | | | | | | | |
|---|---|---|---|---|---|---|---|---|---|---|
| 1300 | HORVU.MOREX.r3.7HG07470 | high | protein_cod | *-* | Kinase fami | E3 ubiquitin | protein hom | GO:000467 | PF00069 | IPR011009,IPR000719 |
| 1301 | HORVU.MOREX.r3.7HG07470 | high | protein_cod | *** | Senescence | senescence | no mercato | NA | PF06911 | IPR009686 |
| 1302 | HORVU.MOREX.r3.7HG07476 | high | protein_cod | *** | Eukaryotic a | Pepsin-type | protein hom | GO:000419 | PF14543,PF | IPR021109,IPR032861,IPR032799 |
| 1303 | HORVU.MOREX.r3.7HG07480 | high | protein_cod | *-* | Heavy meta | hma domair | no mercato | GO:003000 | PF00403 | IPR006121,IPR006121 |
| 1304 | HORVU.MOREX.r3.7HG07480 | high | protein_cod | *-* | Heavy meta | hma domair | no mercato | GO:003000 | PF00403 | IPR006121,IPR006121 |
| 1305 | HORVU.MOREX.r3.7HG07496 | high | protein_cod | *** | Transmemb | transmemb | no mercato | GO:001602 | PF05705 | IPR029058,IPR029058,IPR008547 |
| 1306 | HORVU.MOREX.r3.7HG07498 | low | protein_cod | --* | Protein kina | not classifie | no mercato | NA | NA | NA |
| 1307 | HORVU.MOREX.r3.7HG07509 | high | protein_cod | *** | Dirigent pro | dirigent prot | no mercato | GO:000557 | PF03018,PF | IPR004265,IPR001229,IPR001229 |
| 1308 | HORVU.MOREX.r3.7HG07513 | high | protein_cod | *-* | Ricin B lecti | ricin b lectir | no mercato | GO:003024 | IPR000772,IPR000772 | |
