## Supplementary Table 2 for "A single-cell transcriptome atlas of the barley root apical meristem uncovers conserved and divergent roles of HvWOX5"

**Supplementary Table 2. Top10 marker genes per cluster based on log2FC.**

| No. | gene | X | p_val | avg_log2FC | pct.1 | pct.2 | p_val_adj | cluster | Cell type | ID | Confidence | Biotype | Score | Description | Ontology | pfam | interpro |
|---|---|---|---|---|---|---|---|---|---|---|---|---|---|---|---|---|---|
| 22 | HORVU.MOR | HORVU.MOR | 4.30E-69 | 4.065635817 | 0.332 | 0.088 | 5.37E-65 | 0 | unidentified | HORVU.MOR | low | protein_codir | --* | pH-response re | NA | NA | NA |
| 33 | HORVU.MOR | HORVU.MOR | 1.49E-63 | 4.20526718 | 0.324 | 0.094 | 1.86E-59 | 0 | unidentified | HORVU.MOR | high | protein_codir | *** | Threonine--tRN | GO:0000166, | PF00587,PF0 | IPR012676,IPR002314,IPR018163,IPR012947,IPR004095 |
| 36 | HORVU.MOR | HORVU.MOR | 4.71E-126 | 4.371701669 | 0.308 | 0.038 | 5.88E-122 | 0 | unidentified | HORVU.MOR | low | transposon_r | *-* | Transposon pro | NA | PF03108 | IPR004332 |
| 49 | HORVU.MOR | HORVU.MOR | 2.69E-88 | 4.253820785 | 0.281 | 0.048 | 3.36E-84 | 0 | unidentified | HORVU.MOR | high | protein_codir | --* | myosin heavy c | NA | NA | NA |
| 54 | HORVU.MOR | HORVU.MOR | 1.72E-101 | 4.372008588 | 0.326 | 0.051 | 2.15E-97 | 0 | unidentified | HORVU.MOR | low | protein_codir | --* | alpha/beta-Hy | NA | NA | NA |
| 71 | HORVU.MOR | HORVU.MOR | 7.59E-22 | 7.053516241 | 0.294 | 0.089 | 9.47E-18 | 0 | unidentified | HORVU.MOR | low | transposon_r | *-* | Transposon Ty3 | GO:0003676, | NA | NA |
| 158 | HORVU.MOR | HORVU.MOR | 9.52E-32 | 5.08903966 | 0.316 | 0.071 | 1.19E-27 | 0 | unidentified | HORVU.MOR | high | protein_codir | --* | Helix-loop-heli | GO:0003677, | NA | NA |
| 170 | HORVU.MOR | HORVU.MOR | 2.25E-26 | 4.484076216 | 0.281 | 0.094 | 2.81E-22 | 0 | unidentified | HORVU.MOR | high | protein_codir | *** | Protein TolB | NA | NA | NA |
| 203 | HORVU.MOR | HORVU.MOR | 3.01E-15 | 4.355197453 | 0.34 | 0.154 | 3.75E-11 | 0 | unidentified | HORVU.MOR | high | protein_codir | *** | Beta-glucan bi | GO:0052861 | PF03639 | IPR005200 |
| 218 | HORVU.MOR | HORVU.MOR | 5.85E-64 | 4.117545241 | 0.368 | 0.121 | 7.31E-60 | 0 | unidentified | HORVU.MOR | high | protein_codir | *** | Receptor prote | GO:0000165, | PF13855,PF1 | IPR011009,IPR001611,IPR001611,IPR001611,IPR032675,IPR000719,IPR001611 |
| 30 | HORVU.MOR | HORVU.MOR | 1.70E-154 | 4.688483469 | 0.546 | 0.095 | 2.13E-150 | 1 | Endodermis | HORVU.MOR | low | protein_codir | --* | Serine/threonir | GO:0000159, | NA | NA |
| 31 | HORVU.MOR | HORVU.MOR | 1.99E-48 | 4.721740989 | 0.526 | 0.141 | 2.49E-44 | 1 | Endodermis | HORVU.MOR | high | protein_codir | *** | Polyadenylate- | GO:0003676, | PF00076 | IPR000504,IPR012677 |
| 76 | HORVU.MOR | HORVU.MOR | 1.06E-84 | 4.925193203 | 0.494 | 0.129 | 1.33E-80 | 1 | Endodermis | HORVU.MOR | low | transposon_r | *-* | Retrotranspos | GO:0003676, | NA | NA |
| 117 | HORVU.MOR | HORVU.MOR | 8.25E-99 | 5.972905772 | 0.537 | 0.147 | 1.03E-94 | 1 | Endodermis | HORVU.MOR | low | protein_codir | --* | S-adenosyl-L-r | NA | NA | NA |
| 123 | HORVU.MOR | HORVU.MOR | 5.70E-88 | 4.650259624 | 0.491 | 0.113 | 7.12E-84 | 1 | Endodermis | HORVU.MOR | low | protein_codir | --* | Dual-specificit | GO:0003824, | NA | NA |
| 130 | HORVU.MOR | HORVU.MOR | 1.89E-89 | 5.639289691 | 0.5 | 0.133 | 2.36E-85 | 1 | Endodermis | HORVU.MOR | low | non_coding | --* | 60S ribosomal | GO:0003735, | NA | NA |
| 142 | HORVU.MOR | HORVU.MOR | 4.44E-61 | 5.467206392 | 0.46 | 0.095 | 5.55E-57 | 1 | Endodermis | HORVU.MOR | low | protein_codir | --* | Transcription fa | NA | NA | NA |
| 156 | HORVU.MOR | HORVU.MOR | 6.52E-139 | 5.366435082 | 0.517 | 0.106 | 8.14E-135 | 1 | Endodermis | HORVU.MOR | high | protein_codir | *** | Pectin acetyles | NA | PF03283 | IPR004963,IPR029058 |
| 199 | HORVU.MOR | HORVU.MOR | 4.96E-58 | 4.684857052 | 0.509 | 0.104 | 6.19E-54 | 1 | Endodermis | HORVU.MOR | high | protein_codir | *** | Calmodulin-bi | GO:0005516, | PF07887 | IPR012416 |
| 206 | HORVU.MOR | HORVU.MOR | 1.28E-87 | 5.500088824 | 0.362 | 0.065 | 1.60E-83 | 1 | Endodermis | HORVU.MOR | high | protein_codir | --* | alpha/beta-Hy | NA | NA | NA |
| 64 | HORVU.MOR | HORVU.MOR | 1.39E-217 | 7.940785401 | 0.49 | 0.042 | 1.73E-213 | 2 | QC/Niche | HORVU.MOR | high | protein_codir | *** | Collagen, type | NA | IPR001480 | |
| 67 | HORVU.MOR | HORVU.MOR | 4.11E-100 | 8.084434936 | 0.723 | 0.347 | 5.13E-96 | 2 | QC/Niche | HORVU.MOR | high | protein_codir | *** | IgA FC recepto | NA | NA | NA |
| 81 | HORVU.MOR | HORVU.MOR | 2.66E-42 | 7.97826433 | 0.46 | 0.179 | 3.33E-38 | 2 | QC/Niche | HORVU.MOR | high | protein_codir | *** | CASP-like prot | GO:0005886, | PF04535 | IPR006459,IPR006702 |
| 82 | HORVU.MOR | HORVU.MOR | 1.14E-67 | 8.78216492 | 0.622 | 0.227 | 1.43E-63 | 2 | QC/Niche | HORVU.MOR | high | protein_codir | *** | Leguminosin g | NA | NA | NA |
| 116 | HORVU.MOR | HORVU.MOR | 1.26E-116 | 7.993964059 | 0.44 | 0.085 | 1.57E-112 | 2 | QC/Niche | HORVU.MOR | high | protein_codir | *** | GDSL esterase | GO:0016788 | PF00657 | IPR001087,IPR013830,IPR013830,IPR013830,IPR013830 |
| 125 | HORVU.MOR | HORVU.MOR | 4.37E-31 | 8.038618534 | 0.434 | 0.187 | 5.45E-27 | 2 | QC/Niche | HORVU.MOR | high | protein_codir | *** | Alpha/beta fol | GO:0003824, | PF00561 | IPR029058,IPR000073 |
| 190 | HORVU.MOR | HORVU.MOR | 6.88E-105 | 21.48014013 | 0.304 | 0.001 | 8.59E-101 | 2 | QC/Niche | HORVU.MOR | high | protein_codir | --* | OSBP(Oxyster | NA | NA | NA |
| 221 | HORVU.MOR | HORVU.MOR | 5.72E-227 | 8.33178244 | 0.336 | 0.003 | 7.14E-223 | 2 | QC/Niche | HORVU.MOR | high | protein_codir | *** | CASP-like prot | GO:0005886, | PF04535 | IPR006702,IPR006459 |
| 225 | HORVU.MOR | HORVU.MOR | 3.43E-34 | 7.946635276 | 0.516 | 0.33 | 4.28E-30 | 2 | QC/Niche | HORVU.MOR | high | protein_codir | *** | GDSL esterase | GO:0016788 | PF00657 | IPR001087,IPR013830,IPR013830,IPR013830 |
| 227 | HORVU.MOR | HORVU.MOR | 1.34E-14 | 8.233956903 | 0.295 | 0.077 | 1.67E-10 | 2 | QC/Niche | HORVU.MOR | high | protein_codir | *** | extracellular lig | NA | PF12056 | IPR021924 |
| 1 | HORVU.MOR | HORVU.MOR | 4.95E-175 | 5.332587648 | 0.598 | 0.102 | 6.18E-171 | 3 | unidentified | HORVU.MOR | low | protein_codir | *-* | Gamma-gliadir | GO:0045735 | PF13016 | IPR016140,IPR016140 |
| 42 | HORVU.MOR | HORVU.MOR | 8.52E-194 | 4.726136596 | 0.589 | 0.084 | 1.06E-189 | 3 | unidentified | HORVU.MOR | low | protein_codir | --* | cell wall inverta | NA | NA | NA |
| 72 | HORVU.MOR | HORVU.MOR | 6.94E-52 | 4.846460717 | 0.444 | 0.085 | 8.66E-48 | 3 | unidentified | HORVU.MOR | low | transposon_r | *-* | Transposon Ty3 | GO:0003676, | NA | NA |
| 99 | HORVU.MOR | HORVU.MOR | 1.90E-131 | 8.471199023 | 0.486 | 0.064 | 2.37E-127 | 3 | unidentified | HORVU.MOR | high | protein_codir | *** | Serine/threonir | GO:0000166, | PF00954,PF0 | IPR000858,IPR001480,IPR003609,IPR011009,IPR001480,IPR001245 |
| 118 | HORVU.MOR | HORVU.MOR | 1.71E-115 | 5.033880166 | 0.583 | 0.061 | 2.14E-111 | 3 | unidentified | HORVU.MOR | low | protein_codir | --* | 4-hydroxybenz | GO:0004659, | NA | NA |
| 146 | HORVU.MOR | HORVU.MOR | 7.31E-155 | 4.893162337 | 0.595 | 0.109 | 9.13E-151 | 3 | unidentified | HORVU.MOR | high | protein_codir | *** | Nuclear transc | GO:0005634, | PF00808 | IPR009072,IPR003958 |
| 157 | HORVU.MOR | HORVU.MOR | 1.56E-250 | 4.872072235 | 0.29 | 0.002 | 1.95E-246 | 3 | unidentified | HORVU.MOR | high | protein_codir | *** | AT hook motif I | GO:0003677, | PF03479 | IPR005175 |
| 180 | HORVU.MOR | HORVU.MOR | 1.99E-104 | 5.006855733 | 0.58 | 0.08 | 2.48E-100 | 3 | unidentified | HORVU.MOR | low | protein_codir | *-* | LAG1 longevity | GO:0016020, | PF03798 | IPR006634 |
| 197 | HORVU.MOR | HORVU.MOR | 3.28E-116 | 5.049267152 | 0.565 | 0.081 | 4.09E-112 | 3 | unidentified | HORVU.MOR | low | protein_codir | --* | 5'-AMP-activat | GO:0000166, | NA | NA |
| 226 | HORVU.MOR | HORVU.MOR | 1.12E-61 | 4.601941003 | 0.556 | 0.213 | 1.40E-57 | 3 | unidentified | HORVU.MOR | high | protein_codir | *** | D-ribose-bindi | NA | PF14009 | IPR025322 |
| 9 | HORVU.MOR | HORVU.MOR | 4.08E-26 | 5.050589984 | 0.276 | 0.094 | 5.09E-22 | 4 | unidentified | HORVU.MOR | low | protein_codir | --* | HAT dimerizati | GO:0003677, | NA | NA |
| 16 | HORVU.MOR | HORVU.MOR | 7.05E-64 | 4.468341882 | 0.29 | 0.051 | 8.80E-60 | 4 | unidentified | HORVU.MOR | high | protein_codir | *-* | Harpin inducin | NA | NA | NA |
| 24 | HORVU.MOR | HORVU.MOR | 6.19E-49 | 4.519655263 | 0.276 | 0.059 | 7.73E-45 | 4 | unidentified | HORVU.MOR | low | plastid_relate | --* | Omega-6 fatty | GO:0006629, | NA | NA |
| 25 | HORVU.MOR | HORVU.MOR | 8.16E-42 | 4.702273466 | 0.29 | 0.075 | 1.02E-37 | 4 | unidentified | HORVU.MOR | low | protein_codir | *-* | BED zinc finger | GO:0003677, | PF02892 | IPR003656,IPR012337 |
| 29 | HORVU.MOR | HORVU.MOR | 3.52E-26 | 6.222091601 | 0.287 | 0.095 | 4.40E-22 | 4 | unidentified | HORVU.MOR | high | protein_codir | *** | LINE-1 reverse | NA | PF13966,PF0 | IPR026960,IPR012337,IPR000477 |
| 47 | HORVU.MOR | HORVU.MOR | 2.75E-11 | 4.551383034 | 0.287 | 0.141 | 3.44E-07 | 4 | unidentified | HORVU.MOR | low | protein_codir | --* | ADP-L-glycero- | GO:0003824, | NA | NA |
| 50 | HORVU.MOR | HORVU.MOR | 2.39E-19 | 4.450591482 | 0.283 | 0.084 | 2.98E-15 | 4 | unidentified | HORVU.MOR | low | protein_codir | --* | cytokinin oxida | NA | NA | NA |
| 166 | HORVU.MOR | HORVU.MOR | 1.30E-20 | 4.573187518 | 0.287 | 0.11 | 1.62E-16 | 4 | unidentified | HORVU.MOR | high | protein_codir | *** | Vicilin-like anti | GO:0045735 | PF00190,PF0 | IPR011051,IPR011051,IPR006045,IPR006045 |
| 204 | HORVU.MOR | HORVU.MOR | 0.00584287 | 4.519457193 | 0.287 | 0.165 | 1 | 4 | unidentified | HORVU.MOR | high | protein_codir | *** | Beta-glucan bi | GO:0052861 | PF03639 | IPR005200 |
| 230 | HORVU.MOR | HORVU.MOR | 4.43E-19 | 4.490423524 | 0.297 | 0.131 | 5.53E-15 | 4 | unidentified | HORVU.MOR | low | transposon_r | *-* | Transposon pro | GO:0003677, | PF03108 | IPR004332 |
| 10 | HORVU.MOR | HORVU.MOR | 2.07E-172 | 5.905861257 | 0.544 | 0.079 | 2.59E-168 | 5 | Vasculature | HORVU.MOR | low | protein_codir | --* | Ribosome mat | GO:0005737, | NA | NA |
| 48 | HORVU.MOR | HORVU.MOR | 5.58E-102 | 5.757942051 | 0.445 | 0.063 | 6.97E-98 | 5 | Vasculature | HORVU.MOR | high | protein_codir | --* | Transmembrar | GO:0016021 | NA | NA |
| 87 | HORVU.MOR | HORVU.MOR | 1.93E-22 | 6.025898385 | 0.406 | 0.135 | 2.41E-18 | 5 | Vasculature | HORVU.MOR | high | protein_codir | *** | transmembran | NA | PF03140 | IPR004158 |
| 147 | HORVU.MOR | HORVU.MOR | 1.09E-42 | 5.899627609 | 0.534 | 0.155 | 1.36E-38 | 5 | Vasculature | HORVU.MOR | high | protein_codir | *** | F-box family pr | NA | PF12937 | IPR001810,IPR001810 |
| 149 | HORVU.MOR | HORVU.MOR | 1.16E-184 | 5.654429806 | 0.618 | 0.081 | 1.45E-180 | 5 | Vasculature | HORVU.MOR | low | protein_codir | *-* | Protein IQ-DOI | NA | PF13178 | IPR025064 |
| 167 | HORVU.MOR | HORVU.MOR | 1.46E-127 | 6.477354465 | 0.565 | 0.042 | 1.82E-123 | 5 | Vasculature | HORVU.MOR | high | protein_codir | *-* | Eukaryotic tran | GO:0001732, | NA | NA |
| 171 | HORVU.MOR | HORVU.MOR | 1.60E-144 | 5.92501096 | 0.541 | 0.07 | 1.99E-140 | 5 | Vasculature | HORVU.MOR | low | protein_codir | --* | GRAM-contain | NA | NA | NA |
| 186 | HORVU.MOR | HORVU.MOR | 1.93E-100 | 5.658537715 | 0.558 | 0.049 | 2.41E-96 | 5 | Vasculature | HORVU.MOR | low | protein_codir | *-* | Dynamin-like | GO:0000166, | PF00350 | IPR027417,IPR022812 |
| 210 | HORVU.MOR | HORVU.MOR | 4.41E-163 | 9.193972903 | 0.42 | 0.034 | 5.51E-159 | 5 | Vasculature | HORVU.MOR | low | plastid_relate | --* | Protein PAM68 | GO:0016020, | PF11947 | IPR021855 |
| 223 | HORVU.MOR | HORVU.MOR | 1.37E-184 | 6.001015753 | 0.565 | 0.079 | 1.71E-180 | 5 | Vasculature | HORVU.MOR | high | protein_codir | --* | VQ motif-conta | NA | PF05678 | IPR008889 |

| | | | | | | | | | | | | | | | | | |
|---|---|---|---|---|---|---|---|---|---|---|---|---|---|---|---|---|---|
| 3 | HORVU.MOR | HORVU.MOR | 4.11E-35 | 7.463581169 | 0.647 | 0.446 | 5.13E-31 | 6 | Cell cycle | HORVU.MOR | high | protein_codir | *** | Late embryoge | NA | PF06830 | IPR009646 |
| 21 | HORVU.MOR | HORVU.MOR | 4.30E-194 | 6.649144633 | 0.519 | 0.051 | 5.36E-190 | 6 | Cell cycle | HORVU.MOR | low | protein_codir | --* | Zinc finger prot | GO:0003676, | NA | NA |
| 41 | HORVU.MOR | HORVU.MOR | 4.07E-115 | 6.299151608 | 0.474 | 0.059 | 5.08E-111 | 6 | Cell cycle | HORVU.MOR | low | protein_codir | --* | DNA topoisom | NA | NA | NA |
| 52 | HORVU.MOR | HORVU.MOR | 1.21E-142 | 6.123251506 | 0.564 | 0.074 | 1.51E-138 | 6 | Cell cycle | HORVU.MOR | low | protein_codir | --* | Galactose oxid | NA | NA | NA |
| 88 | HORVU.MOR | HORVU.MOR | 0.00442024 | 9.41522598 | 0.504 | 0.31 | 1 | 6 | Cell cycle | HORVU.MOR | high | protein_codir | *** | Late embryoge | NA | PF06830 | IPR009646 |
| 91 | HORVU.MOR | HORVU.MOR | 8.09E-05 | 6.635980961 | 0.541 | 0.346 | 1 | 6 | Cell cycle | HORVU.MOR | high | protein_codir | *** | Expansin prote | GO:0005576, | PF01357,PF0 | IPR009009,IPR007117,IPR007117,IPR009009 |
| 98 | HORVU.MOR | HORVU.MOR | 3.25E-57 | 5.979711507 | 0.365 | 0.083 | 4.05E-53 | 6 | Cell cycle | HORVU.MOR | high | protein_codir | *-* | Micronuclear li | GO:0016020, | NA | NA |
| 151 | HORVU.MOR | HORVU.MOR | 1.32E-05 | 9.801693856 | 0.158 | 0.276 | 0.1647223 | 6 | Cell cycle | HORVU.MOR | high | protein_codir | *** | Late embryoge | NA | PF06830 | IPR009646 |
| 152 | HORVU.MOR | HORVU.MOR | 8.58E-18 | 11.04780228 | 0.053 | 0.302 | 1.07E-13 | 6 | Cell cycle | HORVU.MOR | high | protein_codir | *** | Late embryoge | NA | PF06830 | IPR009646 |
| 189 | HORVU.MOR | HORVU.MOR | 3.00E-162 | 5.787736663 | 0.583 | 0.071 | 3.74E-158 | 6 | Cell cycle | HORVU.MOR | high | protein_codir | --* | THAP domain-c | GO:0003676, | NA | NA |
| 20 | HORVU.MOR | HORVU.MOR | 2.35E-75 | 5.536417652 | 0.477 | 0.108 | 2.94E-71 | 7 | Cortex | HORVU.MOR | high | protein_codir | *** | Flavin-containi | GO:0004497, | PF00743 | IPR020946,IPR023753,IPR023753 |
| 28 | HORVU.MOR | HORVU.MOR | 3.69E-125 | 5.607535492 | 0.387 | 0.027 | 4.60E-121 | 7 | Cortex | HORVU.MOR | low | protein_codir | *-* | Ribosomal pro | GO:0005739, | NA | NA |
| 44 | HORVU.MOR | HORVU.MOR | 6.68E-26 | 5.666590306 | 0.256 | 0.066 | 8.34E-22 | 7 | Cortex | HORVU.MOR | high | protein_codir | *** | Benzyl alcohol | GO:0016740, | PF02458 | IPR003480 |
| 84 | HORVU.MOR | HORVU.MOR | 2.41E-80 | 5.103439542 | 0.331 | 0.044 | 3.01E-76 | 7 | Cortex | HORVU.MOR | low | protein_codir | --* | 2-hydroxymucc | GO:0008152, | NA | NA |
| 112 | HORVU.MOR | HORVU.MOR | 7.17E-21 | 7.145945623 | 0.271 | 0.088 | 8.94E-17 | 7 | Cortex | HORVU.MOR | high | protein_codir | *** | BTR2-like prote | NA | NA | NA |
| 120 | HORVU.MOR | HORVU.MOR | 9.63E-64 | 5.114329998 | 0.346 | 0.092 | 1.20E-59 | 7 | Cortex | HORVU.MOR | low | protein_codir | --* | Restriction of t | GO:0003674, | NA | NA |
| 129 | HORVU.MOR | HORVU.MOR | 1.96E-86 | 5.910188861 | 0.508 | 0.106 | 2.45E-82 | 7 | Cortex | HORVU.MOR | low | protein_codir | --* | R-spondin-2 | GO:0001649, | NA | NA |
| 133 | HORVU.MOR | HORVU.MOR | 1.21E-59 | 5.662870597 | 0.342 | 0.126 | 1.51E-55 | 7 | Cortex | HORVU.MOR | high | protein_codir | *** | Pectinesterase | GO:0004857, | PF04043 | IPR006501,IPR006501,IPR006501 |
| 135 | HORVU.MOR | HORVU.MOR | 2.76E-44 | 10.79753479 | 0.44 | 0.111 | 3.44E-40 | 7 | Cortex | HORVU.MOR | high | protein_codir | *** | Apoptosis-indu | GO:0016491, | PF07992 | IPR023753,IPR023753 |
| 161 | HORVU.MOR | HORVU.MOR | 2.22E-18 | 5.596584717 | 0.365 | 0.135 | 2.77E-14 | 7 | Cortex | HORVU.MOR | high | protein_codir | *** | Scarecrow trar | GO:0003700, | PF03514 | IPR005202 |
| 61 | HORVU.MOR | HORVU.MOR | 0.00872243 | 3.937630439 | 0.218 | 0.371 | 1 | 8 | unidentified | HORVU.MOR | high | protein_codir | *** | WRKY transcrip | GO:0003677, | PF03106 | IPR003657,IPR003657 |
| 79 | HORVU.MOR | HORVU.MOR | 3.40E-10 | 3.895585684 | 0.129 | 0.251 | 4.25E-06 | 8 | unidentified | HORVU.MOR | low | protein_codir | --* | Heme-degradi | GO:0004392, | NA | NA |
| 89 | HORVU.MOR | HORVU.MOR | 0.00123816 | 3.625994319 | 0.452 | 0.512 | 1 | 8 | unidentified | HORVU.MOR | high | protein_codir | *** | Xyloglucan enc | NA | PF06955,PF0 | IPR013320,IPR010713,IPR000757 |
| 95 | HORVU.MOR | HORVU.MOR | 1.13E-63 | 3.643286408 | 0.782 | 0.508 | 1.41E-59 | 8 | unidentified | HORVU.MOR | high | protein_codir | *** | Beta-fructofura | GO:0004553, | PF00251,PF0 | IPR013320,IPR013320,IPR023296,IPR013148,IPR013189 |
| 108 | HORVU.MOR | HORVU.MOR | 1.42E-21 | 3.799676052 | 0.605 | 0.566 | 1.78E-17 | 8 | unidentified | HORVU.MOR | high | protein_codir | *** | GDSL esterase | GO:0016788 | PF00657 | IPR001087,IPR013830,IPR013830,IPR013830,IPR013830 |
| 110 | HORVU.MOR | HORVU.MOR | 9.24E-36 | 3.837051007 | 0.669 | 0.544 | 1.15E-31 | 8 | unidentified | HORVU.MOR | high | protein_codir | --* | GMP synthase | GO:0000166, | PF07722 | IPR029062,IPR029062,IPR011697 |
| 145 | HORVU.MOR | HORVU.MOR | 5.61E-08 | 4.834042389 | 0.444 | 0.443 | 0.00069988 | 8 | unidentified | HORVU.MOR | high | protein_codir | *** | 3'-N-debenzoy | GO:0016740, | PF02458 | IPR003480 |
| 217 | HORVU.MOR | HORVU.MOR | 0.00134043 | 3.715137981 | 0.21 | 0.397 | 1 | 8 | unidentified | HORVU.MOR | high | protein_codir | *-* | Laccase | GO:0005507, | PF07732,PF0 | IPR008972,IPR011707,IPR008972,IPR008972,IPR011706,IPR001117 |
| 229 | HORVU.MOR | HORVU.MOR | 0.00018167 | 3.903517341 | 0.181 | 0.274 | 1 | 8 | unidentified | HORVU.MOR | low | non_coding | *** | Cytochrome P4 | GO:0004497, | PF00067 | IPR001128,IPR001128 |
| 233 | HORVU.MOR | HORVU.MOR | 2.89E-05 | 3.76141838 | 0.181 | 0.336 | 0.36066656 | 8 | unidentified | HORVU.MOR | high | protein_codir | --* | DNA topoisom | GO:0003676, | PF06839 | IPR010666 |
| 18 | HORVU.MOR | HORVU.MOR | 2.37E-106 | 5.294906924 | 0.416 | 0.054 | 2.96E-102 | 9 | Cortex | HORVU.MOR | low | protein_codir | *-* | Pyruvate dehyc | GO:0003824, | PF01693,PF0 | IPR011320,IPR009027,IPR029061,IPR005475 |
| 119 | HORVU.MOR | HORVU.MOR | 1.78E-67 | 5.186450644 | 0.521 | 0.19 | 2.22E-63 | 9 | Cortex | HORVU.MOR | low | protein_codir | *-* | zinc finger MYN | NA | PF14291 | IPR025398 |
| 122 | HORVU.MOR | HORVU.MOR | 3.71E-90 | 5.305200117 | 0.29 | 0.014 | 4.64E-86 | 9 | Cortex | HORVU.MOR | low | transposon_r | *-* | Transposon pro | GO:0008270, | PF04434 | IPR007527 |
| 126 | HORVU.MOR | HORVU.MOR | 3.83E-153 | 5.435052835 | 0.315 | 0.041 | 4.78E-149 | 9 | Cortex | HORVU.MOR | low | protein_codir | *-* | Serine/threonir | NA | PF10536 | IPR019557 |
| 137 | HORVU.MOR | HORVU.MOR | 2.20E-11 | 6.163015917 | 0.311 | 0.104 | 2.74E-07 | 9 | Cortex | HORVU.MOR | low | protein_codir | *-* | Retrovirus-rela | NA | NA | NA |
| 160 | HORVU.MOR | HORVU.MOR | 1.04E-36 | 5.379466366 | 0.513 | 0.129 | 1.30E-32 | 9 | Cortex | HORVU.MOR | high | protein_codir | *** | Scarecrow trar | GO:0003700, | PF03514 | IPR005202 |
| 174 | HORVU.MOR | HORVU.MOR | 6.63E-39 | 6.054808266 | 0.504 | 0.099 | 8.28E-35 | 9 | Cortex | HORVU.MOR | low | non_coding | --* | Mediator of RN | GO:0001104, | NA | NA |
| 208 | HORVU.MOR | HORVU.MOR | 1.51E-71 | 5.790203898 | 0.576 | 0.145 | 1.89E-67 | 9 | Cortex | HORVU.MOR | high | protein_codir | *** | YABBY protein | GO:0007275 | PF04690 | IPR009071,IPR006780 |
| 215 | HORVU.MOR | HORVU.MOR | 1.14E-122 | 6.695683166 | 0.315 | 0.007 | 1.42E-118 | 9 | Cortex | HORVU.MOR | low | non_coding | *** | Cytochrome P4 | GO:0005506, | PF00067 | IPR001128,IPR001128 |
| 228 | HORVU.MOR | HORVU.MOR | 1.49E-71 | 5.314107057 | 0.45 | 0.122 | 1.87E-67 | 9 | Cortex | HORVU.MOR | high | protein_codir | *-* | Zinc finger MYN | NA | NA | NA |
| 38 | HORVU.MOR | HORVU.MOR | 1.63E-79 | 10.84355268 | 0.549 | 0.052 | 2.04E-75 | 10 | Metaxlem | HORVU.MOR | high | protein_codir | *** | Mitochondrial t | GO:0003690, | PF02536,PF0 | IPR003690,IPR003690 |
| 43 | HORVU.MOR | HORVU.MOR | 2.36E-62 | 11.38198642 | 0.319 | 0.065 | 2.95E-58 | 10 | Metaxlem | HORVU.MOR | high | protein_codir | *** | Benzyl alcohol | GO:0016740, | PF02458 | IPR003480 |
| 97 | HORVU.MOR | HORVU.MOR | 1.42E-101 | 6.384974736 | 0.712 | 0.165 | 1.78E-97 | 10 | Metaxlem | HORVU.MOR | low | protein_codir | --* | basic helix-loo | NA | NA | NA |
| 141 | HORVU.MOR | HORVU.MOR | 5.58E-126 | 6.61456724 | 0.544 | 0.078 | 6.97E-122 | 10 | Metaxlem | HORVU.MOR | high | protein_codir | *** | Amino acid per | GO:0016020, | PF01490 | IPR013057 |
| 153 | HORVU.MOR | HORVU.MOR | 1.69E-115 | 6.508939514 | 0.469 | 0.085 | 2.11E-111 | 10 | Metaxlem | HORVU.MOR | high | protein_codir | *** | NBS-LRR-like r | GO:0043531 | PF00931 | IPR002182,IPR027417,IPR032675 |
| 168 | HORVU.MOR | HORVU.MOR | 2.57E-67 | 6.074880071 | 0.376 | 0.038 | 3.20E-63 | 10 | Metaxlem | HORVU.MOR | high | protein_codir | *** | Receptor-like p | GO:0004672, | PF12819,PF0 | IPR011009,IPR024788,IPR001245 |
| 179 | HORVU.MOR | HORVU.MOR | 2.12E-136 | 6.382754249 | 0.54 | 0.097 | 2.65E-132 | 10 | Metaxlem | HORVU.MOR | high | protein_codir | *** | Receptor kinas | GO:0000166, | PF08263,PF0 | IPR011009,IPR013210,IPR001611,IPR001611,IPR001611,IPR000719 |
| 185 | HORVU.MOR | HORVU.MOR | 1.13E-56 | 8.021450948 | 0.593 | 0.062 | 1.41E-52 | 10 | Metaxlem | HORVU.MOR | high | protein_codir | *** | Cytokinin ribos | GO:0009691, | PF03641 | IPR031100,IPR005269 |
| 224 | HORVU.MOR | HORVU.MOR | 9.74E-133 | 6.854979885 | 0.942 | 0.32 | 1.22E-128 | 10 | Metaxlem | HORVU.MOR | high | protein_codir | *** | Pathogenesis-r | GO:0006952, | PF00407 | IPR000916 |
| 240 | HORVU.MOR | HORVU.MOR | 3.76E-185 | 6.00867291 | 0.619 | 0.061 | 4.69E-181 | 10 | Metaxlem | HORVU.MOR | high | protein_codir | *** | SPX domain-cc | GO:0016036, | PF03105 | IPR004331 |
| 46 | HORVU.MOR | HORVU.MOR | 4.53E-52 | 7.492096629 | 0.536 | 0.144 | 5.65E-48 | 11 | Cell cycle | HORVU.MOR | high | protein_codir | *-* | Chitinase | GO:0004568, | PF00187,PF0 | IPR001002,IPR001002,IPR001002,IPR001002,IPR001002,IPR001002,IPR001002,IPR |
| 60 | HORVU.MOR | HORVU.MOR | 3.60E-166 | 10.2798446 | 0.377 | 0.045 | 4.49E-162 | 11 | Cell cycle | HORVU.MOR | high | protein_codir | -** | Glycine-rich ce | NA | NA | NA |
| 78 | HORVU.MOR | HORVU.MOR | 1.80E-58 | 8.368722271 | 0.609 | 0.105 | 2.25E-54 | 11 | Cell cycle | HORVU.MOR | high | protein_codir | *** | Serine incorpo | GO:0016020, | PF03348 | IPR005016 |
| 92 | HORVU.MOR | HORVU.MOR | 1.84E-180 | 7.113163676 | 0.264 | 0.006 | 2.30E-176 | 11 | Cell cycle | HORVU.MOR | high | protein_codir | *** | B3 domain-cor | GO:0003677, | PF02362,PF0 | IPR015300,IPR003340,IPR003340,IPR015300 |
| 102 | HORVU.MOR | HORVU.MOR | 1.47E-111 | 7.131983814 | 0.482 | 0.037 | 1.83E-107 | 11 | Cell cycle | HORVU.MOR | high | protein_codir | *** | Peroxidase | GO:0004601, | PF00141 | IPR010255,IPR002016 |
| 105 | HORVU.MOR | HORVU.MOR | 8.67E-239 | 7.229903612 | 0.555 | 0.043 | 1.08E-234 | 11 | Cell cycle | HORVU.MOR | high | protein_codir | --* | Auxin-binding p | GO:0005783, | NA | NA |
| 106 | HORVU.MOR | HORVU.MOR | 3.70E-241 | 6.470475605 | 0.668 | 0.061 | 4.62E-237 | 11 | Cell cycle | HORVU.MOR | high | protein_codir | --* | Transcription e | GO:0000398, | NA | NA |
| 107 | HORVU.MOR | HORVU.MOR | 8.19E-33 | 7.112565656 | 0.495 | 0.145 | 1.02E-28 | 11 | Cell cycle | HORVU.MOR | high | protein_codir | --* | Trypsin inhibitc | GO:0004867, | PF00228,PF0 | IPR000877,IPR000877,IPR000877,IPR000877,IPR000877,IPR000877 |
| 172 | HORVU.MOR | HORVU.MOR | 1.14E-264 | 6.875958421 | 0.591 | 0.03 | 1.42E-260 | 11 | Cell cycle | HORVU.MOR | high | protein_codir | *** | Glycosyltransfe | GO:0016020, | PF04577 | IPR007657 |
| 239 | HORVU.MOR | HORVU.MOR | 1.85E-204 | 6.339852551 | 0.736 | 0.097 | 2.31E-200 | 11 | Cell cycle | HORVU.MOR | high | protein_codir | *** | Cellulose synth | GO:0016020, | PF03552,PF0 | IPR029044,IPR029044,IPR029044,IPR005150,IPR005150 |
| 14 | HORVU.MOR | HORVU.MOR | 1.19E-248 | 4.983742992 | 0.441 | 0.016 | 1.49E-244 | 12 | Cortex | HORVU.MOR | high | protein_codir | --* | Reverse transc | GO:0003964, | NA | NA |
| 15 | HORVU.MOR | HORVU.MOR | 1.57E-94 | 5.379374042 | 0.592 | 0.115 | 1.95E-90 | 12 | Cortex | HORVU.MOR | high | protein_codir | *** | Endoglucanase | GO:0000272, | PF00759 | IPR008928,IPR001701 |

| | | | | | | | | | | | | | | | | | |
|---|---|---|---|---|---|---|---|---|---|---|---|---|---|---|---|---|---|
| 94 | HORVU.MOR | HORVU.MOR | 5.02E-120 | 5.767791099 | 0.545 | 0.076 | 6.27E-116 | 12 | Cortex | HORVU.MOR | low | protein_codir | --* | Orotidine 5'-ph | GO:0003824, | NA | NA |
| 101 | HORVU.MOR | HORVU.MOR | 2.11E-57 | 5.195500401 | 0.338 | 0.027 | 2.63E-53 | 12 | Cortex | HORVU.MOR | high | protein_codir | *** | Acetyl-coenzyr | GO:0000166, | PF13193,PF0 | IPR025110,IPR000873 |
| 132 | HORVU.MOR | HORVU.MOR | 6.09E-93 | 6.805810102 | 0.432 | 0.077 | 7.60E-89 | 12 | Cortex | HORVU.MOR | high | protein_codir | *** | Leucine-rich re | GO:0000166, | PF13855,PF1 | IPR001611,IPR001611,IPR013210,IPR032675 |
| 134 | HORVU.MOR | HORVU.MOR | 7.49E-32 | 5.946163571 | 0.441 | 0.114 | 9.35E-28 | 12 | Cortex | HORVU.MOR | high | protein_codir | *** | Apoptosis-indu | GO:0016491, | PF07992 | IPR023753,IPR023753 |
| 143 | HORVU.MOR | HORVU.MOR | 1.51E-50 | 6.374721071 | 0.272 | 0.114 | 1.89E-46 | 12 | Cortex | HORVU.MOR | low | protein_codir | --* | Transcription fa | NA | NA | NA |
| 192 | HORVU.MOR | HORVU.MOR | 1.40E-77 | 5.117713377 | 0.634 | 0.125 | 1.75E-73 | 12 | Cortex | HORVU.MOR | low | protein_codir | --* | Homeobox-leu | NA | NA | NA |
| 222 | HORVU.MOR | HORVU.MOR | 5.92E-13 | 5.10070794 | 0.286 | 0.11 | 7.39E-09 | 12 | Cortex | HORVU.MOR | low | transposon_r | *-* | Retrovirus-rela | GO:0003676, | NA | NA |
| 231 | HORVU.MOR | HORVU.MOR | 1.65E-80 | 7.677331419 | 0.512 | 0.08 | 2.06E-76 | 12 | Cortex | HORVU.MOR | low | protein_codir | --* | Glutathione S-t | GO:0016740 | NA | NA |
| 5 | HORVU.MOR | HORVU.MOR | 1.19E-20 | 3.493839862 | 0.63 | 0.333 | 1.48E-16 | 13 | Cortex | HORVU.MOR | high | protein_codir | *** | Chymotrypsin i | GO:0004867, | PF00280 | IPR000864,IPR000864 |
| 8 | HORVU.MOR | HORVU.MOR | 1.78E-05 | 3.428262819 | 0.481 | 0.281 | 0.22264072 | 13 | Cortex | HORVU.MOR | high | protein_codir | *** | Chymotrypsin i | GO:0004867, | PF00280 | IPR000864,IPR000864 |
| 19 | HORVU.MOR | HORVU.MOR | 7.27E-38 | 3.607274576 | 0.741 | 0.539 | 9.08E-34 | 13 | Cortex | HORVU.MOR | high | protein_codir | *** | Lactoylglutathi | GO:0016829 | PF00903 | IPR029068,IPR004360 |
| 53 | HORVU.MOR | HORVU.MOR | 0.00576789 | 3.420390724 | 0.196 | 0.331 | 1 | 13 | Cortex | HORVU.MOR | low | protein_codir | --* | SET domain-co | NA | NA | NA |
| 68 | HORVU.MOR | HORVU.MOR | 0.00713363 | 3.484623694 | 0.201 | 0.298 | 1 | 13 | Cortex | HORVU.MOR | low | protein_codir | --* | formin 3 | NA | NA | NA |
| 74 | HORVU.MOR | HORVU.MOR | 3.37E-05 | 4.86973467 | 0.18 | 0.302 | 0.42061813 | 13 | Cortex | HORVU.MOR | high | protein_codir | *** | Peroxidase | GO:0004601, | PF00141 | IPR010255,IPR002016 |
| 140 | HORVU.MOR | HORVU.MOR | 3.62E-05 | 3.612435818 | 0.122 | 0.25 | 0.45183863 | 13 | Cortex | HORVU.MOR | low | protein_codir | --* | 2-oxoglutarate | NA | NA | NA |
| 175 | HORVU.MOR | HORVU.MOR | 3.30E-07 | 3.407333404 | 0.132 | 0.302 | 0.00411538 | 13 | Cortex | HORVU.MOR | low | protein_codir | --* | Meiosis arrest | GO:0005777, | NA | NA |
| 200 | HORVU.MOR | HORVU.MOR | 7.04E-07 | 4.418613049 | 0.058 | 0.293 | 0.00879417 | 13 | Cortex | HORVU.MOR | high | protein_codir | *-* | Glutathione S-t | GO:0016740 | NA | NA |
| 207 | HORVU.MOR | HORVU.MOR | 0.00659878 | 3.78135214 | 0.196 | 0.27 | 1 | 13 | Cortex | HORVU.MOR | low | protein_codir | --* | Mortality factor | GO:0000724, | NA | NA |
| 11 | HORVU.MOR | HORVU.MOR | 3.07E-44 | 6.914105605 | 0.311 | 0.048 | 3.83E-40 | 14 | Cell cycle | HORVU.MOR | low | protein_codir | --* | Potassium volt | GO:0005216, | NA | NA |
| 23 | HORVU.MOR | HORVU.MOR | 7.80E-133 | 6.018401829 | 0.578 | 0.055 | 9.74E-129 | 14 | Cell cycle | HORVU.MOR | low | protein_codir | *-* | Peroxidase | GO:0004601, | PF00141 | IPR002016,IPR010255 |
| 37 | HORVU.MOR | HORVU.MOR | 8.91E-99 | 5.675013779 | 0.567 | 0.091 | 1.11E-94 | 14 | Cell cycle | HORVU.MOR | low | transposon_r | *-* | Retrotranspos | GO:0003676, | PF00098 | IPR001878,IPR001878 |
| 115 | HORVU.MOR | HORVU.MOR | 9.22E-20 | 6.012839071 | 0.367 | 0.05 | 1.15E-15 | 14 | Cell cycle | HORVU.MOR | low | protein_codir | --* | VQ motif-conta | NA | NA | NA |
| 121 | HORVU.MOR | HORVU.MOR | 2.37E-125 | 5.85317977 | 0.561 | 0.065 | 2.96E-121 | 14 | Cell cycle | HORVU.MOR | low | protein_codir | --* | Elongation fact | GO:0000166, | NA | NA |
| 165 | HORVU.MOR | HORVU.MOR | 1.84E-82 | 5.960211757 | 0.4 | 0.065 | 2.30E-78 | 14 | Cell cycle | HORVU.MOR | high | protein_codir | *-* | Zinc finger fam | GO:0003676, | PF13912,PF1 | IPR007087,IPR007087 |
| 176 | HORVU.MOR | HORVU.MOR | 1.76E-115 | 6.06805306 | 0.506 | 0.04 | 2.19E-111 | 14 | Cell cycle | HORVU.MOR | low | protein_codir | --* | villin 3 | NA | NA | NA |
| 177 | HORVU.MOR | HORVU.MOR | 2.33E-71 | 5.502335625 | 0.367 | 0.024 | 2.90E-67 | 14 | Cell cycle | HORVU.MOR | low | transposon_r | *-* | Retrotranspos | NA | IPR005135 | |
| 191 | HORVU.MOR | HORVU.MOR | 1.59E-78 | 5.966570029 | 0.594 | 0.083 | 1.99E-74 | 14 | Cell cycle | HORVU.MOR | high | protein_codir | --* | RNA-binding (F | NA | PF11321 | IPR021470 |
| 237 | HORVU.MOR | HORVU.MOR | 1.53E-61 | 6.241423225 | 0.267 | 0.027 | 1.91E-57 | 14 | Cell cycle | HORVU.MOR | high | protein_codir | *-* | Chitinase | GO:0004568, | PF00187,PF0 | IPR001002,IPR001002,IPR001002,IPR001002,IPR001002,IPR001002,IPR001002,IPR |
| 27 | HORVU.MOR | HORVU.MOR | 1.25E-08 | 3.515912083 | 0.101 | 0.272 | 0.00015663 | 15 | unidentified | HORVU.MOR | high | protein_codir | *** | LIGHT-DEPENI | NA | PF04852 | IPR006936 |
| 55 | HORVU.MOR | HORVU.MOR | 1.01E-05 | 3.600128845 | 0.107 | 0.267 | 0.12587785 | 15 | unidentified | HORVU.MOR | high | protein_codir | *** | Disease resista | NA | IPR027417 | |
| 63 | HORVU.MOR | HORVU.MOR | 2.14E-08 | 4.071924643 | 0.112 | 0.319 | 0.00026673 | 15 | unidentified | HORVU.MOR | high | protein_codir | --* | F-box family pr | NA | PF00646 | IPR001810,IPR001810 |
| 70 | HORVU.MOR | HORVU.MOR | 3.24E-05 | 3.813189116 | 0.112 | 0.276 | 0.40497813 | 15 | unidentified | HORVU.MOR | low | transposon_r | *-* | Retrovirus-rela | GO:0003676, | NA | NA |
| 73 | HORVU.MOR | HORVU.MOR | 1.97E-13 | 3.84988292 | 0.065 | 0.305 | 2.46E-09 | 15 | unidentified | HORVU.MOR | high | protein_codir | *** | Peroxidase | GO:0004601, | PF00141 | IPR010255,IPR002016 |
| 80 | HORVU.MOR | HORVU.MOR | 3.04E-09 | 4.018822175 | 0.101 | 0.25 | 3.79E-05 | 15 | unidentified | HORVU.MOR | low | protein_codir | --* | Heme-degradi | GO:0004392, | NA | NA |
| 85 | HORVU.MOR | HORVU.MOR | 2.14E-06 | 4.806185352 | 0.107 | 0.255 | 0.02665368 | 15 | unidentified | HORVU.MOR | high | protein_codir | --* | Cortactin-bind | GO:0005737, | NA | NA |
| 90 | HORVU.MOR | HORVU.MOR | 8.87E-10 | 4.50368792 | 0.083 | 0.277 | 1.11E-05 | 15 | unidentified | HORVU.MOR | low | protein_codir | --* | dynamin relate | NA | NA | NA |
| 150 | HORVU.MOR | HORVU.MOR | 8.64E-08 | 3.499045149 | 0.083 | 0.26 | 0.00107902 | 15 | unidentified | HORVU.MOR | high | protein_codir | *** | SKP1-like prote | GO:0006511 | PF03931,PF0 | IPR016072,IPR011333,IPR016073,IPR016072 |
| 182 | HORVU.MOR | HORVU.MOR | 2.59E-05 | 4.054201304 | 0.118 | 0.25 | 0.3234027 | 15 | unidentified | HORVU.MOR | high | protein_codir | *** | Pre-mRNA-spli | GO:0006396 | IPR011990,IPR011990,IPR011990 | |
| 6 | HORVU.MOR | HORVU.MOR | 1.68E-32 | 5.619981459 | 0.256 | 0.042 | 2.09E-28 | 16 | unidentified | HORVU.MOR | low | protein_codir | --* | tRNA (uracil(54 | GO:0003723, | NA | NA |
| 39 | HORVU.MOR | HORVU.MOR | 4.90E-06 | 5.538789653 | 0.25 | 0.14 | 0.06118554 | 16 | unidentified | HORVU.MOR | low | protein_codir | --* | Oligopeptide-b | GO:0005886, | NA | NA |
| 58 | HORVU.MOR | HORVU.MOR | 0.00010085 | 5.691397207 | 0.341 | 0.273 | 1 | 16 | unidentified | HORVU.MOR | low | non_coding | *** | Cytochrome P4 | GO:0004497, | NA | NA |
| 83 | HORVU.MOR | HORVU.MOR | 1.10E-09 | 6.657957453 | 0.28 | 0.068 | 1.38E-05 | 16 | unidentified | HORVU.MOR | low | non_coding | *** | Cytochrome P4 | GO:0004497, | PF00067 | IPR001128,IPR001128 |
| 100 | HORVU.MOR | HORVU.MOR | 5.46E-10 | 5.566710685 | 0.268 | 0.081 | 6.82E-06 | 16 | unidentified | HORVU.MOR | high | protein_codir | *-* | Exocyst compl | GO:0000145, | PF03081 | IPR004140,IPR016159 |
| 113 | HORVU.MOR | HORVU.MOR | 1.58E-10 | 5.605196374 | 0.25 | 0.093 | 1.97E-06 | 16 | unidentified | HORVU.MOR | high | protein_codir | *** | BTR2-like prote | NA | NA | NA |
| 209 | HORVU.MOR | HORVU.MOR | 3.82E-10 | 5.462218586 | 0.293 | 0.151 | 4.77E-06 | 16 | unidentified | HORVU.MOR | high | protein_codir | *** | Peptide transp | GO:0005215, | PF00854 | IPR000109,IPR020846,IPR020846,IPR020846 |
| 212 | HORVU.MOR | HORVU.MOR | 4.07E-17 | 5.9144147 | 0.317 | 0.137 | 5.08E-13 | 16 | unidentified | HORVU.MOR | high | protein_codir | *-* | Receptor-kinas | GO:0000166, | PF08263,PF0 | IPR013210,IPR032675,IPR000719,IPR001611,IPR011009,IPR001611,IPR032675 |
| 234 | HORVU.MOR | HORVU.MOR | 1.19E-21 | 5.536580608 | 0.268 | 0.068 | 1.49E-17 | 16 | unidentified | HORVU.MOR | high | protein_codir | *** | post-illuminati | NA | NA | NA |
| 235 | HORVU.MOR | HORVU.MOR | 5.73E-26 | 5.75124929 | 0.25 | 0.066 | 7.15E-22 | 16 | unidentified | HORVU.MOR | high | protein_codir | *** | Yellow stripe 1 | GO:0005886, | PF03169 | IPR004813,IPR004813 |
| 32 | HORVU.MOR | HORVU.MOR | 1.07E-40 | 5.614571175 | 0.254 | 0.033 | 1.33E-36 | 17 | Cortex | HORVU.MOR | low | protein_codir | *-* | Protein FAR1-F | GO:0003676, | PF10551,PF0 | IPR018289,IPR004330 |
| 35 | HORVU.MOR | HORVU.MOR | 5.18E-37 | 5.709708617 | 0.261 | 0.04 | 6.47E-33 | 17 | Cortex | HORVU.MOR | low | protein_codir | --* | PDZ and LIM do | GO:0005515, | NA | NA |
| 56 | HORVU.MOR | HORVU.MOR | 8.68E-39 | 5.315124157 | 0.254 | 0.033 | 1.08E-34 | 17 | Cortex | HORVU.MOR | high | protein_codir | *** | Werner Syndro | GO:0003676, | PF01612 | IPR012337,IPR002562 |
| 65 | HORVU.MOR | HORVU.MOR | 6.68E-22 | 5.753200887 | 0.275 | 0.066 | 8.34E-18 | 17 | Cortex | HORVU.MOR | low | protein_codir | *-* | DNA-directed I | GO:0003677, | PF01191 | IPR000783,IPR000783,IPR005571 |
| 69 | HORVU.MOR | HORVU.MOR | 1.59E-18 | 5.752396855 | 0.261 | 0.072 | 1.98E-14 | 17 | Cortex | HORVU.MOR | high | protein_codir | --* | Phosphoinositi | GO:0016020, | NA | NA |
| 75 | HORVU.MOR | HORVU.MOR | 1.06E-17 | 5.335467662 | 0.261 | 0.072 | 1.32E-13 | 17 | Cortex | HORVU.MOR | low | transposon_r | *-* | Transposon Ty3 | GO:0003676, | NA | NA |
| 131 | HORVU.MOR | HORVU.MOR | 0.00980604 | 6.403771267 | 0.283 | 0.192 | 1 | 17 | Cortex | HORVU.MOR | high | protein_codir | *** | Ankyrin repeat- | NA | PF00023,PF1 | IPR002110,IPR020683,IPR020683,IPR020683 |
| 198 | HORVU.MOR | HORVU.MOR | 8.59E-11 | 5.532920359 | 0.254 | 0.084 | 1.07E-06 | 17 | Cortex | HORVU.MOR | low | protein_codir | --* | P-loop contain | NA | NA | NA |
| 213 | HORVU.MOR | HORVU.MOR | 1.10E-20 | 5.411125723 | 0.261 | 0.067 | 1.37E-16 | 17 | Cortex | HORVU.MOR | high | protein_codir | *** | Germin-like pro | GO:0005576, | PF00190 | IPR011051,IPR006045 |
| 219 | HORVU.MOR | HORVU.MOR | 2.22E-18 | 5.562582961 | 0.261 | 0.072 | 2.77E-14 | 17 | Cortex | HORVU.MOR | high | protein_codir | --* | Methyltransfer | GO:0008168, | PF07109 | IPR010940,IPR029063,IPR010251 |
| 2 | HORVU.MOR | HORVU.MOR | 2.50E-90 | 6.819996261 | 0.346 | 0.115 | 3.12E-86 | 18 | Cell cycle | HORVU.MOR | high | protein_codir | *** | Phosphoinositi | GO:0016020, | PF02383 | IPR002013 |
| 62 | HORVU.MOR | HORVU.MOR | 5.31E-162 | 7.611706527 | 0.638 | 0.074 | 6.63E-158 | 18 | Cell cycle | HORVU.MOR | high | protein_codir | *** | Peroxidase | NA | PF00141 | IPR010255,IPR002016 |
| 109 | HORVU.MOR | HORVU.MOR | 2.40E-94 | 6.712378715 | 1 | 0.278 | 3.00E-90 | 18 | Cell cycle | HORVU.MOR | high | protein_codir | *** | Pectin lyase-lik | GO:0004650, | PF00295 | IPR011050,IPR000743 |
| 111 | HORVU.MOR | HORVU.MOR | 3.31E-271 | 6.601504553 | 0.819 | 0.042 | 4.14E-267 | 18 | Cell cycle | HORVU.MOR | low | protein_codir | --* | Retinal guanyly | GO:0000166, | NA | NA |

| | | | | | | | | | | | | | | | | | |
|---|---|---|---|---|---|---|---|---|---|---|---|---|---|---|---|---|---|
| 164 | HORVU.MOR | HORVU.MOR | 2.98E-200 | 7.847052775 | 0.827 | 0.068 | 3.72E-196 | 18 | Cell cycle | HORVU.MOR | low | protein_codir | --* | T-lymphoma in | GO:0005085, | NA | NA |
| 173 | HORVU.MOR | HORVU.MOR | 5.34E-153 | 6.593766119 | 0.291 | 0.028 | 6.67E-149 | 18 | Cell cycle | HORVU.MOR | high | protein_codir | *** | Disease resista | GO:0043531 | PF00931 | IPR027417,IPR002182 |
| 183 | HORVU.MOR | HORVU.MOR | 1.70E-88 | 6.901511429 | 0.827 | 0.102 | 2.13E-84 | 18 | Cell cycle | HORVU.MOR | high | protein_codir | *** | Purine permea | GO:0005215, | NA | NA |
| 184 | HORVU.MOR | HORVU.MOR | 1.02E-184 | 6.505971189 | 0.787 | 0.055 | 1.27E-180 | 18 | Cell cycle | HORVU.MOR | low | protein_codir | *-* | RING finger pro | NA | NA | NA |
| 205 | HORVU.MOR | HORVU.MOR | 8.37E-245 | 6.848578486 | 0.323 | 0.017 | 1.04E-240 | 18 | Cell cycle | HORVU.MOR | high | protein_codir | *** | Kinase family p | GO:0004672, | PF00069 | IPR000719,IPR011009 |
| 216 | HORVU.MOR | HORVU.MOR | 4.43E-76 | 6.94023377 | 0.63 | 0.083 | 5.53E-72 | 18 | Cell cycle | HORVU.MOR | high | protein_codir | *** | 3-ketoacyl-CoA | GO:0003824, | PF08392,PF0 | IPR013601,IPR016039,IPR016039,IPR013747 |
| 13 | HORVU.MOR | HORVU.MOR | 8.87E-208 | 10.41869102 | 0.573 | 0.011 | 1.11E-203 | 19 | Phloem | HORVU.MOR | high | protein_codir | --* | Phytochrome E | GO:0006355, | NA | NA |
| 96 | HORVU.MOR | HORVU.MOR | 9.41E-92 | 10.35873961 | 0.991 | 0.28 | 1.18E-87 | 19 | Phloem | HORVU.MOR | high | protein_codir | *-* | Heavy metal tra | NA | IPR006121 | |
| 139 | HORVU.MOR | HORVU.MOR | 1.21E-103 | 10.22987571 | 0.973 | 0.226 | 1.51E-99 | 19 | Phloem | HORVU.MOR | high | protein_codir | --* | Myosin IC heav | GO:0000166, | NA | NA |
| 148 | HORVU.MOR | HORVU.MOR | 3.02E-94 | 10.45263049 | 0.773 | 0.157 | 3.77E-90 | 19 | Phloem | HORVU.MOR | high | protein_codir | *** | Annexin | NA | PF00191,PF0 | IPR018502,IPR018502,IPR018502,IPR018502 |
| 154 | HORVU.MOR | HORVU.MOR | 4.48E-76 | 10.65935599 | 0.864 | 0.273 | 5.59E-72 | 19 | Phloem | HORVU.MOR | high | protein_codir | --* | Receptor-type | GO:0000166, | NA | NA |
| 155 | HORVU.MOR | HORVU.MOR | 3.63E-119 | 10.12455545 | 0.827 | 0.151 | 4.54E-115 | 19 | Phloem | HORVU.MOR | high | protein_codir | *** | Metallothionei | GO:0046872 | PF01439 | IPR000347 |
| 181 | HORVU.MOR | HORVU.MOR | 3.95E-279 | 10.31879364 | 0.636 | 0.024 | 4.93E-275 | 19 | Phloem | HORVU.MOR | high | protein_codir | *-* | ATP-dependen | GO:0000166, | PF14363,PF0 | IPR025753,IPR003959,IPR027417,IPR027417 |
| 193 | HORVU.MOR | HORVU.MOR | 2.35E-279 | 10.307063 | 0.645 | 0.023 | 2.93E-275 | 19 | Phloem | HORVU.MOR | high | protein_codir | *** | Protein NRT1/ I | GO:0016020, | PF00854 | IPR020846,IPR020846,IPR000109 |
| 196 | HORVU.MOR | HORVU.MOR | 9.31E-147 | 10.25106631 | 0.6 | 0.048 | 1.16E-142 | 19 | Phloem | HORVU.MOR | low | non_coding | --* | 30S ribosomal | GO:0003735, | NA | NA |
| 238 | HORVU.MOR | HORVU.MOR | 5.85E-275 | 10.37295392 | 0.964 | 0.06 | 7.30E-271 | 19 | Phloem | HORVU.MOR | high | protein_codir | *** | Myb family trar | GO:0003677, | PF14379,PF0 | IPR009057,IPR025756,IPR001005,IPR006447 |
| 12 | HORVU.MOR | HORVU.MOR | 1.33E-72 | 6.265711772 | 0.71 | 0.125 | 1.67E-68 | 20 | Endodermis | HORVU.MOR | low | protein_codir | --* | NADH-quinone | GO:0005886, | NA | NA |
| 26 | HORVU.MOR | HORVU.MOR | 3.53E-71 | 6.284921727 | 0.699 | 0.118 | 4.41E-67 | 20 | Endodermis | HORVU.MOR | low | transposon_r | *-* | Transposon pro | NA | NA | NA |
| 34 | HORVU.MOR | HORVU.MOR | 6.39E-75 | 6.259889917 | 0.72 | 0.13 | 7.97E-71 | 20 | Endodermis | HORVU.MOR | low | protein_codir | *-* | Subtilisin-like p | GO:0004252, | NA | NA |
| 40 | HORVU.MOR | HORVU.MOR | 9.56E-42 | 6.047268319 | 0.43 | 0.106 | 1.19E-37 | 20 | Endodermis | HORVU.MOR | low | protein_codir | --* | NADH dehydro | NA | NA | NA |
| 86 | HORVU.MOR | HORVU.MOR | 1.10E-52 | 6.109942087 | 0.441 | 0.094 | 1.37E-48 | 20 | Endodermis | HORVU.MOR | high | protein_codir | --* | Coiled-coil dor | NA | NA | NA |
| 124 | HORVU.MOR | HORVU.MOR | 7.78E-73 | 6.587481659 | 0.742 | 0.128 | 9.71E-69 | 20 | Endodermis | HORVU.MOR | low | protein_codir | --* | Dual-specificit | GO:0003824, | NA | NA |
| 136 | HORVU.MOR | HORVU.MOR | 5.32E-07 | 6.774741842 | 0.355 | 0.124 | 0.006646 | 20 | Endodermis | HORVU.MOR | high | protein_codir | *** | Apoptosis-indu | GO:0016491, | PF07992 | IPR023753,IPR023753 |
| 162 | HORVU.MOR | HORVU.MOR | 0.0001088 | 6.236073591 | 0.28 | 0.145 | 1 | 20 | Endodermis | HORVU.MOR | high | protein_codir | *** | Scarecrow trar | GO:0003700, | PF03514 | IPR005202 |
| 178 | HORVU.MOR | HORVU.MOR | 1.91E-78 | 7.023870697 | 0.312 | 0.007 | 2.38E-74 | 20 | Endodermis | HORVU.MOR | low | protein_codir | --* | kinase with ade | NA | NA | NA |
| 202 | HORVU.MOR | HORVU.MOR | 1.20E-35 | 6.214242284 | 0.398 | 0.024 | 1.50E-31 | 20 | Endodermis | HORVU.MOR | low | protein_codir | --* | Eukaryotic tran | GO:0001732, | NA | NA |
| 7 | HORVU.MOR | HORVU.MOR | 6.93E-48 | 7.841304062 | 0.702 | 0.1 | 8.65E-44 | 21 | CSC | HORVU.MOR | high | protein_codir | *** | Bidirectional su | GO:0005886, | PF03083,PF0 | IPR004316,IPR004316 |
| 77 | HORVU.MOR | HORVU.MOR | 2.68E-36 | 7.646737764 | 0.447 | 0.031 | 3.35E-32 | 21 | CSC | HORVU.MOR | high | protein_codir | --* | glutathione S-t | NA | NA | NA |
| 93 | HORVU.MOR | HORVU.MOR | 2.10E-39 | 7.877522495 | 0.66 | 0.044 | 2.63E-35 | 21 | CSC | HORVU.MOR | high | protein_codir | *** | GDSL esterase | GO:0016788 | PF00657 | IPR013830,IPR013830,IPR001087 |
| 163 | HORVU.MOR | HORVU.MOR | 7.97E-77 | 7.540664622 | 0.426 | 0.026 | 9.95E-73 | 21 | CSC | HORVU.MOR | low | protein_codir | --* | Ribonuclease F | NA | NA | NA |
| 169 | HORVU.MOR | HORVU.MOR | 2.31E-36 | 7.685559887 | 0.979 | 0.359 | 2.89E-32 | 21 | CSC | HORVU.MOR | high | protein_codir | *** | Cysteine prote | GO:0006508, | PF00112,PF0 | IPR000668,IPR013201 |
| 194 | HORVU.MOR | HORVU.MOR | 4.55E-50 | 7.553735469 | 0.787 | 0.127 | 5.68E-46 | 21 | CSC | HORVU.MOR | high | protein_codir | *** | Protein DETOX | NA | PF01554,PF0 | IPR002528,IPR002528,IPR002528 |
| 201 | HORVU.MOR | HORVU.MOR | 2.82E-07 | 8.444904309 | 0.277 | 0.05 | 0.00351903 | 21 | CSC | HORVU.MOR | low | protein_codir | --* | Rad23 UV excis | NA | NA | NA |
| 211 | HORVU.MOR | HORVU.MOR | 1.10E-25 | 7.510271004 | 0.511 | 0.052 | 1.37E-21 | 21 | CSC | HORVU.MOR | low | plastid_relate | --* | Protein PAM68 | GO:0016020, | PF11947 | IPR021855 |
| 214 | HORVU.MOR | HORVU.MOR | 8.37E-34 | 8.483297046 | 0.957 | 0.396 | 1.04E-29 | 21 | CSC | HORVU.MOR | high | protein_codir | *** | Bidirectional su | GO:0005886, | PF03083,PF0 | IPR004316,IPR004316 |
| 220 | HORVU.MOR | HORVU.MOR | 7.44E-49 | 7.662334327 | 0.426 | 0.078 | 9.29E-45 | 21 | CSC | HORVU.MOR | high | protein_codir | *-* | Calcium unipo | GO:0016020, | PF04678,PF0 | IPR006769,IPR006769 |
| 17 | HORVU.MOR | HORVU.MOR | 5.10E-153 | 8.307243205 | 0.698 | 0.024 | 6.37E-149 | 22 | unidentified | HORVU.MOR | high | protein_codir | *** | Kinase-like | GO:0004672, | PF00069 | IPR011009,IPR000719 |
| 45 | HORVU.MOR | HORVU.MOR | 2.08E-07 | 8.498822054 | 0.302 | 0.074 | 0.00259925 | 22 | unidentified | HORVU.MOR | high | protein_codir | *** | Benzyl alcohol | GO:0016740, | PF02458 | IPR003480 |
| 51 | HORVU.MOR | HORVU.MOR | 2.58E-100 | 8.116291617 | 0.558 | 0.028 | 3.22E-96 | 22 | unidentified | HORVU.MOR | high | protein_codir | *** | WRKY transcrip | GO:0003677, | PF03106 | IPR003657,IPR003657 |
| 57 | HORVU.MOR | HORVU.MOR | 3.68E-143 | 8.277263309 | 0.628 | 0.026 | 4.59E-139 | 22 | unidentified | HORVU.MOR | low | non_coding | *-* | Cytochrome P4 | GO:0004497, | NA | NA |
| 114 | HORVU.MOR | HORVU.MOR | 5.66E-31 | 8.098610531 | 0.721 | 0.149 | 7.07E-27 | 22 | unidentified | HORVU.MOR | high | protein_codir | *** | BTB/POZ doma | NA | PF00651 | IPR000210,IPR011333,IPR008974 |
| 127 | HORVU.MOR | HORVU.MOR | 3.49E-152 | 8.159048915 | 0.581 | 0 | 4.35E-148 | 22 | unidentified | HORVU.MOR | low | protein_codir | --* | Outer arm dyne | NA | NA | NA |
| 128 | HORVU.MOR | HORVU.MOR | 3.49E-152 | 8.159048915 | 0.581 | 0 | 4.35E-148 | 22 | unidentified | HORVU.MOR | low | protein_codir | --* | Outer arm dyne | NA | NA | NA |
| 138 | HORVU.MOR | HORVU.MOR | 1.32E-14 | 8.71481135 | 0.581 | 0.109 | 1.65E-10 | 22 | unidentified | HORVU.MOR | low | protein_codir | *-* | Retrovirus-rela | NA | NA | NA |
| 187 | HORVU.MOR | HORVU.MOR | 1.05E-79 | 8.193322329 | 0.744 | 0.058 | 1.31E-75 | 22 | unidentified | HORVU.MOR | high | protein_codir | *** | Receptor-like k | GO:0004672, | PF00069 | IPR011009,IPR000719 |
| 232 | HORVU.MOR | HORVU.MOR | 0 | 8.593749727 | 0.698 | 0.008 | 0 | 22 | unidentified | HORVU.MOR | low | transposon_r | *-* | Transposon pro | GO:0003677, | PF02992,PF1 | IPR004242,IPR029480 |
| 4 | HORVU.MOR | HORVU.MOR | 8.74E-27 | 11.27777616 | 1 | 0.325 | 1.09E-22 | 23 | Xylem | HORVU.MOR | high | protein_codir | *** | Cysteine prote | GO:0006508, | PF08246,PF0 | IPR013201,IPR000668 |
| 59 | HORVU.MOR | HORVU.MOR | 6.08E-143 | 12.14284146 | 0.594 | 0.01 | 7.59E-139 | 23 | Xylem | HORVU.MOR | high | protein_codir | *** | Dirigent protei | GO:0005576, | PF03018 | IPR004265 |
| 66 | HORVU.MOR | HORVU.MOR | 1.39E-26 | 10.92969385 | 1 | 0.346 | 1.73E-22 | 23 | Xylem | HORVU.MOR | high | protein_codir | *** | Dirigent protei | GO:0005576, | PF03018 | IPR004265 |
| 103 | HORVU.MOR | HORVU.MOR | 2.48E-47 | 11.47926484 | 0.906 | 0.134 | 3.09E-43 | 23 | Xylem | HORVU.MOR | high | protein_codir | *** | NAC domain-c | GO:0003677, | PF02365 | IPR003441,IPR003441 |
| 104 | HORVU.MOR | HORVU.MOR | 8.70E-99 | 11.00355794 | 0.906 | 0.052 | 1.09E-94 | 23 | Xylem | HORVU.MOR | high | protein_codir | *** | NAC domain-c | GO:0003677, | PF02365 | IPR003441,IPR003441 |
| 144 | HORVU.MOR | HORVU.MOR | 9.46E-29 | 11.86517629 | 0.875 | 0.212 | 1.18E-24 | 23 | Xylem | HORVU.MOR | high | protein_codir | *-* | ROP-interactiv | NA | PF00786 | IPR000095 |
| 159 | HORVU.MOR | HORVU.MOR | 1.27E-109 | 12.05720308 | 0.812 | 0.039 | 1.59E-105 | 23 | Xylem | HORVU.MOR | high | protein_codir | *** | WEB family pro | NA | NA | NA |
| 188 | HORVU.MOR | HORVU.MOR | 7.26E-40 | 13.17268643 | 0.844 | 0.136 | 9.06E-36 | 23 | Xylem | HORVU.MOR | high | protein_codir | *-* | Endo-1,4-beta- | GO:0004553, | PF02018,PF0 | IPR008979,IPR003305,IPR017853,IPR001000 |
| 195 | HORVU.MOR | HORVU.MOR | 3.92E-52 | 11.16517293 | 0.844 | 0.098 | 4.89E-48 | 23 | Xylem | HORVU.MOR | high | protein_codir | *-* | transcription fa | NA | PF04784,PF1 | IPR006869,IPR025757 |
| 236 | HORVU.MOR | HORVU.MOR | 1.65E-25 | 10.86075504 | 0.875 | 0.12 | 2.05E-21 | 23 | Xylem | HORVU.MOR | high | protein_codir | *** | transmembran | NA | PF05078 | IPR007770 |

001002

001002
