## Supplementary Table 3 for "A single-cell transcriptome atlas of the barley root apical meristem uncovers conserved and divergent roles of HvWOX5"

**Supplementary Table 3. Agreements after cross-species comparisons.**

| barley cluster | number of agreeing predictions | agreement | consensus cell type | arabidopsis annotation | tomato annotation | wheat annotation | maize annotation | rice annotation |
|---|---|---|---|---|---|---|---|---|
| 0 | 2 | No | Epidermis | Lateral root cap | ATRICHOBLAST | 1 - Epidermis | 2 - Cortex, 13 - Cortex | Stele |
| 1 | 5 | Very strong | Cortex | Cortex | CORTEX | 2 - Cortex | 2 - Cortex | Cortex |
| 2 | 4 | Strong | Endodermis | Endodermis | ENDODERMIS | 9 - Endodermis | 9 - Stele, 11 - Pith, 5 - Pith | Endodermis |
| 3 | 3 | Weak | Cortex | Endodermis | CORTEX, EXODERMIS | 2 - Cortex | 2 - Cortex | Stele |
| 4 | 4 | Strong | Cortex | Cortex | PERICYLCE, CORTEX | 2 - Cortex | 1 - Cortex | Epidermis (near root hair) |
| 5 | 4 | Strong | Stele | Columella, Initials, Phloem | PHLOEM & PROCAMB | 11 - Endodermis/ Phloem, 4 - Provascular Cells | 16 - Endodermis, 12 - Endodermal initials | Metaxylem |
| 6 | 4 | Strong | Epidermis | Atrichoblast | ATRICHOBLAST | 1 - Epidermis | 7 - Epidermis/LRC | Root cap |
| 7 | 5 | Very strong | Cortex | Cortex | CORTEX | 2 - Cortex, 0 - Unknown | 1 - Cortex, 19 - Cortex | Cortex |
| 8 | 2 | No | Epidermis / stele | Lateral root cap | ATRICHOBLAST | 1 - Epidermis | 3 - Stele | Stele |
| 9 | 5 | Very strong | Cortex | Cortex | CORTEX | 2 - Cortex | 2 - Cortex | Cortex |
| 10 | 5 | Very strong | Stele | QC, Pericycle, Procambium | PHLOEM & PROCAMB, RC & COL & QC | 11 - Endodermis/ Phloem | 10 - Phloem, 0 - Stele | Metaxylem |
| 11 | 4 | Strong | Initials | Initials | MERIST ZONE | 3 - G1/S, 12 - G2/M, 7 - Epidermis | 20 - Initials | Endodermis |
| 12 | 5 | Very strong | Cortex | Cortex | CORTEX | 2 - Cortex | 1 - Cortex | Cortex |
| 13 | 5 | Very strong | Cortex | Cortex | CORTEX | 2 - Cortex | 2 - Cortex, 8 - Cortex, 1 - Cortex | Cortex |
| 14 | 5 | Very strong | Epidermis | Atrichoblast | TRICHOBLAST | 1 - Epidermis | 17 - Cortex initials, 7 - Epidermis/LRC | Epidermis (near root hair), Root hair |
| 15 | 3 | Weak | Epidermis | Atrichoblast | ATRICHOBLAST | 1 - Epidermis | 17 - Cortex initials | Root cap |
| 16 | 3 | Weak | Epidermis / LRC | Trichoblast, Lateral root cap | ATRICHOBLAST | 1 - Epidermis, 14 - Root Cap | 3 - Stele | Stele |
| 17 | 2 | No | Initials / cortex | Cortex | MERIST ZONE | 3 - G1/S | 14 - Cortex | Root cap |
| 18 | 4 | Strong | Epidermis | Atrichoblast | TRICHOBLAST | 8 - Root Hair | 7 - Epidermis/LRC | Endodermis |
| 19 | 3 | Weak | Phloem | Phloem | PHLOEM & PROCAMB | 6 - Pericycle, 13 - Phloem | 6 - Endodermis | Metaxylem |
| 20 | 3 | Weak | Cortex | Endodermis | CORTEX | 2 - Cortex | 8 - Cortex | Epidermis |
| 21 | 3 | Weak | Phloem / endodermis | Phloem, Epidermis | PHLOEM & PROCAMB | 11 - Endodermis/ Phloem | 4 - Pericycle, 16 - Endodermis | Endodermis |
| 22 | 2 | No | Stele | Lateral root cap | ATRICHOBLAST | 5 - Unknown | 3 - Stele | Stele |
| 23 | 5 | Very strong | Xylem | Xylem | XYLEM | 10 - Xylem | 15 - Xylem | Metaxylem, Epidermis |

| Source | DOI |
|---|---|
| Arabidopsis | https://doi.org/10.1126/science.aay4970 |
| Tomato | https://doi.org/10.1038/s41477-023-01567-x |
| Maize | https://doi.org/10.1126/science.abj2327 |
| Rice | https://doi.org/10.1016/j.molp.2020.12.014 |
| Wheat | https://doi.org/10.1016/j.celrep.2025.115240 |
