## Supplementary Table 4 for "A single-cell transcriptome atlas of the barley root apical meristem uncovers conserved and divergent roles of HvWOX5"

**Supplementary Table 4. Root length measurements of GPF and *hvwox5* -mutant roots [mm].**

| GPF | wox5 |
|---|---|
| 122,349 | 102,202 |
| 117,516 | 113,12 |
| 132,489 | 64,312 |
| 125,555 | 88,723 |
| 130,818 | 119,164 |
| 118,496 | 109,296 |
| 110,351 | 84,132 |
| 122,607 | 92,93 |
| 108,493 | 84,089 |
| 129,363 | 78,063 |
| 131,548 | 81,963 |
| 129,763 | 56,026 |
| 130,654 | 97,436 |
| 95,336 | 95,537 |
| 126,479 | 38,033 |
| 94,079 | 116,373 |
| 95,967 | 69,175 |
| 84,124 | 62,696 |
| 118,149 | 59,966 |
| 93,381 | 102,392 |
| 119,284 | 106,063 |
| 125,943 | 84,169 |
| 93,281 | 98,743 |
| 108,781 | 70,688 |
| 112,425 | 74,401 |
| 123,988 | 102,565 |
| 111,178 | 96,5 |
| 108,434 | 64,344 |
| 119,931 | 121,902 |
| 127,095 | 93,146 |
| 102,178 | 51,013 |
| 74,233 | 77,852 |
| 105,249 | 82,466 |
| 107,109 | 69,081 |
| 127,527 | 86,487 |
| 111,924 | 73,26 |
| 112,145 | 88,479 |
| 151,742 | 55,309 |
| 118,273 | 69,501 |
| | 99,211 |
| | 110,895 |
| | 69,305 |
| | 43,874 |
| | 50,187 |
| | 111,304 |
| | 102,016 |
| | 58,735 |
| | 85,879 |
| | 87,418 |
| | 41,366 |
| | 71,536 |
