## Supplementary Table 5 for "A single-cell transcriptome atlas of the barley root apical meristem uncovers conserved and divergent roles of HvWOX5"

**Supplementary Table 5. Meristematic zone length measurements of GPF and *hvwox5* -mutant roots [µm].**

| GPF | wox5 |
|---|---|
| 861,949 | 631,571 |
| 710,942 | 699,348 |
| 714,256 | 730,929 |
| 891,556 | 816,653 |
| 595,926 | 565,238 |
| 876,108 | 543,816 |
| 832,716 | 409,715 |
| 892,102 | 330,988 |
| | 429,294 |
| | 700,532 |
| | 429,76 |
| | 600,758 |
