## Supplementary Table 6 for "A single-cell transcriptome atlas of the barley root apical meristem uncovers conserved and divergent roles of HvWOX5"

**Supplementary Table 6. Scoring of metaxylem phenotypes in GPF and *hvwox5* -mutant roots.**

| Genotype | Root nr. | Normal | Division | Double |
|---|---|---|---|---|
| GPF | 1 | 1 | | |
| GPF | 2 | | 1 | |
| GPF | 3 | 1 | | |
| GPF | 4 | 1 | | |
| GPF | 5 | | 1 | |
| GPF | 6 | 1 | | |
| GPF | 7 | 1 | | |
| GPF | 8 | | 1 | |
| GPF | 9 | 1 | | |
| GPF | 10 | | 1 | |
| GPF | 11 | | 1 | |
| GPF | 12 | 1 | | |
| GPF | 13 | | 1 | |
| GPF | 14 | 1 | | |
| GPF | 15 | 1 | | |
| GPF | 16 | | 1 | |
| wox5 | 1 | | 1 | |
| wox5 | 2 | | 1 | |
| wox5 | 3 | | | 1 |
| wox5 | 4 | 1 | | |
| wox5 | 5 | | | 1 |
| wox5 | 6 | | | 1 |
| wox5 | 7 | | 1 | |
| wox5 | 8 | | | 1 |
| wox5 | 9 | | 1 | |
| wox5 | 10 | | | 1 |
| wox5 | 11 | 1 | | |
| wox5 | 12 | | | 1 |
| wox5 | 13 | | 1 | |
| wox5 | 14 | | 1 | |
| wox5 | 15 | | | 1 |
| wox5 | 16 | | | 1 |
| wox5 | 17 | | | 1 |
| wox5 | 18 | | 1 | |
