## Supplementary Table 7 for "A single-cell transcriptome atlas of the barley root apical meristem uncovers conserved and divergent roles of HvWOX5"

**Supplementary Table 7. EdU/mPSPI staining of GPF and *hvwox5* -mutant roots.** Number of QC divisions and CSC [illegible]

| Genotype | Root # | QC divisions | CSC layers |
|---|---|---|---|
| GPF | 1 | 0 | 2 |
| GPF | 2 | 1 | 2 |
| GPF | 3 | 0 | 2 |
| GPF | 4 | 3 | 1 |
| GPF | 5 | 0 | 2 |
| GPF | 6 | 2 | 2 |
| GPF | 7 | 0 | 3 |
| GPF | 8 | 1 | 2 |
| GPF | 9 | 0 | 2 |
| wox5 | 1 | 0 | 2 |
| wox5 | 2 | 1 | 1 |
| wox5 | 3 | 2 | 2 |
| wox5 | 4 | 0 | 3 |
| wox5 | 5 | 4 | 3 |
| wox5 | 6 | 1 | 3 |
| wox5 | 7 | 3 | 1 |
| wox5 | 8 | 3 | 1 |
| wox5 | 9 | 3 | 0 |
| wox5 | 10 | 3 | 1 |
| wox5 | 11 | 2 | 3 |
| wox5 | 12 | 1 | 2 |
| wox5 | 13 | 1 | 1 |
| wox5 | 14 | 1 | 2 |
| wox5 | 15 | 2 | 1 |
| wox5 | 16 | 3 | 1 |
| wox5 | 17 | 2 | 1 |
| wox5 | 18 | 1 | 2 |
| wox5 | 19 | 1 | 1 |

layers as indicated.
